# Czech Republic butterfly barcoding reveals that distribution of genetic lineages depends on species traits

**DOI:** 10.1101/2024.01.17.576072

**Authors:** Alena Sucháčková Bartoňová, Patrik Škopek, Martin Konvička, Jiří Beneš, Lukáš Spitzer, Claudio Sbaraglia, Vladimír Vrabec, Jana Papp Marešová, Hana Konvičková, Zdeněk Faltýnek Fric

## Abstract

**Aim:** The distribution of within-species lineages has been affected by Quaternary climate changes, and population differentiation has been influenced by species life histories. We investigated whether the distribution of individual mitochondrial genetic lineages reflects the constituent species’ traits. Using the functionally diverse group of butterflies, we examined which lineages are present in Central Europe, an important suture zone.

**Location:** Czech Republic and Western Palearctic.

**Taxon:** A total of 140 butterfly species.

**Methods:** We sequenced DNA barcodes (cytochrome c oxidase 1) (959 sequences) of the entire Czech Republic butterfly fauna and used BOLD data to visualize the species’ biogeographic patterns across Europe. We categorised the distribution patterns of lineages inhabiting the Czech Republic, and used multivariate statistics to interpret these categories by the butterflies’ habitats, life histories, and threat levels.

**Results:** Open habitat dwellers with specialist traits belonged to Eastern, Southern, and temperate lineages. Habitat generalists and woodland dwellers belonged to the Western lineage, formed several lineages, or displayed low genetic diversity; they often developed on woody plants, were large-winged, and had long flight periods. The most threatened species were the specialists of Southern, Eastern, and temperate lineages.

**Main conclusions:** The distribution of lineages in Central Europe reflects the history of Quaternary ecosystems: during cold periods of the Pleistocene, the diverse open habitats prevailed, and species could expand westwards. Such species also suffer the most under the current anthropogenic habitat alteration. On the other hand, the mobile generalists and woodland dwellers expanded to Central Europe during the Holocene. Our approach of linking the distribution of lineages with species traits can be transferred to other study systems, and we show that DNA barcoding of under-sampled areas represents a powerful tool for discovering the driving forces of biogeography.

## INTRODUCTION

Population genetic differentiation is affected by species’ range sizes, life histories, and varies across geographic regions (Dapporto et al., 2019; Gamba & Muchhala, 2020; Miller et al., 2021). In temperate regions, the distribution of different genetic lineages, i.e., the groups of sequences with shared history, has been massively influenced by Quaternary climate change (Hewitt, 2000; 2004; Schmitt, 2007; Hofreiter & Stewart, 2009). Thus, a population inhabiting an area might be the result of combining the landscape history with trait-mediated divergence.

Under the Quaternary’s dynamic conditions, Europe’s unique geographic idiosyncrasy affected the distribution of genetic lineages. In Europe, there are three peninsulas of a limited area with a Mediterranean climate functioning as speciation centres, east-west oriented mountain ridges acting as barriers, and a gradient between oceanic climate in the west and continental climate in the east. These factors influenced species’ ranges: during the shorter warm stages of the Quaternary, warm-adapted species expanded from Mediterranean peninsulas or other spatially restricted areas (refugia), while cold-adapted species retreated uphill and northwards (de Lattin, 1967; Hewitt, 1996, 1999; Schmitt, 2007; Schmitt & Varga, 2012). Simultaneously, continental species tended to retreat eastwards (Stewart, Lister, Barnes, & Dalén, 2010). Notably, Central Europe represents a crossroad of different lineages (Janoušek et al., 2012; Pfäffle, Bolfíková, Hulva, & Petney, 2014; Nürnberger, Lohse, Fijarczyk, Szymura, & Blaxter, 2016). This area went through immense changes during the Quaternary, with an exchange of biota between the cold and warm stages (Horáček & Ložek, 1988), while the landscape varied from a cold steppe-tundra to open park-like woodlands (Kahlke, 2014; Sandom, Ejrnæs, Hansen, & Svenning, 2014; Vera, 2000; Pearce et al., 2023).

A feasible method to evaluate genetic differentiation and lineage distribution on large-scale data is DNA barcoding, i.e., sequencing a short standardized gene fragment, which originally aimed for a simple tool for species identification and discovery (Hebert, Cywinska, Ball, & deWaard, 2003). An important step towards this goal was building well-curated world-wide databases covering as many species as possible, such as the Barcoding of Life Data System (BOLD; Ratnasingham & Hebert 2007). The DNA barcode widely used in various animal groups, mitochondrial gene cytochrome c oxidase subunit I (COI), proved effective not only for species identification (Hebert, Ratnasingham, & de Waard, 2003), but also for uncovering cryptic diversity including discovery of new species (Hernández-Roldán et al., 2016), and phylogeographic and population genetic studies (Kühne, Kosuch, Hochkirch, & Schmitt, 2017; Maresova et al., 2021). The existence of large database data allowed testing the DNA barcoding performance across large scales on entire faunal groups, summarizing genetic diversity and uncovering biogeographic patterns (e.g., Meier et al., 2006; Geiger et al., 2014; Hendrich et al., 2015; Weigand et al., 2019; Dincă et al., 2021; D’Ercole et al., 2021; Galimberti et al., 2021; Dapporto et al., 2022).

European butterflies are one of the best DNA barcoded groups of organisms, represented by several national and international DNA barcoding libraries (Dincă, Zakharov, Hebert, & Vila, 2011; Dincă et al., 2015, 2021; Hausmann et al., 2011; Huemer, Mutanen, Sefc, & Hebert, 2014; Huemer et al., 2018; Litman et al., 2018; Dapporto et al., 2022). Geographic patterns of diversity and differentiation are extraordinarily well-explored, especially for south-western Europe (Dapporto et al., 2019, 2022; Dincă et al., 2021).

In this study, we focused on the functionally diverse butterfly fauna of the Czech Republic and provide the DNA barcoding library for the country. We view the country, situated in Central Europe, as a sieve of species that either reached the area during the Holocene, or survived there since the last glaciation. The Czech butterfly fauna consists of both generalists and specialists of diverse habitats, dry and wet, lowland and highland, and of different stages of habitat openness, whose life histories are known in considerable detail (Bartonova, Benes, & Konvicka, 2014; Macek, Laštůvka, Beneš, & Traxler, 2015), and at the same time have one of the highest threat values in Europe (Warren et al., 2021). We DNA barcoded samples of Czech populations of 140 species (∼98% of extant Czech butterfly fauna), accessed BOLD data for these species across their ranges, analysed their mitochondrial phylogeographic structure, and related them to species-specific traits. We predict that lineages do not inhabit the region randomly, but do so with respect to individual species’ habitat affinities and life histories. In particular, we expect differences between generalist and specialist species, and between species associated with closed (woodland) and open (grasslands, steppe, wetlands) habitats. Moreover, the lineage presence can be related to the species’ current threat levels, thus, the genetic backgrounds can predetermine individual species’ future.

## MATERIAL AND METHODS

### Sampling and DNA sequencing

Czech butterfly fauna consists of 141 species including migrants, plus six irregular vagrants, and a further 16 species that are considered extinct (Benes et al., 2002; Wiemers et al., 2018). We collated samples from 140 butterfly species originating from the Czech Republic (CZ) (Figure 1a), from the years 2004–2022 (Supplementary Materials: figshare repository DOI 10.6084/m9.figshare.24559852). The dataset comprises all the extant species except for *Melitaea phoebe* (recent colonizer), *Parnassius apollo* (existing as a conservation-dependent reintroduced population), and *Hyponephele lycaon* (recently extinct in the country), but including *Pieris mannii*, which newly recolonized the country, and a rare vagrant, *Leptotes pirithous*. We also sequenced one specimen of *Lampides boeticus*, imported with fruit from Spain (not included in analyses).

**Figure 1.**
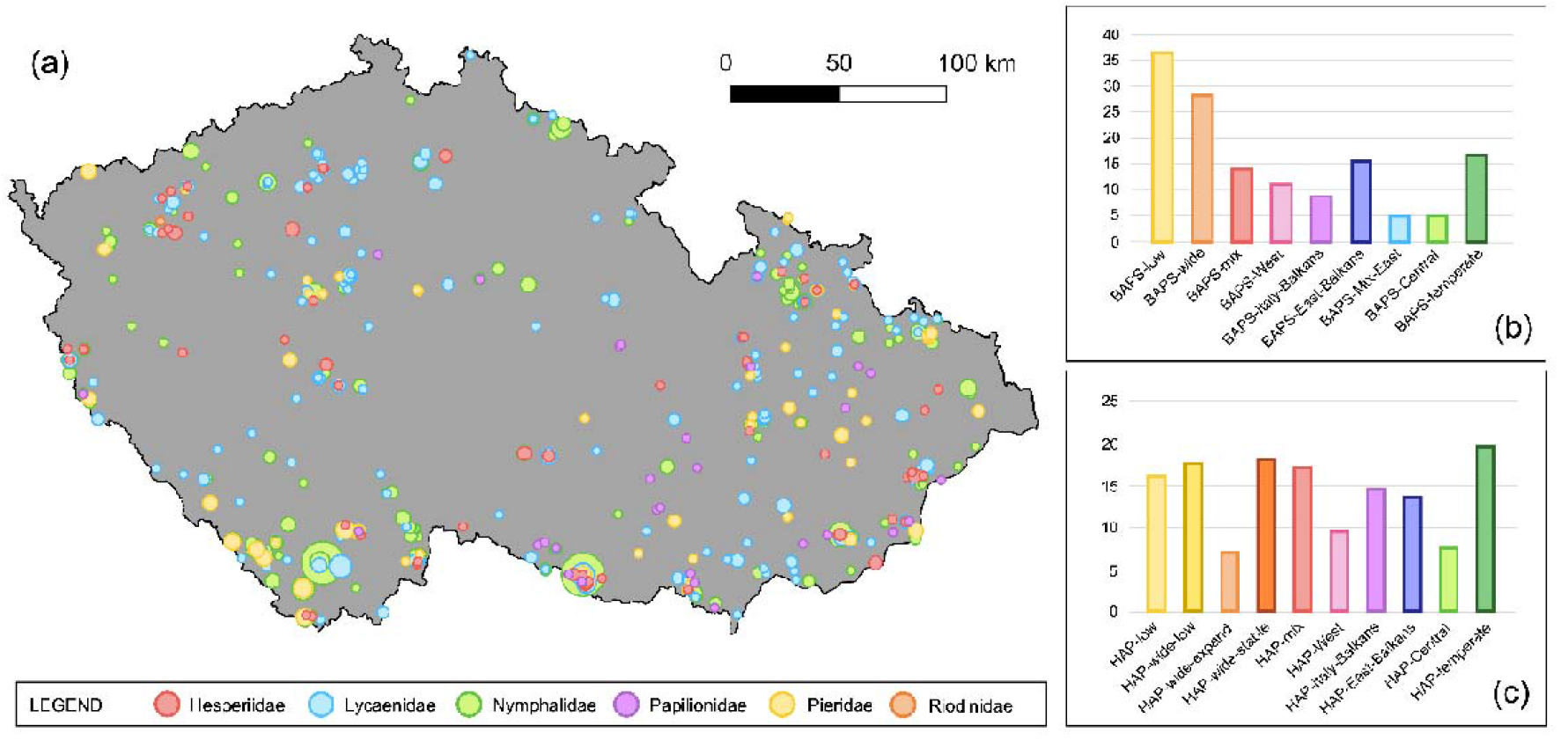
**(a)** Map of the Czech Republic with samples obtained for Czech butterfly DNA barcoding (sequencing of the mitochondrial gene COI). The points’ sizes reflect the number of samples. (b–c) Numbers of species with different genetic patterns, as revealed by **(b)** Bayesian Analysis of Population Structure (BAPS), and **(c)** haplotype networks. The map projection is S-JTSK/Krovak East North.

We extracted DNA from the butterfly legs using Genomic DNA Mini Kit (Tissue) (Geneaid Biotech Ltd.). We amplified the barcode, first part of COI, using primers hybLCO (5′-TAATACGACTCACTATAGGGGGTCAACAAATCATAAAGATATTGG-3′) and hybHCO (5′-ATTAACCCTCACTAAAGGGTAAACTTCAGGGTGACCAAAAAATCA-3′) (Folmer, Black, Hoeh, Lutz, & Vrijenhoek, 1994). In the case of fragmented DNA, we used two pairs of primers: [LCO-K699] + [Ron-HCO] (K699: 5′-ATTAACCCTCACTAAAGGGWGGGGGGTAAACTGTTCATCC-3′; Ron: 5′-TAATACGACTCACTATAGGGGGATCACCTGATATAGCATTCCC-3′) (www.nymphalidae.net; Monteiro & Pierce 2001). The mixture for each sample consisted of 4 µl of PCR H_2_O, 0.625 µl of each primer, 6.25 µl of Bioline 2x MyTaq HS Red Mix (Meridian Bioscience Inc.), and 2 µl of extracted DNA. The polymerase chain reaction protocol consisted of 95°C for 5 min; followed by 40 cycles of 94°C for 30 s, 50°C for 30 s, and 72°C for 90 s; with a final extension of 72°C for 10 min. PCR products were cleaned using FastAP and Exo I enzymes (Thermo Fisher Scientific). Sanger sequencing was performed in Macrogen Inc. (Amsterdam, Netherlands) on 3730xl DNA Analyzer, in one direction from the 5′ end. The sequences were checked, trimmed, and aligned in Geneious v. 8.0.5 (Kearse et al., 2012). We downloaded all COI sequences for each species occurring in the Czech Republic from the BOLD database (Ratnasingham & Hebert, 2007) (for 12 October 2022) which were georeferenced and sequenced from the 5′ end. We aligned the samples together with the newly sequenced data.

We analysed each species dataset separately. Members of species pairs known to share haplotypes or not monophyletic in the DNA barcode (Dincă et al., 2011) (specifically *Colias crocea/erate, Cupido decoloratus/alcetas, Erebia euryale/ligea, Pieris napi*/*bryoniae, Polyommatus bellargus/coridon, Pseudophilotes baton/vicrama*) entered the analyses as separate species because of their unique species traits.

### Phylogeographic patterns

For each species, we performed two phylogeographic analyses: (1) Bayesian Analysis of Population Structure (BAPS) (Cheng, Connor, Sirén, Aanensen, & Corander, 2013), which sorts data into clusters of related sequences (*genetic lineages*), in R package ‘rhierbaps’ (Tonkin-Hill, Lees, Bentley, Frost, & Corander, 2018), on level 1, automatically estimating the number of clusters; and (2) TCS haplotype networks in the program POPART (Leigh & Bryant, 2015), a parsimony method which, additionally, shows putative evolutionary relationships among samples. In both cases, we checked the results visually and cleaned the dataset as to omit database samples which were either shorter than 500 bp, or potentially misidentified (dissimilarity of sequences represented by long branches in haplotype networks was evaluated using the BLAST algorithm, Altschul et al., 1990, in the NCBI database).

We plotted the distribution of BAPS clusters on a map of Europe using the geographic coordinates of the samples. Then, we manually assigned the lineages present in the Czech Republic to one of the following categories, or to two, if two specific lineages were present: (1) *BAPS-low*: low genetic diversity – only a single BAPS cluster in the entire species; (2) *BAPS-wide*: a lineage widely distributed across Europe also present in CZ (but more lineages are present within the species); (3) *BAPS-mix*: two or more widespread European lineages are present in CZ; (4) *BAPS-West*: the CZ samples related to samples from Western Europe; (5) *BAPS-Italy-Balkans*: the CZ samples related to those from Italy or Italy plus the Balkans; (6) *BAPS-East-Balkans*: the CZ samples are related to those from the Balkans or Balkans plus Eastern Europe/Asia; (7) *BAPS-mix-East*: CZ samples consist of more lineages distributed in Italy/Balkans or East/Balkans; (8) *BAPS-Central*: the Central European mountains or Pannonian lowlands produced a separate lineage to which CZ samples belong; and (9) *BAPS-temperate*: the CZ samples are related to lineages inhabiting a similar latitudinal belt, not present in the Mediterranean peninsulas.

The results of haplotype networks were scored based on the geographic area inhabited by haplotypes found in CZ and haplotypes distant from them with maximum of two mutations. When the Czech samples were part of a widespread lineage, the supposed demographic scenario from the net was visually estimated. The categories were: (1) *HAP-low*: lack of differentiation (either low diversity or expansion -star-like haplotype structure) in the entire species. Then, in categories (2)–(4), the Czech samples are part of a widespread European lineage (another, diverged lineage exists in the network): (2) *HAP-wide-low*: CZ samples belong to a widespread European lineage with low haplotype diversity (supposed bottleneck; maximum of 10 closely related haplotypes); (3) *HAP-wide-expand*: part of expanding widespread European lineage (star-like structure, with eight and more haplotypes diverging from the main one); (4) *HAP-wide-complex*: part of a complex widespread lineage (showing either a stable network with several connections and even sequences in haplotypes, or combination of a stable network and expansion. (5) *HAP-mix*: divergent haplotypes were present in CZ with distance >2 mutations. The following categories were defined similarly to BAPS classification: (6) *HAP-West*; (7) *HAP-Italy-Balkans*; (8) *HAP-East-Balkans*; (9) *HAP-Central*; and (10) *HAP-temperate*.

### Species traits

We prepared data on the following three groups of species traits: (1) Habitat affinity, defining nine categories: *ubiquitous* (generalist), *mesophilic 1* (grassland), *mesophilic 2* (shrubland), *mesophilic 3* (woodland), *xerothermophilic 1* (steppe), *xerothermophilic 2* (dry shrubland), *hydrophilic* (wetland), *tyrphophilic* (bog/peatland), and *alpine* (Benes et al., 2002). If two categories applied for a species, these were assigned as “0.5” per category.

1. Life history traits linked to dispersal and landscape scale survival (Bartonova, Benes, Fric, Chobot, & Konvicka, 2016): *density*, the number of individuals which can occur per area of habitat (adapted from area demand) [ranked 1–9, from sparse to dense] (Reinhardt, Sbieschne, Settele, Fischer, & Fiedler, 2007); *feeding index*, the trophic range, defined as an index that weights the number of consumed host plant families [F] and genera [G] in the Czech Republic: (G*F^a^)^1/2^ where a=F/2G (Benes et al., 2002; index modified after Garcia-Barros 2000); *fertility*, the number of eggs per female at eclosion [ranked 1–9] (Reinhardt et al., 2007); *flight period length*, the number of months of adult flight, summed across generations and excluding hibernation (Benes et al., 2002); *forewing length*, approximation for body size [mm] (Higgins & Riley, 1970); *host plant form*, express prevailing host plant apparency [ranked 1–4, small forbs, large forbs/grasses, bushes, trees] (Cizek, Fric, & Konvicka, 2006); *mobility*, the propensity to disperse [ranked 1–9, from sedentary to mobile] (Reinhardt et al., 2007); *overwintering stage* [ranked 1–5, from egg to migrating adult] (Tolman & Lewington, 2008); and *voltinism*, average number of generations per year in the Czech Republic (Benes et al., 2002). In the case of species not covered in Reinhardt et al. (2007) (N=39), missing values of density and fertility were substituted by averages, and missing values of mobility were scored based on authors’ expertise (similarly to Bartonova et al.,2016).
2. Threat status, based on national Red List categories (ordinal scale: 1 – least concern, 2 – near threatened, 3 – vulnerable, 4 – endangered, and 5 – critically endangered) (Hejda, Farkač, & Chobot, 2017).

### Relating phylogeographic patterns to species traits

To investigate the relationships between phylogeographic patterns and traits, we used a multivariate approach, the unimodal canonical correspondence analyses (CCA) in CANOCO v.5.0 (Ter Braak & Šmilauer, 2012), which extracts variation in multivariate dependent variables even if obscured by other larger sources of variation (Ter Braak & Verdonschot, 1995), and tests the significance of the ordinations via the Monte-Carlo permutation tests (999 runs). In these analyses, individual species were the samples, either BAPS or haplotype network categories were responses, and species traits were predictors. The response variables were categorical as they identified the group to which a sample (butterfly species) belong. Since response variables could have been represented by more categories, they entered the analysis in the form of dummy variables with fuzzy coding (0, 0.5 or 1), and CCA in this case represented a generalization of linear discriminant analysis (LDA), optimal for categorical variables, with chi-square distances as generalization of Mahalanobis distances among observations (Ter Braak & Verdonschot, 1995; Ter Braak & Šmilauer, 2012). CANOCO v.5.0 automatically applies standardization to unit variance (bringing their means to zero and variances to one) to control for the different dimensions of the predictors. In the case of habitat affinities and life history traits, two tests were performed: (a) global test, using all variables and (b) a forward selection procedure, picking a combination of traits that significantly fit the phylogeographic patterns. Threat status, representing a single explanatory variable, was also tested using CCA.

## RESULTS

### Dataset

The final dataset consisted of 1,110 sequences of the 140 species found in the Czech Republic (Fig. 1a) (excluding *L. boeticus*); 151 sequences from BOLD and GenBank, and 959 newly generated (mean=8±5.1 SE sequence per species, range 1–83), which can be found in BOLD as project BBCZ. We added 20,695 sequences for these species from BOLD originating in other locations. Together, we used 21,805 sequences (mean=156±86.5 SE sequence per species, range 18–854) (Supplementary Materials). The number of species assigned to each BAPS and haplotype network category are summarized in Fig. 1b and c, and the categories assigned for each species are found in Supporting Information 1 and Supplementary Materials. The BAPS category (Fig. 2) with the highest number of species was *BAPS-low* (N=36), followed by *BAPS-wide* (N=285). The haplotype network category (Fig. 3) represented by highest number of species was *HAP-temperate* (N=19.5), followed by *HAP-wide-complex* (N=18), *HAP-wide-low* (N=17.5), and *HAP-mix* (N=17).

**Figure 2.**
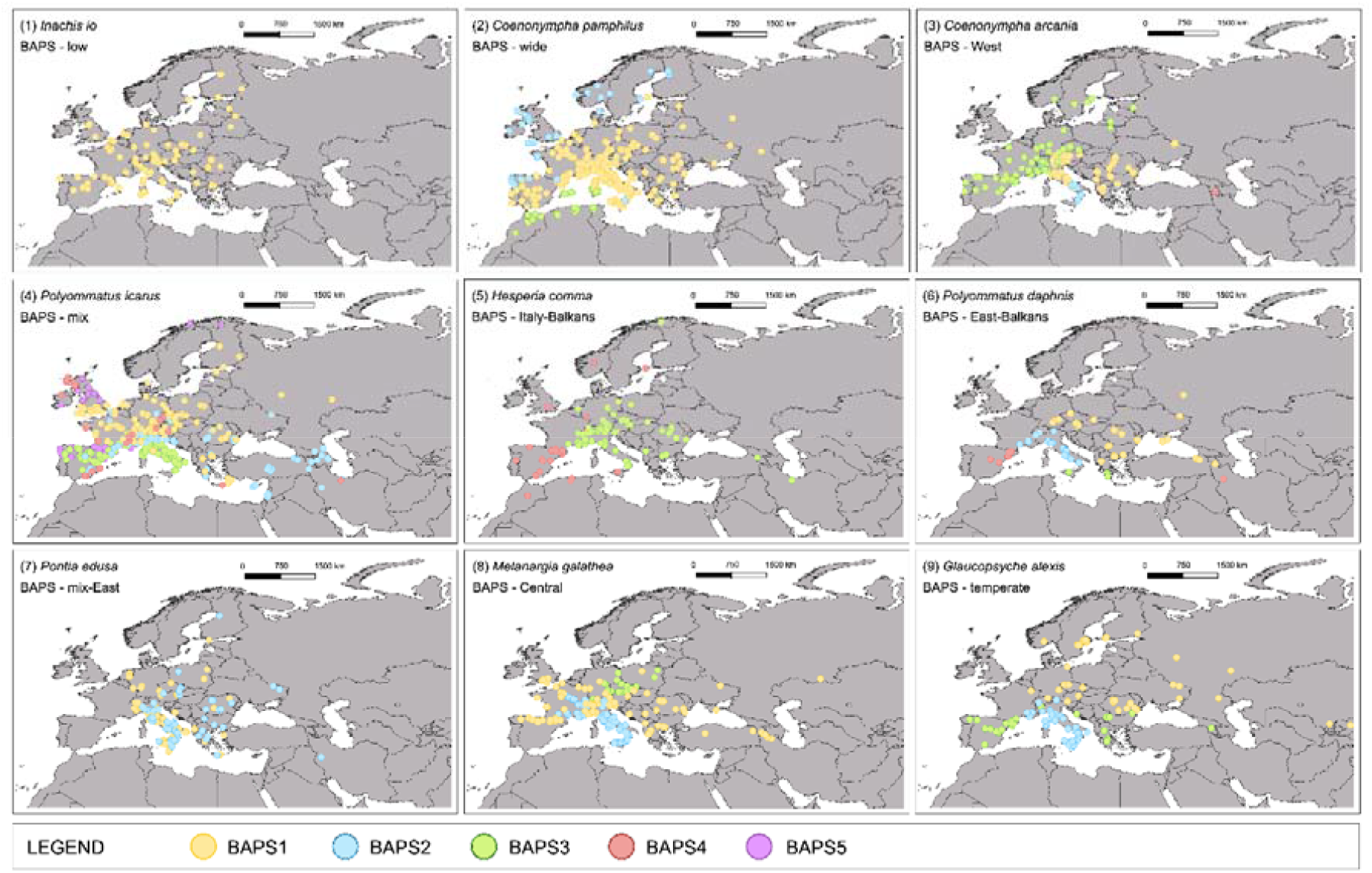
Examples of Czech Republic butterfly species displaying different mitochondrial genetic patterns according to Bayesian Analysis of Population Structure (BAPS categories), depicted on the map of Europe. The method is sorting sequences into clusters of genetically similar individuals (BAPS1–BAPS5, termed here as *genetic lineages*); the number of clusters is estimated automatically by the ‘rhierbaps’ R package (Tonkin-Hill *et al*. 2018) and differs among species. Examining the map, based on the presence of genetic lineages in the Czech Republic, we assigned each of the 140 species into one of nine categories (*BAPS-low, BAPS-wide, BAPS-West, BAPS-mix, BAPS-Italy-Balkans, BAPS-East-Balkans, BAPS-mix-East, BAPS-Central*, and *BAPS-temperate*). Maps were produced in QGIS 3.28.1 (http://qgis.org) with WGS84 projection using country boundaries available at Natural Earth Free vector and raster map data (https://www.naturalearthdata.com/).

**Figure 3.**
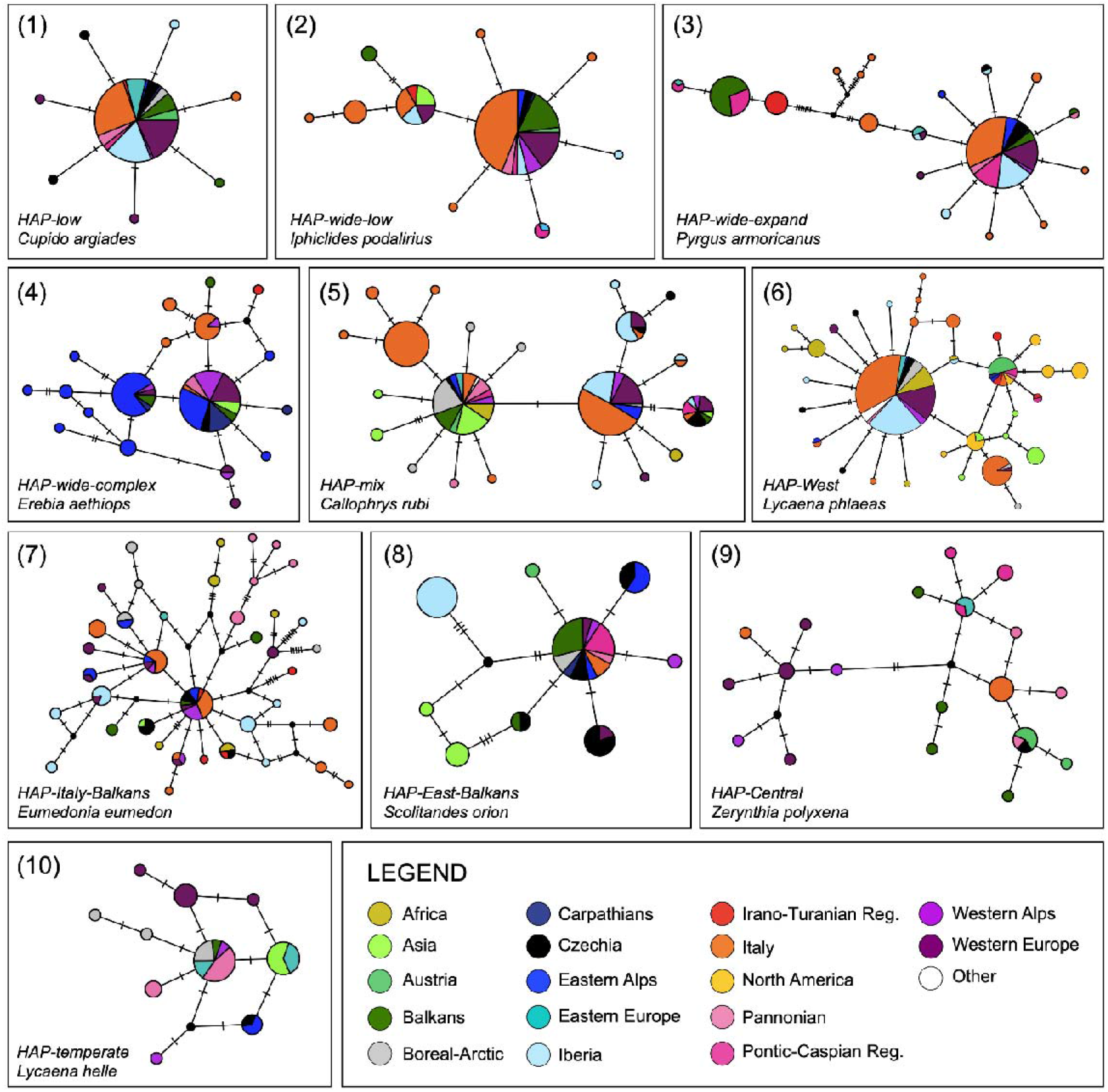
Examples of the Czech butterfly species displaying different mitochondrial genetic patterns, obtained by scoring haplotype networks (HAP). Mutations are depicted as black dots and hatch marks. Examining the network, based on the relations of haplotypes present in the Czech Republic to the other regions, we assigned each of the 140 species into one of ten categories (*HAP-low, HAP-wide-low, HAP-wide-expand, HAP-wide-complex, HAP-mix, HAP-West, HAP-Italy-Balkans, HAP-East-Balkans, HAP-Central*, and *HAP-temperate*).

### Relating BAPS categories to species traits

In the CCA relating BAPS categories to (1) habitat affinity, the global test revealed a significant relationship (eigenvalues: 0.23, 0.14, 0.11, 0.09; first axis var. [explained variation] =3.1%, F=4.2, p=0.046; all axes var.=9.0%, F=1.6, p=0.005). The forward selection identified *ubiquitous* (i.e., generalist, var.=2.2%, F=3.1, p=0.001) and *xerothermophilic 1* (i.e., steppe, var.=1.5%, pseudo-F=2.1, p=0.034) affinities (model’s canonical eigenvalues: 0.18, 0.09; total var.=3.7%). The first canonical axis in the forward selection CCA distinguished *BAPS-East-Balkans, BAPS-Italy-Balkans, BAPS-wide, BAPS-temperate*, and *BAPS-Central* patterns from *BAPS-mix, BAPS-mix-East, BAPS-West*, and *BAPS-low* patterns, and showed that the former tend to be steppe species, whereas the latter are habitat generalists (Fig. 4a).

**Figure 4.**
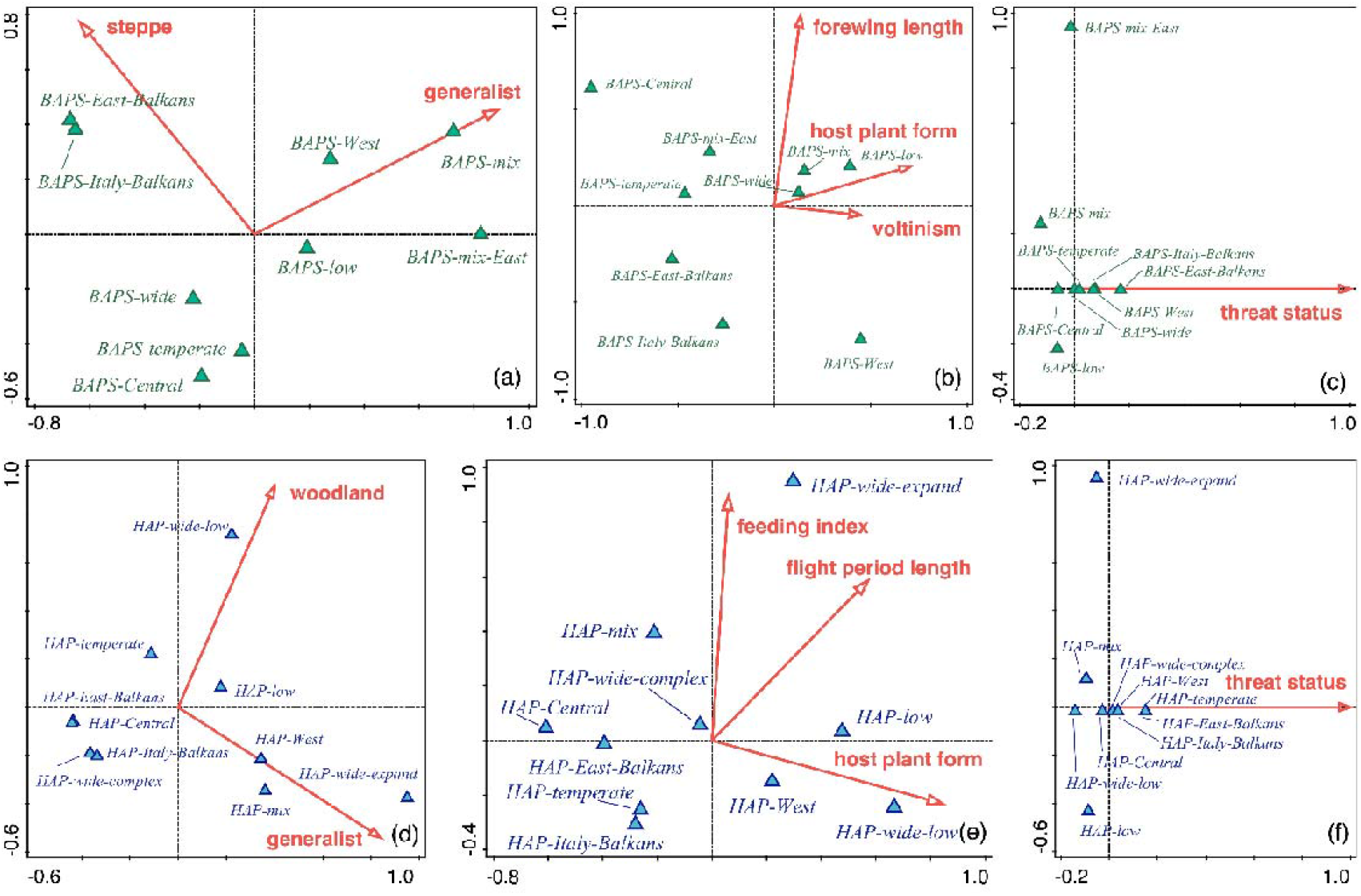
Canonical correspondence analyses (CCA) biplots relating the mitochondrial genetic patterns found in Czech Republic butterflies to individual species traits. The explanatory variables (species traits) were forward-selected. Upper line: (a–c) patterns from Bayesian Analysis of Population Structure (BAPS) interpreted by **(a)** habitats, **(b)** life histories, **(c)** Red List categories (threat status) of individual species. Bottom line: (d–f) patterns from haplotype networks (HAP) interpreted by **(d)** habitats, **(e)** life histories, **(f)** Red List categories (threat status) of the species.

The CCA relating BAPS categories to (2) life history traits also produced significant ordination (global test eigenvalues: 0.22, 0.20, 0.11, 0.07; first axis var. =3.1%, F=4.1, p=0.066; all axes var.=9.0%, F=1.4, p=0.015). The forward-selected traits were *host plant form* (var.=1.7%, F=2.3, p=0.014), *forewing length* (var.=1.6%, F=2.3, p=0.015), and *voltinism* (var.=1.6%, F=2.3, p=0.017) (model’s canonical eigenvalues: 0.19, 0.12, 0.05; total var.=4.9%). The first canonical axis in the forward selection CCA separated species belonging to categories *BAPS-low, BAPS-West, BAPS-mix*, and *BAPS-wide*, from those belonging to *BAPS-Central, BAPS-East-Balkans, BAPS-temperate*, BAPS-*mix-East*, and *BAPS-Italy-Balkans*, indicating that the former tend to develop on woody plants, have more generations per year and a large wingspan; whereas the latter develop on herbs, have less generations and a small wingspan (Fig. 4b).

The BAPS categories were also related to (3) threat status (model’s canonical eigenvalue: 0.13; var.=1.8%, F=2.5, p=0.014). The analysis indicated that species with pattern *BAPS-East-Balkans* tend to be the most threatened, followed by *BAPS-Italy-Balkans*, and *BAPS-West*; and those of *BAPS-mix* are the least endangered (Fig. 4c).

### Relating haplotype network categories to species traits

The global test relating haplotype network categories to (1) habitat affinity was significant (global test eigenvalues: 0.23, 0.19, 0.12, 0.06; first axis var.=2.7%, F=3.6, p=0.068; all axes var.= 8.8%, F=1.4, p=0.012). Two significant habitat affinities were forward-selected: *ubiquitous* (var.=2.2%, F=3.1, p=0.002) and *mesophilic 3* (*i*.*e*., woodland; var.=2.0%, F=2.9, p=0.001) (model’s canonical eigenvalues: 0.21, 0.16; total var.=4.2%). The first canonical axis in the forward selection CCA separated species of patterns *HAP-wide-expand, HAP-mix, HAP-West, HAP-wide-low, HAP-low*, from *HAP-Central, HAP-East-Balkans, HAP-wide-complex, HAP-Italy-Balkans*, and *HAP-temperate*, revealing that the former are habitat generalists and woodland species, whereas the latter are non-woodland specialists (Fig. 4d).

Explaining haplotype network categories by (2) life history traits was significant (global test eigenvalues: 0.35, 0.21, 0.12, 0.10; first axis var.=4.1%, F=5.5, p=0.001; all axes var.=11.2%, F=1.8, p=0.001). The significant forward-selected variables were *host plant form* (var.=2.3%, F=3.2, p=0.001), *flight period length* (var.=1.6%, F=2.3, p=0.02), and *feeding index* (var.=1.3%, F=1.9, p=0.042) (model’s canonical eigenvalues: 0.25, 0.15, 0.05; total var.=5.2%). The forward-selected ordination separated patterns *HAP-wide-low, HAP-low, HAP-wide-expand*, and *HAP-West* from *HAP-Central, HAP-East-Balkans, HAP-Italy-Balkans, HAP-temperate, HAP-mix*, and *HAP-wide complex*. The former species have long flight periods, develop on tree or shrub host plans, and have a broader feeding niche, in contrast to the latter species (Fig. 4e).

The haplotype network categories were also explicable by (3) threat status (model’s canonical eigenvalue: 0.14; var.=1.7%, F=2.3, p=0.008). Species with patterns *HAP-temperate* and *HAP-East-Balkans* tend to be the most threatened, whereas *HAP-wide-low, HAP-mix*, and *HAP-low* are the least threatened (Fig. 4f).

## DISCUSSION

We DNA barcoded 140 species of butterflies occurring in the Czech Republic, used database data to visualize the species’ biogeographic patterns across Europe, and related the mitochondrial genetic structure of Czech samples to species traits. We showed that species’ habitat use and life histories influence which of the mitochondrial lineages inhabit a specific area, and these factors in turn influence their current threat status.

Several authors have related quantitative measures of overall species’ genetic diversity and differentiation to species’ range sizes, life histories and samples’ geographic origins. Genetic diversity varies greatly in European butterflies, as measured by both mtDNA (Dincă et al., 2021) and genome-wide markers (Mackintosh et al., 2019). Mitochondrial COI haplotype diversity itself could not be explained by life history traits determining dispersal and colonization abilities (Dapporto et al., 2019). Generalists do not have greater levels of genomic diversity than specialists; only smaller butterflies tend to have higher genomic diversity (Mackintosh et al., 2019; but see Habel et al., 2013 on landscape level). Population differentiation, on the other hand, is explicable by life histories. Lower differentiation applies to species with longer flight periods and higher number of generations. Accordingly, a higher differentiation was discovered in species with smaller wings, utilising a lower number of host plants, and with short flight periods, i.e., in specialists (Habel et al., 2013; Dapporto et al., 2017, 2019; Scalercio et al., 2020). Widespread species also showed high population differentiation (Dapporto et al., 2019).

We use a more detailed approach here, explaining geographic distribution of individual genetic lineages by species’ traits. The Czech Republic well represents temperate Europe as a whole, owing to its diverse butterfly fauna. We are aware that classification of the lineage distribution is tied to the target region. It is also influenced by the amount of available sequence data and their geographical coverage for a given species. The habitat affinities, and such life history traits as flight period length or host plant use, and of course the level of threat, may also vary across species’ ranges (e.g., Korb et al., 2016; Lindestad et al., 2022). Changing the target region could reveal different lineages for species analysed here, in parallel with changing regional fauna composition and its life histories. Still, our approach offers a template for analysing the constitution of regional faunas.

Both BAPS and haplotype networks categories of lineages present in the Czech butterfly species revealed matching patterns (Fig. 4) when associated with habitats, and analyses of life histories contributed to the understanding of the system (cf. Potocký et al., 2018). In principle, generalists and woodlands dwellers displayed different genetic patterns than specialists and open habitats species. Specifically, generalists and species of wooded habitats formed mixed lineages, or displayed shallow genetic patterns, or had Western affinities (Fig. 4 a, d). Such species frequently develop on shrubs or trees, form multiple generations per year, and have long flight periods, long wings, and wide host plant ranges (Fig. 4 b, e). Species of low genetic diversity (and hence low population differentiation in this case) belong to this group, in agreement with Dapporto et al. (2019). In contrast, open habitats specialists (i.e., steppe species) displayed Southern and/or Eastern affinities, or had temperate patterns, or formed Central European genetic lineages. Such species tended to develop on herbs and grasses and displayed such specialist traits as small wings, low numbers of generations, short flight periods, and narrow ranges of host plants (Bartonova et al., 2014). Such species were linked with high genetic differentiation in Dapporto et al. (2019).

The findings for open habitat species can be attributed to the Quaternary history of biomes in Central Europe. During the mid to late Quaternary, the habitat prevailing in both time and space was the cold mammoth steppe (Kahlke, 2014), associated with diverse types of open habitats. Evidence points to *in situ* survival of steppe lineages in Central Europe through the climatic cycles (Kajtoch et al., 2016; Kirschner et al., 2020; Sucháčková Bartoňová et al., 2021) including the Holocene (Feurdean, Ruprecht, Molnár, Hutchinson, & Hickler, 2018). That long-term persistence could have produced higher population differentiation (Kajtoch, Kubisz, Gutowski, & Babik, 2014; Dapporto et al., 2019; Hawlitschek et al. 2023). In some butterflies targeted here, we indeed observed the Central European genetic patterns (e.g., one lineage in *Melanargia galathea* and *Pyrgus malvae, Polyommatus thersites*).

The association of steppe butterflies with Southern and Eastern lineages is attributable to the geography of the Czech Republic, with lowland corridors situated in the southeast and east (e.g., in *Melitaea didyma, Polyommatus daphnis, Pyrgus carthami*). The Pannonian lowlands, with natural vegetation formed by forest steppes, are connecting the Czech Republic with the Balkan Peninsula. An alternative route is through the Moravian Gate, connecting Czech territory with the lowlands of southern Poland, continuing to the Pontic steppes (the “Sarmatian route”; Mařan 1958; Sternberg 1998). On the other hand, the warm lowlands in Bohemia (Elbe river valley) were likely colonized by steppe elements from Moravia, blocked by mountain chains from similar habitats in Western Europe. This scenario was documented, e.g., for the ground squirrel *Spermophilus citellus* (Říčanová et al., 2013).

Multiple species had a genetic lineage distributed along a single latitudinal belt (temperate patterns), lacking the Mediterranean distribution. These were species of open habitats and habitat specialists. Many genetic lineages could have inhabited the Palearctic continent alongside a temperature belt (Maresova et al., 2021), with no barriers to dispersal in the east-west direction. These species might follow the east-west oceanic-continental gradient of glacial-interglacial faunal interchange related to moisture (Stewart et al., 2010). Continental species might have expanded west during the arid glacials, establishing stable widespread populations, with Mediterranean peninsulas serving as areas of endemism rather than refugia for them (e.g., in *Agriades optilete, Boloria euphrosyne, Glaucopsyche alexis*).

Woodlands re-expanded across Europe during the Holocene. Thus, most of the populations of forest species are relatively newly established and shallow genetic structures, which we observed for woodland inhabitants and tree feeders, are expected (e.g., in the European woodland species *Pararge aegeria*, Livraghi et al., 2018; or in *Nymphalis polychloros, Celastrina argiolus*). Habitat generalists display similar genetic patterns to the woodland species, linked especially with a mix of lineages inhabiting the focal area, which likely diversified in multiple refugia (e.g., *Pieris napi, Polyommatus icarus*). Longer flight periods, multiple generations and larger bodies, expressed as longer wings, might facilitate such swift expansions, making these species good long-distance dispersers (Stevens, Trochet, Van Dyck, Clobert, & Baguette, 2012). Species with Western patterns (lineages connected to Iberia, or Iberia and Italy) possess similar traits (e.g., *Lycaena phlaeas, Lasiommata megera*). Better dispersers might be equipped to cross the mountains (Fric, Hula, Konvička, & Pavlíčko, 2000), and expansions of woodland butterflies could have been facilitated by the earlier onset of the Holocene near the Atlantic coast (Heiri et al., 2014), and slowed down by continental climates surrounding the Balkans, and/or East Asian refugia.

The most threatened species in the Czech Republic are those having multiple differentiated lineages across Europe with a single lineage inhabiting the focal region (*BAPS-East-Balkans, BAPS-West, BAPS-Italy-Balkans* and *BAPS-temperate*) (e.g., *Pyrgus alveus, Hipparchia hermione, Polyommatus dorylas*, and *Polyommatus damon*), whereas the generalists (*BAPS-mix, HAP-mix* or *HAP-low*) are the least endangered (Fig. 4 c, f). This can be related to the dispersal abilities of the species (Dapporto et al., 2019): the higher the differentiation, the more specialized the butterfly is, and the less likely to survive in simplified modern landscapes (Bartonova et al., 2016; Ockinger et al., 2010) because of reduced possibility of rescue effect (Brückmann, Krauss, & Steffan-Dewenter, 2010). This is supported by the evidence that butterfly species with a higher population diversification disappeared first from islands (Dapporto et al., 2017). The species that display multiple lineages across Europe are also threatened in other European countries (Maes et al., 2019).

The existence of distinct lineages across the species ranges was noted already by de Lattin (1967), and, with the advent of genetic data, investigated in many case studies. Only now, however, does the vast DNA database data allow us to test predictions on the resolution of entire faunas. For a better understanding of lineage distribution and suture zones, it would be necessary to properly cover the whole of Europe, and extend the sampling beyond, to obtain information for entire species ranges. DNA barcoding of specimens in under-sampled areas, even if not particularly biodiversity-rich, still represents a powerful tool for discovering the driving forces of biogeography on large scales, such as the links between phylogeography and species-specific traits.

## DATA AVAILABILITY

The newly generated sequences for Czech butterfly species are deposited in the BOLD database as the BBCZ Project [CZBB001–CZBB959], and in the NCBI database [accession codes OR890444–OR891402]. The Supplementary Materials (specimen metadata, Nexus alignments for each species, R script for data handling, species traits and assigned phylogeographic patterns) are stored at the figshare repository DOI 10.6084/m9.figshare.24559852.

## ACKNOWLEDGEMENTS

The work was supported by the Technology Agency of the Czech Republic (SS01010526). We would like to thank the Nature Conservation Agency of the Czech Republic and Ministry of the Environment of the Czech Republic, as well as all the local conservation authorities (National Parks and Protected Landscape Areas). The following colleagues helped us with material: Oldřich Čížek, Marek Fišer, Vladimír Hula, Tomáš Kadlec, František Kopeček, Tomáš Kuras, Dan Leština, Zdeněk Mráček, Alois Pavlíčko, David Ričl, Ondřej Sedláček, Pavel Skala, Přemysl Tájek, Pavel Vrba, and Jan Walter. We would also like to thank Petr Šmilauer for consultation on statistics, Pedro de G. Ribeiro and Marie Drábková for comments on the manuscripts, and Matthew Sweney for English correction.

## BIOSKETCH

The authors are experts in insect ecology, phylogeny, phylogeography, and distribution patterns. The authors focus on the history, the current times and a future of Quaternary ecosystems. They utilize butterflies, and other insects, as examples, to study the impact of climate change, extinct megafauna, and habitat alterations on communities. They combine the diverse scientific approaches and knowledge of the past to contribute to current habitat and insect conservation. Author contributions: ASB designed the study, collected data, performed analyses, and wrote the first draft of the manuscript. PŠ performed laboratory work and analyses. MK collected data and contributed to the writing of the first draft. JB, LS, and VV collected data. CS and HK performed laboratory work. JPM prepared all graphics in the manuscript and supplements. ZFF designed the study, collected data, and performed analyses. All authors contributed substantially to revisions and approved the final version of the manuscript.

## SUPPORTING INFORMATION

**Supporting Information 1**. Detailed summary and results of the Czech Republic butterfly barcoding (sequencing cytochrome c oxidase subunit 1, COI) for each species occurring in the country.

