## Supporting Information 1. for "Czech Republic butterfly barcoding reveals that distribution of genetic lineages depends on species traits" (from another version of this preprint: version 1, 2024-01-22)

Detailed summary and results of the Czech Republic butterfly barcoding (sequencing cytochrome c oxidase subunit 1, COI) for each species occurring in the country.

**The maps** show results of the Bayesian Analysis of Population Structure (BAPS) across Europe (genetic lineages BAPS1–BAPS6) **(a)**, and a detailed sampling in the Czech Republic **(b)**. Haplotype networks **(c)** show phylogenetic relationships of samples. The backgrounds of the networks are shaded based on lineages revealed by BAPS. In some cases, a haplotype encompasses more BAPS lineages, which is caused by the computational differences of the two methods (missing data handling). The grid map **(d)** shows results from the Czech Butterflies and Moths Recording [CBMR] valid for 2016: the colour dots stands for the newest records in a particular cell: **blue**: ≤ 1950, **green**: 1951–1980, **yellow**: 1981–1996, **orange**: 1997–2001; **red**: 2002–2016. All maps were created using Free and Open Source QGIS 3.28.1 (http://qgis.org) and the graphics were edited in Graphic for Mac v. 3.1 (https://www.graphic.com/mac/) and Inkscape 1.0.2 (https://inkscape.org/).

**The caption** of each Figure includes categories of species' habitat (Benes et al., 2002), national Red List (Hejda, Farkač, & Chobot, 2017), assigned BAPS and haplotype network (HAP) categories, and range, and commented current distribution in the Czech Republic, including comments on the genetic structure when needed.

The **habitat** categories: ubiquitous (generalist), mesophilic 1 (grassland), mesophilic 2 (shrubland), mesophilic 3 (woodland), xerothermophilic 1 (steppe), xerothermophilic 2 (dry shrubland), hydrophilic (wetland), tyrphophilic (bog/peatland), and alpine.

The **Red List** categories: least concern (LC), near threatened (NT), vulnerable (VU), endangered (EN), and critically endangered (CR).

The **BAPS** categories, based on the presence of genetic lineages (BAPS1–BAPS6) in the Czech Republic: BAPS-low, BAPS-wide, BAPS-mix, BAPS-West, BAPS-Italy-Balkans, BAPS-East-Balkans, BAPS-mix-East, BAPS-Central, and BAPS-temperate.

The **HAP** categories, based on the position of Czech samples in the network: HAP-low, HAP-wide-low, HAP-wide-expand, HAP-wide-complex, HAP-mix, HAP-West, HAP-Italy-Balkans, HAP-East-Balkans, HAP-Central, and HAP-temperate.

The **range**: cosmopolite, Holarctic, Palearctic, Euro-Siberian (not inhabiting southern Palearctic), Western Palearctic (west of Altai Mts.), European (west of Ural Mts.), Pontic-Mediterranean, Atlantic-Mediterranean, and Mediterranean.

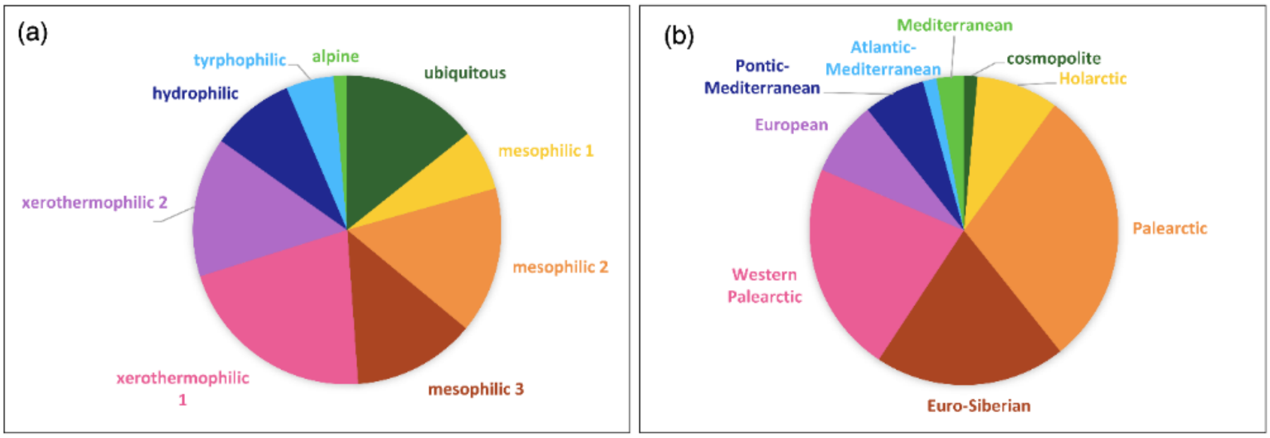


**Figure S1.1.** The distribution of categories of species' habitat **(a)** and range **(b)** in 140 Czech butterflies (~98% of fauna).

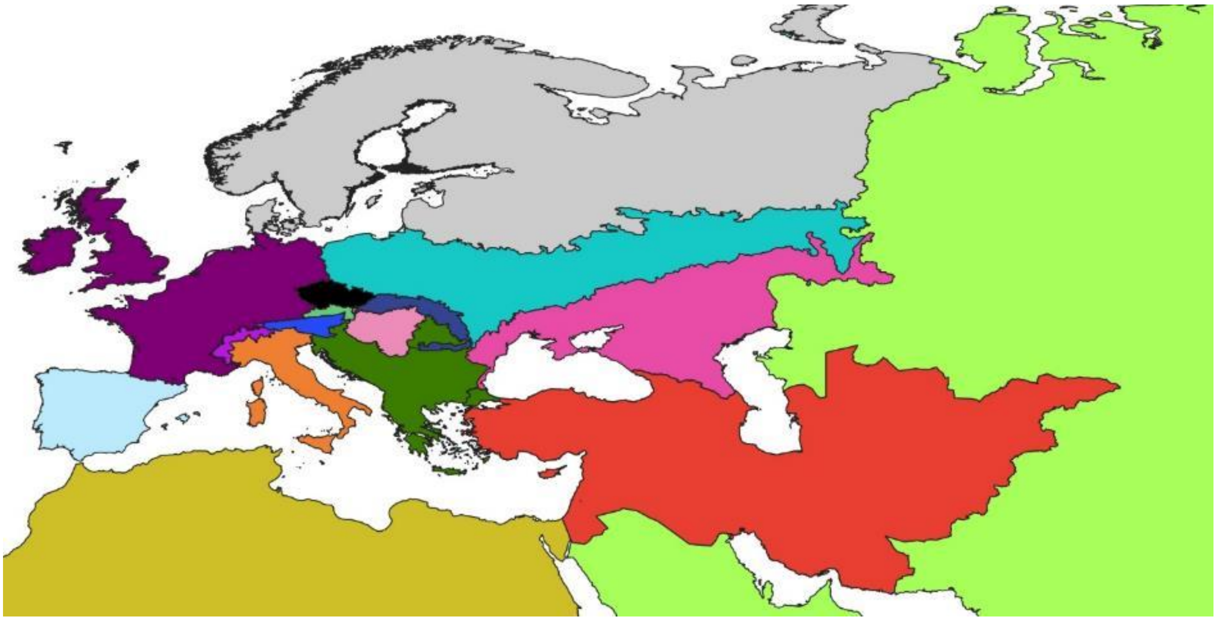

**Figure S1.2.** The boundaries of regions used to visualize haplotypes' origin in the haplotype networks (Western Palearctic part). The shapefile was created using QGIS 3.28.1, as a combination of country boundaries (Natural Earth Free vector and raster map data: naturalearthdata.com) and biogeographic regions of Europe (© EuroGeographics, © FAO (UN), © TurkStat Source: European Commission – Eurostat/GISCO, available at https://www.eea.europa.eu/en/datahub/datahubitem-view/11db8d14-f167-4cd5-9205-95638dfd9618).

**The Czech Republic** (Czechia, CZ), situated in Central Europe, has an area of 78,871 km$^2$. The country consists of three historical lands: Bohemia, covering western two thirds of the area; Moravia, covering most of the eastern third of the area; and Silesia, which reaches CZ from northeast. The western part belongs to old Palaeozoic Bohemian Massif, forming the borderland mountains, with its northern, highest parts Sudeten Mts. Some of the inner parts of Bohemian Massif are formed by Tertiary volcanic highlands, such as Central Bohemian Uplands. The Carpathian Mountains arc of Tertiary origin reaches the east (White Carpathians, Beskids). Warm Pannonian lowlands are reaching the country in the southeast (South Moravia). The warm (thermophilic) parts are present also in northwestern and central parts of Bohemia due to the rain shadow of the borderland mountains.

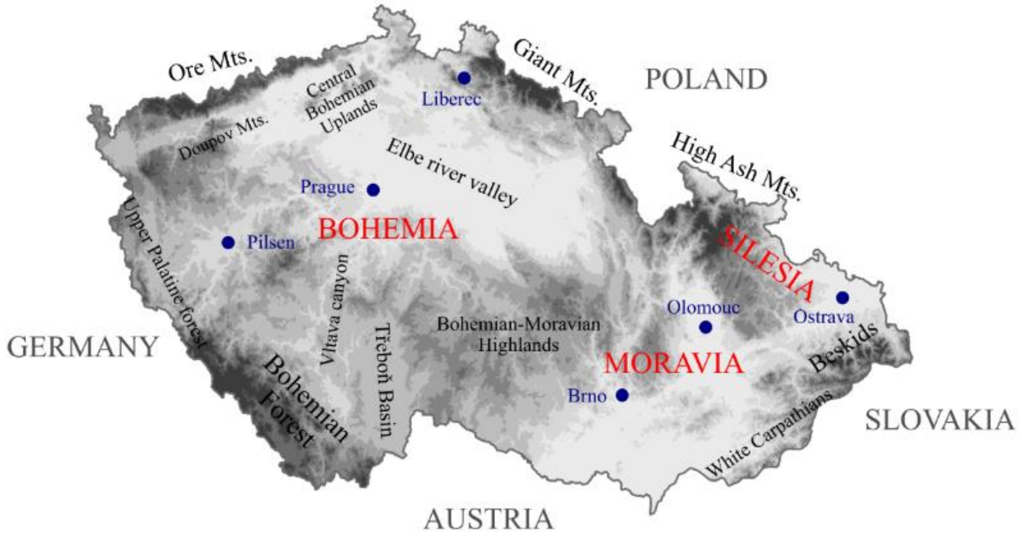


**Figure S1.3.** The topography map of the Czech Republic. The topography was obtained from free resource database web GIS (http://www.webgis.com/terr_world.html).

Glossary of Czech and English geographical names used in the Figure captions:

Beskydy = Beskids
Bílé Karpaty = White Carpathians
České středohoří = Central Bohemian Uplands
Českomoravská vrchovina = Bohemian-Moravian Highlands
Český les = Upper Palatine Forest
Doupovské hory = Doupov Mountains
Hrubý Jeseník = High Ash Mountains
Krkonoše = Giant Mountains
Krušné hory = Ore Mountains
Labe = Elbe river
Šumava = Bohemian Forest
Třeboňsko = Třeboň Basin

**Papilionidae**

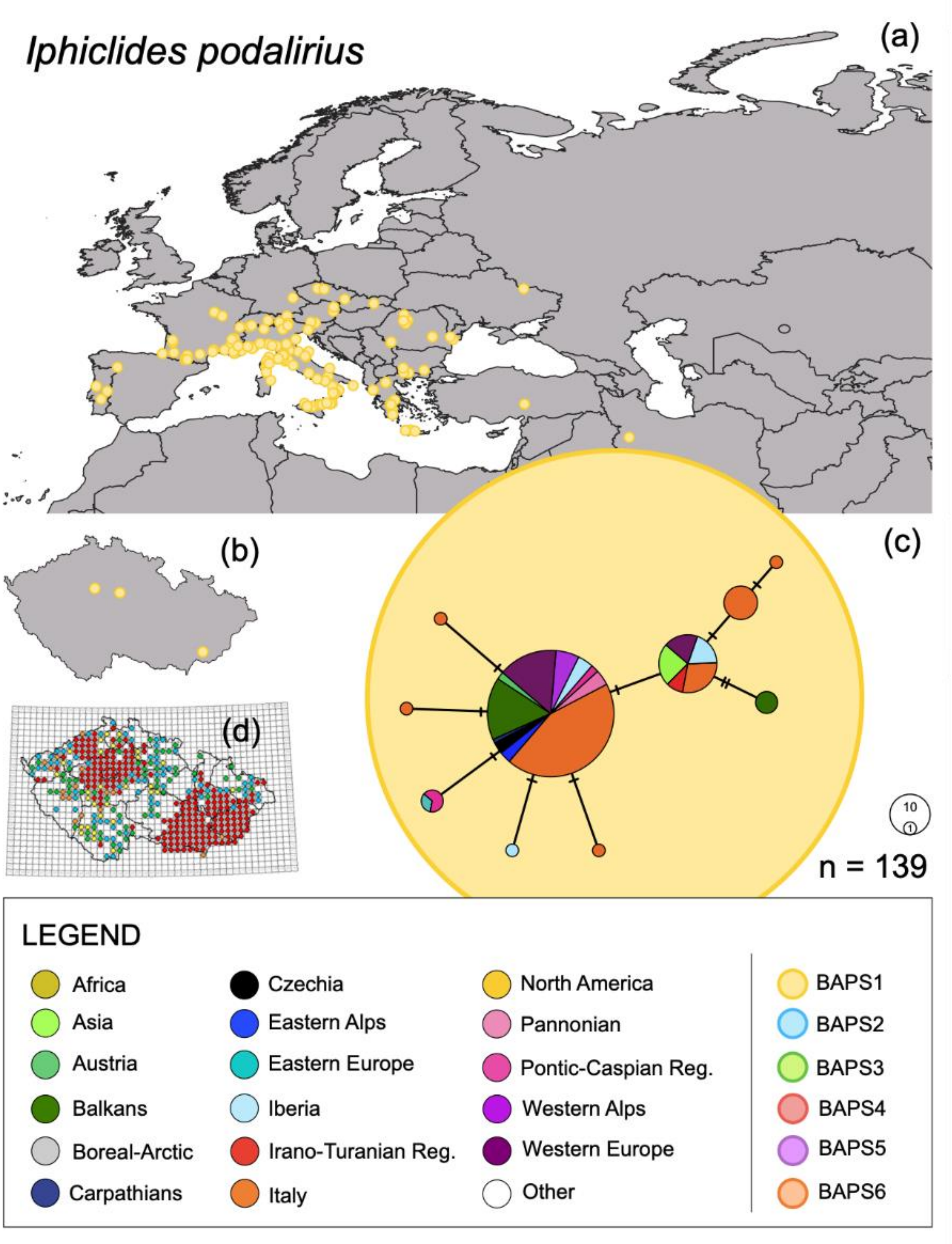


*Iphiclides podalirius* (Linnaeus, 1758): **Habitat:** xerothermophilic 2. **Red List:** NT

**BAPS:** BAPS-low. **HAP:** HAP-wide-low

**Range:** Western Palearctic. **Distribution CZ:** warm sunny rocky outcrops, dry scrub and forest steppes. Thermophilic regions of CZ, retreated in the past but now expanding with warmer climate.

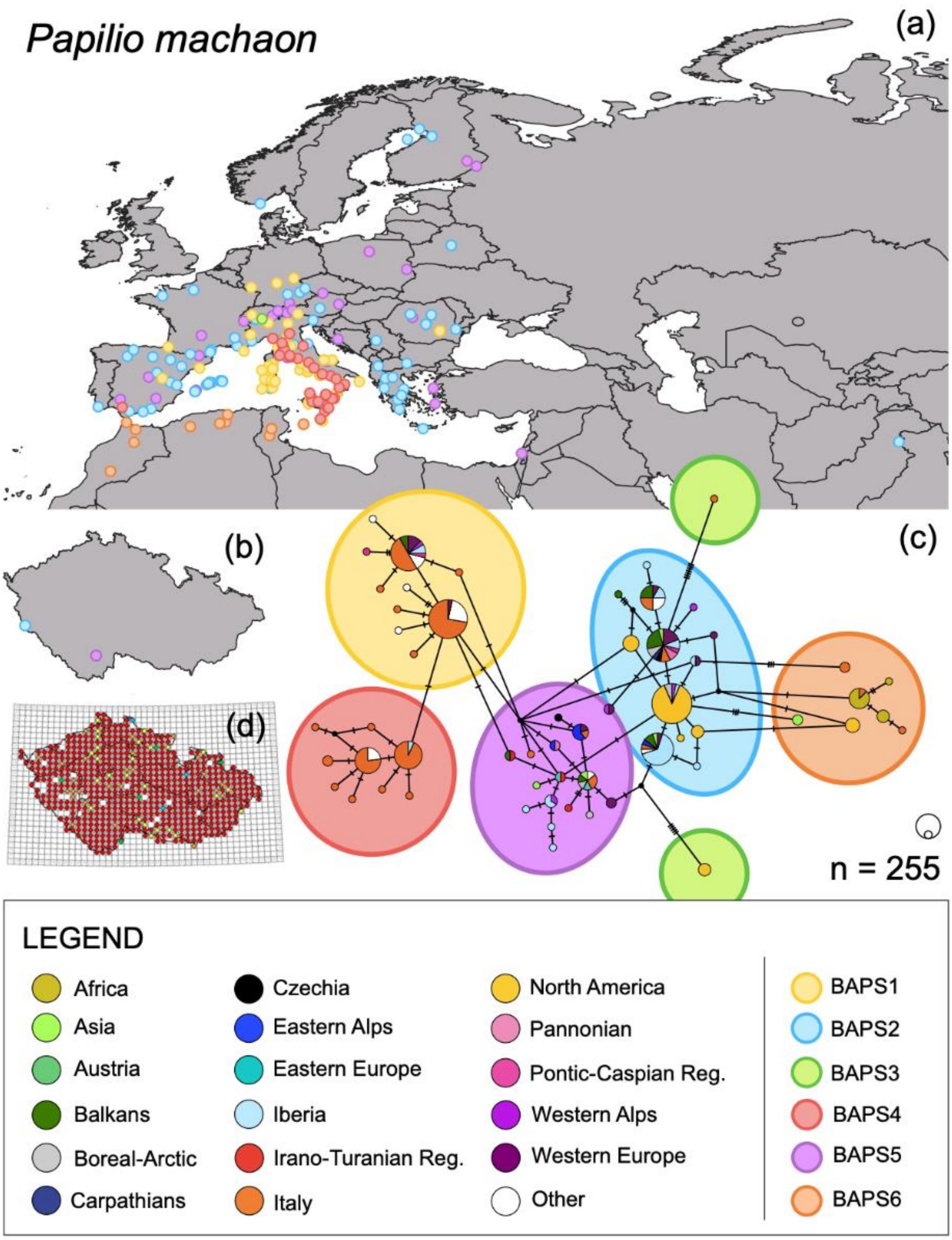


*Papilio machaon* Linnaeus, 1758: **Habitat:** ubiquitous. **Red List:** LC

**BAPS:** BAPS-mix. **HAP:** HAP-mix

**Range:** Holarctic. **Distribution CZ:** diverse habitats, widespread across the country.

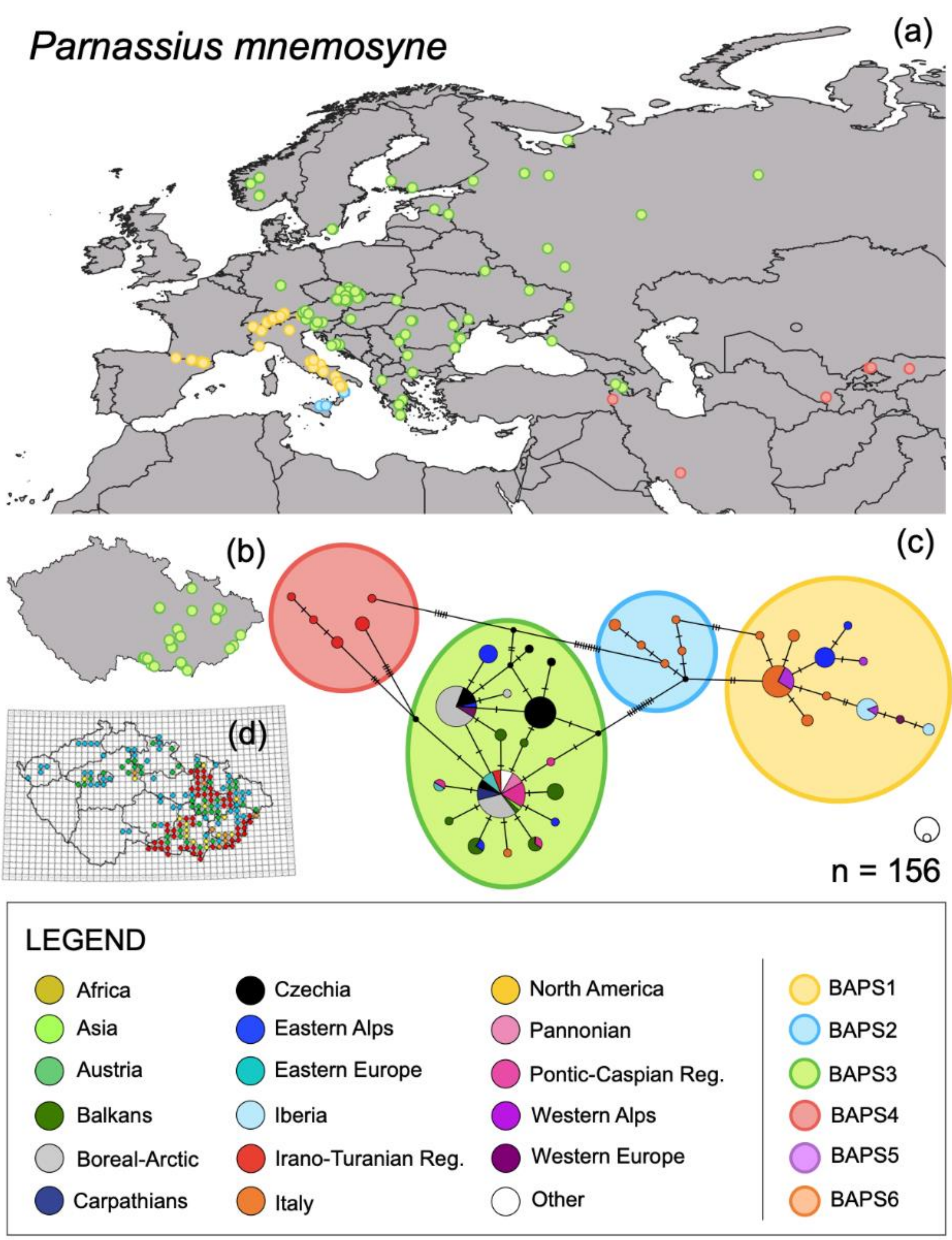


*Parnassius mnemosyne* (Linnaeus, 1758): **Habitat:** mesophilic 1. **Red List:** EN

**BAPS:** BAPS-East-Balkans. **HAP:** HAP-East-Balkans

**Range:** Western Palearctic. **Distribution CZ:** open deciduous woodlands, coppiced woodlands. Severely declined (loss of open woodlands), extinct from Bohemia, several metapopulations still hold across Moravia.

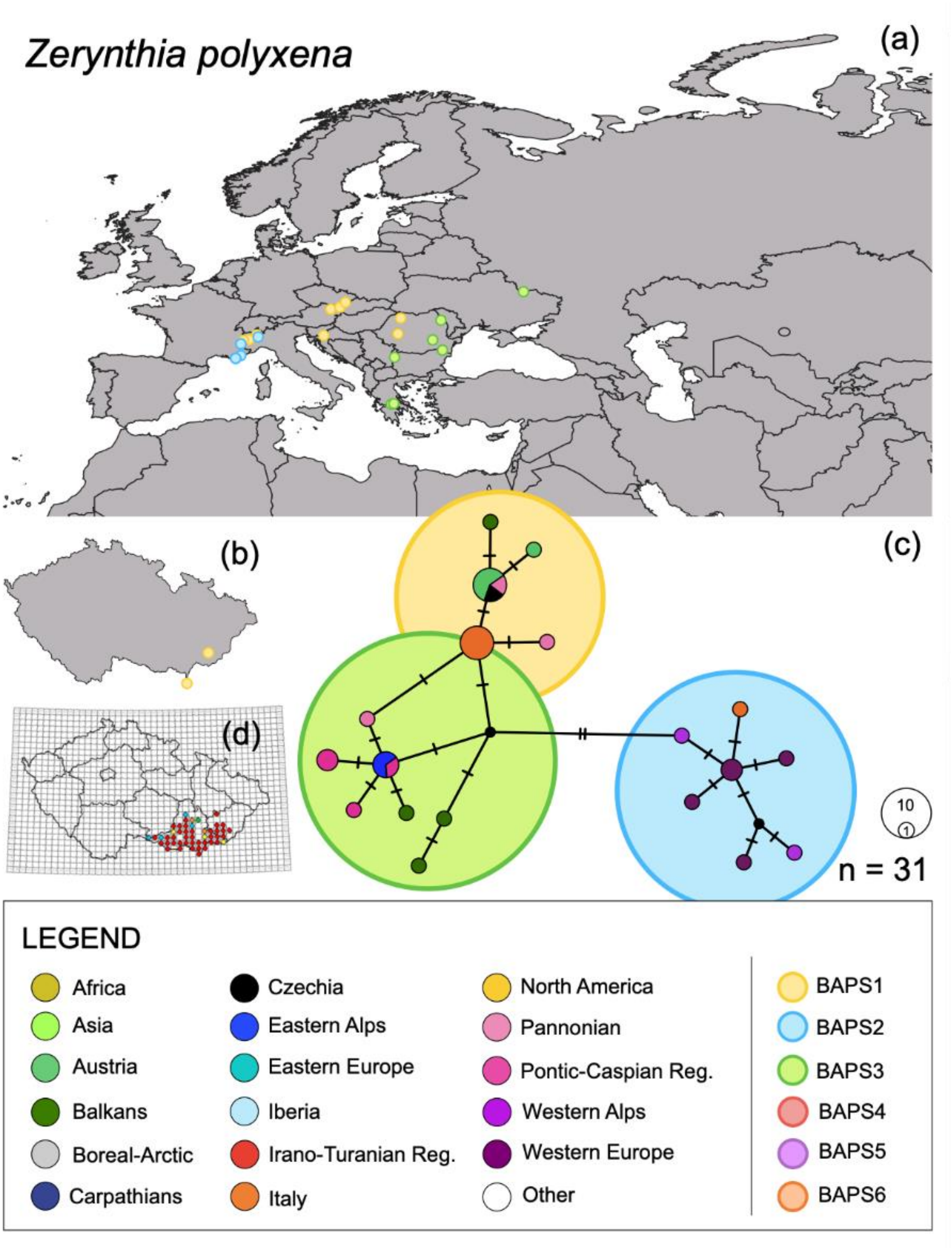


*Zerynthia polyxena* ([Denis & Schiffermüller], 1775): **Habitat:** mesophilic 2 + xerothermophilic 2. **Red List:** NT

**BAPS:** BAPS-Central. **HAP:** HAP-Central

**Range:** Pontic-Mediterranean. **Distribution CZ:** abandoned woodland margins, river valleys. Both historically and currently restricted to South Moravia, decreased with agricultural intensification.

### Hesperiidae

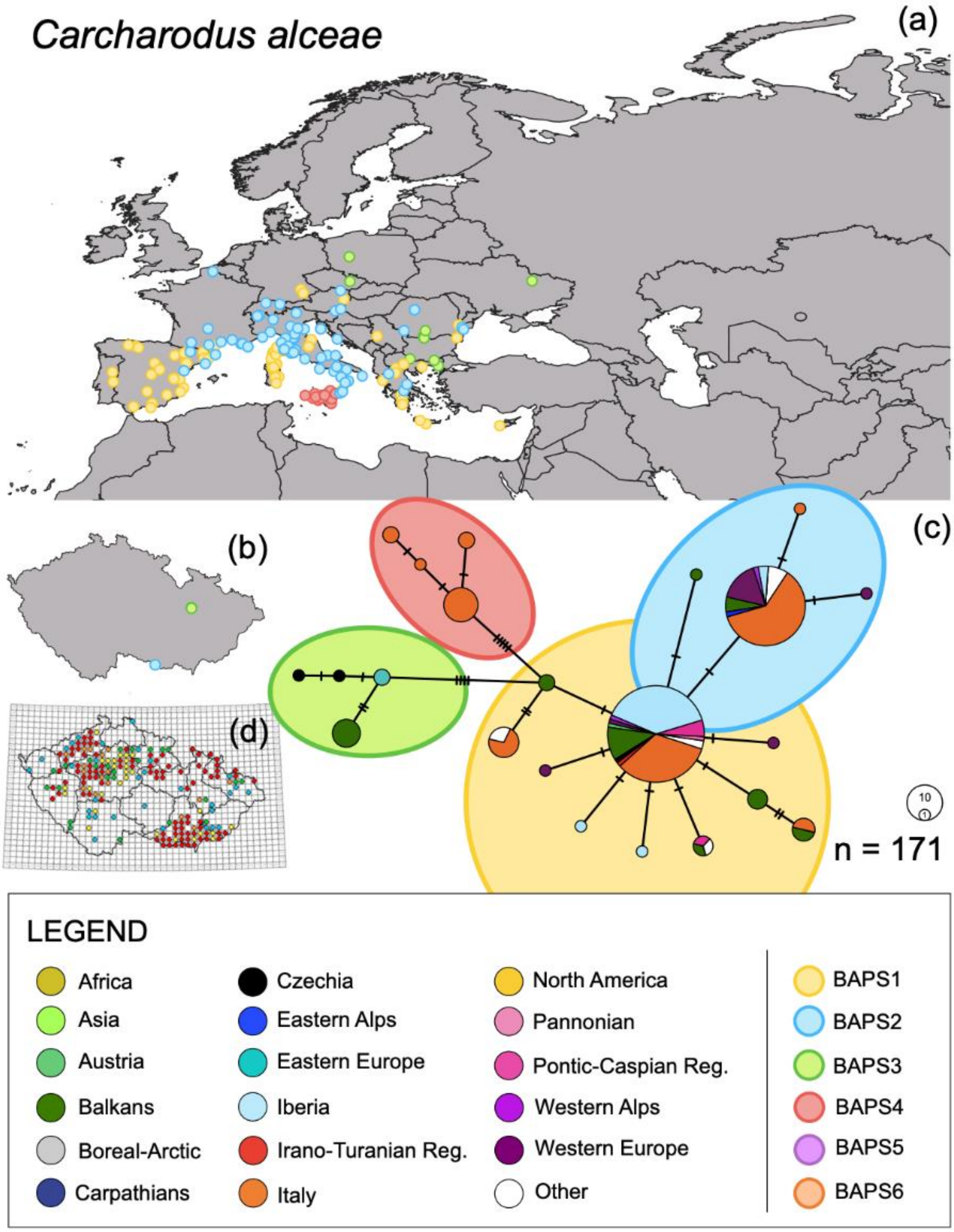


*Carcharodus alceae* (Esper, 1780): **Habitat:** xerothermophilic 2, **Red List:** NT

**BAPS:** BAPS-wide + BAPS-East-Balkans. **HAP:** HAP-wide-complex + HAP-East-Balkans

**Range:** Western Palearctic. **Distribution CZ:** forest steppes and abandoned habitats in warmer regions (Central Bohemia, Central Bohemian Uplands, South Moravia) and random records from other regions in the last decades.

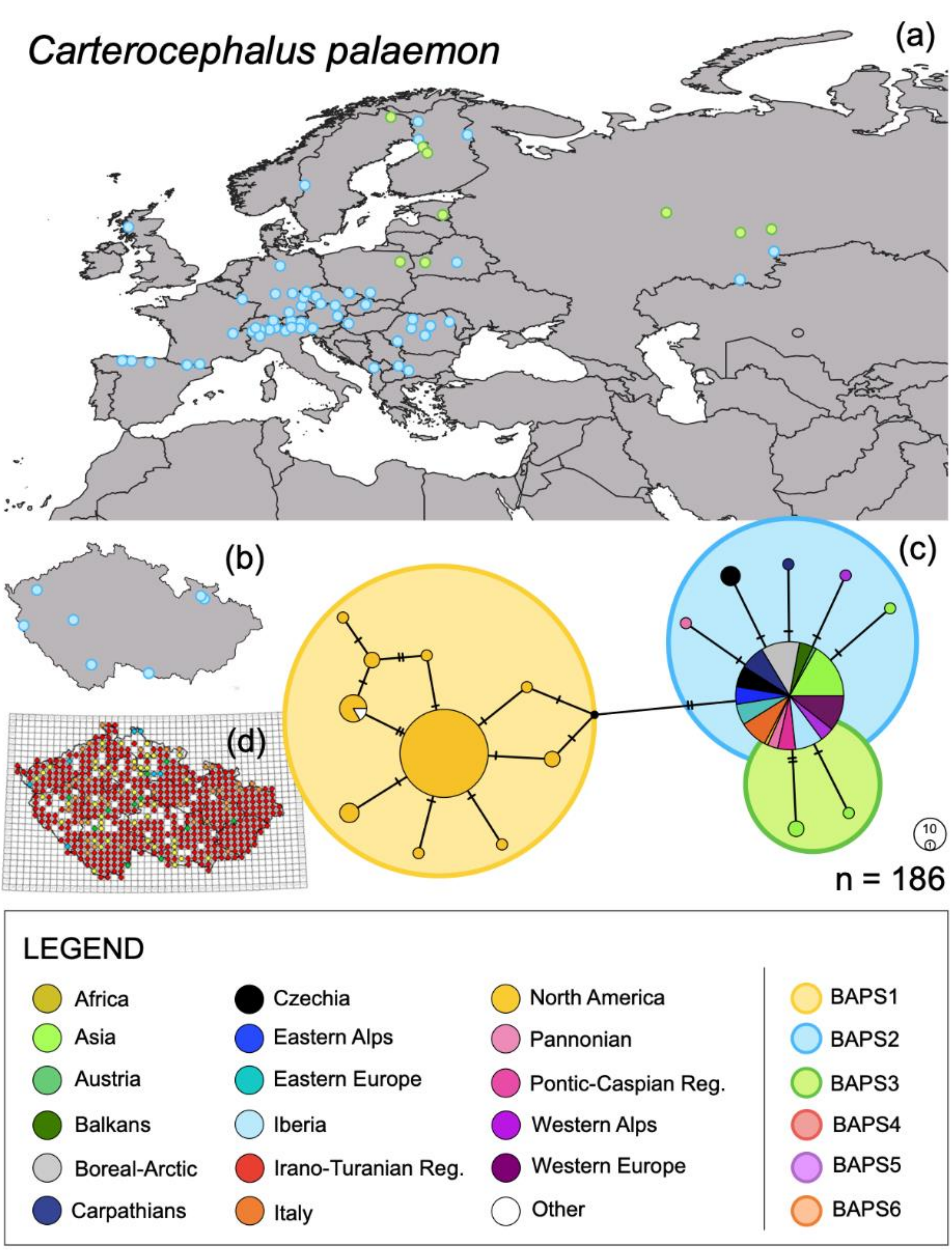


*Carterocephalus palaemon* (Pallas, 1771): **Habitat:** mesophilic 2 + hydrophilic. **Red List:** LC

**BAPS:** BAPS-temperate. **HAP:** HAP-temperate (such distribution).

**Range:** Holarctic. **Distribution CZ:** wet meadows, woodland margins and clearings. Widespread across the country, common.

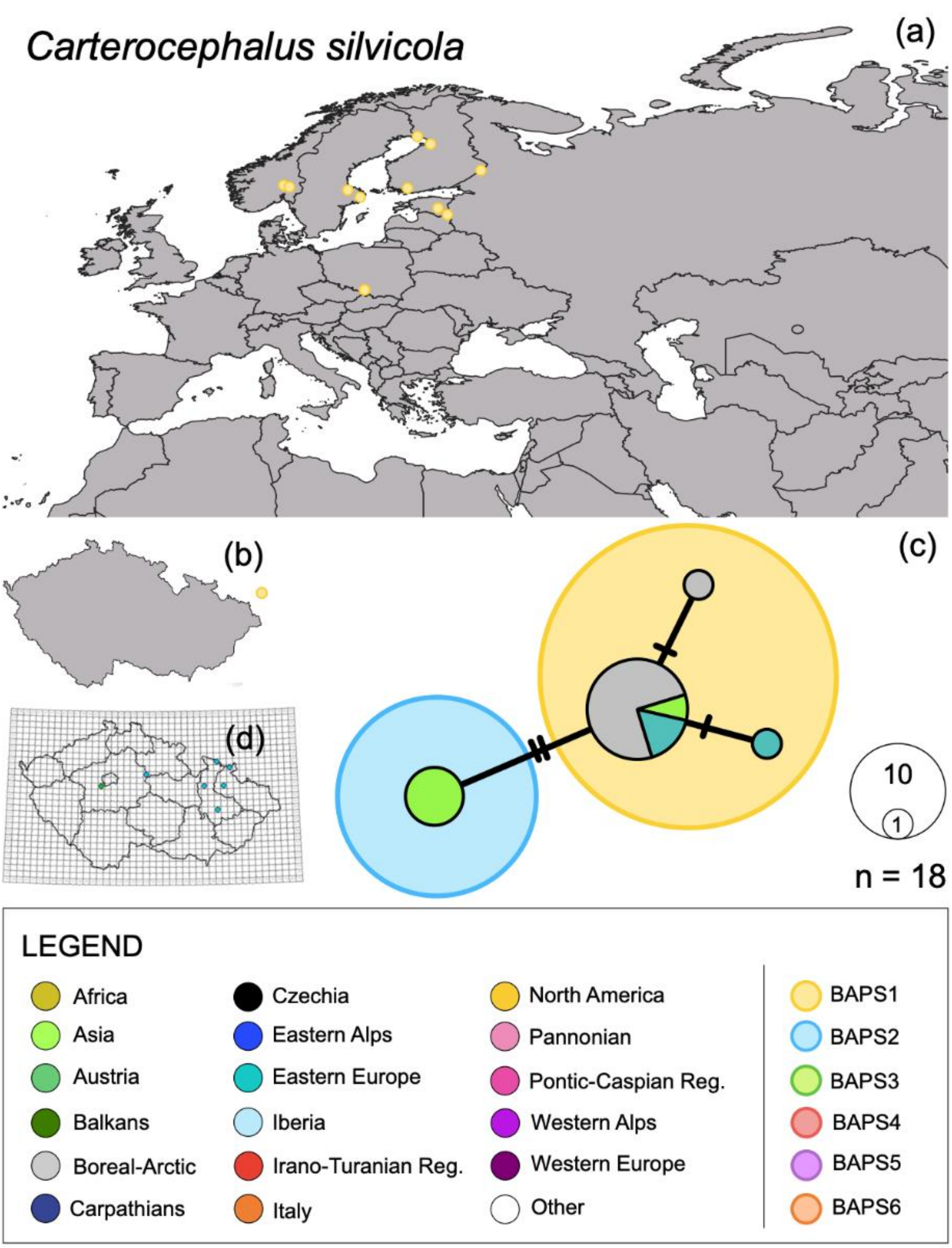


*Carterocephalus silvicola* (Meigen, 1829): **Habitat:** hydrophilic. **Red List:** EX
**Range:** Euro-Siberian. **Distribution CZ:** Extinct, not included in analyses. The analysed sample was collected close to the Czech border, in southern Poland, of probable similar origin to the population that once inhabited the Northeast of CZ. The species was in its southern margin of distribution in CZ. Observed for the last time in 1950s.

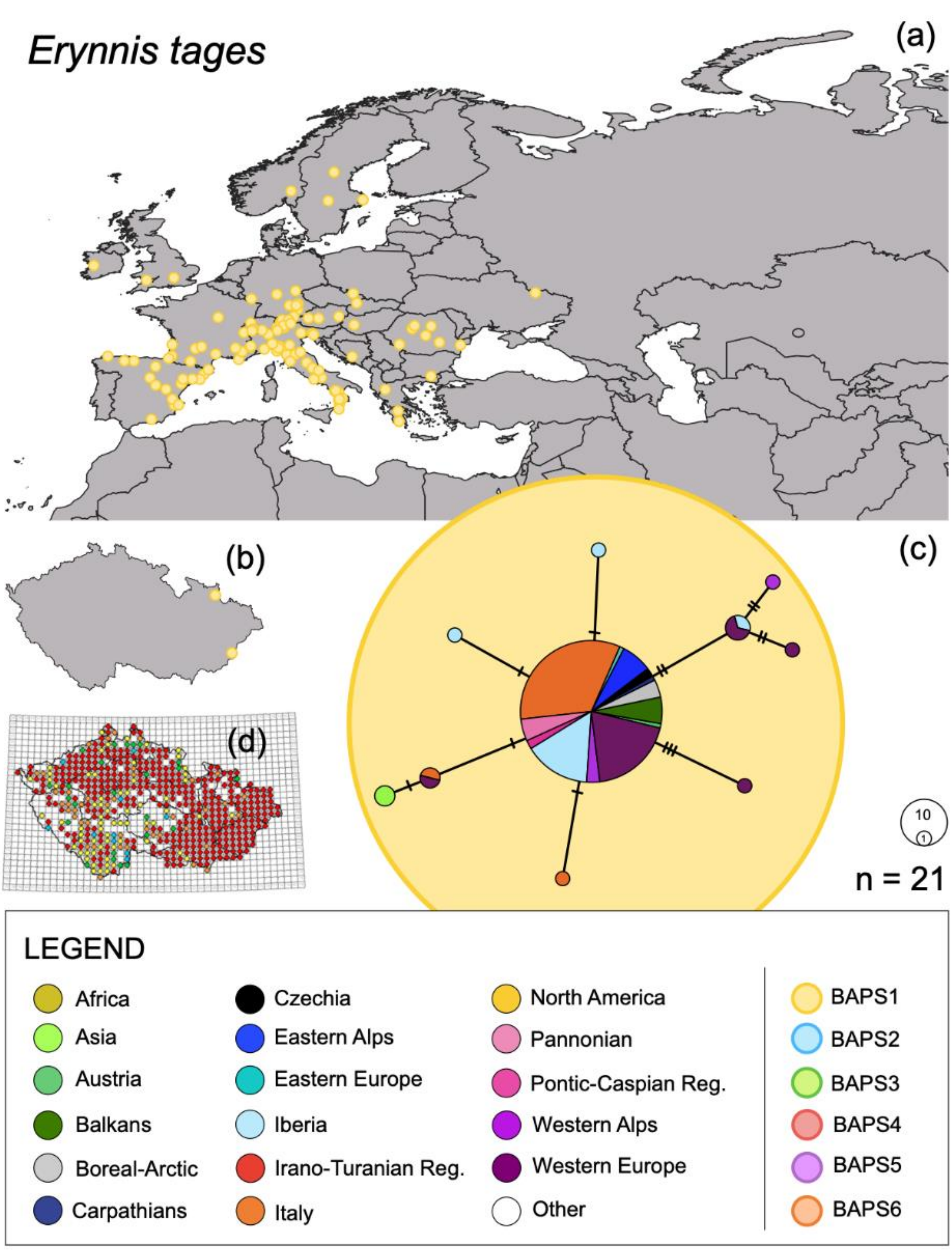


*Erynnis tages* (Linnaeus, 1758): **Habitat:** xerothermophilic 1. **Red List:** LC

**BAPS:** BAPS-low. **HAP:** HAP-wide-low

**Range:** Euro-Siberian. **Distribution CZ:** open places, dry grasslands and margins. Widespread across the country.

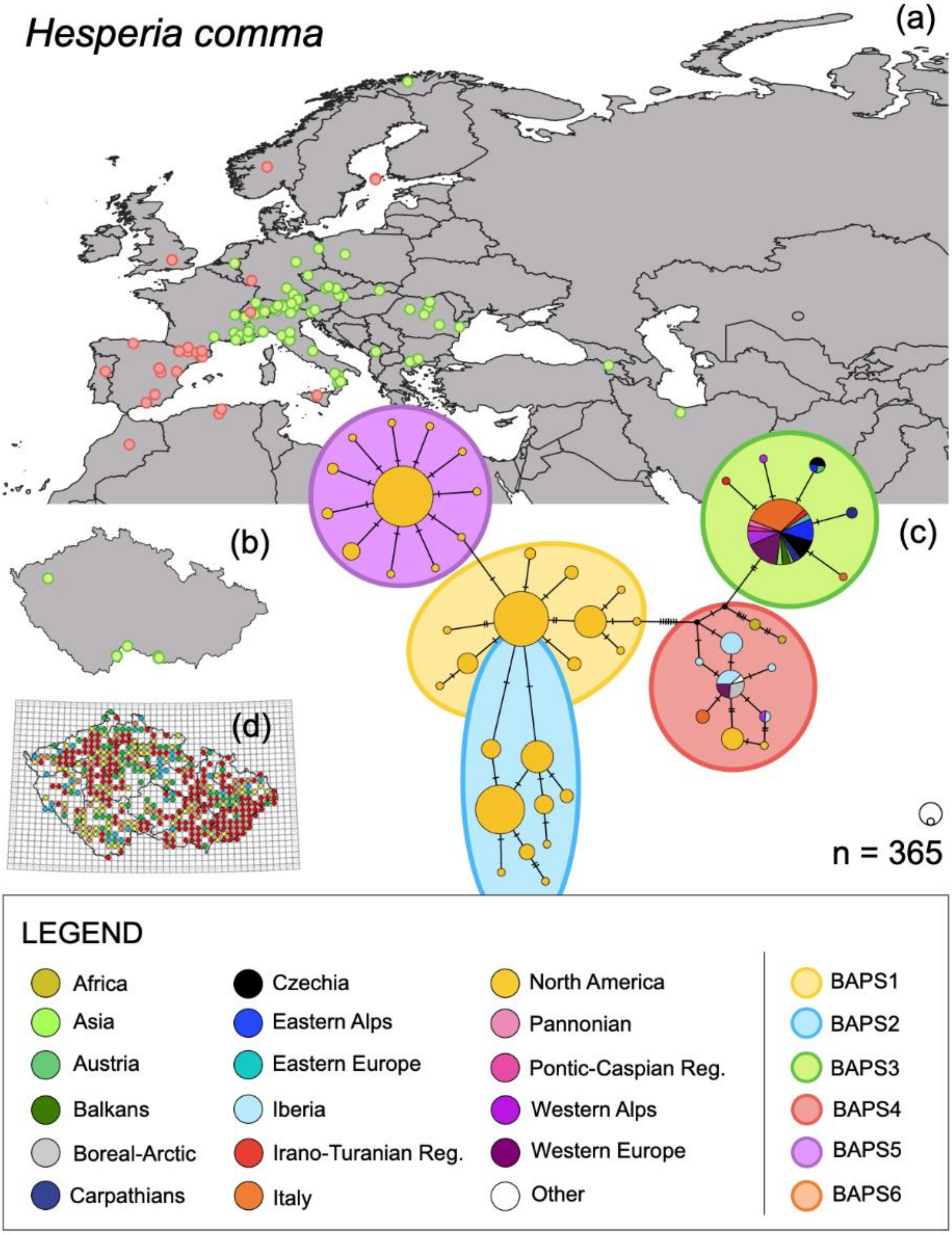


*Hesperia comma* (Linnaeus, 1758): **Habitat:** xerothermophilic 1, **Red List:** VU

**BAPS:** BAPS-Italy-Balkans. **HAP:** HAP-Italy-Balkans

**Range:** Holarctic. **Distribution CZ:** dry flowery grasslands. Entire area but localized, currently returning back after a decrease.

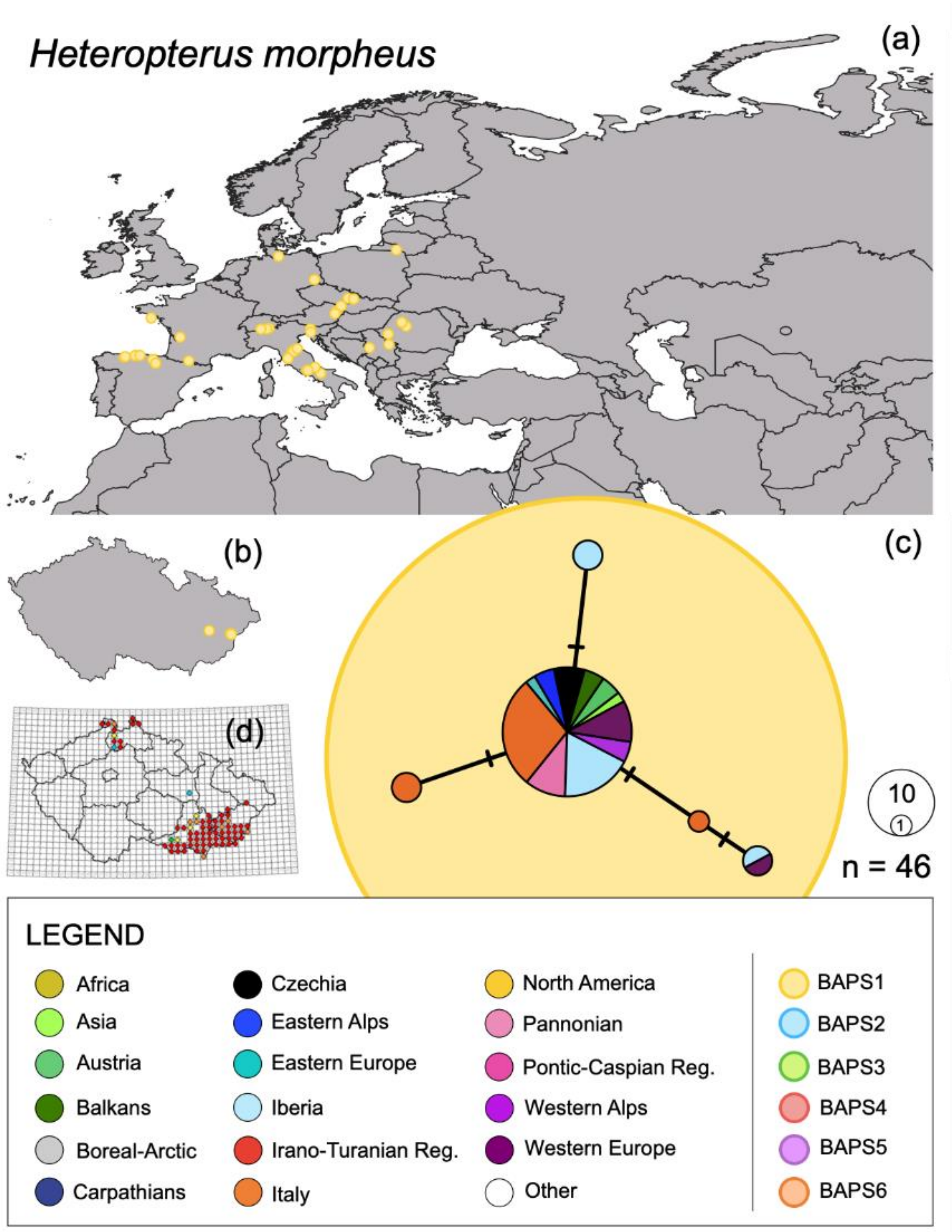


*Heteropterus morpheus* (Pallas, 1771): **Habitat:** xerothermophilic 2 + hydrophilic. **Red List:** LC

**BAPS:** BAPS-low. **HAP:** HAP-low

**Range:** Euro-Siberian. **Distribution CZ:** wet abandoned meadows, woodland margins and clearings, close to water bodies. Disjunctive distribution in CZ: in south and southeast of Moravia, and in north of Bohemia. Expanding.

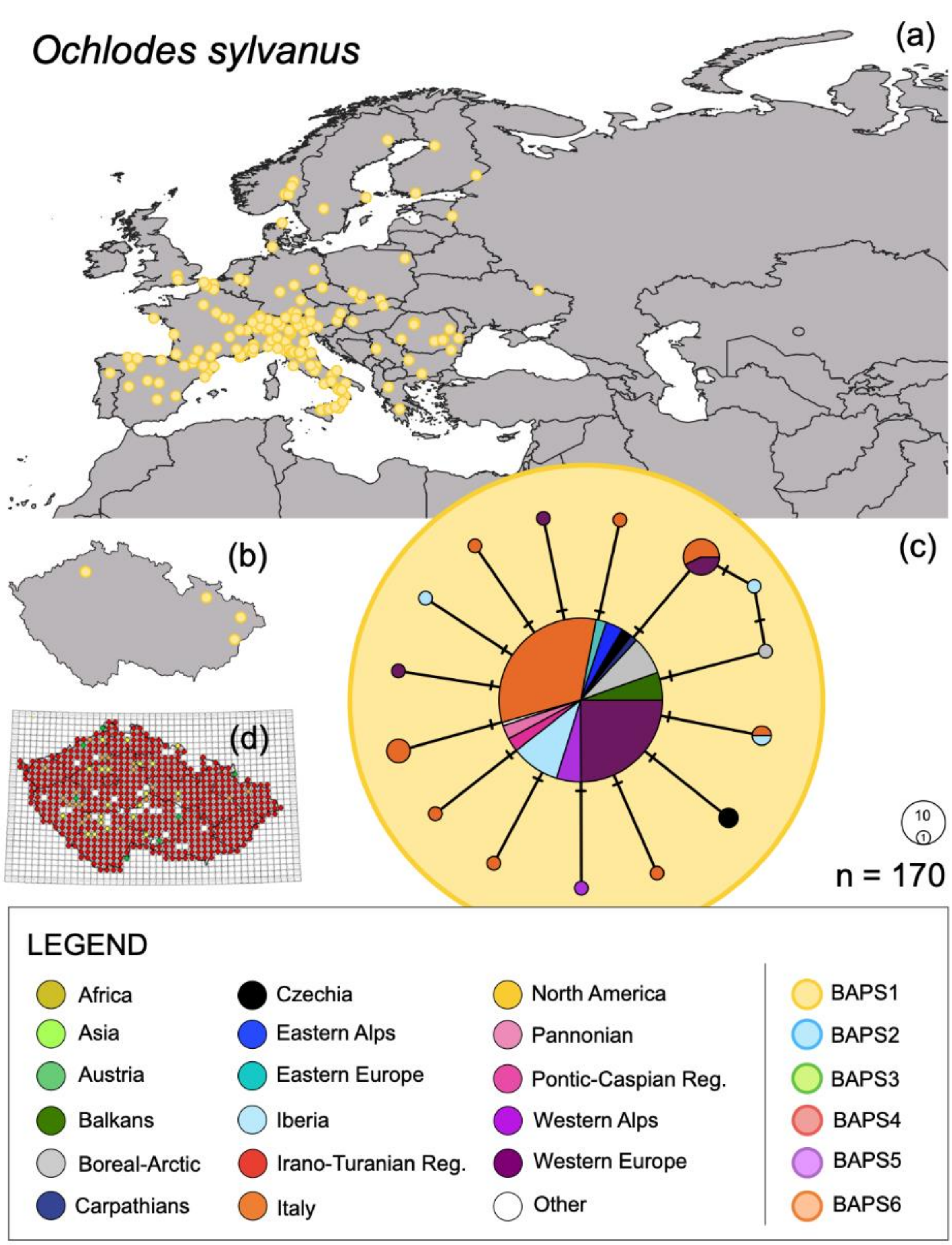


*Ochlodes sylvanus* (Esper, 1777): **Habitat:** ubiquitous. **Red List:** LC

**BAPS:** BAPS-low. **HAP:** HAP-wide-expand

**Range:** Palearctic. **Distribution CZ:** diverse habitats. Widespread and common.

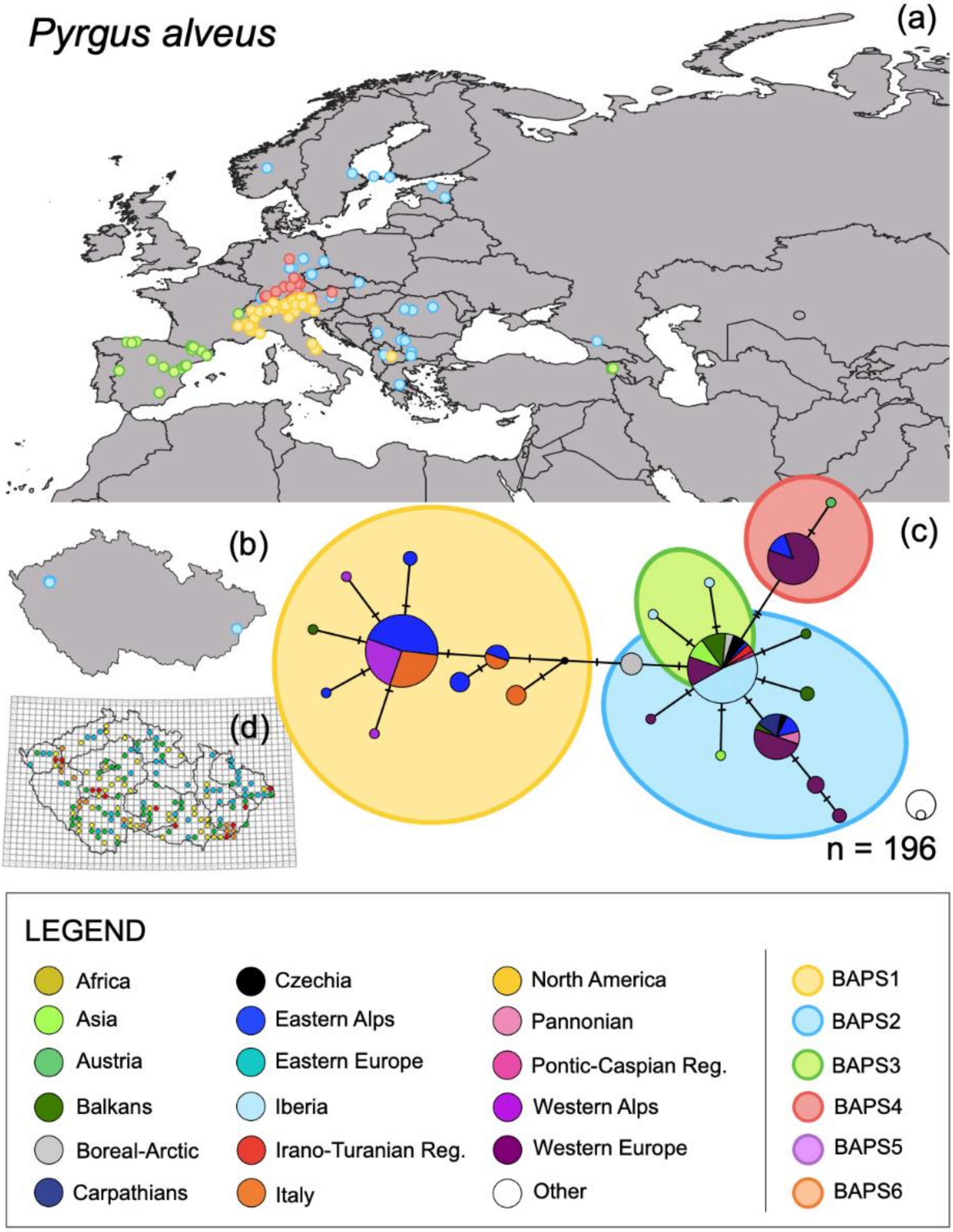


*Pyrgus alveus* (Hübner, 1803): **Habitat:** mesophilic 2 + xerothermophilic 2, **Red List:** CR
**BAPS:** BAPS-East-Balkans. **HAP:** HAP-wide-complex
**Range:** Palearctic. **Distribution CZ:** dry short flowery grasslands. Localized and rare, uplands and hill slopes, decreasing.

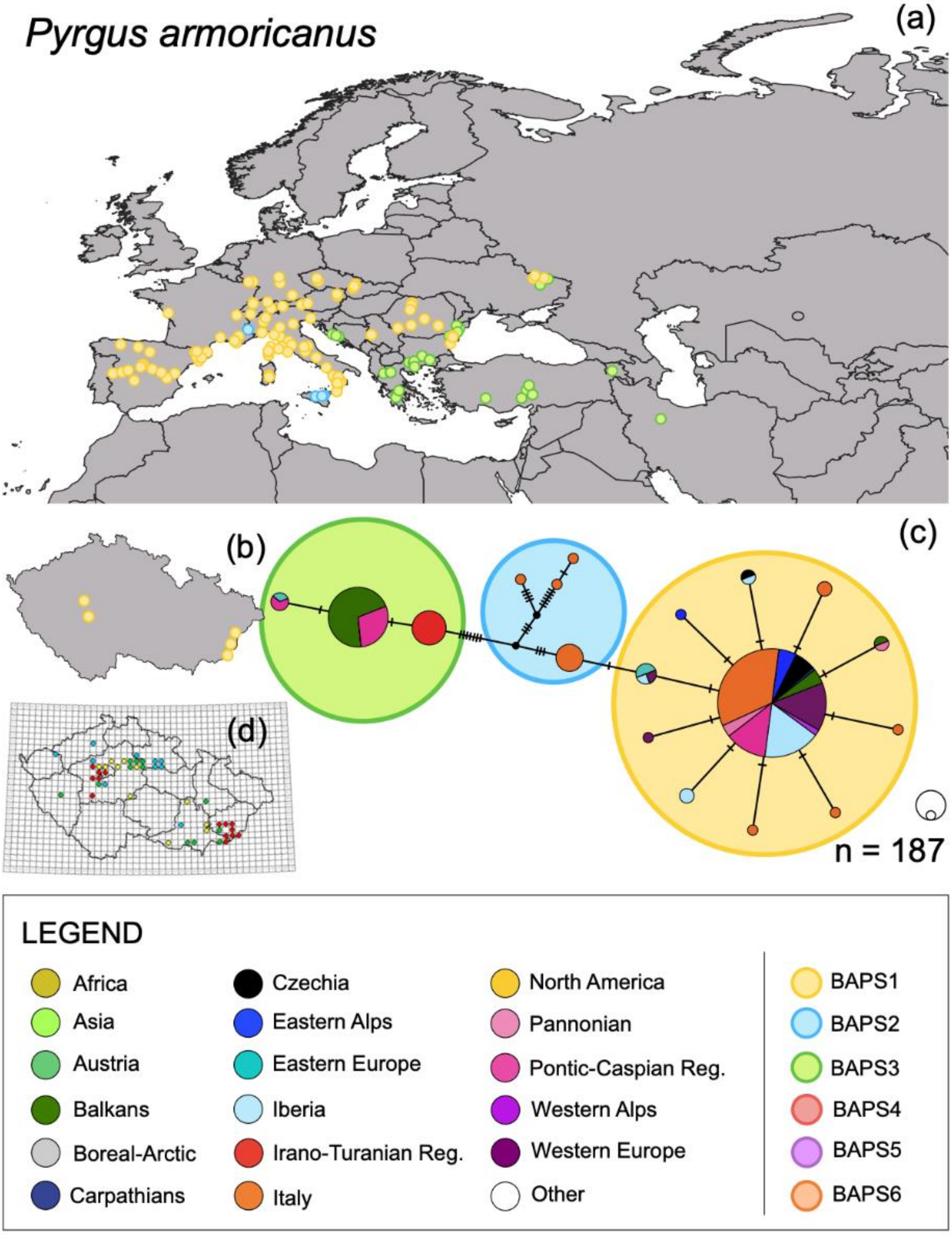


*Pyrgus armoricanus* (Oberthür, 1910): **Habitat:** xerothermophilic 1, **Red List:** CR

**BAPS:** BAPS-West. **HAP:** HAP-wide-expand

**Range:** Western Palearctic. **Distribution CZ:** warm dry sunny flowery grasslands. Rare, decreased in a past, but currently expanding. Currently Central Bohemia, White Carpathians and Vizovice Highlands (part of Carpathian arc).

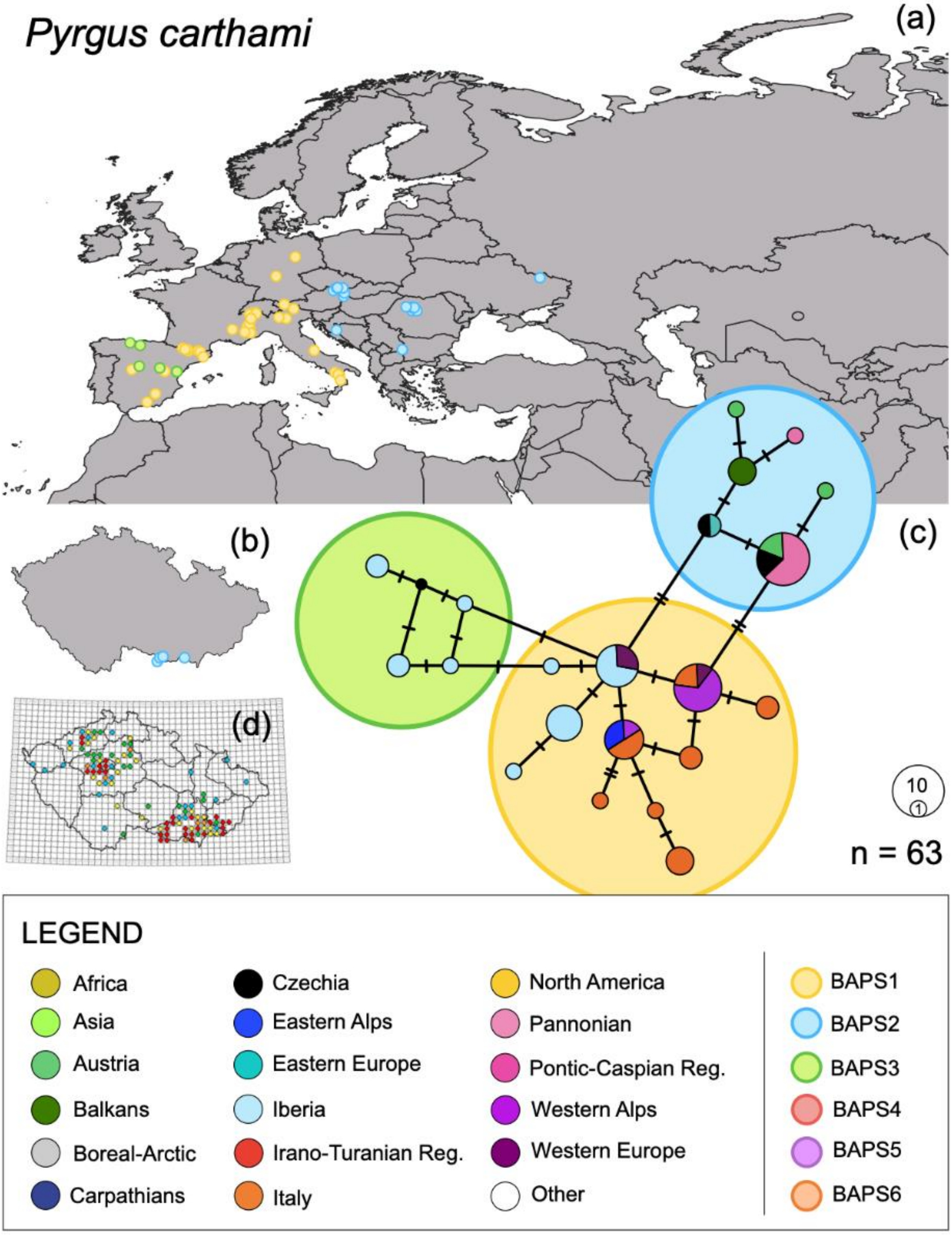


*Pyrgus carthami* (Hübner, 1813): **Habitat:** xerothermophilic 1, **Red List:** VU

**BAPS:** BAPS-East-Balkans. **HAP:** HAP-East-Balkans

**Range:** Western Palearctic. **Distribution CZ:** rocky steppes in warm regions, rare. Currently Central Bohemia, South Bohemia, South and South-East Moravia.

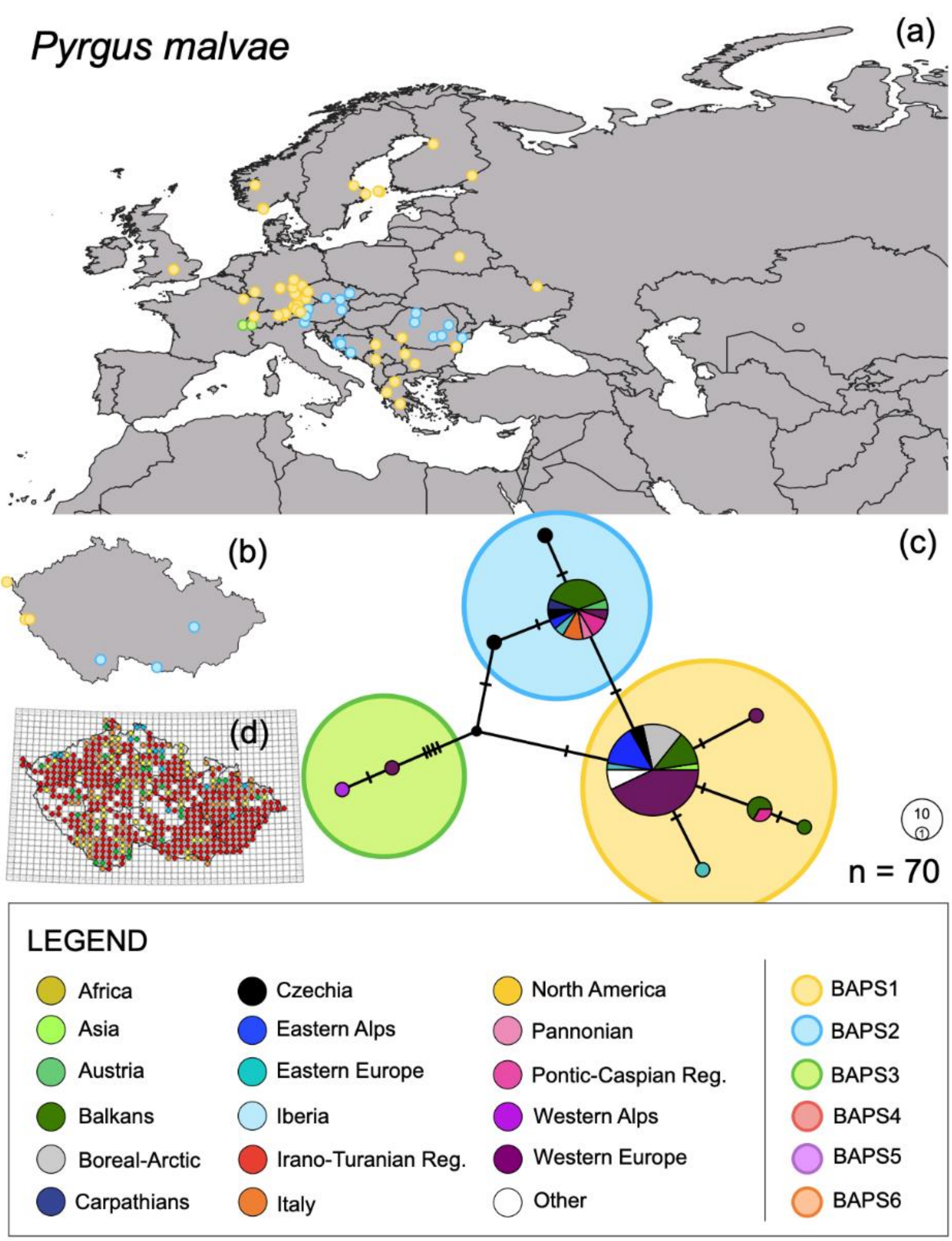


*Pyrgus malvae* (Linnaeus, 1758): **Habitat:** mesophilic 2. **Red List:** LC

**BAPS:** BAPS-East-Balkans + BAPS-Central. **HAP:** HAP-East-Balkans + HAP-Central

**Range:** Palearctic. **Distribution CZ:** flowery meadows, woodland clearings and sunny margins. Widespread, retreated from areas with dense woodlands.

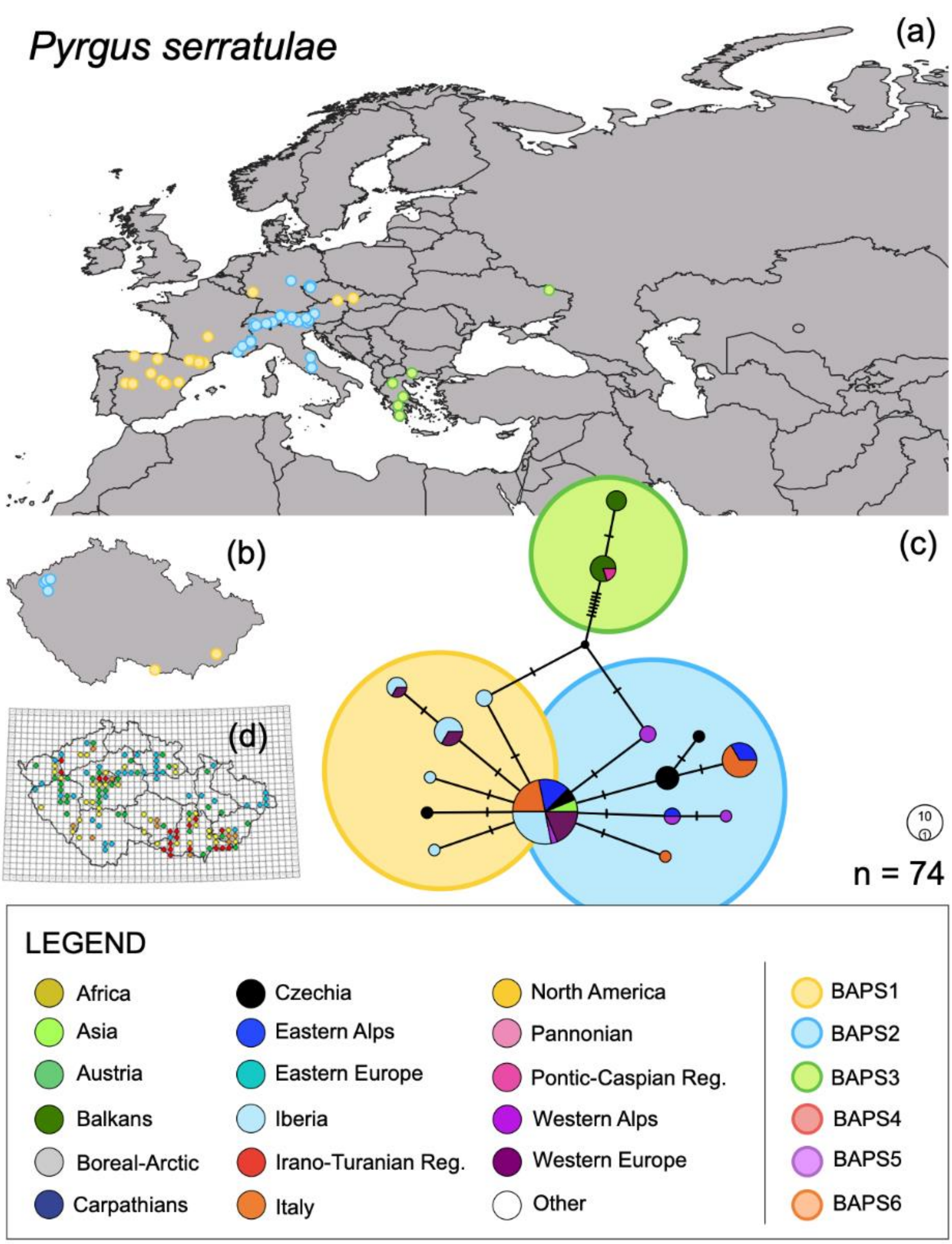


*Pyrgus serratulae* (Rambur, 1839): **Habitat:** xerothermophilic 1. **Red List:** EN

**BAPS:** BAPS-West + BAPS-Italy-Balkans. **HAP:** HAP-wide-complex

**Range:** Euro-Siberian. **Distribution CZ:** rocky dry grasslands and sand lands. Decreasing and localized.

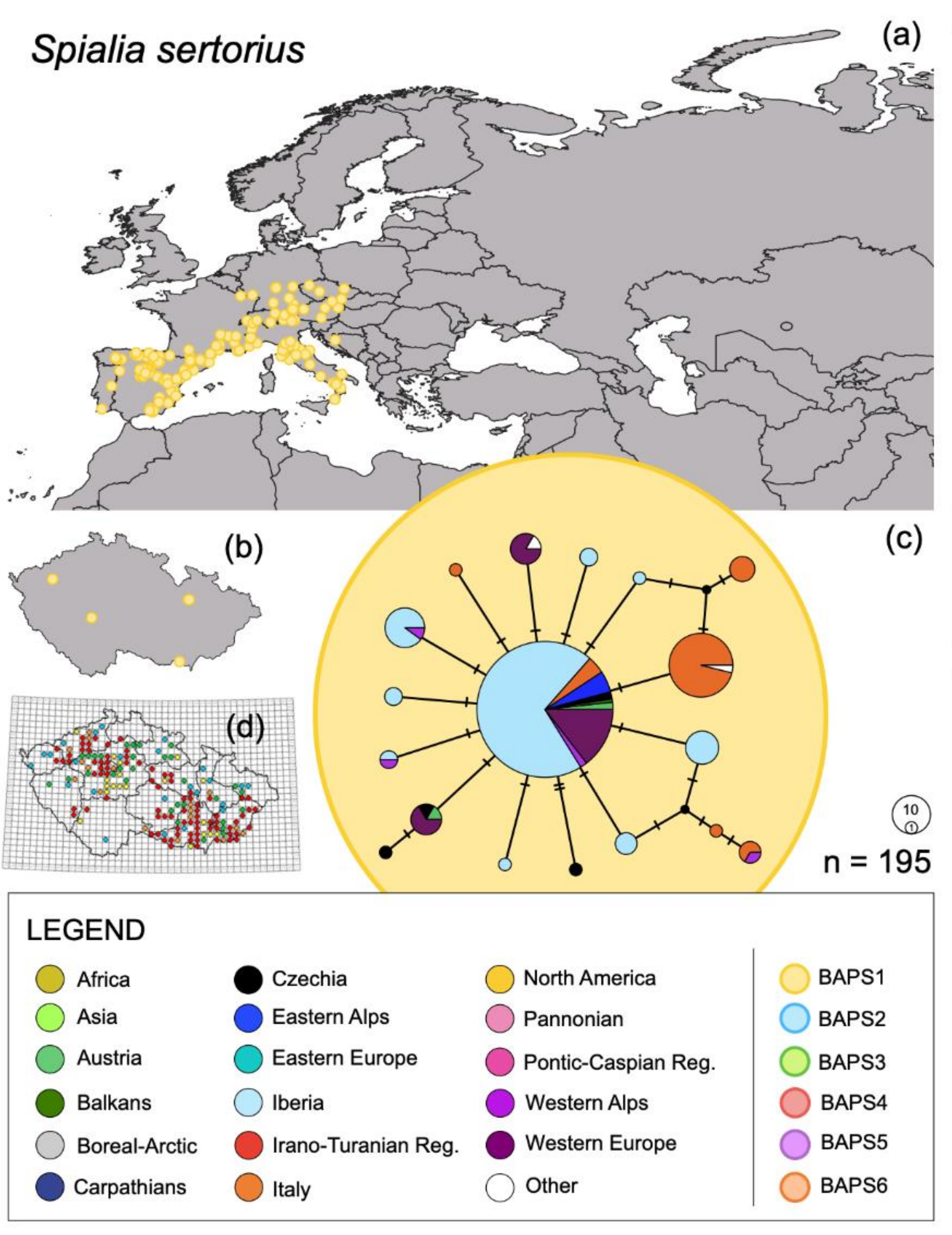


*Spialia sertorius* (Hoffmansegg, 1804): **Habitat:** xerothermophilic 1. **Red List:** VU

**BAPS:** BAPS-low. **HAP:** HAP-wide-complex

**Range:** Atlantic-Mediterranean. **Distribution CZ:** rocky steppes, dry short grasslands. Warm lowlands and uplands in thermophilic regions, localized. Decreased with loss of traditional land use.

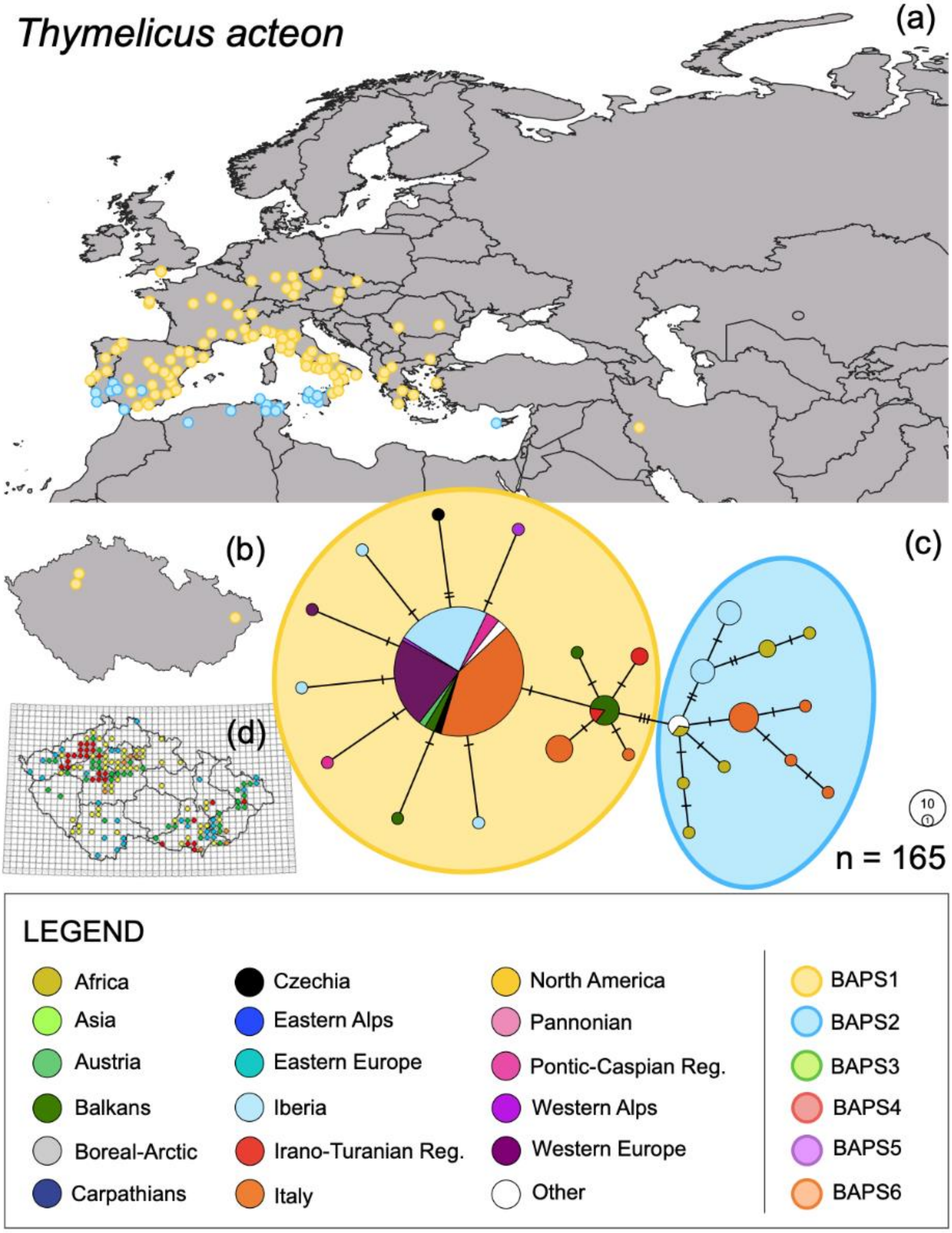


*Thymelicus acteon* (Rottemburg, 1775): **Habitat:** xerothermophilic 2, **Red List:** EN

**BAPS:** BAPS-wide. **HAP:** HAP-wide-expand

**Range:** Mediterranean. **Distribution CZ:** forest steppes and dry scrublands in warm regions, decreased. Thermophilic regions of Bohemia, South Moravia, and two isolated localities in the east (Olomouc and Moravian-Silesian Regions).

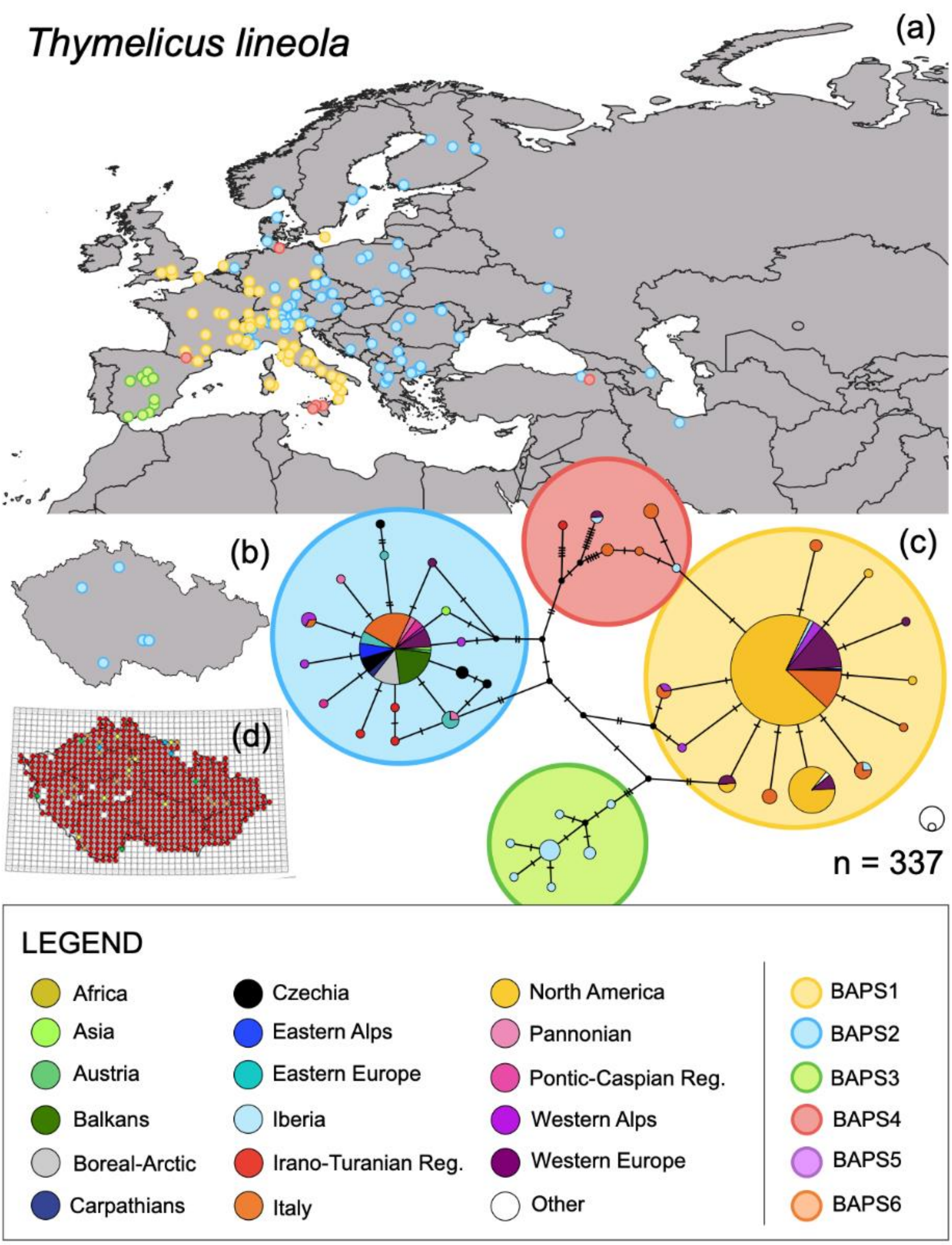


*Thymelicus lineola* (Ochsenheimer, 1808): **Habitat:** mesophilic 1. **Red List:** LC

**BAPS:** BAPS-East-Balkans. **HAP:** HAP-East-Balkans

**Range:** Palearctic, invasively in North America. **Distribution CZ:** grasslands. Widespread and common.

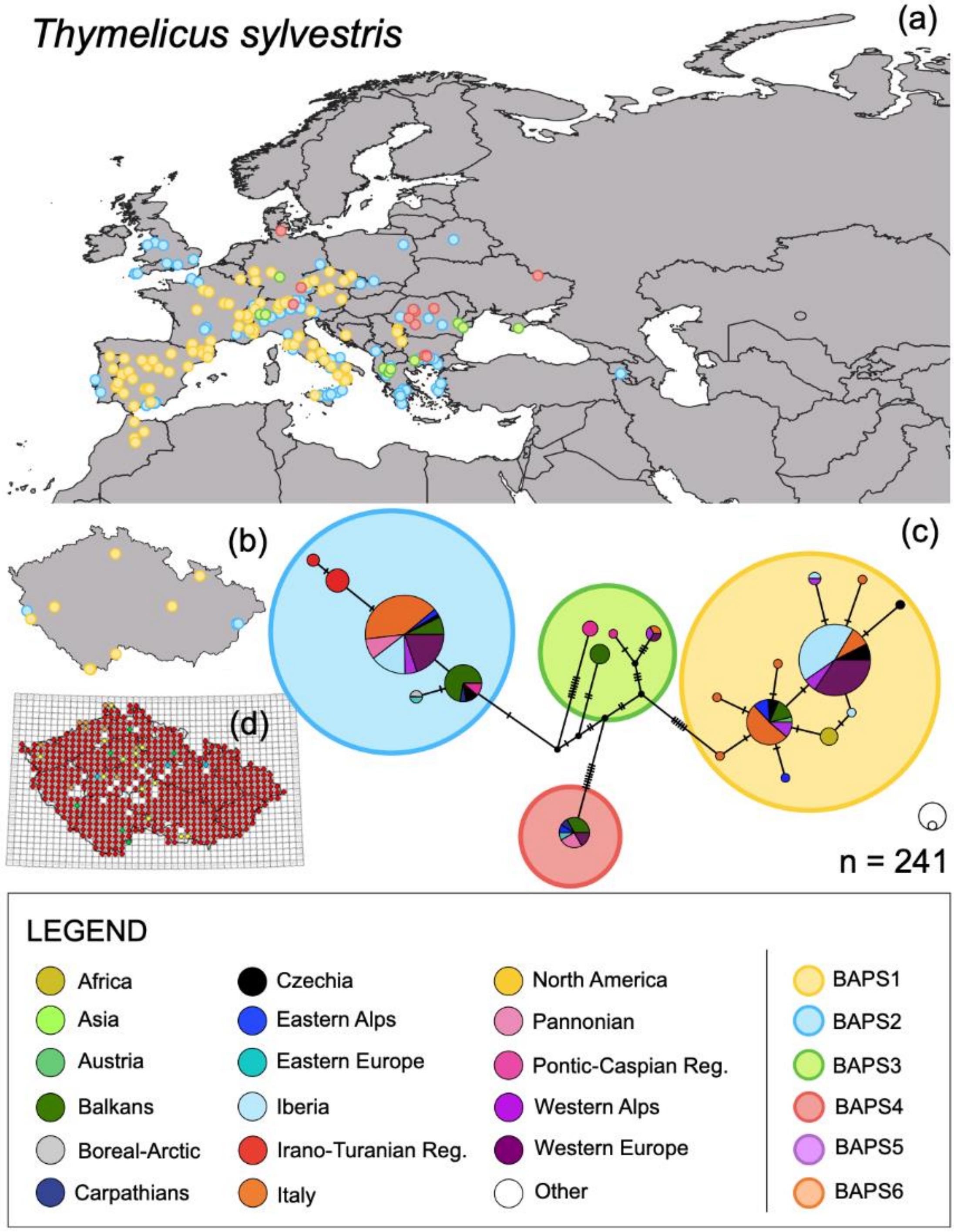


*Thymelicus sylvestris* (Poda, 1761): **Habitat:** mesophilic 2. **Red List:** LC

**BAPS:** BAPS-mix. **HAP:** HAP-mix

**Range:** Western Palearctic. **Distribution CZ:** abandoned meadows, scrublands, woodland clearings and margins. Widespread and common, avoids areas with intensive agriculture. The observed deeply-structured and mixed mitochondrial lineages were not recovered with genome-wide markers, suggesting that mtDNA underwent a different evolutionary history (Hinojosa et al., 2019).

### Pieridae

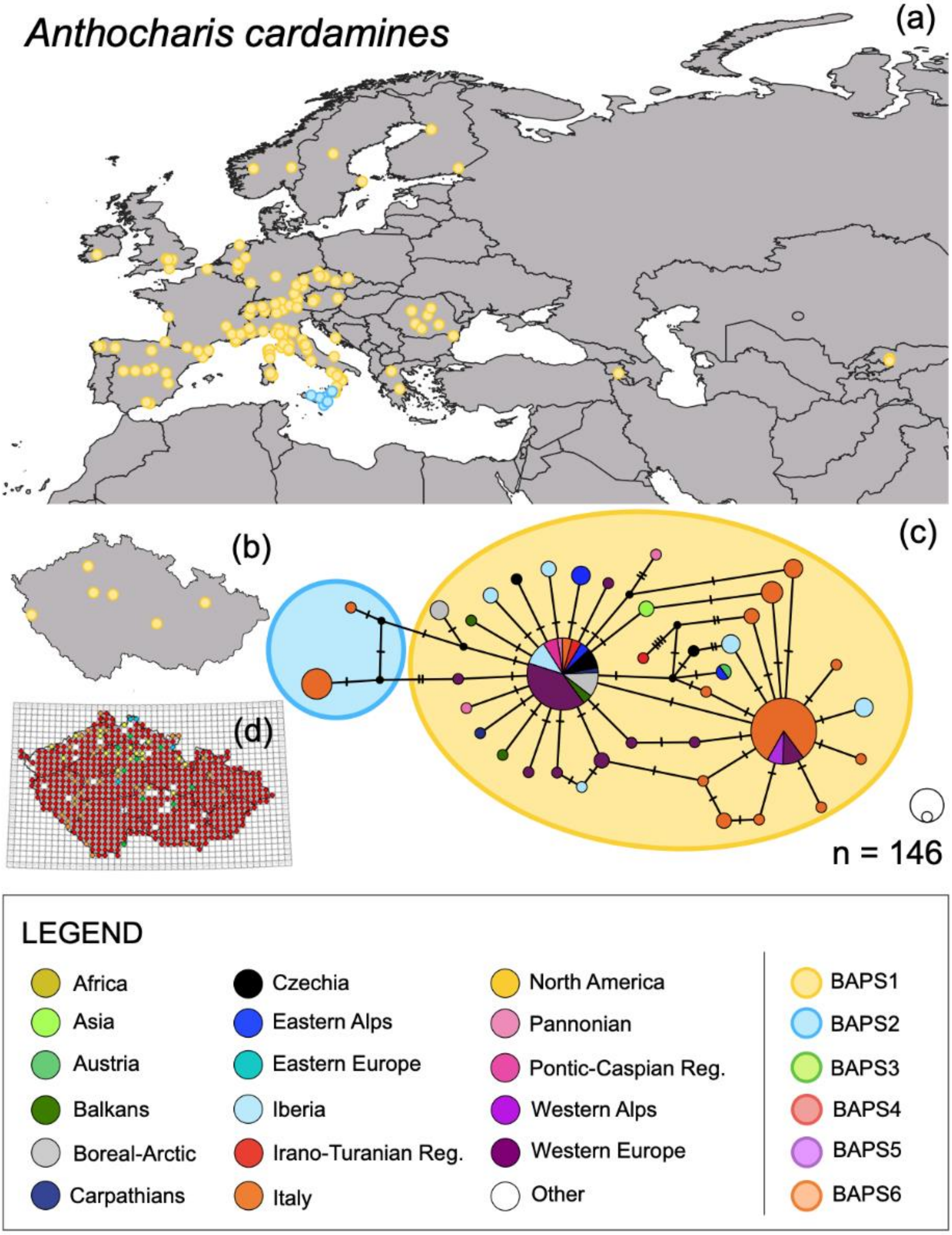


*Anthocharis cardamines* (Linnaeus, 1758): **Habitat:** mesophilic 1, **Red List:** LC

**BAPS:** BAPS-wide. **HAP:** HAP-wide-complex

**Range:** Palearctic. **Distribution CZ:** wet meadows, river banks, woodland margins. Widespread and common.

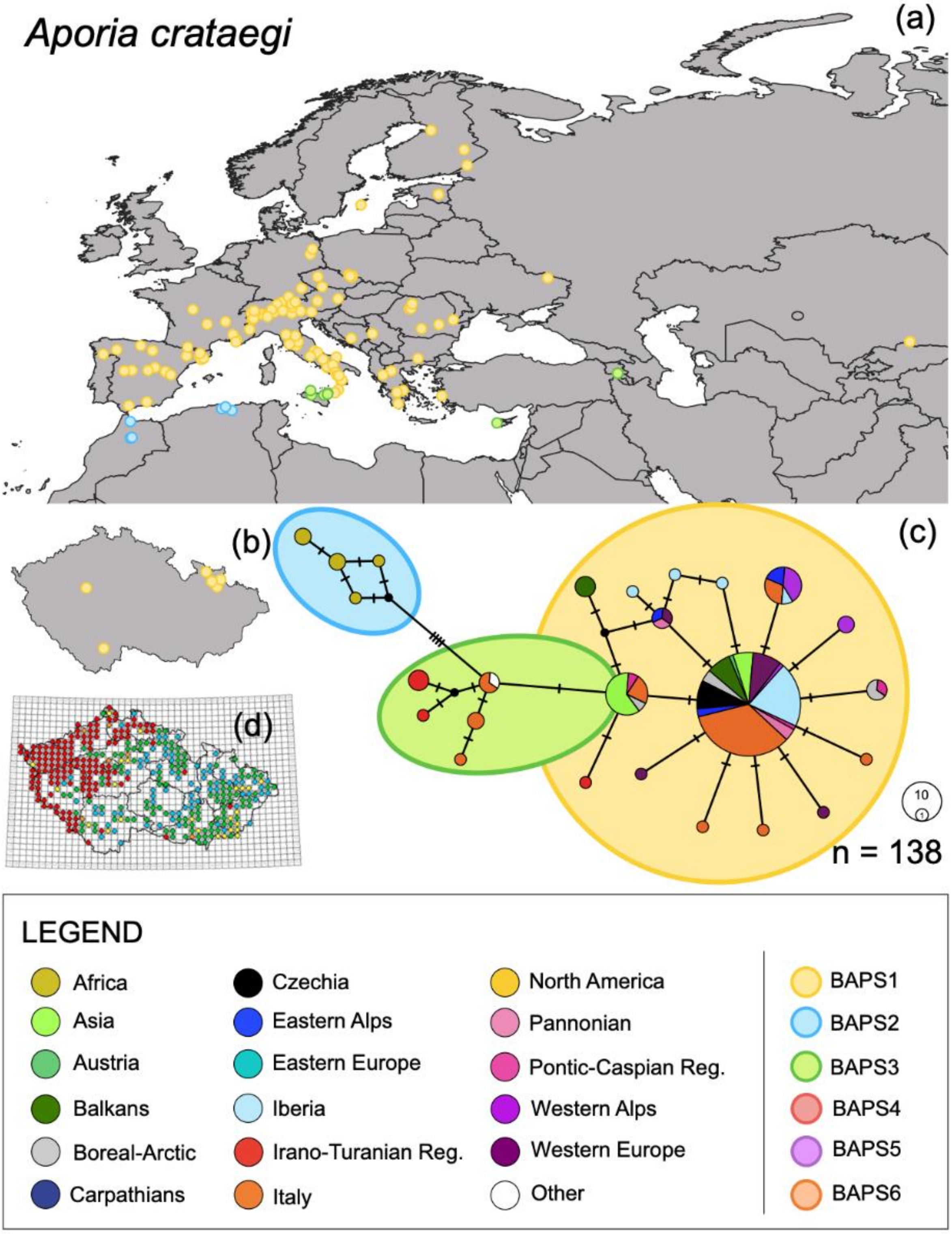


*Aporia crataegi* (Linnaeus, 1758): **Habitat:** mesophilic 2 + xerothermophilic 2, **Red List:** LC
**BAPS:** BAPS-wide. **HAP:** HAP-wide-complex

**Range:** Palearctic. **Distribution CZ:** forest steppes and scrublands. Widespread in past and missing since 1990s. Now expanding from the west (Z. Fric, Hula, Konvička, & Pavlíčko, 2000), reaching Bohemian-Moravian Highlands and High Ash Mts.

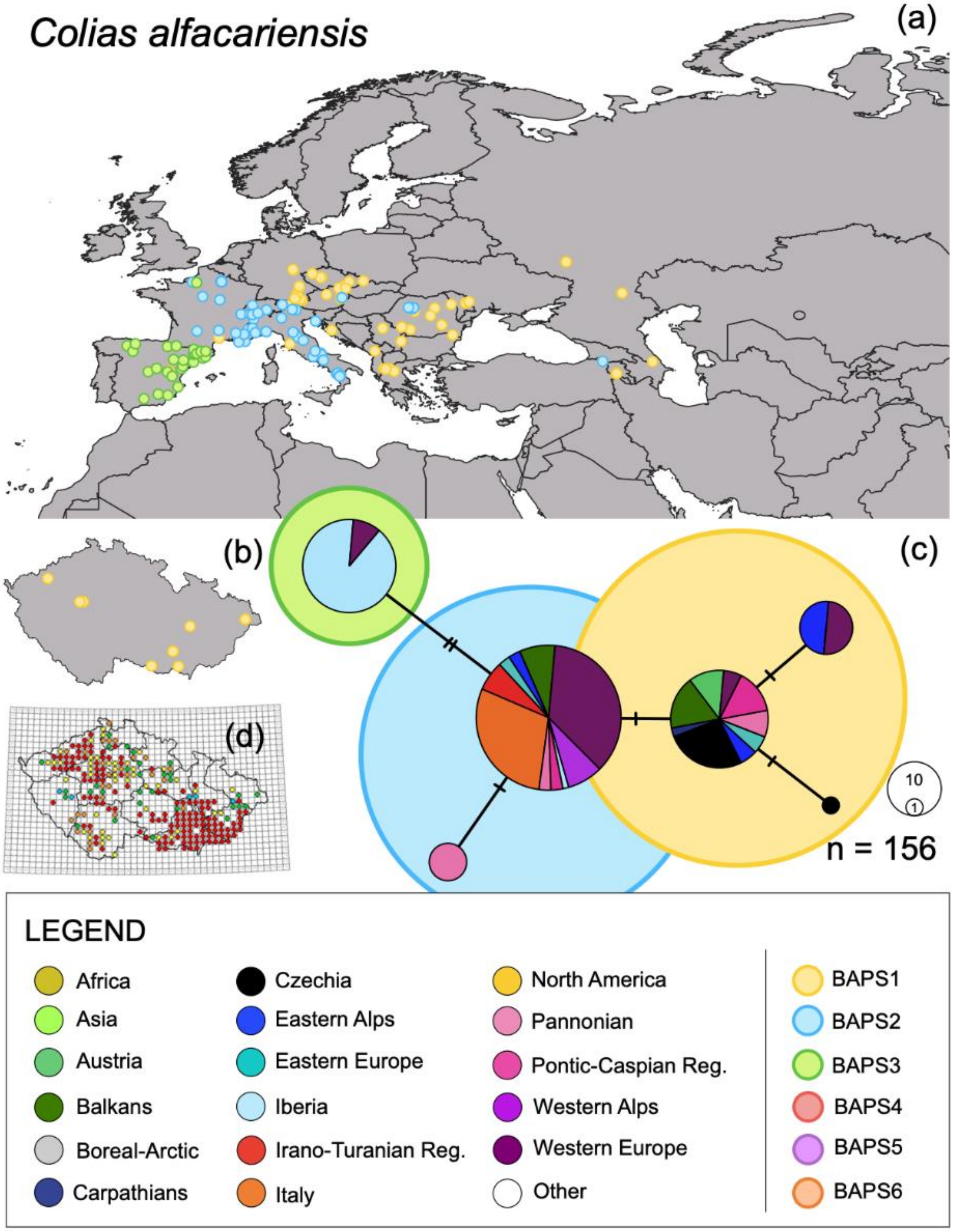


*Colias alfacariensis* Ribbe, 1905: **Habitat:** xerothermophilic 1, **Red List:** VU

**BAPS:** BAPS-East-Balkans. **HAP:** HAP-East-Balkans

**Range:** Western Palearctic. **Distribution CZ:** warm dry grasslands across the country, forest steppes.

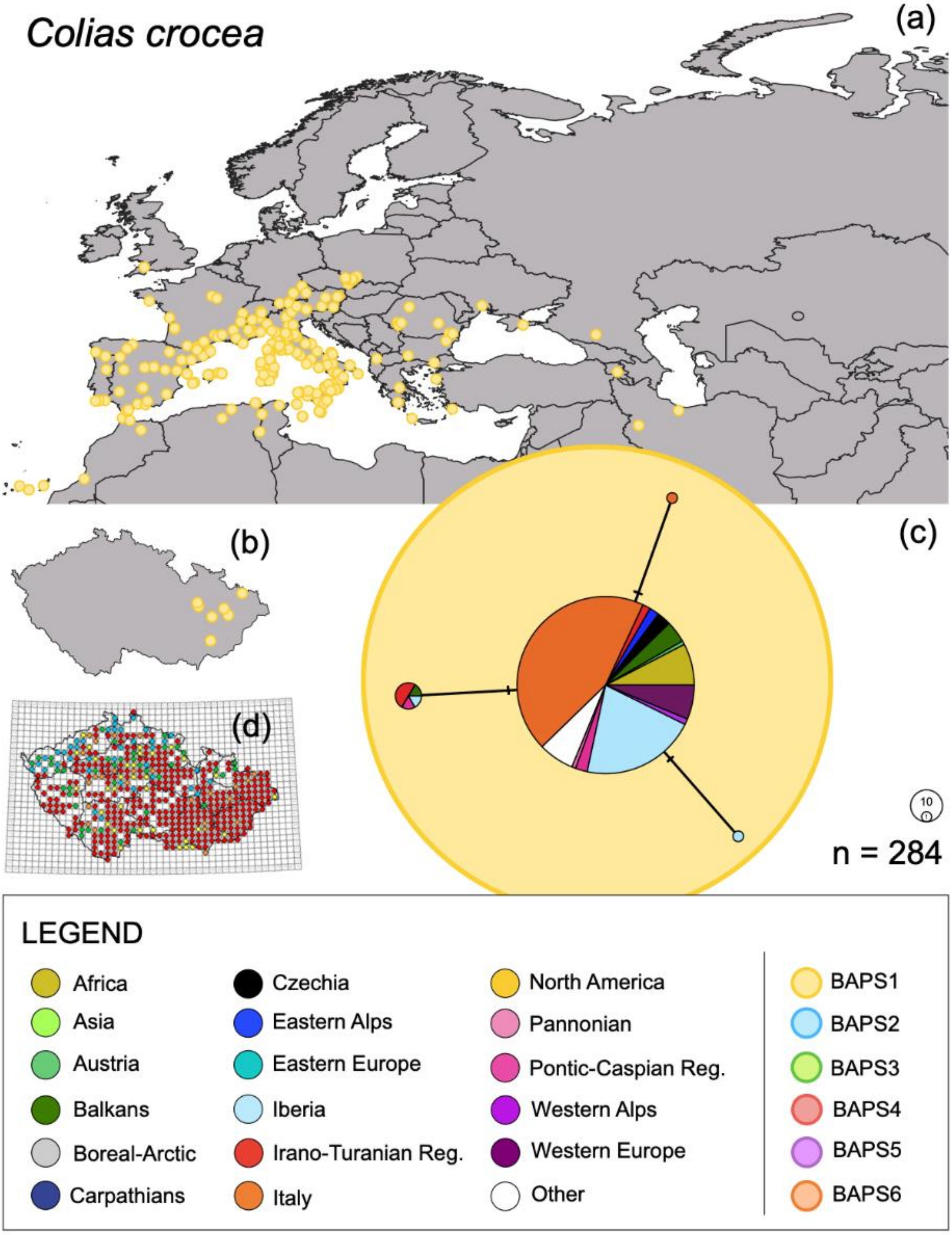


*Colias crocea* (Geoffroy, 1785): **Habitat:** ubiquitous, **Red List:** LC

**BAPS:** BAPS-low. **HAP:** HAP-low

**Range:** Mediterranean. **Distribution CZ:** migrant found on meadows and ruderal sites. More common in Moravia (migration corridor).

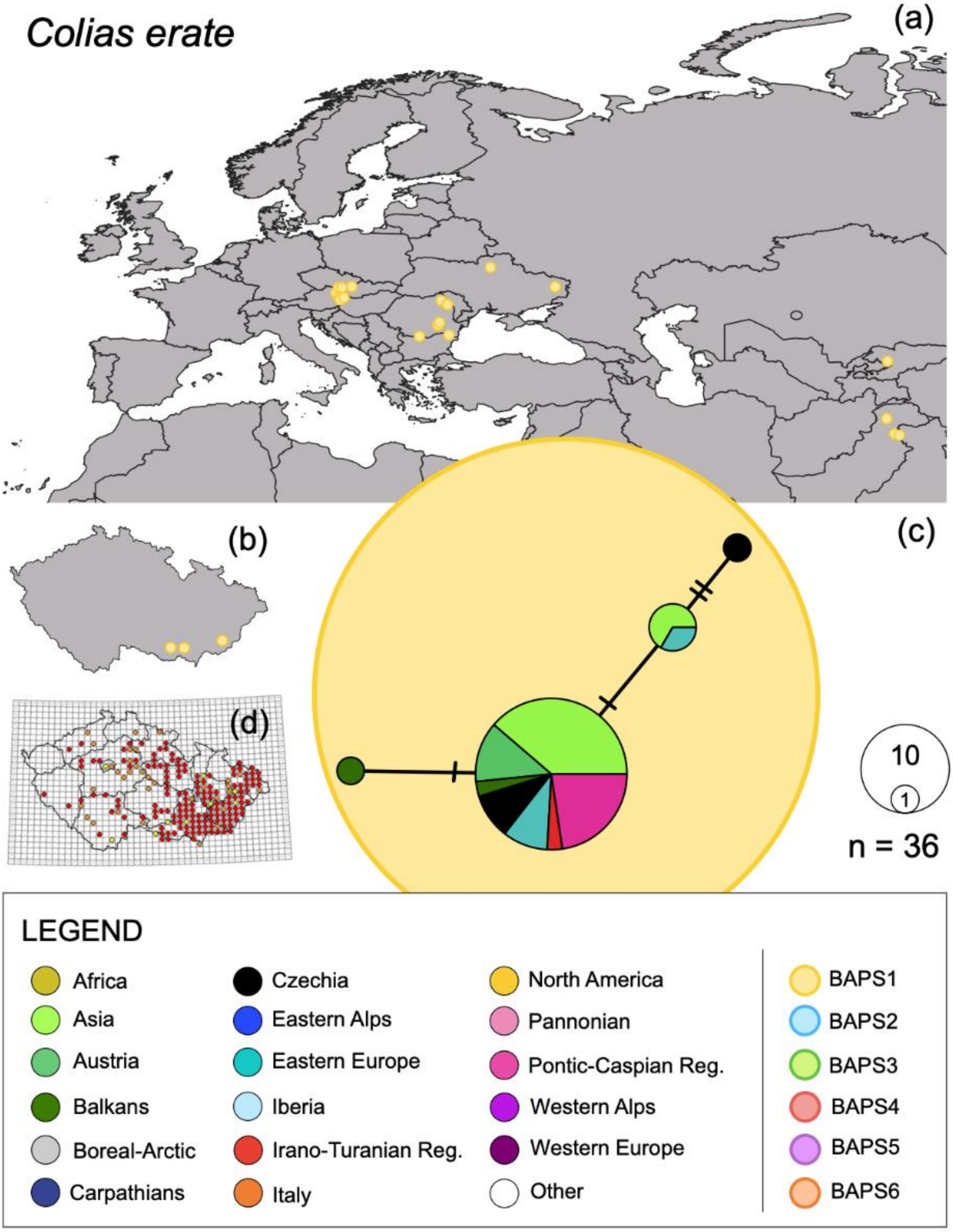


*Colias erate* (Esper, 1805): **Habitat:** ubiquitous, **Red List:** LC

**BAPS:** BAPS-low. **HAP:** HAP-low

**Range:** Palearctic (not in Western Europe). **Distribution CZ:** ruderal places. Migrant, reaching the country since the 1990s, regularly found in Moravia, rarer in Bohemia.

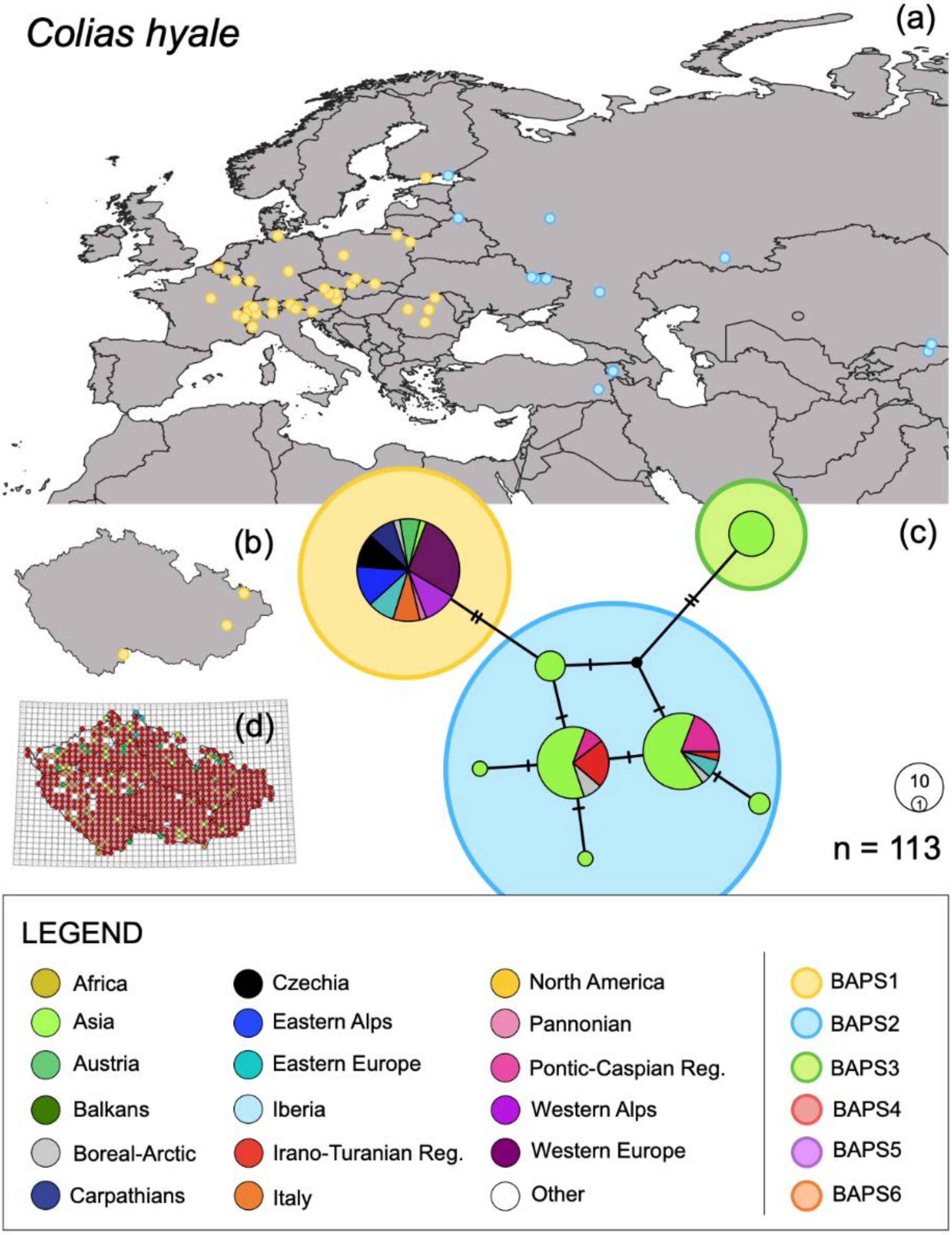


*Colias hyale* (Linnaeus, 1758): **Habitat:** ubiquitous, **Red List:** LC

**BAPS:** BAPS-temperate. **HAP:** HAP-temperate

**Range:** Euro-Siberian (not in Southern Europe). **Distribution CZ:** widespread and common in agricultural landscapes. The phylogeographic patterns correspond to the species distribution, with a suture zone located further eastwards.

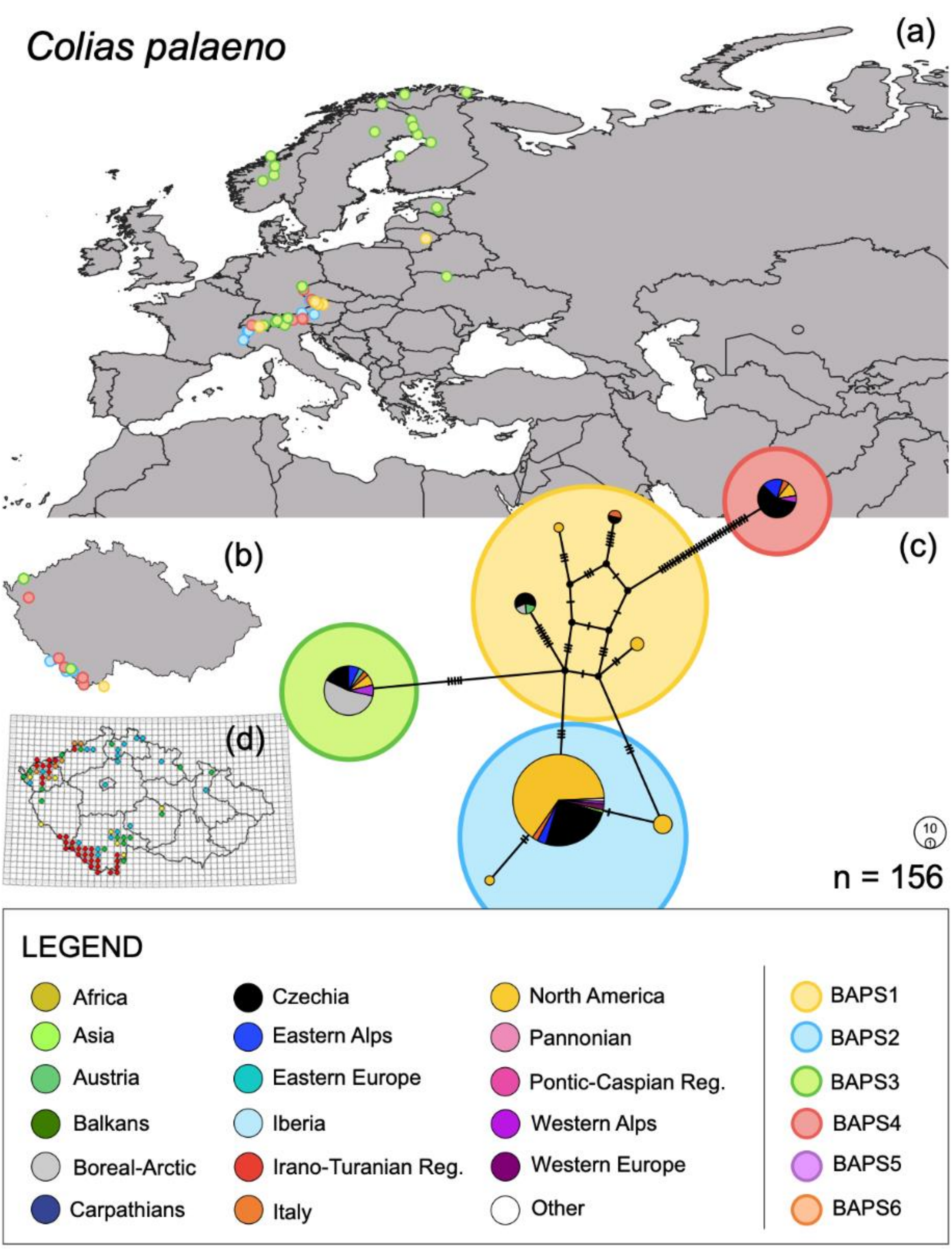


*Colias palaeno* (Linnaeus, 1760): **Habitat:** tyrphophilic, **Red List:** VU

**BAPS:** BAPS-mix. **HAP:** HAP-mix

**Range:** Holarctic. **Distribution CZ:** peat bogs. Currently present only in borderland mountains of Bohemia (south and southwest, west), extinct from Moravia. Threatened by successional changes of bogs.

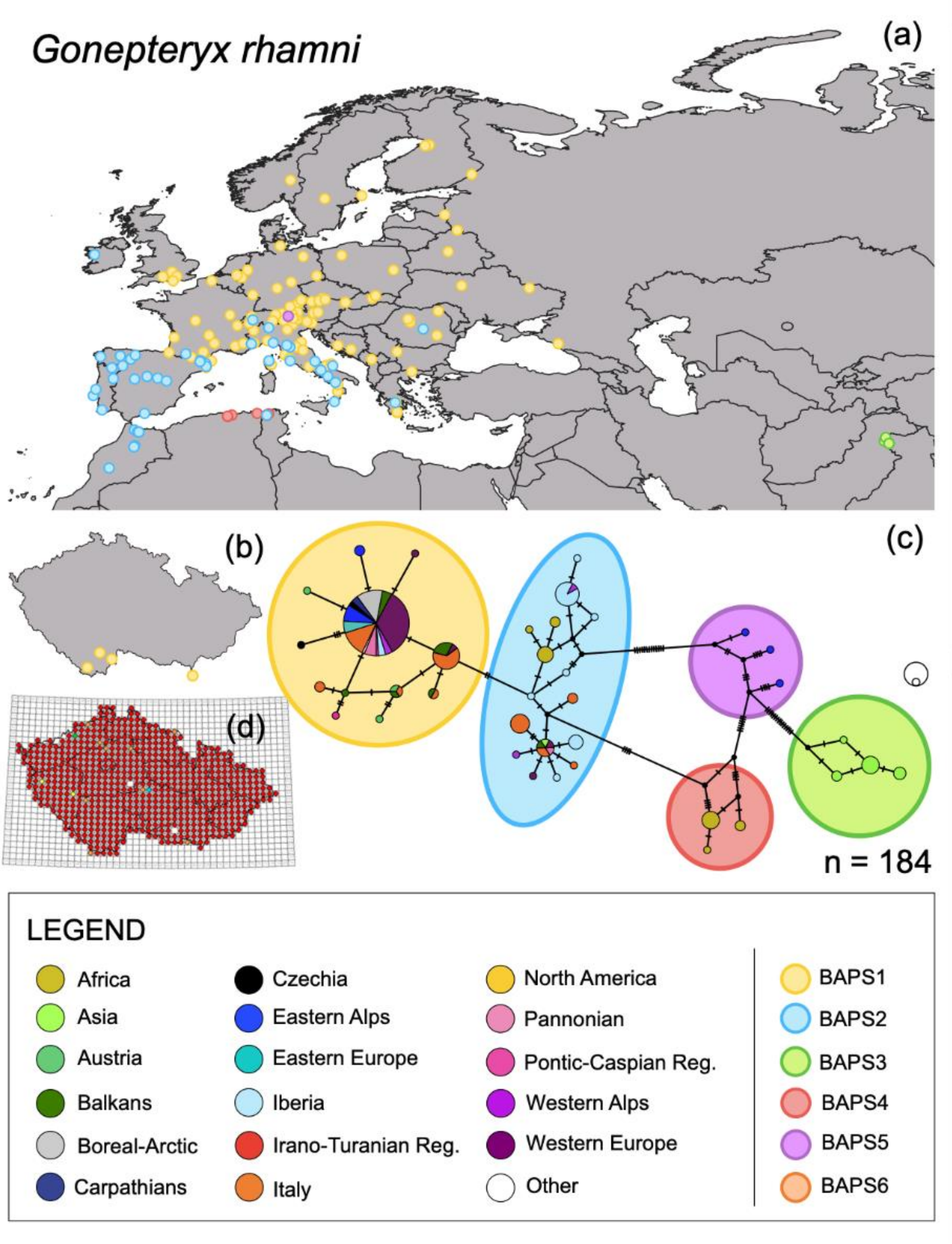


*Gonepteryx rhamni* (Linnaeus, 1758): **Habitat:** mesophilic 2, **Red List:** LC

**BAPS:** BAPS-wide. **HAP:** HAP-wide-complex

**Range:** Western Palearctic. **Distribution CZ:** woodland margins and clearings, woodland meadows. Widespread and common.

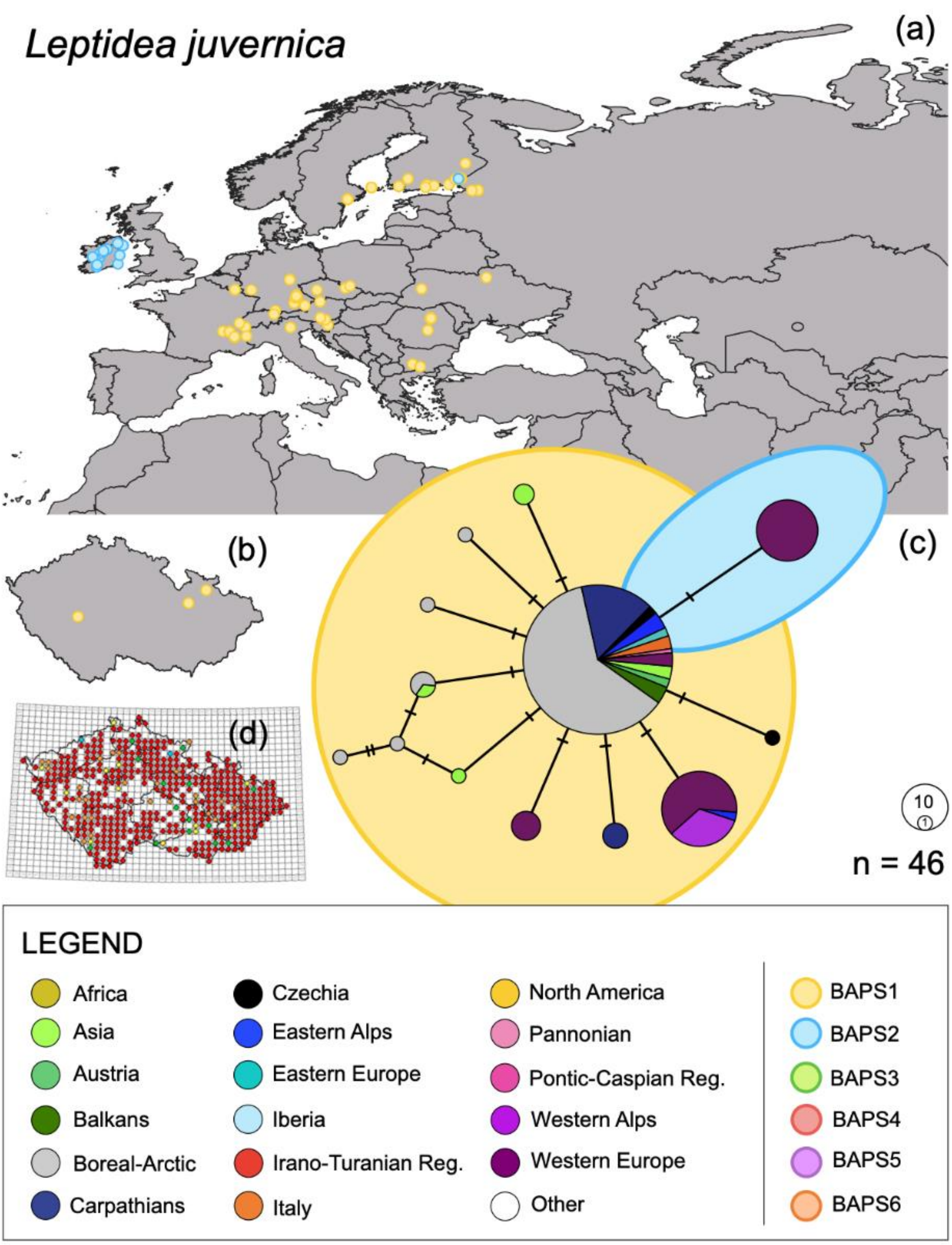


*Leptidea juvernica* Williams, 1946: **Habitat:** mesophilic 2 + hydrophilic, **Red List:** LC

**BAPS:** BAPS-temperate. **HAP:** HAP-wide-complex

**Range:** Western Palearctic. **Distribution CZ:** meadows in alluvia, river banks, wet flowery meadows. Widespread across the country, retreated from agricultural landscapes.

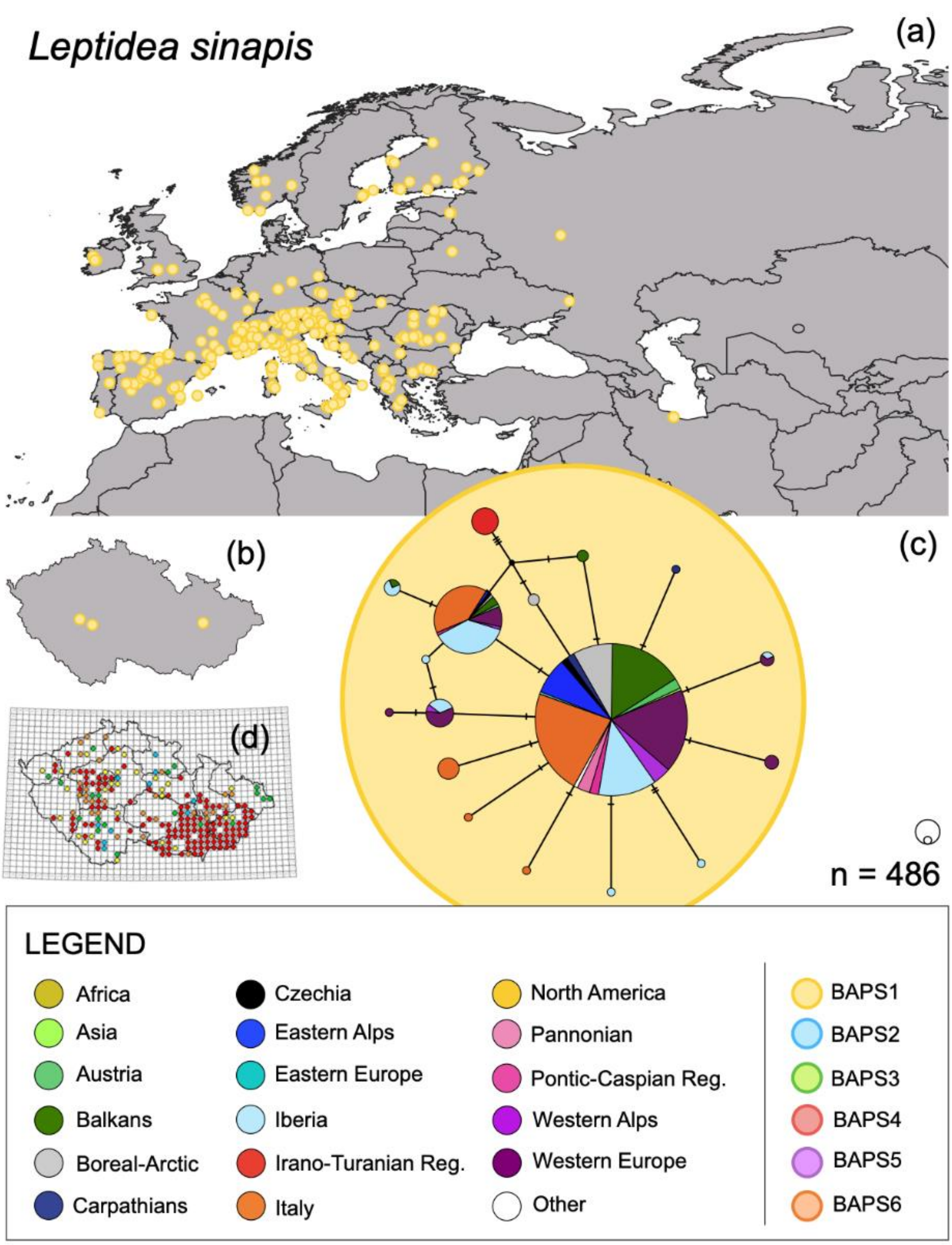


*Leptidea sinapis* (Linnaeus, 1758): **Habitat:** mesophilic 2 + xerothermophilic 2, **Red List:** NT

**BAPS:** BAPS-low. **HAP:** HAP-wide-complex

**Range:** Western Palearctic. **Distribution CZ:** dry scrublands and forest steppes. Present in warm regions across the country, localized.

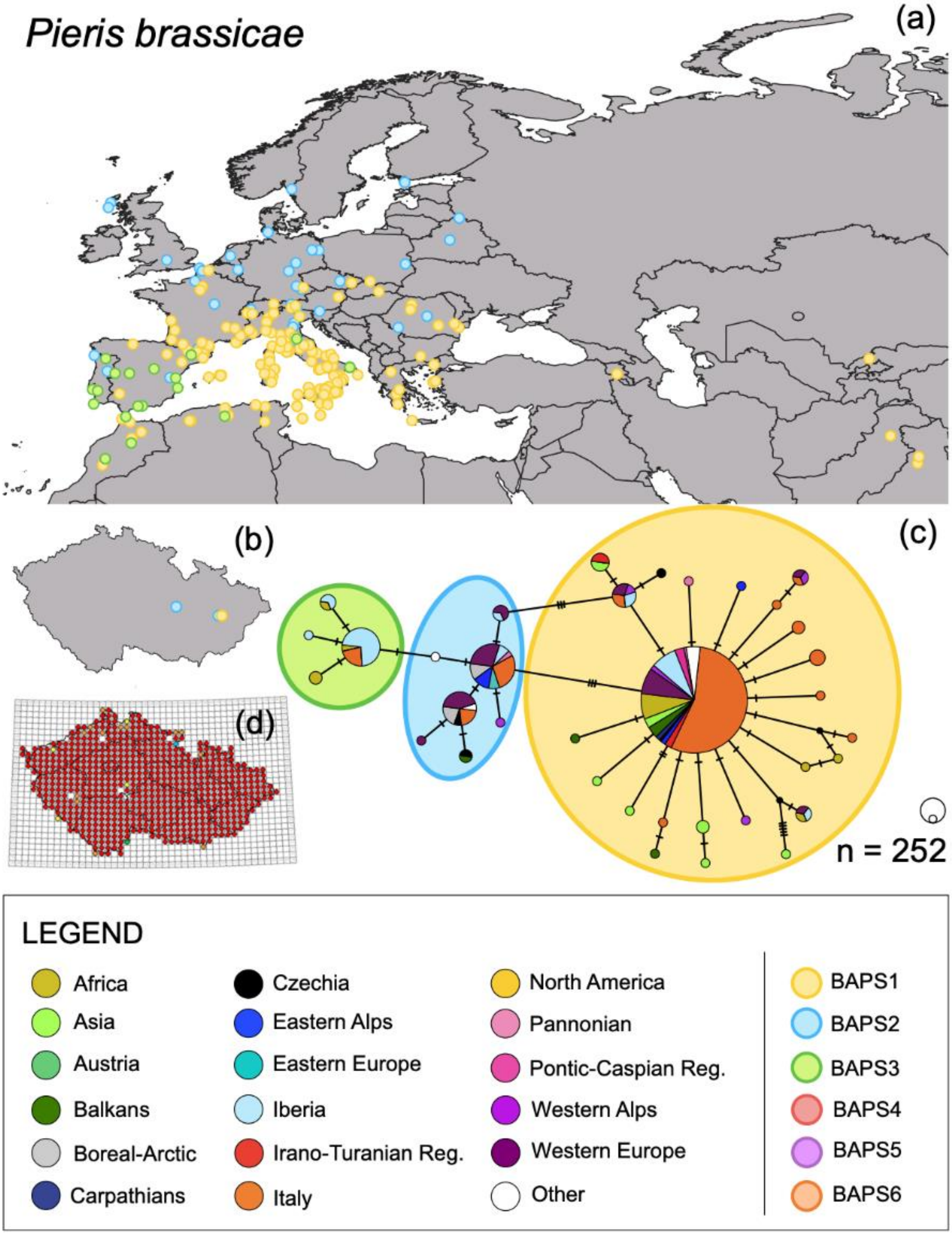


*Pieris brassicae* (Linnaeus, 1758): **Habitat:** ubiquitous, **Red List:** LC

**BAPS:** BAPS-mix. **HAP:** HAP-mix

**Range:** Western Palearctic, introduced to America and South Africa. **Distribution CZ:** agricultural landscapes. Widespread and common, migratory (in spring northwards, in late summer southwards). Two genetic lineages are widespread across Europe (and in CZ), one more southern (BAPS1) and one more northern (BAPS2).

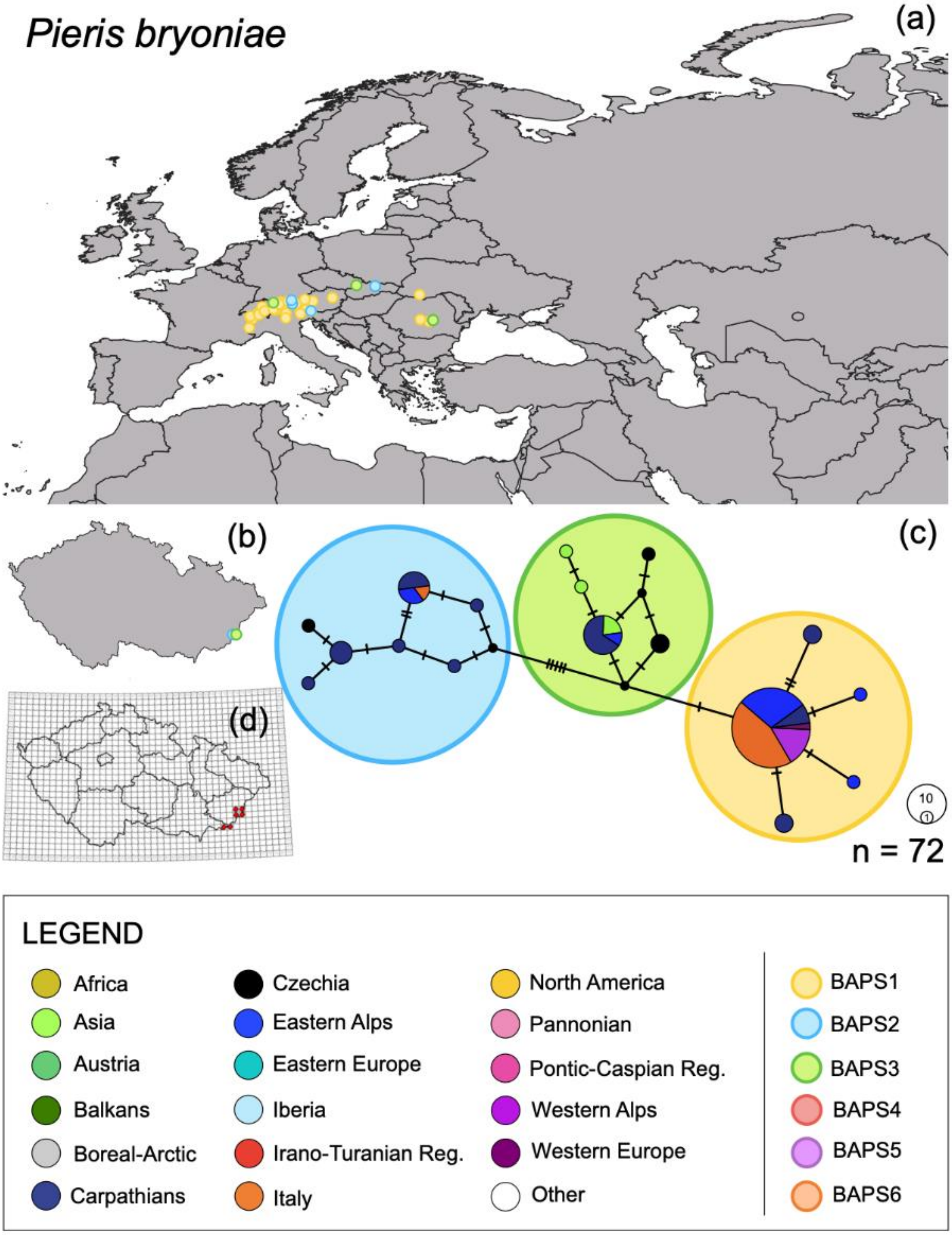


*Pieris bryoniae* (Hübner, 1806): **Habitat:** mesophilic 2+3, **Red List:** NT

**BAPS:** BAPS-Central + BAPS-temperate. **HAP:** HAP-Central + HAP-temperate

**Range:** (probably) Palearctic, mountains. **Distribution CZ:** White Carpathians: mountain meadows, woodland clearings and edges. BAPS3 also found in East Asia (*Pieris bryoniae kamtschadalis* Röber, 1907). *Pieris bryoniae* shares haplotypes with *P. napi*, but Czech *P. bryoniae* samples are distinctive, with unique haplotypes.

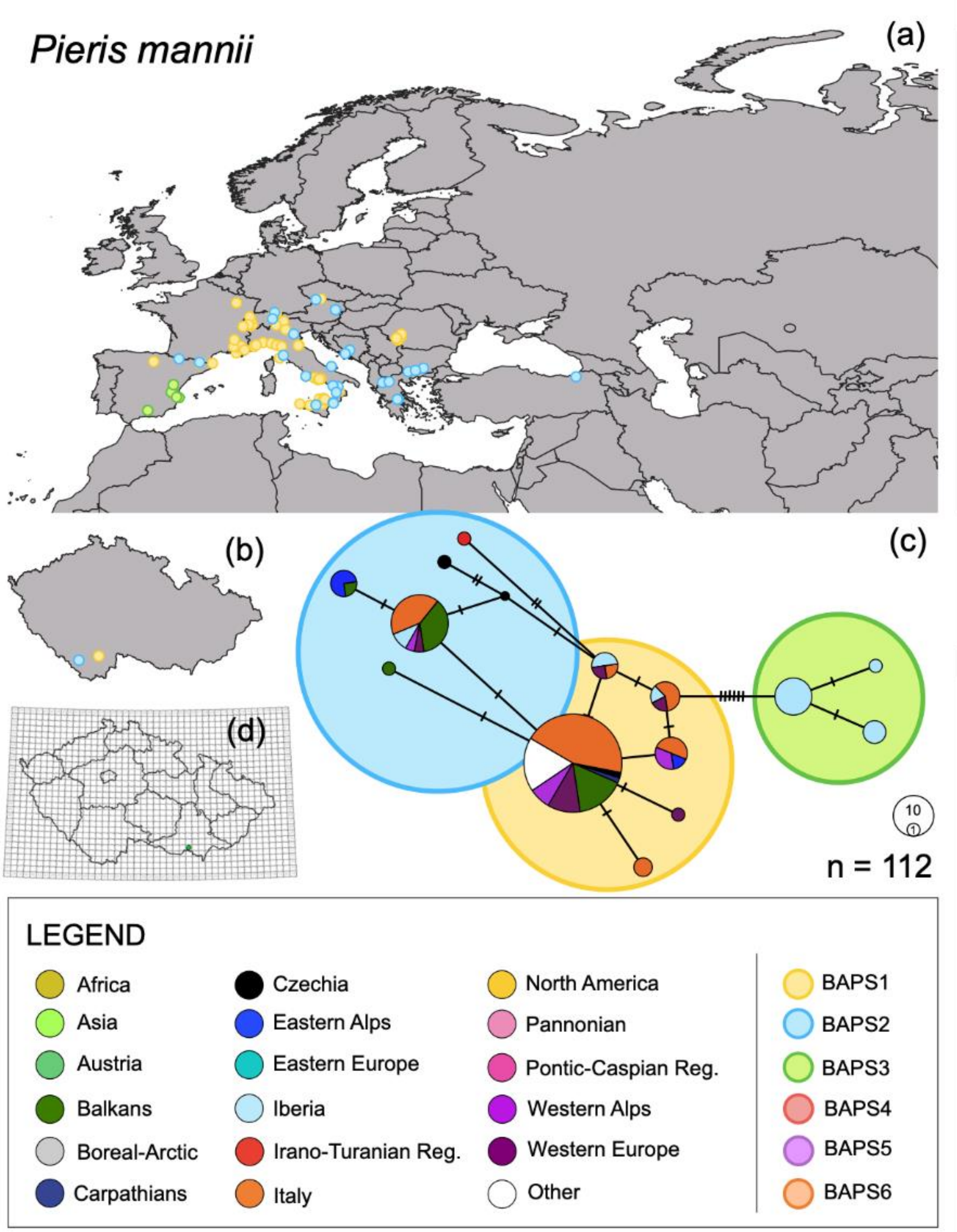


*Pieris mannii* (Mayer, 1851): **Habitat:** xerothermophilic 1, **Red List:** LC

**BAPS:** BAPS-mix. **HAP:** HAP-mix

**Range:** Mediterranean. **Distribution CZ:** rocky dry grasslands and forest steppes, pastures, gardens and parks. Extinct from its only locality in south Moravia in the 1960s. With the warming climate, it is currently expanding across Western Europe. First recent records from southern Bohemia in 2018, afterwards recorded from several other localities across Bohemia (e.g., (Z. F. Fric, 2019).

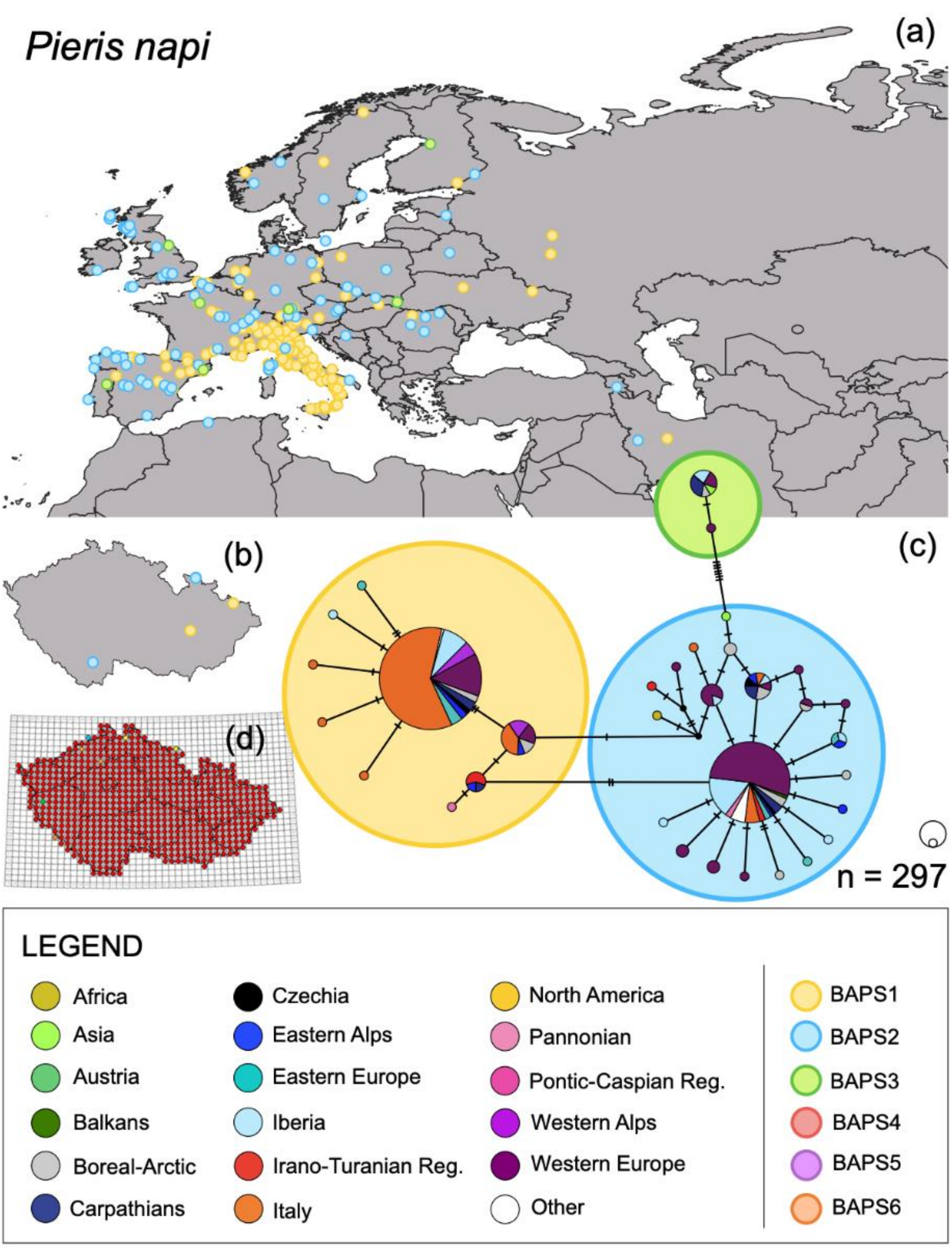


*Pieris napi* (Linnaeus, 1758): **Habitat:** ubiquitous, **Red List:** LC

**BAPS:** BAPS-mix. **HAP:** HAP-mix

**Range:** Palearctic. **Distribution CZ:** diverse habitats, widespread and common.

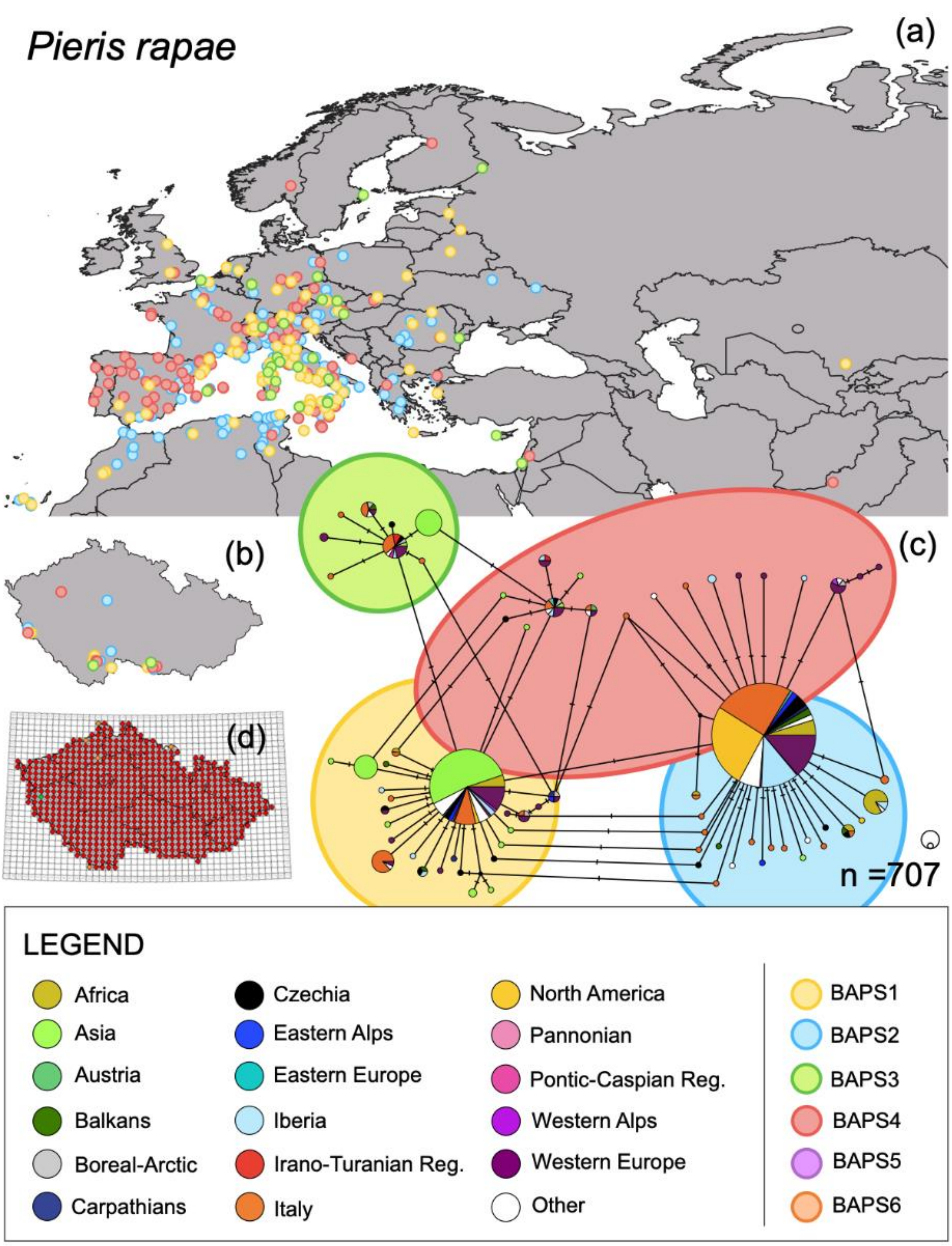


*Pieris rapae* (Linnaeus, 1758): **Habitat:** ubiquitous, **Red List:** LC

**BAPS:** BAPS-mix. **HAP:** HAP-mix

**Range:** Cosmopolite. **Distribution CZ:** diverse habitats, common even on farmlands, widespread.

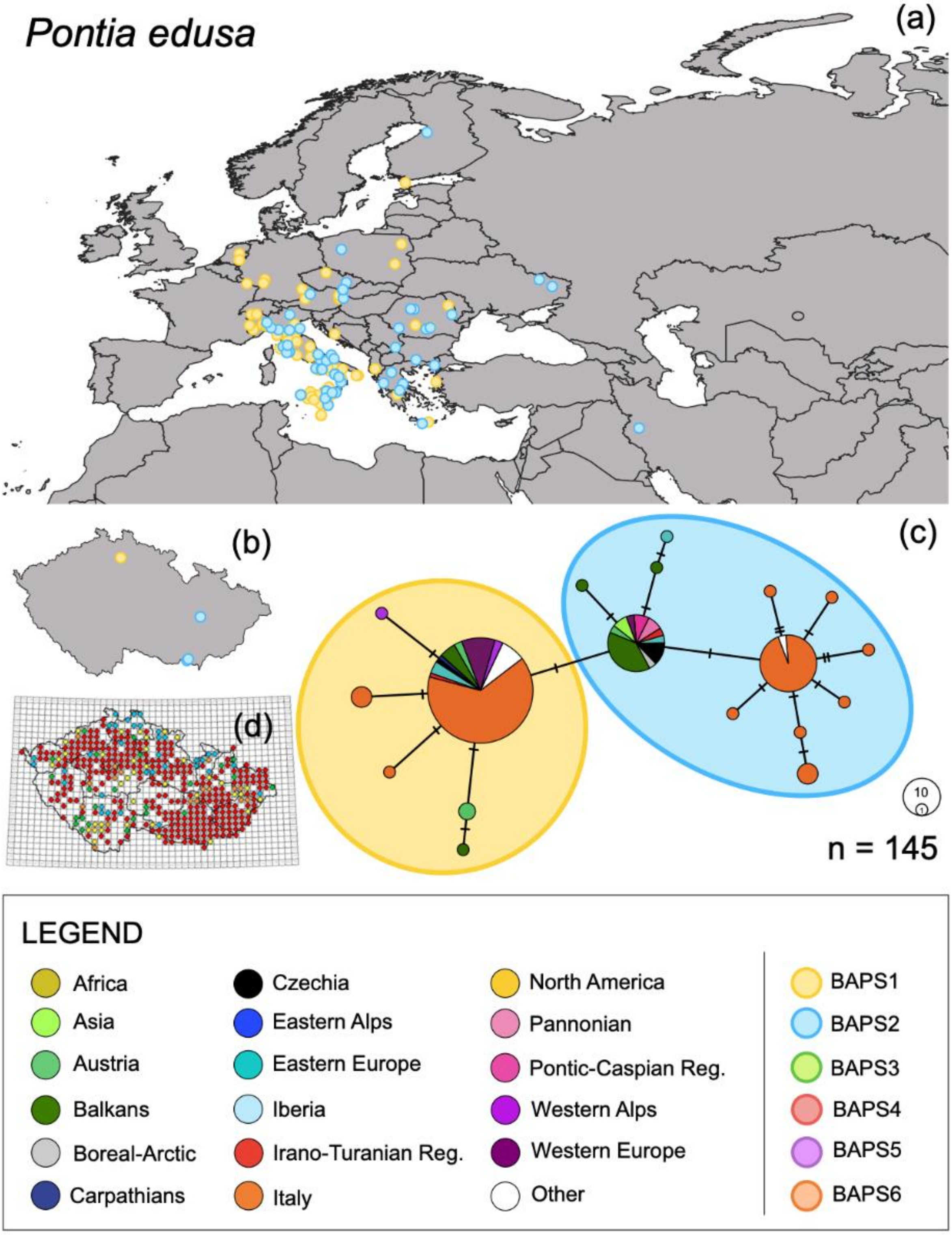


*Pontia edusa* (Fabricius, 1777): **Habitat:** ubiquitous, **Red List:** LC

**BAPS:** BAPS-mix-East. **HAP:** HAP-Italy-Balkans

**Range:** Palearctic, vicariant with *P. daplidice* (west of Rhine River). **Distribution CZ:** warm ruderal places, migratory with open population structure. The genetic patterns are related to its distribution.

### Riodinidae

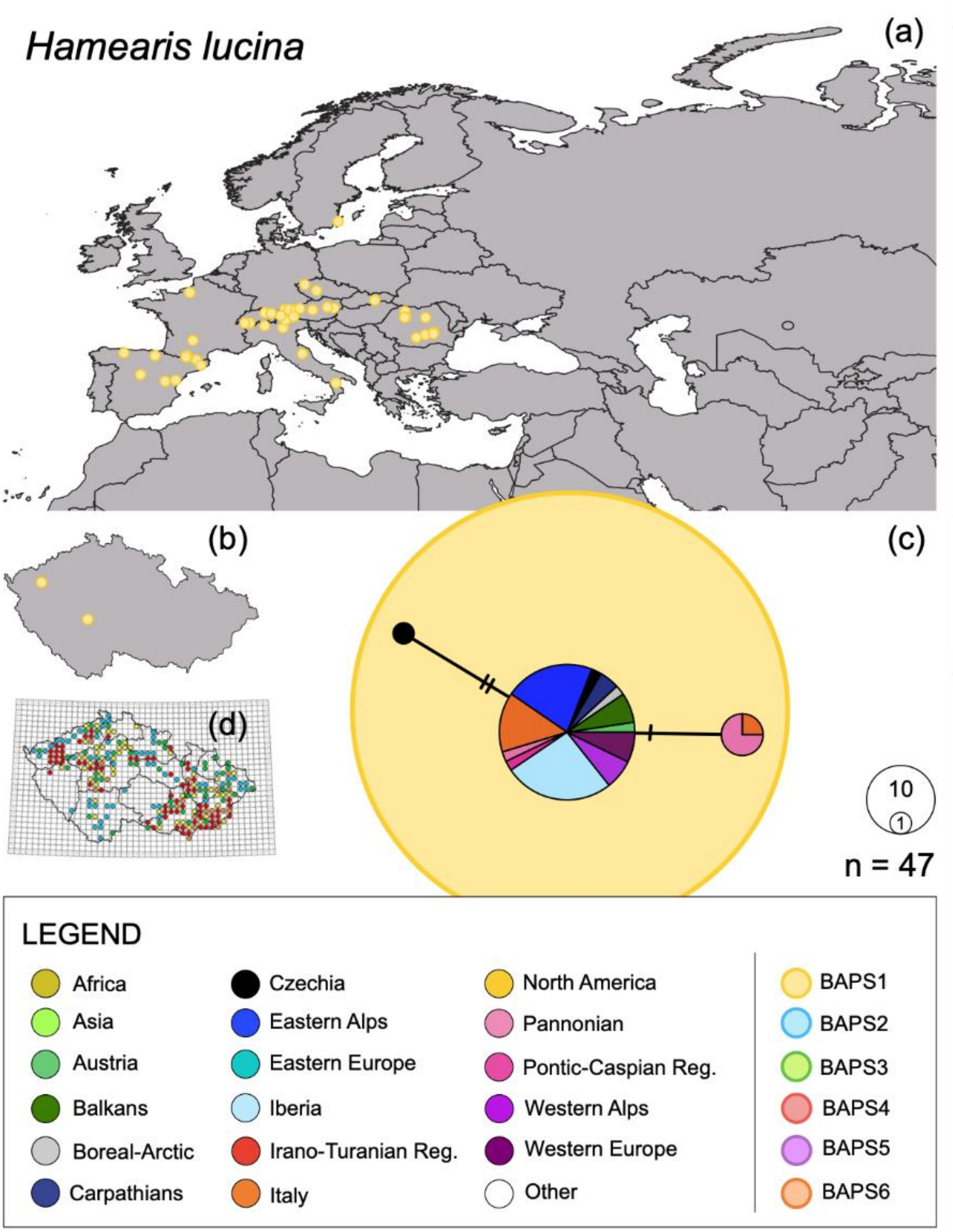


*Hamearis lucina* (Linnaeus, 1758): **Habitat:** mesophilic 2, **Red List:** EN

**BAPS:** BAPS-low. **HAP:** HAP-low

**Range:** European. **Distribution CZ:** flowery meadows in deciduous woodlands. Decreased (loss of open woodlands), currently distributed in open woodlands of both Bohemia and Moravia.

### Lycaenidae

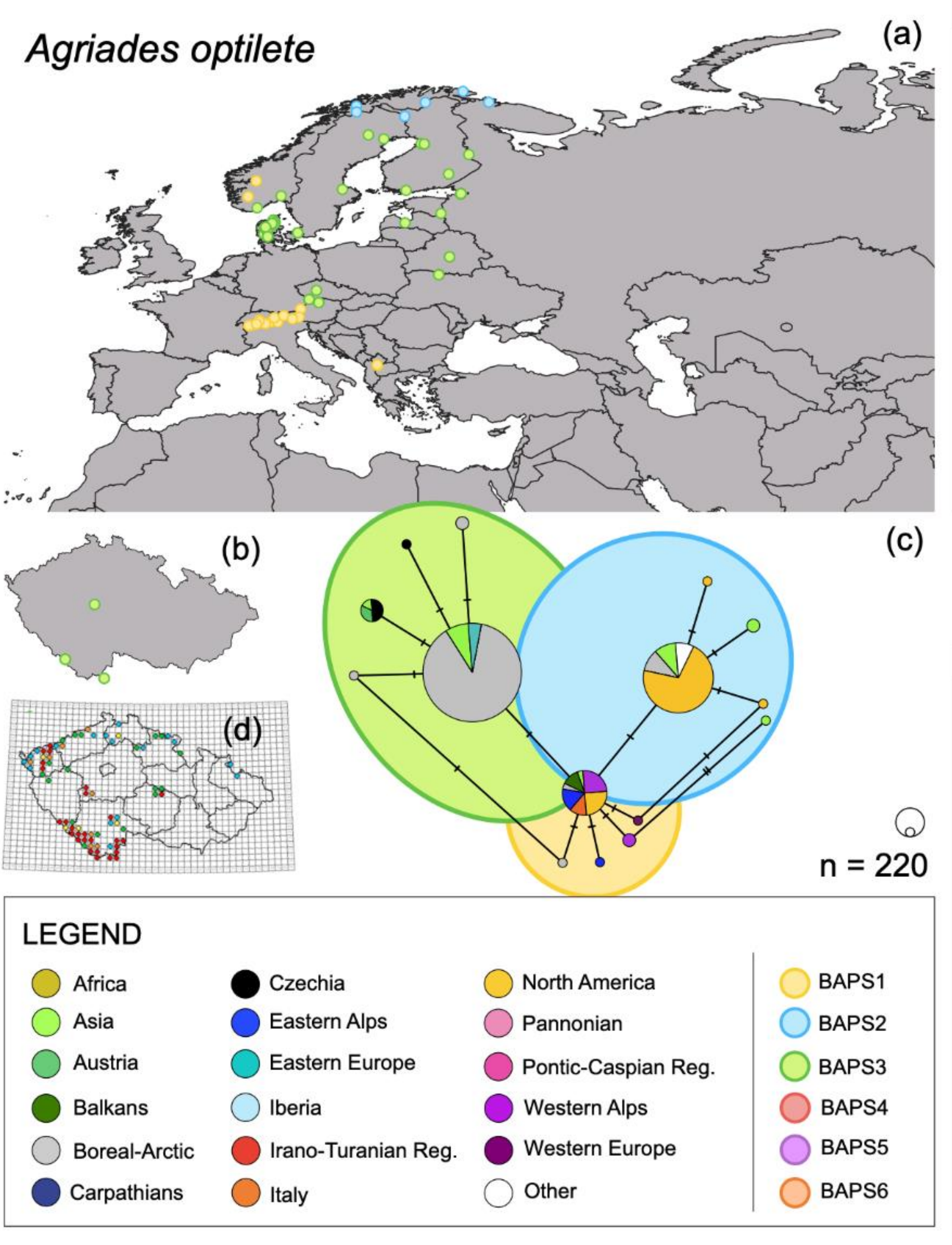


*Agriades optilete* (Knoch, 1781): **Habitat:** tyrphophilic, **Red List:** VU

**BAPS:** BAPS-temperate. **HAP:** HAP-temperate

**Range:** Holarctic. **Distribution CZ:** peat bogs. Isolated localities in south and west Bohemia, Brdy Mountains and Bohemian-Moravian Highlands. Decreased with bogs terrestrification and woodland encroachment.

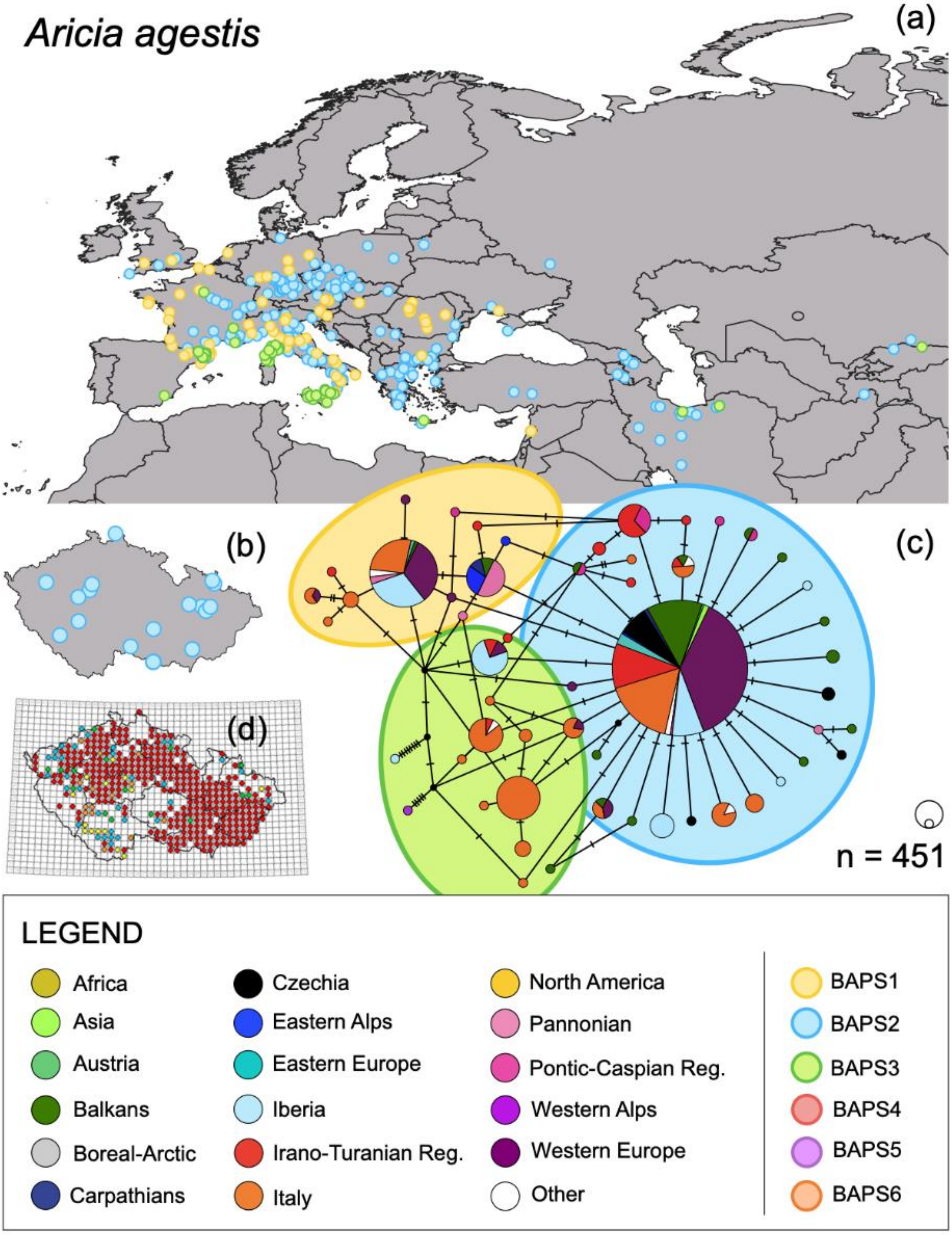


*Aricia agestis* ([Denis & Schiffermüller], 1775): **Habitat:** xerothermophilic 1, **Red List:** LC
**BAPS:** BAPS-wide. **HAP:** HAP-wide-complex

**Range:** Palearctic. **Distribution CZ:** dry warm places, including ruderals. Czech samples belong to an expanding part, although closely related also to the more stable network structure.

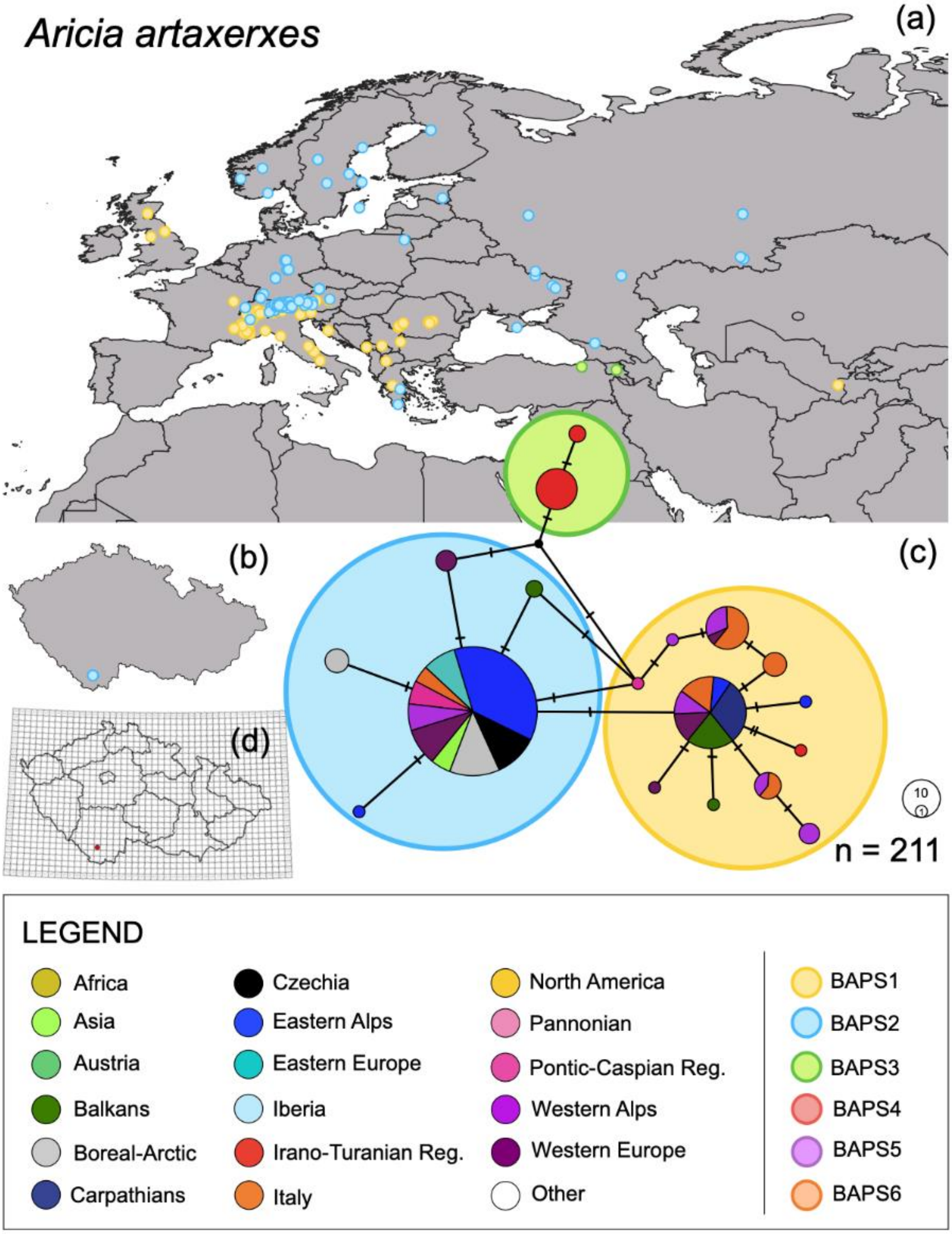


*Aricia artaxerxes* (Fabricius, 1793): **Habitat:** xerothermophilic 1, **Red List:** CR
**BAPS:** BAPS-East-Balkans. **HAP:** HAP-temperate (different haplotype in the Balkans)
**Range:** Palearctic. **Distribution CZ:** documented only from a small karstic area in South Bohemia so far, other records from CZ represent dark form of *A. agestis* (Bartoňová, Beneš, Fric, & Konvička, 2019). The two BAPS lineages found in Europe harbour different

*Wolbachia* strains in most of the cases (Sucháčková Bartoňová, Konvička, Marešová, Wiemers, et al., 2021).

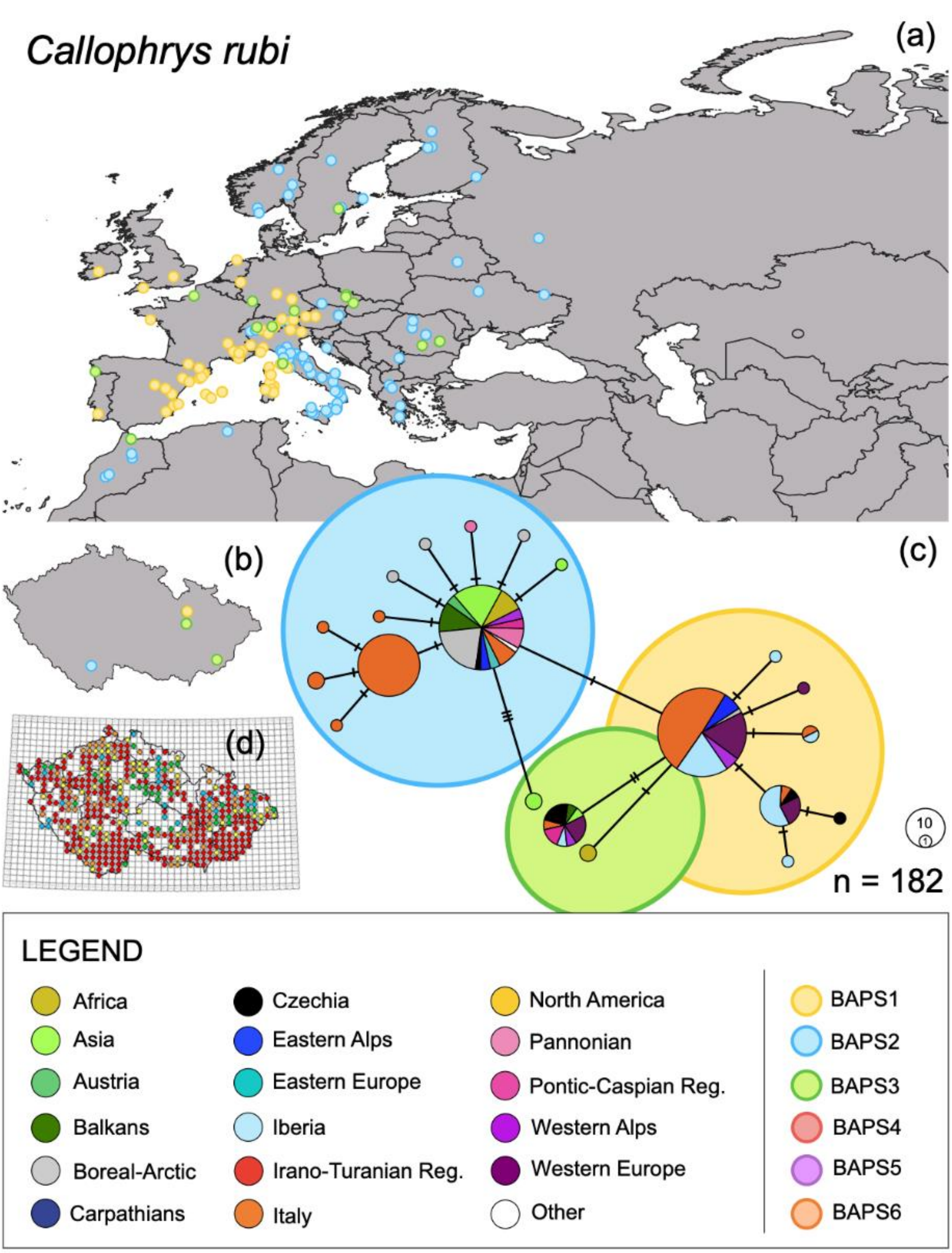


*Callophrys rubi* (Linnaeus, 1758): **Habitat:** mesophilic 2 + tyrphophilic, **Red List:** NT

**BAPS:** BAPS-mix. **HAP:** HAP-mix

**Range:** Palearctic. **Distribution CZ:** two habitat optima – forest-steppes and light evergreen forests on sandy or boggy substrates. Widespread in lowlands and highlands.

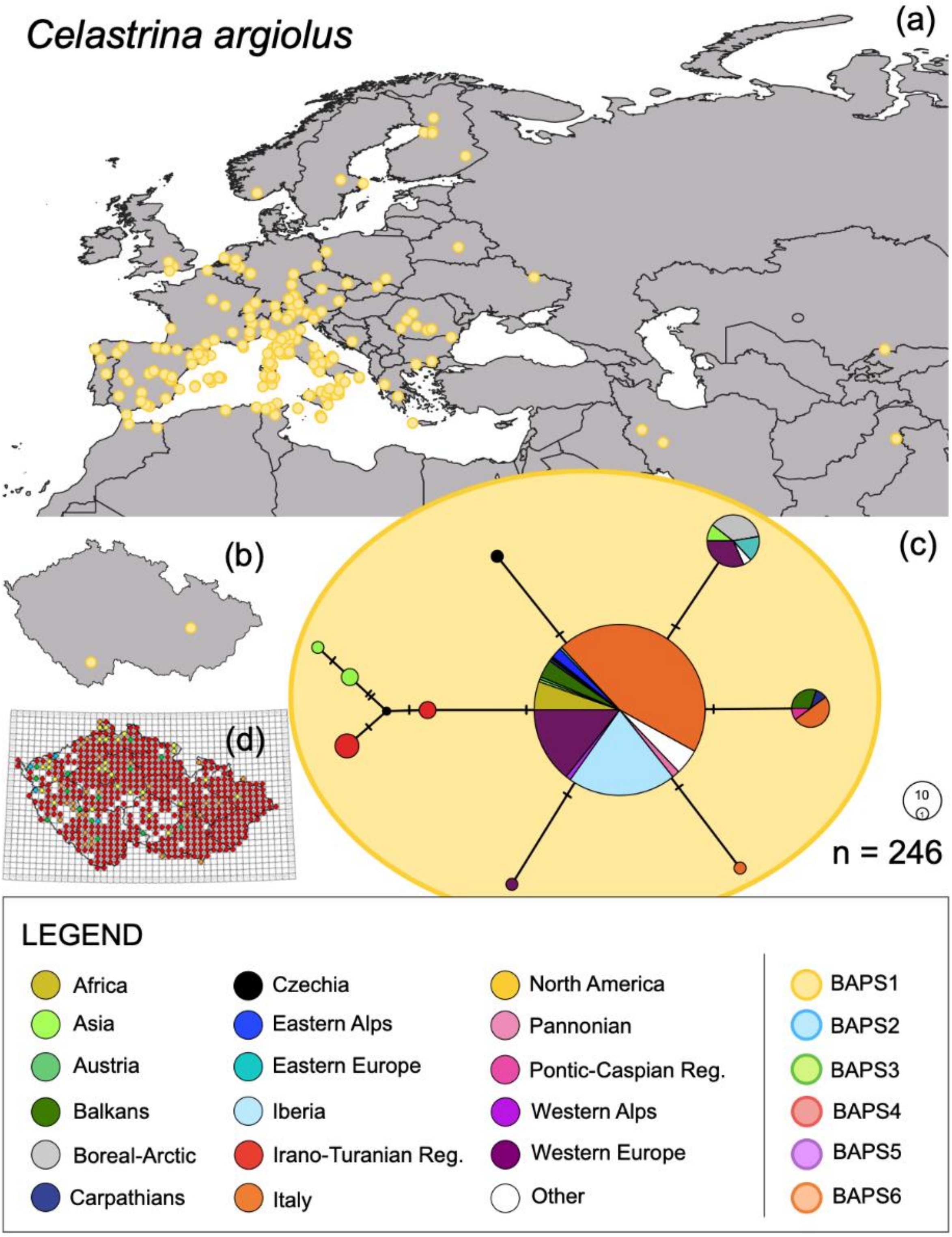


*Celastrina argiolus* (Linnaeus, 1758): **Habitat:** mesophilic 3, **Red List:** LC

**BAPS:** BAPS-low. **HAP:** HAP-wide-low

**Range:** Palearctic. **Distribution CZ:** widespread in all woodland types and shrubby meadows.

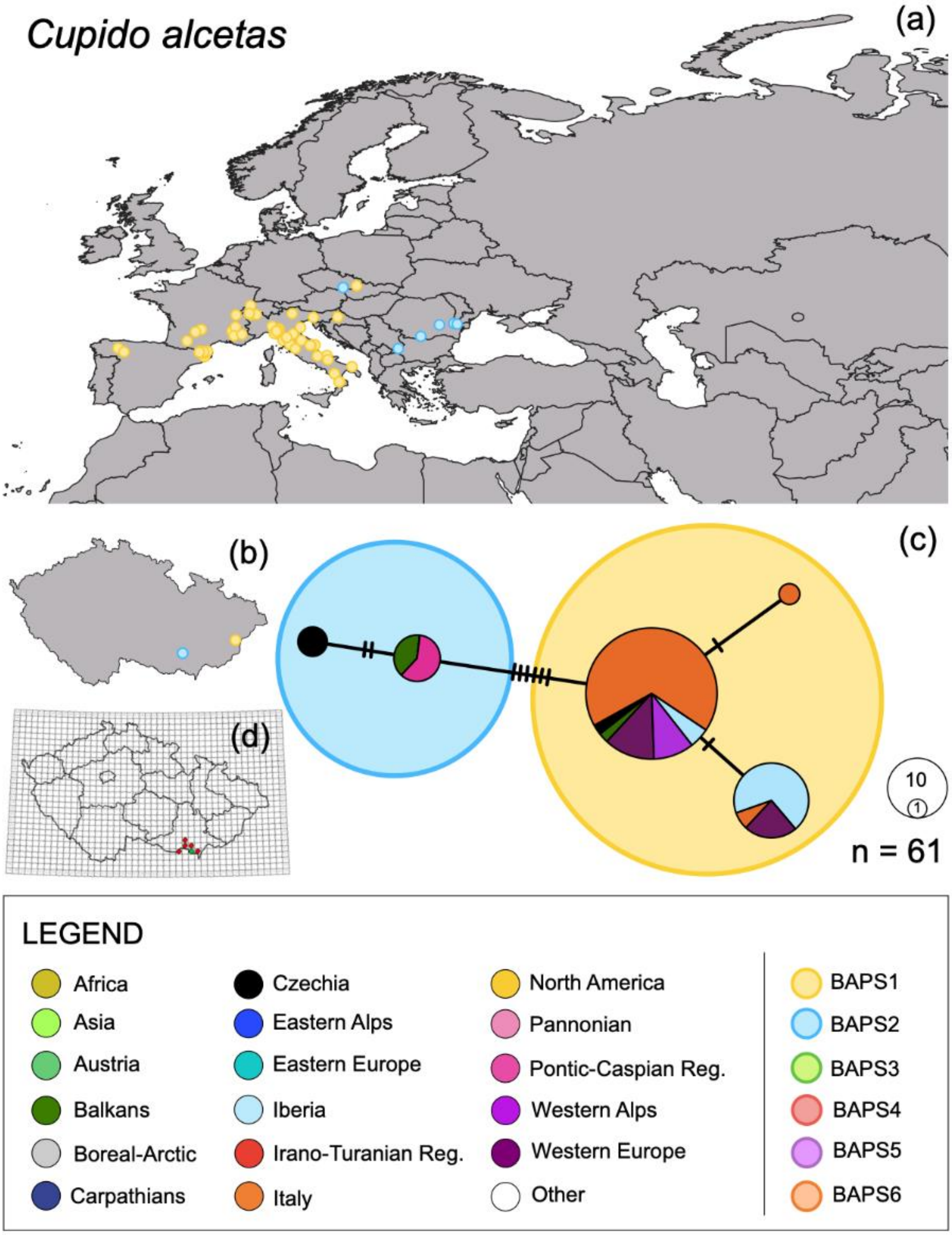


*Cupido alcetas* (Hoffmansegg, 1804): **Habitat:** xerothermophilic 2 + hydrophilic, **Red List:** CR

**BAPS:** BAPS-West + BAPS-East-Balkans. **HAP:** HAP-West + HAP-East-Balkans

**Range:** Pontic-Mediterranean. **Distribution CZ:** South Moravia. Warm shrubby edges near streams, wet meadows and clearings in alluvial woodlands. *Cupido decoloratus* (distributed east of the Alps), shares haplotypes with western *C. alcetas* (BAPS1).

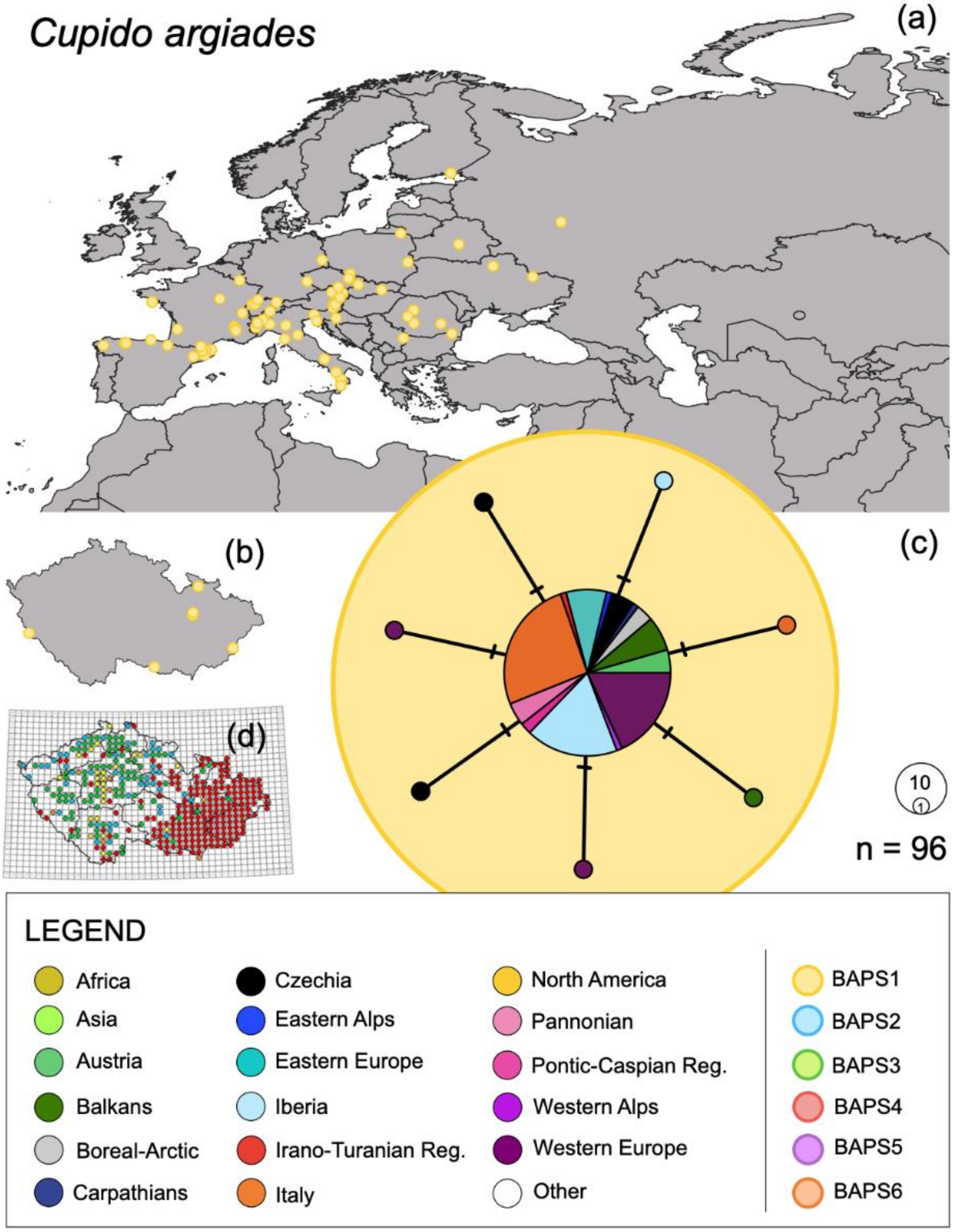


*Cupido argiades* (Pallas, 1771): **Habitat:** xerothermophilic 1, **Red List:** LC

**BAPS:** BAPS-low. **HAP:** HAP-low

**Range:** Palearctic. **Distribution CZ:** across entire area, more common in Moravia. Warm, dry grasslands, currently expanding.

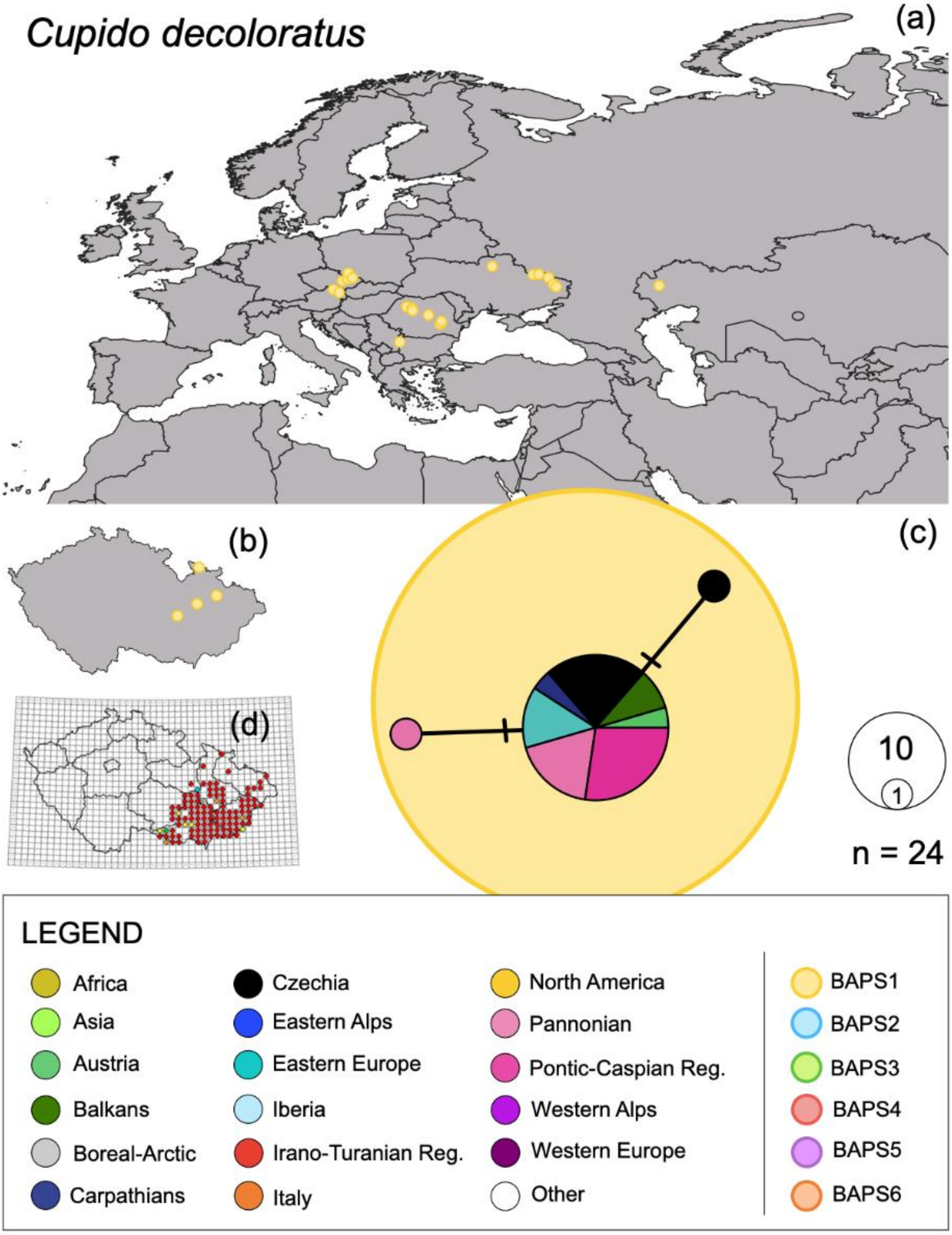


*Cupido decoloratus* (Staudinger, 1886): **Habitat:** xerothermophilic 1, **Red List:** LC

**BAPS:** BAPS-low. **HAP:** HAP-low

**Range:** Pontic-Mediterranean (from South Austria to Turkey). **Distribution CZ:** Moravia. Tall dry ruderal grasslands, anthropogenic localities. Expanding from South Moravia towards the north. *Cupido decoloratus* shares haplotypes with western *C. alcetas* (BAPS1).

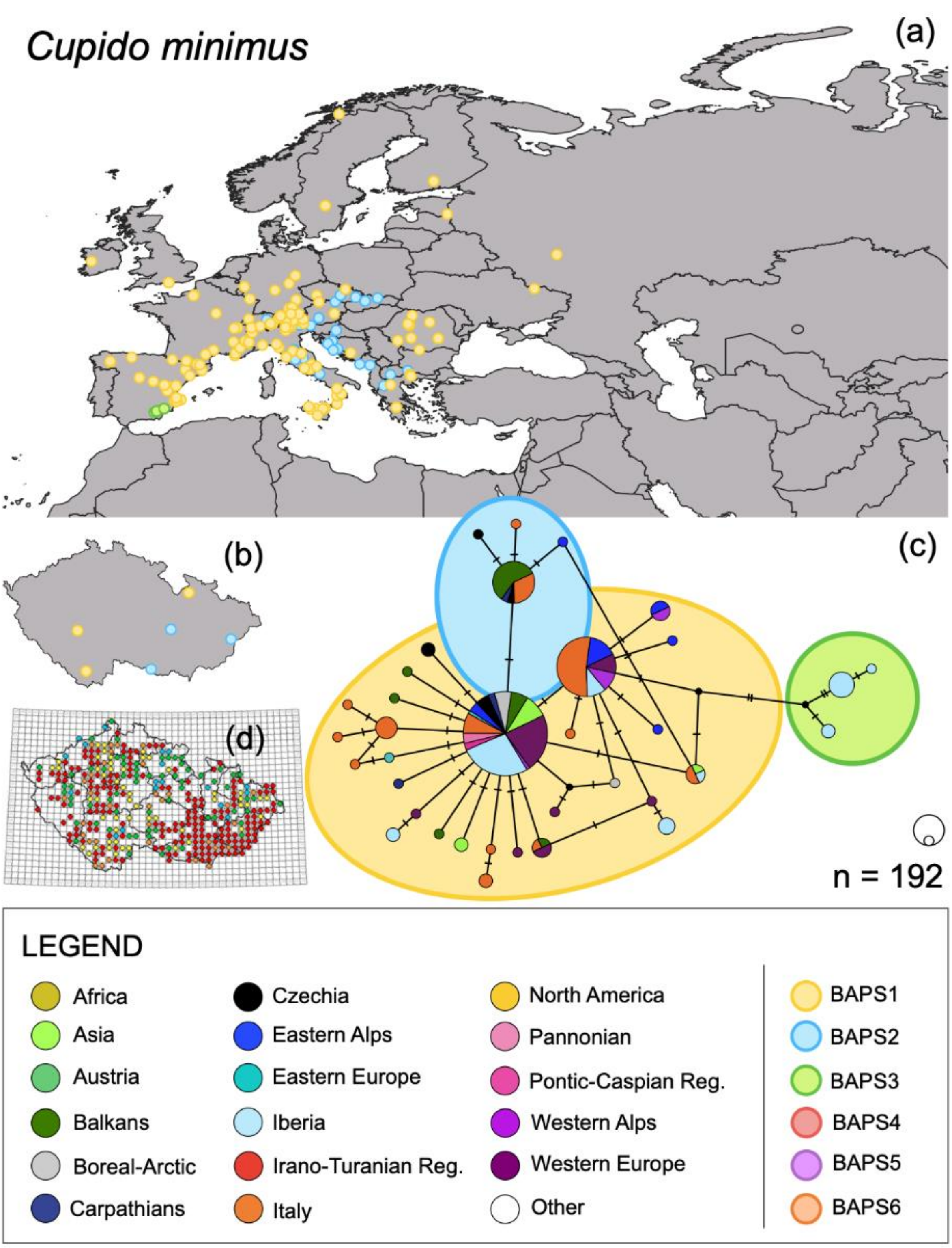


*Cupido minimus* (Fuessly, 1775): **Habitat:** xerothermophilic 1, **Red List:** VU

**BAPS:** BAPS-wide + BAPS-Italy-Balkans. **HAP:** HAP-wide-complex + HAP-Italy-Balkans

**Range:** Euro-Siberian. **Distribution CZ:** dry grasslands. Scattered across the entire country in warmer regions, decreased with abandonment of traditional land use.

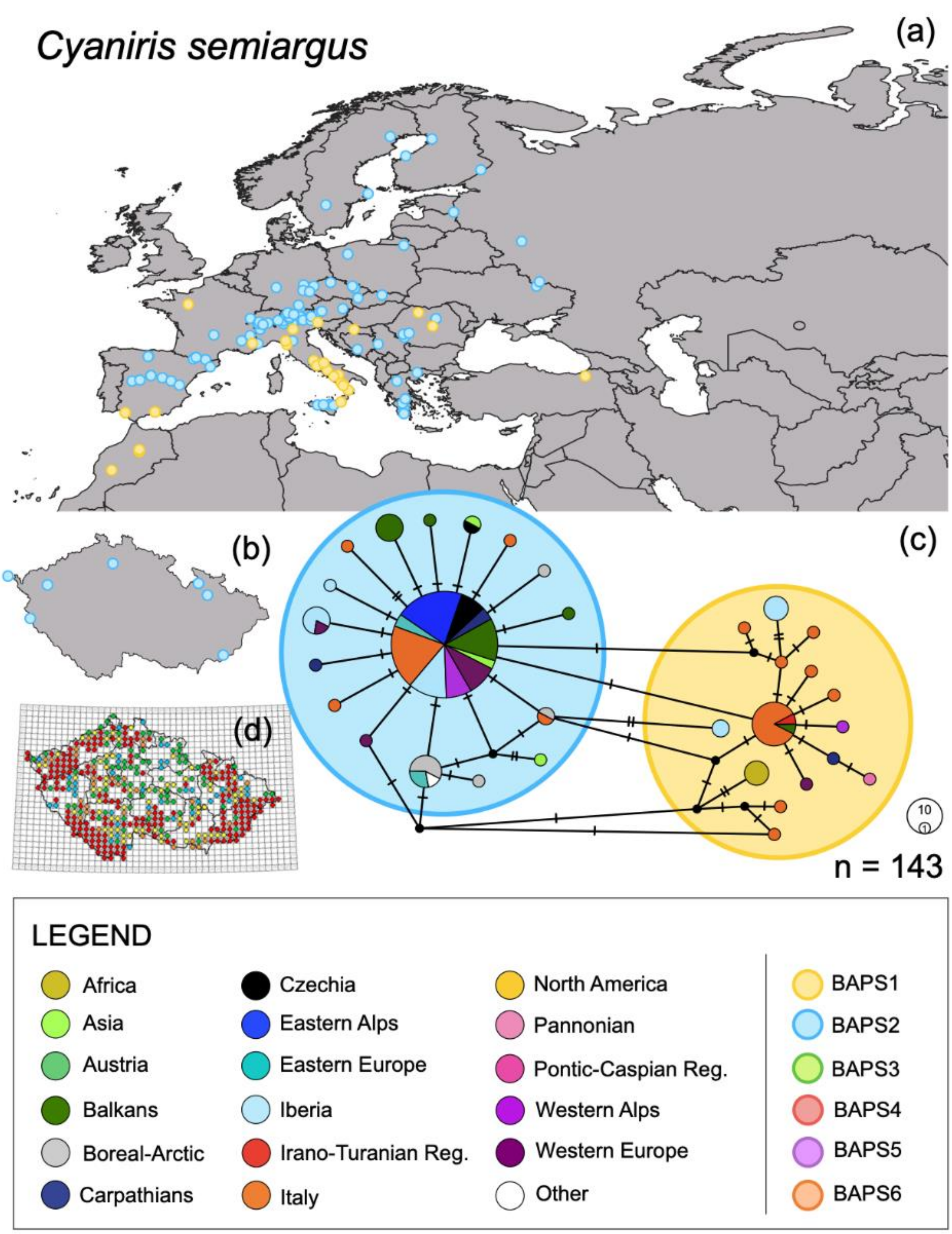


*Cyaniris semiargus* (Rottemburg, 1775): **Habitat:** mesophilic 1 + hydrophilic, **Red List:** VU
**BAPS:** BAPS-wide. **HAP:** HAP-wide-complex

**Range:** Palearctic. **Distribution CZ:** various grassland types, mostly in highlands and mountains across the country. Retreated from interior of the country due to grassland improvements, drainage and afforestation activities.

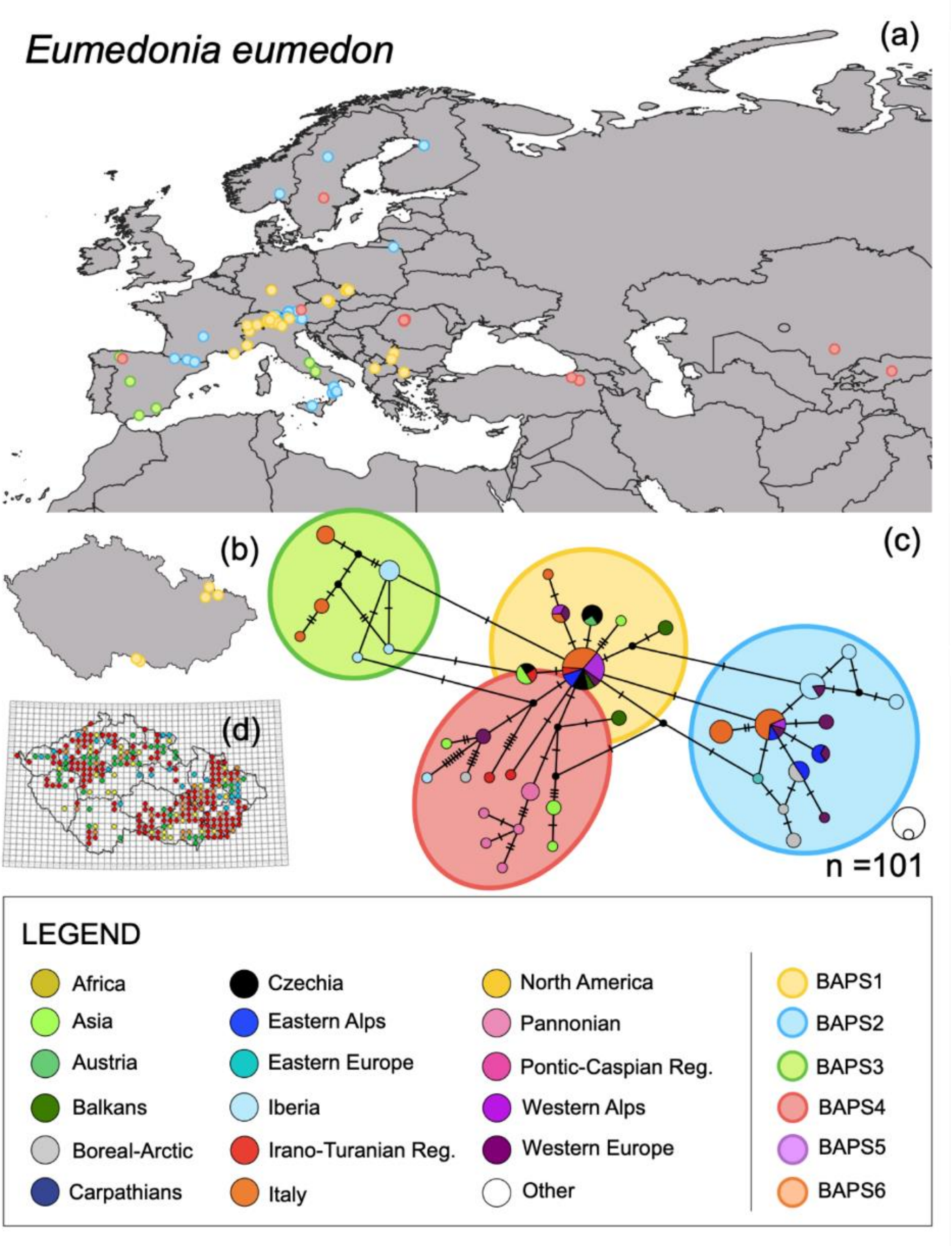


*Eumedonia eumedon* (Esper, 1780): **Habitat:** mesophilic 1 + hydrophilic, **Red List:** NT

**BAPS:** BAPS-Italy-Balkans. **HAP:** HAP-Italy-Balkans

**Range:** Palearctic, dot-like distribution. **Distribution CZ:** Wet meadows in river valleys, scattered distribution across the country, including urban areas.

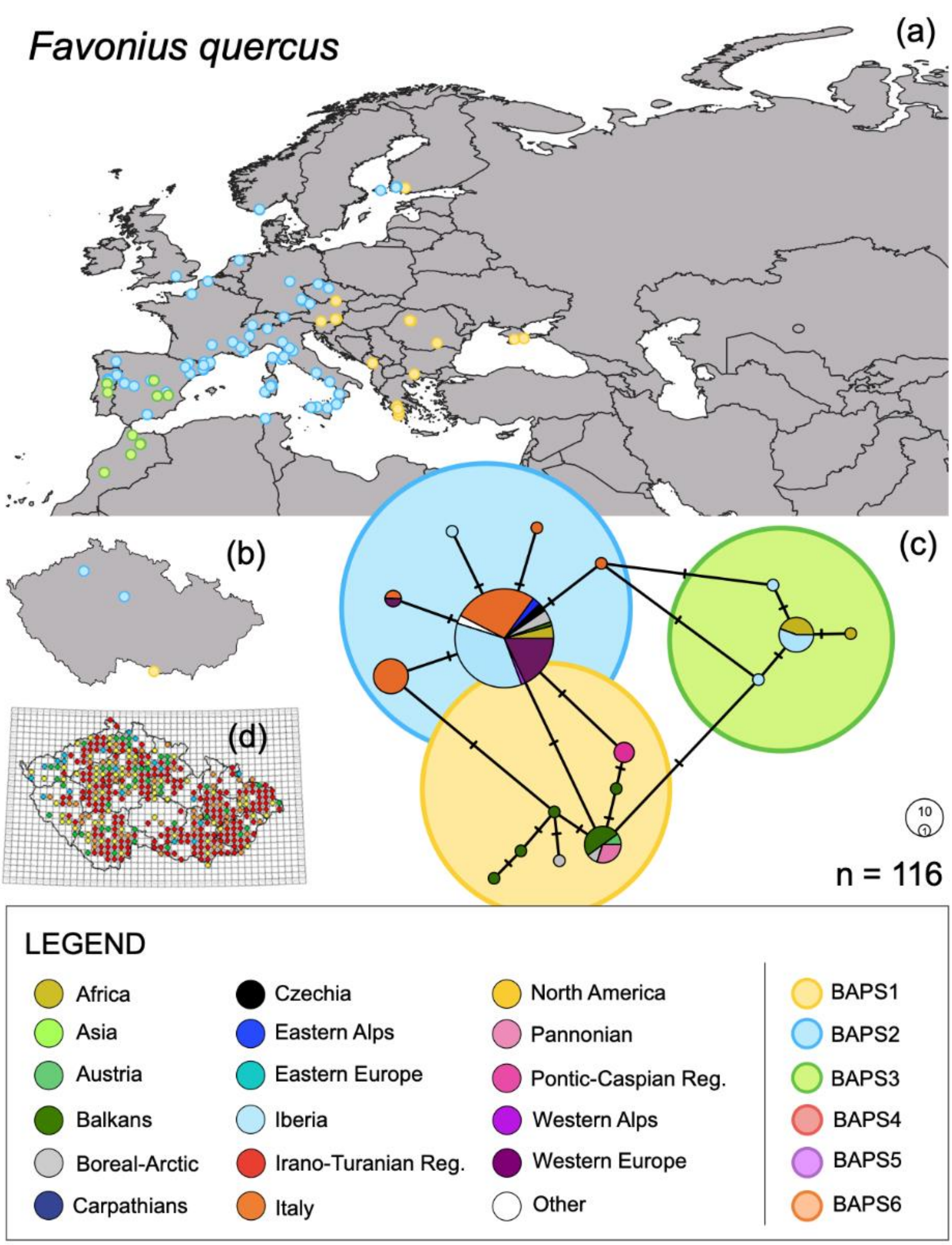


*Favonius quercus* (Linnaeus, 1758): **Habitat:** mesophilic 3, **Red List:** LC

**BAPS:** BAPS-West. **HAP:** HAP-West

**Range:** Western Palearctic. **Distribution CZ:** oak woodlands. Widespread across the country, missing in mountains.

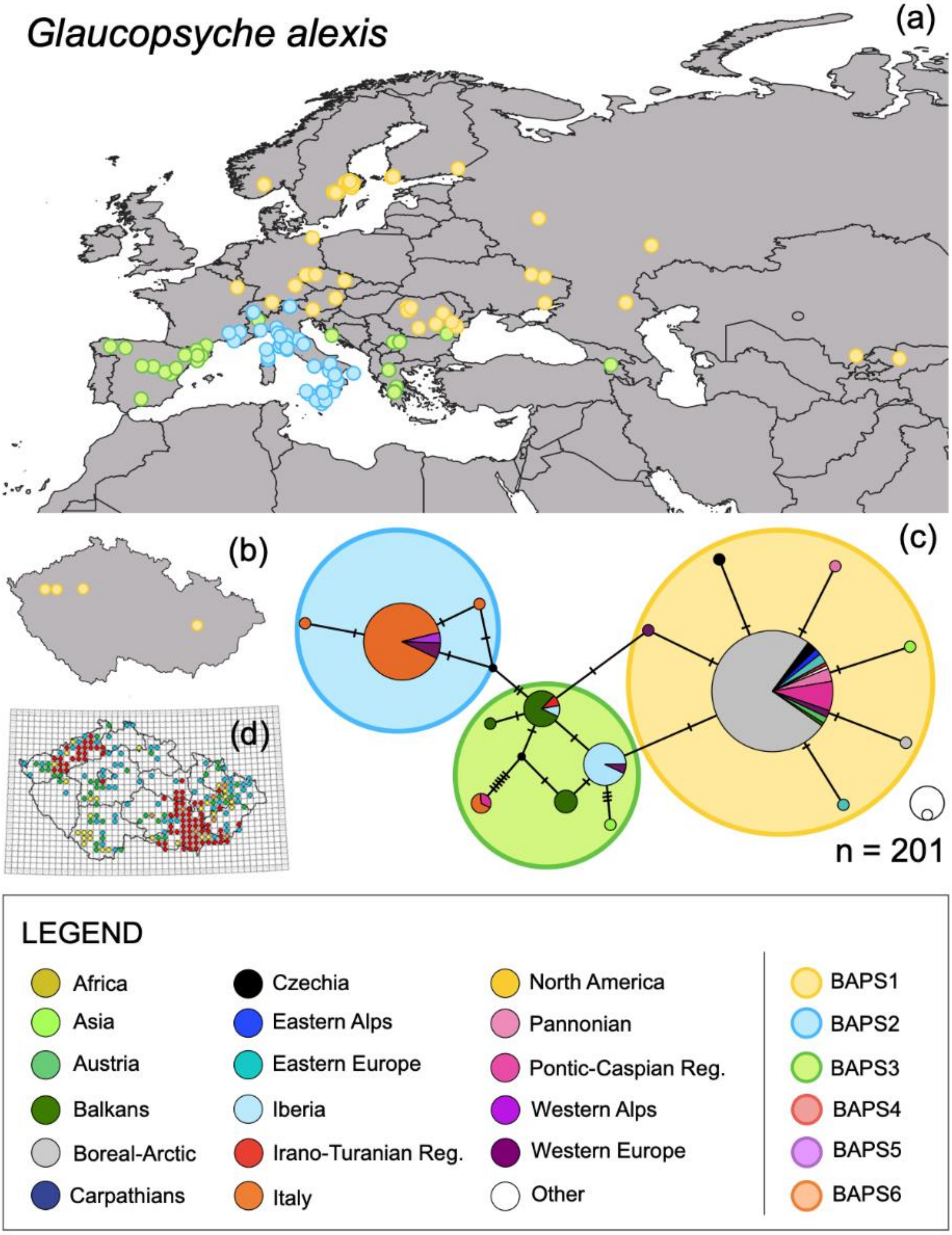


*Glaucopsyche alexis* (Poda, 1761): **Habitat:** xerothermophilic 2, **Red List:** VU

**BAPS:** BAPS-temperate. **HAP:** HAP-temperate

**Range:** Palearctic. **Distribution CZ:** overgrown dry grasslands. Lowland parts of Moravia, Central Bohemian Uplands and Doupov Mountains, decreased.

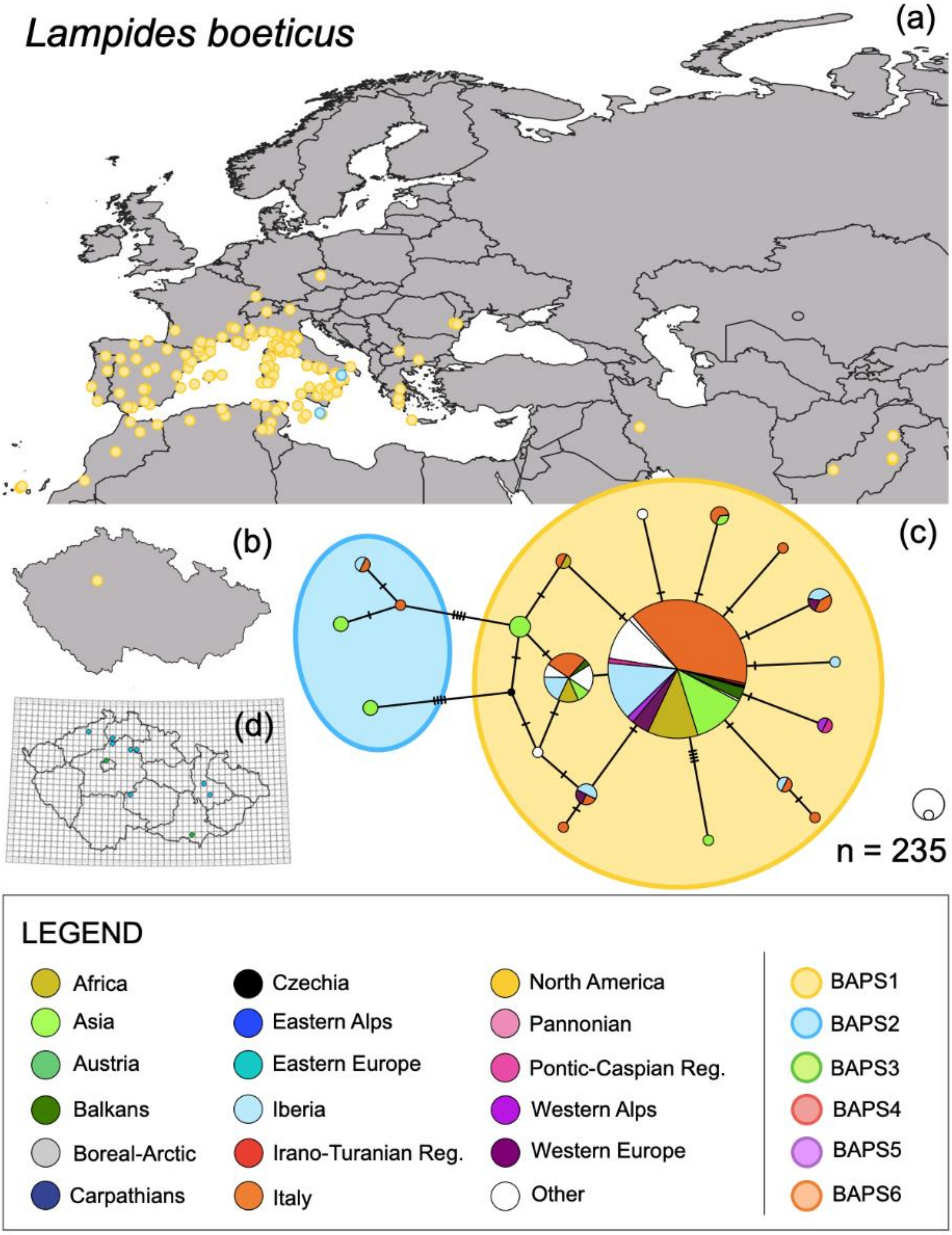


*Lampides boeticus* (Linnaeus, 1767): very rare vagrant (the latest observation in 1950s). The sequenced specimen was introduced to Prague with fruits. Not a part of Czech fauna, excluded from analyses.

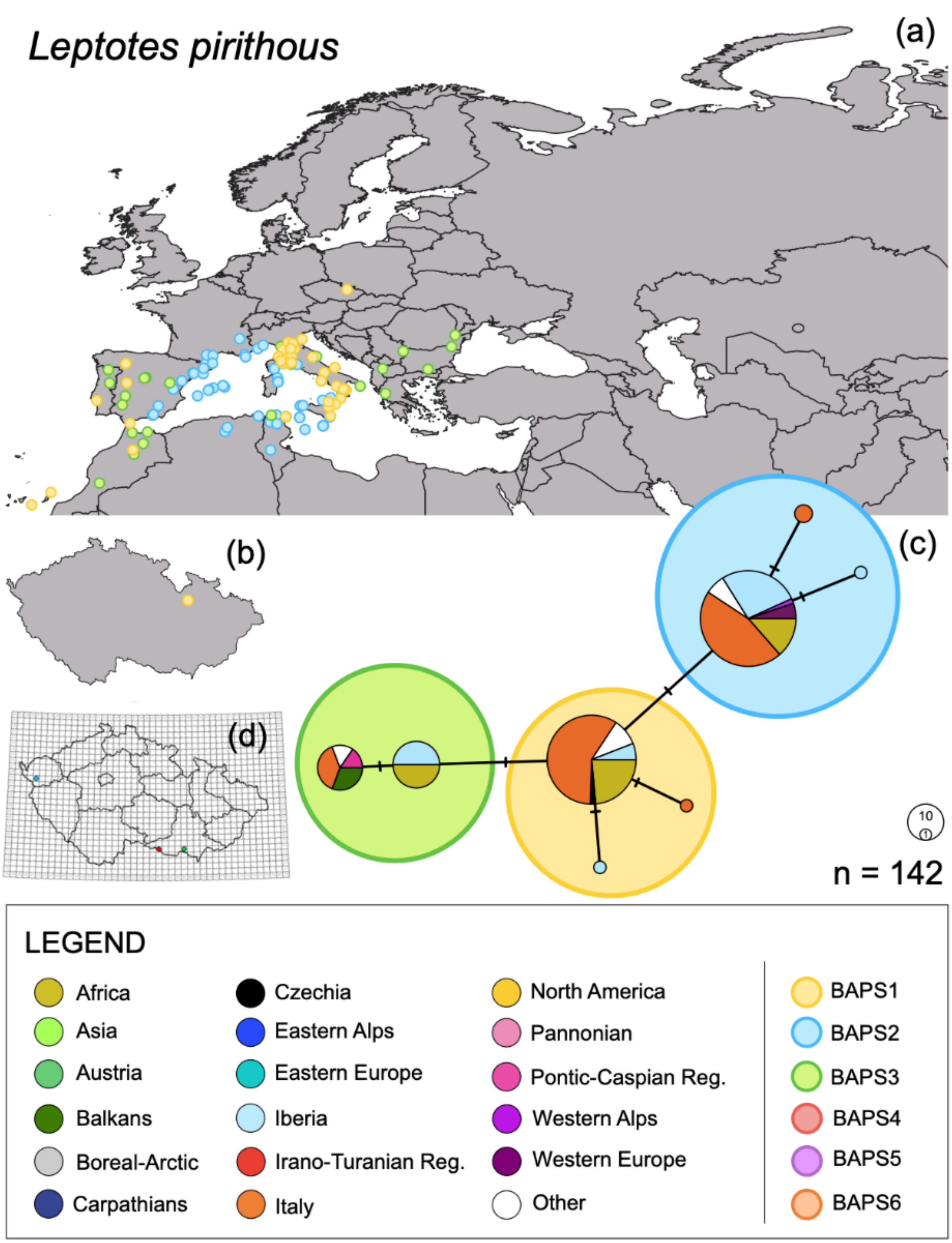


*Leptotes pirithous* (Linnaeus, 1767): **Habitat:** ubiquitous, **Red List:** LC

**BAPS:** BAPS-West. **HAP:** HAP-West

**Range:** Afrotropical, Mediterranean. **Distribution CZ:** dry grasslands and agricultural landscapes. Rare migrant in CZ, several records from recent years (both Bohemia and Moravia).

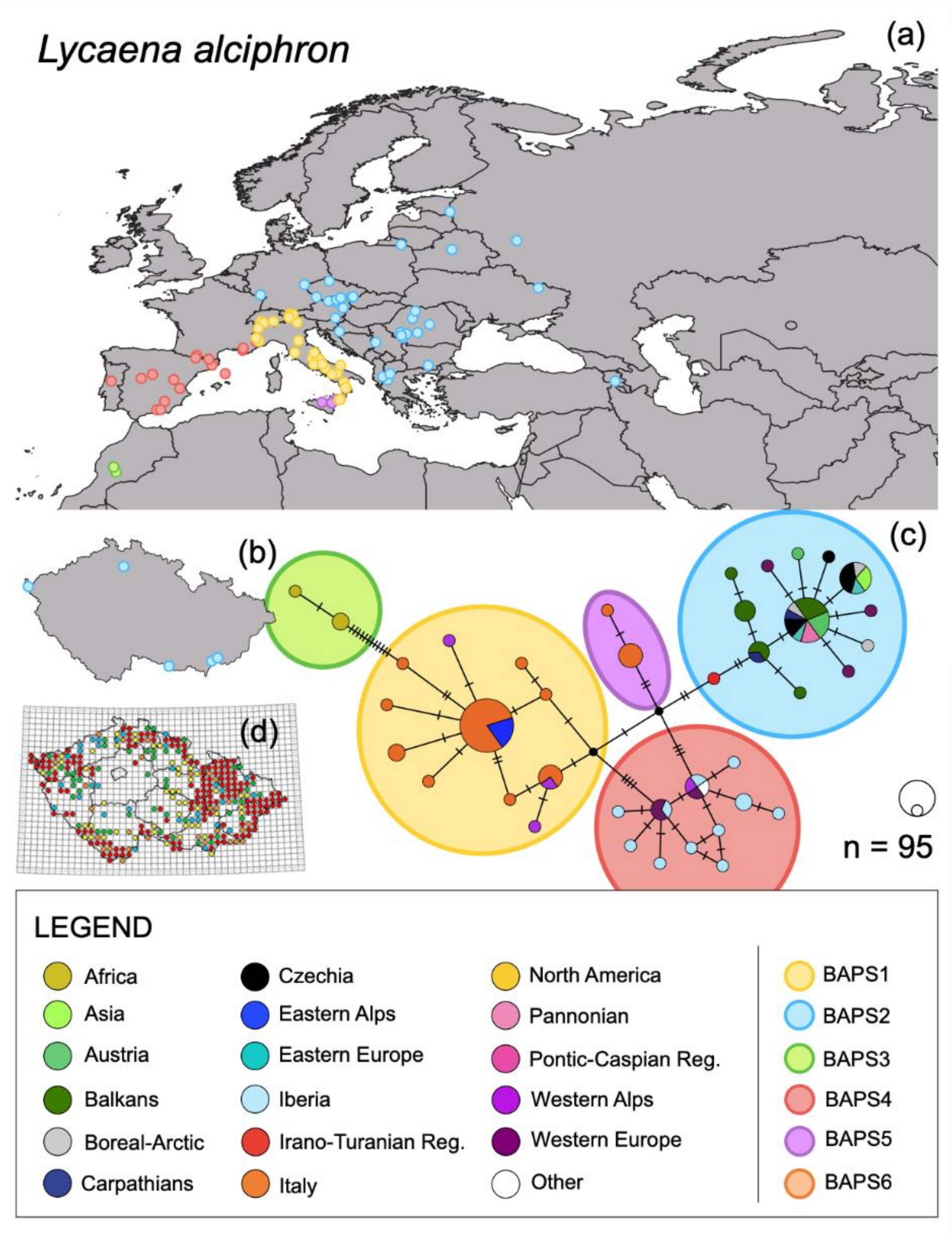


*Lycaena alciphron* (Rottemburg, 1775): **Habitat:** mesophilic 1 + hydrophilic, **Red List:** VU
**BAPS:** BAPS-East-Balkans. **HAP:** HAP-East-Balkans
**Range:** Western Palearctic. **Distribution CZ:** in piedmont areas, cooler flowery meadows and bogs, also heathlands and sands in lowlands.

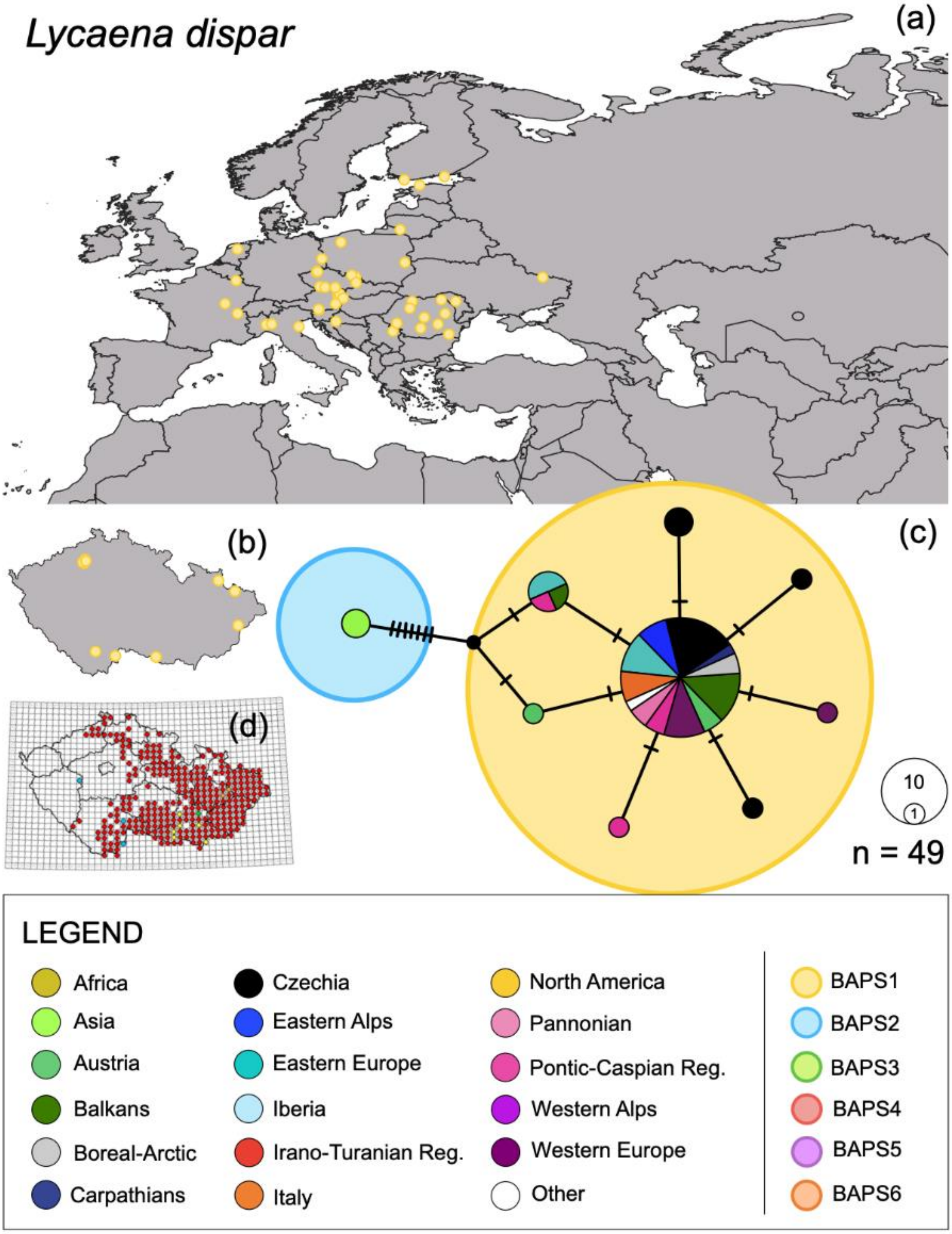


*Lycaena dispar* ([Haworth], 1802): **Habitat:** hydrophilic. **Red List:** LC

**BAPS:** BAPS-temperate, **HAP:** HAP-temperate (such distribution; several haplotypes found in CZ)

**Range:** Euro-Siberian (missing in Southern Europe). **Distribution CZ:** wet meadows and stream banks, wet ruderal places. Rapidly expanding from the south, currently across almost entire country.

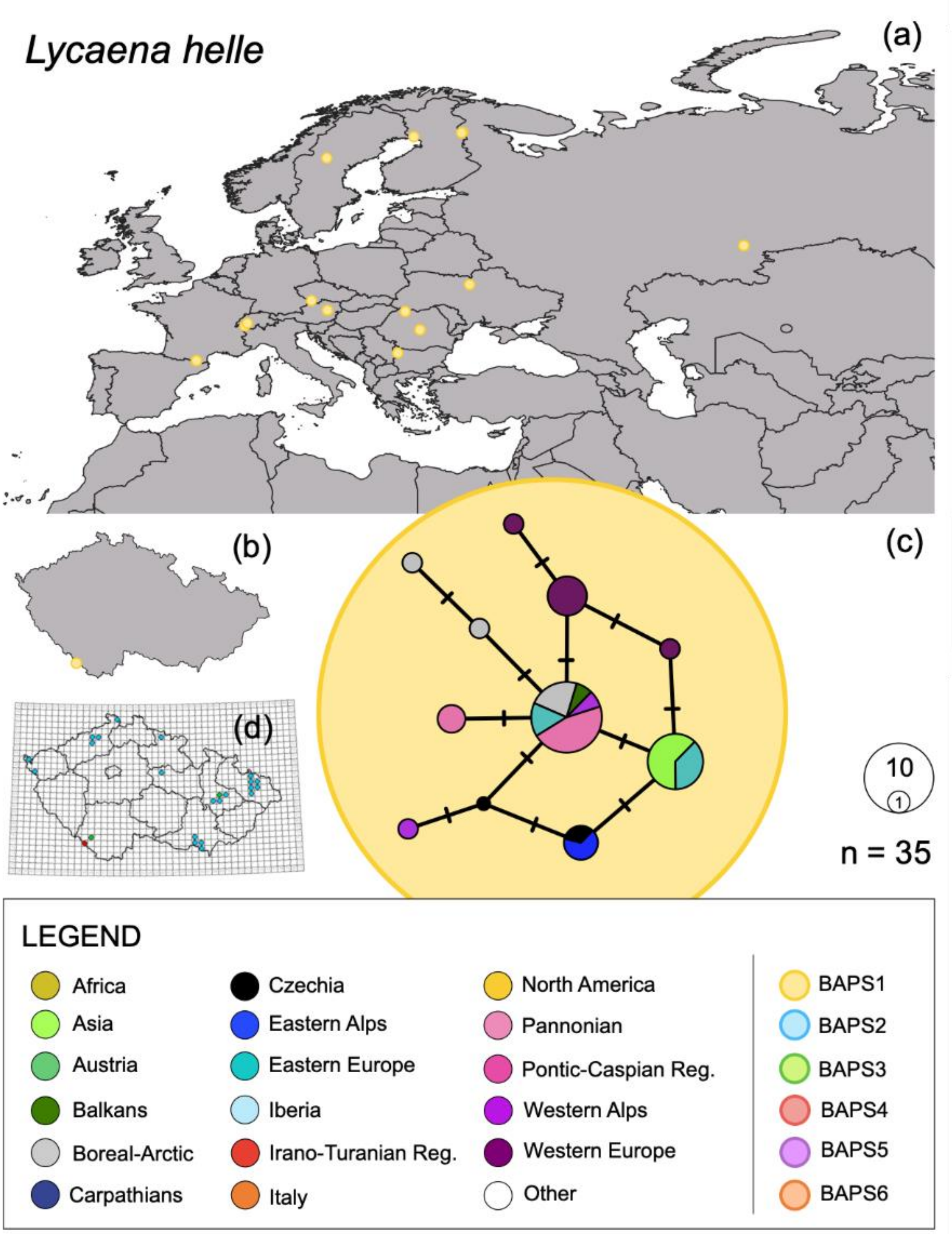


*Lycaena helle* ([Denis & Schiffermüller], 1775): **Habitat:** hydrophilic. **Red List:** CR
**BAPS:** BAPS-low. **HAP:** HAP-temperate
**Range:** Euro-Siberian (missing in Southern Europe). **Distribution CZ:** historically distributed in wet meadows and bogs with *Polygonium bistorta*. Extinct since 1952. In early 2000s reintroduced to a single locality in Bohemian Forest from a population close to Mariazell, Austria (Předotová, 2013). In the haplotype network, this originally Eastern Alpine population is closely related to haplotypes from Eastern Europe and continental Asia.

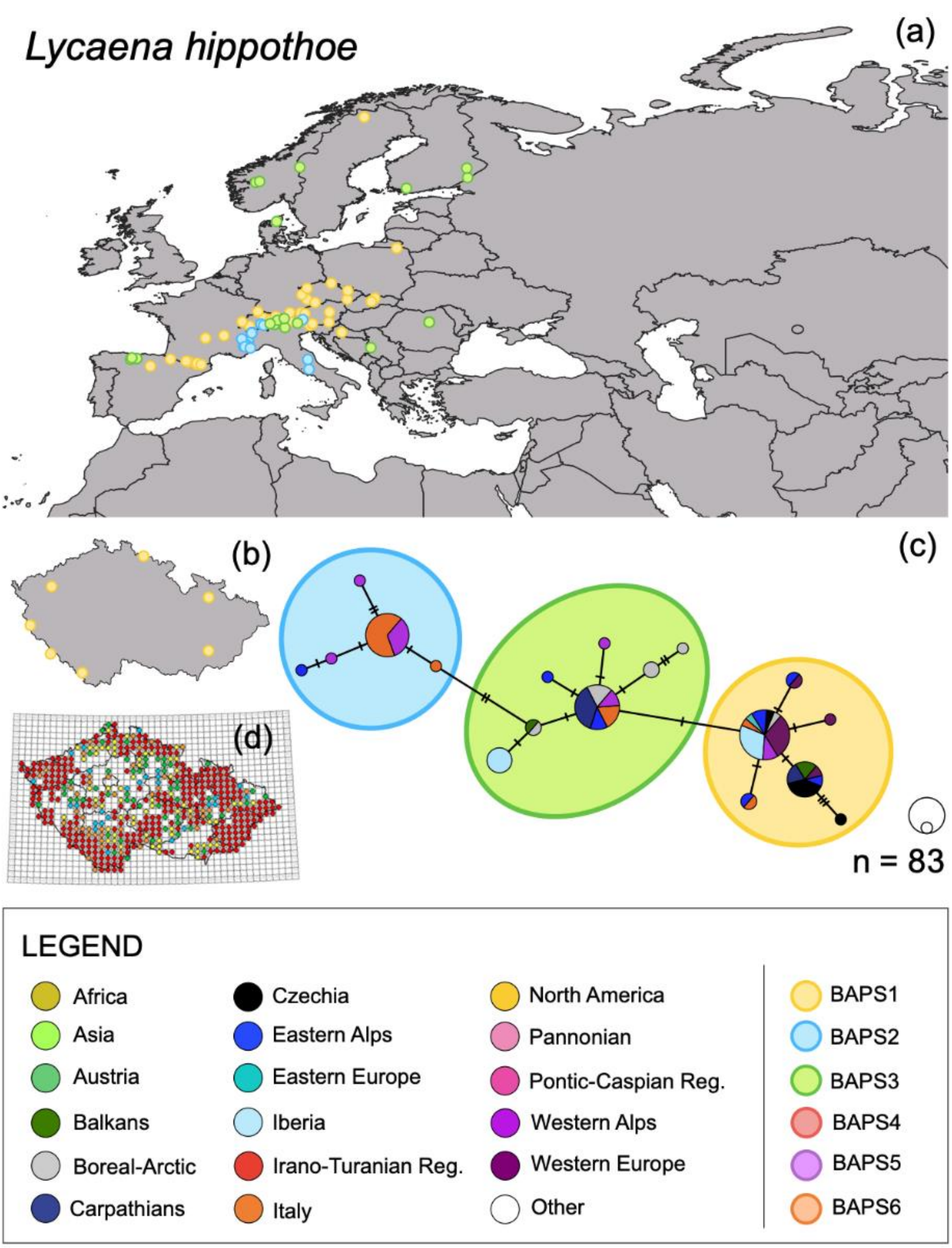


*Lycaena hippothoe* (Linnaeus, 1760): **Habitat:** mesophilic 1 + hydrophilic. **Red List:** NT

**BAPS:** BAPS-West. **HAP:** HAP-West

**Range:** Euro-Siberian. **Distribution CZ:** mountain and piedmont areas across entire country, wet meadows and pastures, slowly decreasing.

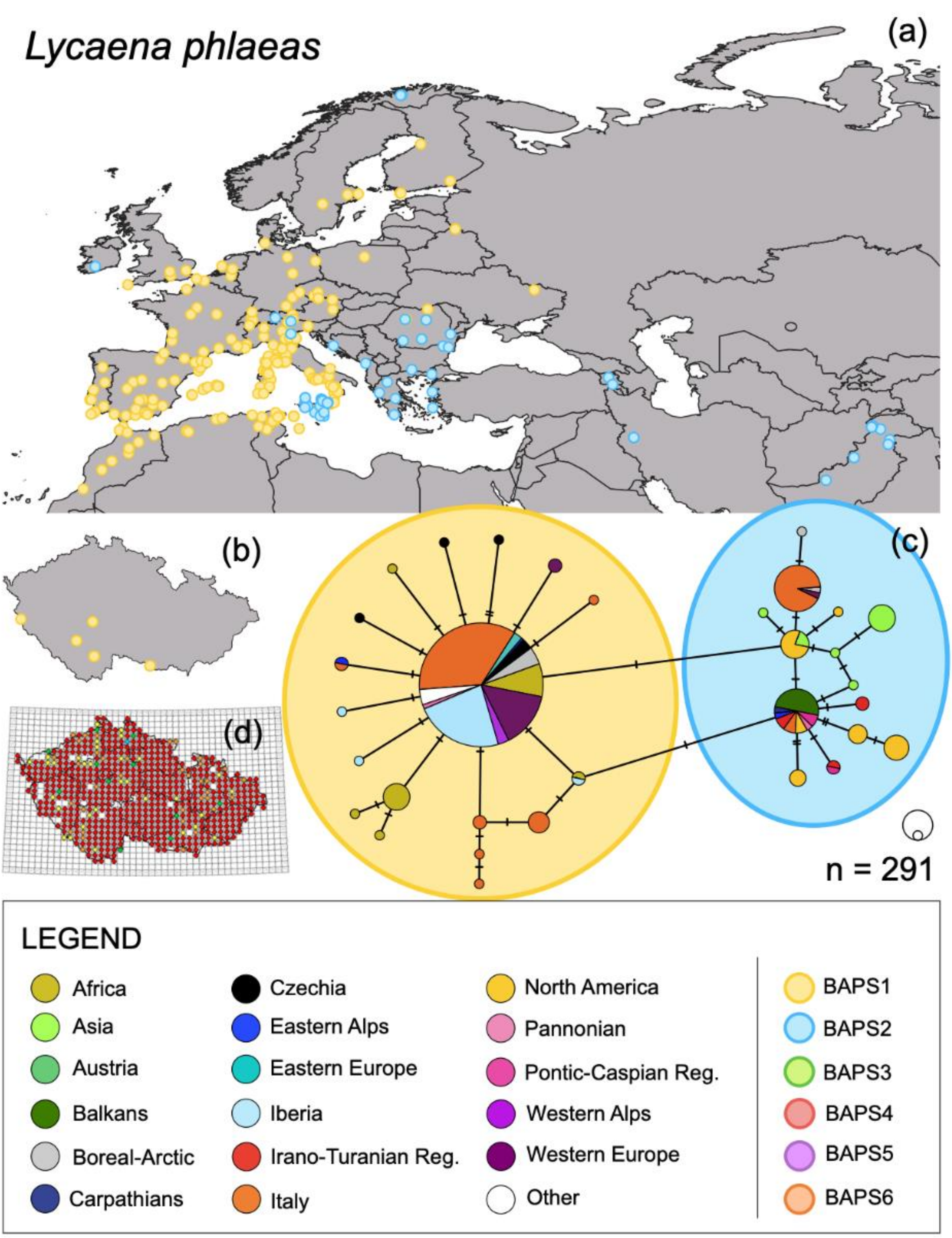


*Lycaena phlaeas* (Linnaeus, 1760): **Habitat:** ubiquitous. **Red List:** LC

**BAPS:** BAPS-West. **HAP:** HAP-West

**Range:** Holarctic. **Distribution CZ:** diverse habitats, widespread and common.

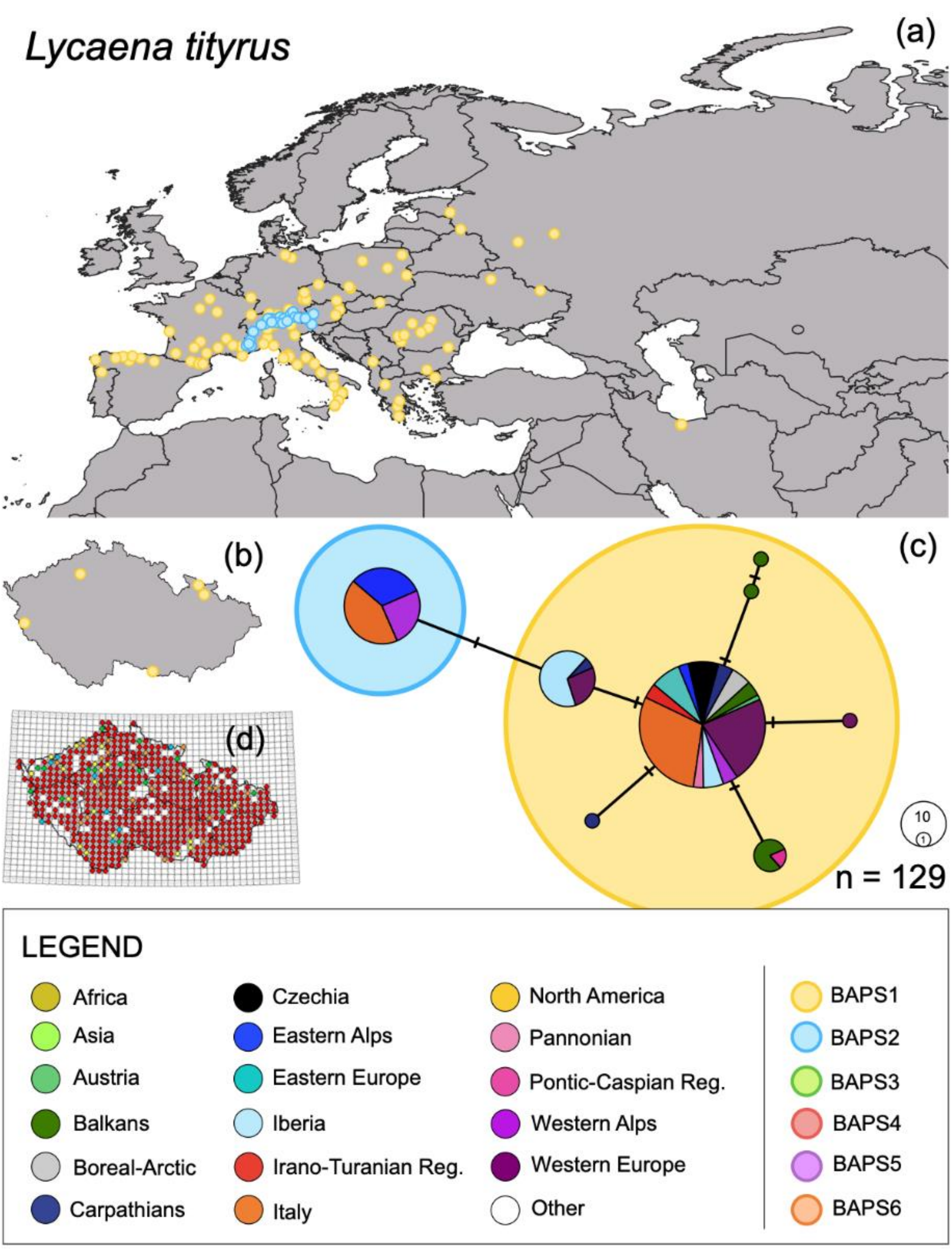


*Lycaena tityrus* (Poda, 1761): **Habitat:** mesophilic 1 + xerothermophilic 1. **Red List:** LC
**BAPS:** BAPS-wide. **HAP:** HAP-wide-low

**Range:** Western Palearctic. **Distribution CZ:** dry shrubby grasslands, mesic and wet flowery meadows. Widespread across the country, avoiding high mountains and intensive agricultural landscapes.

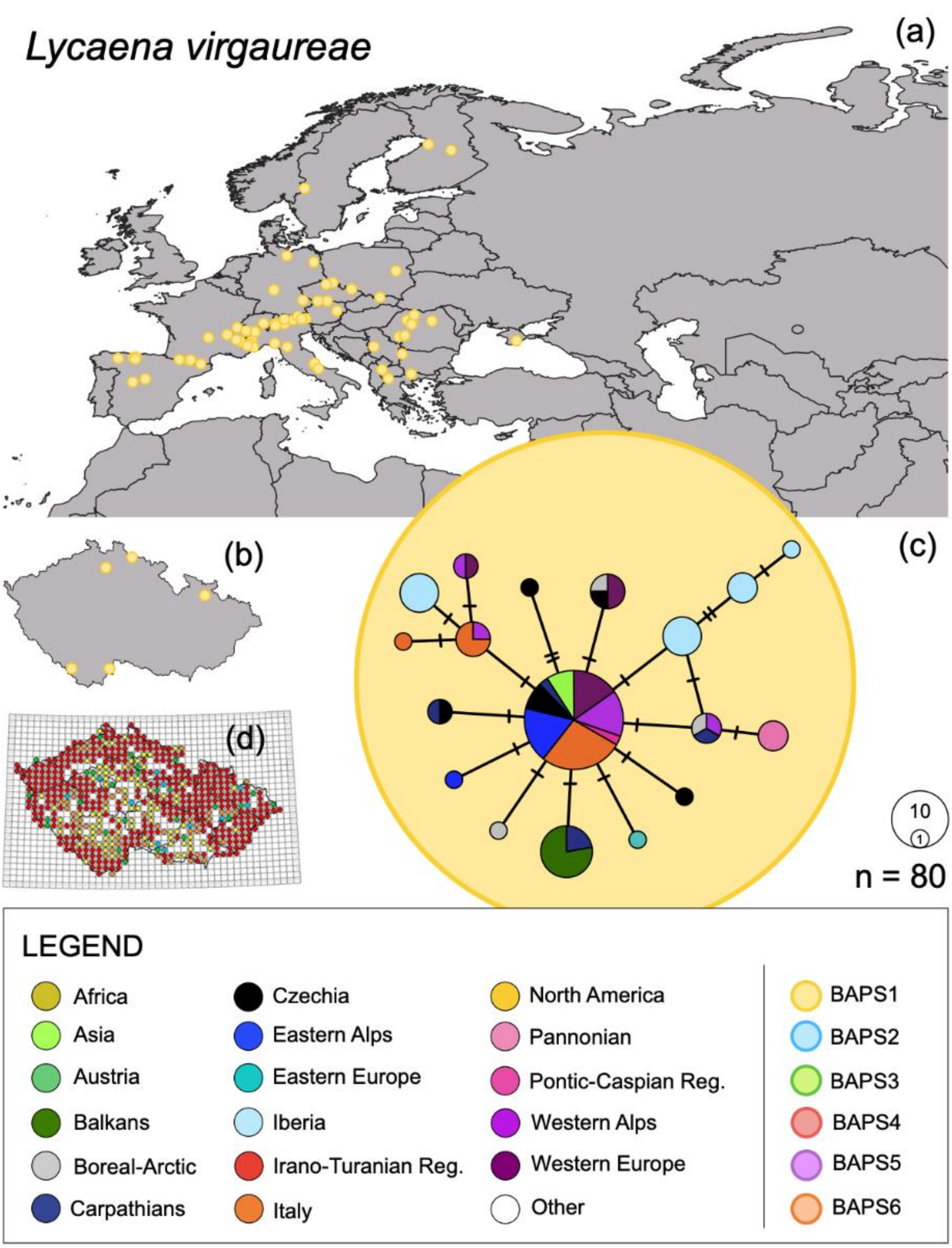


*Lycaena virgaureae* (Linnaeus, 1758): **Habitat:** mesophilic 2. **Red List:** NT

**BAPS:** BAPS-low. **HAP:** HAP-Italy-Balkans (Iberia is divergent)

**Range:** Palearctic. **Distribution CZ:** open wet places, scrublands, mainly in highlands and mountains. Widespread, but retreated from lowlands with intensive agriculture.

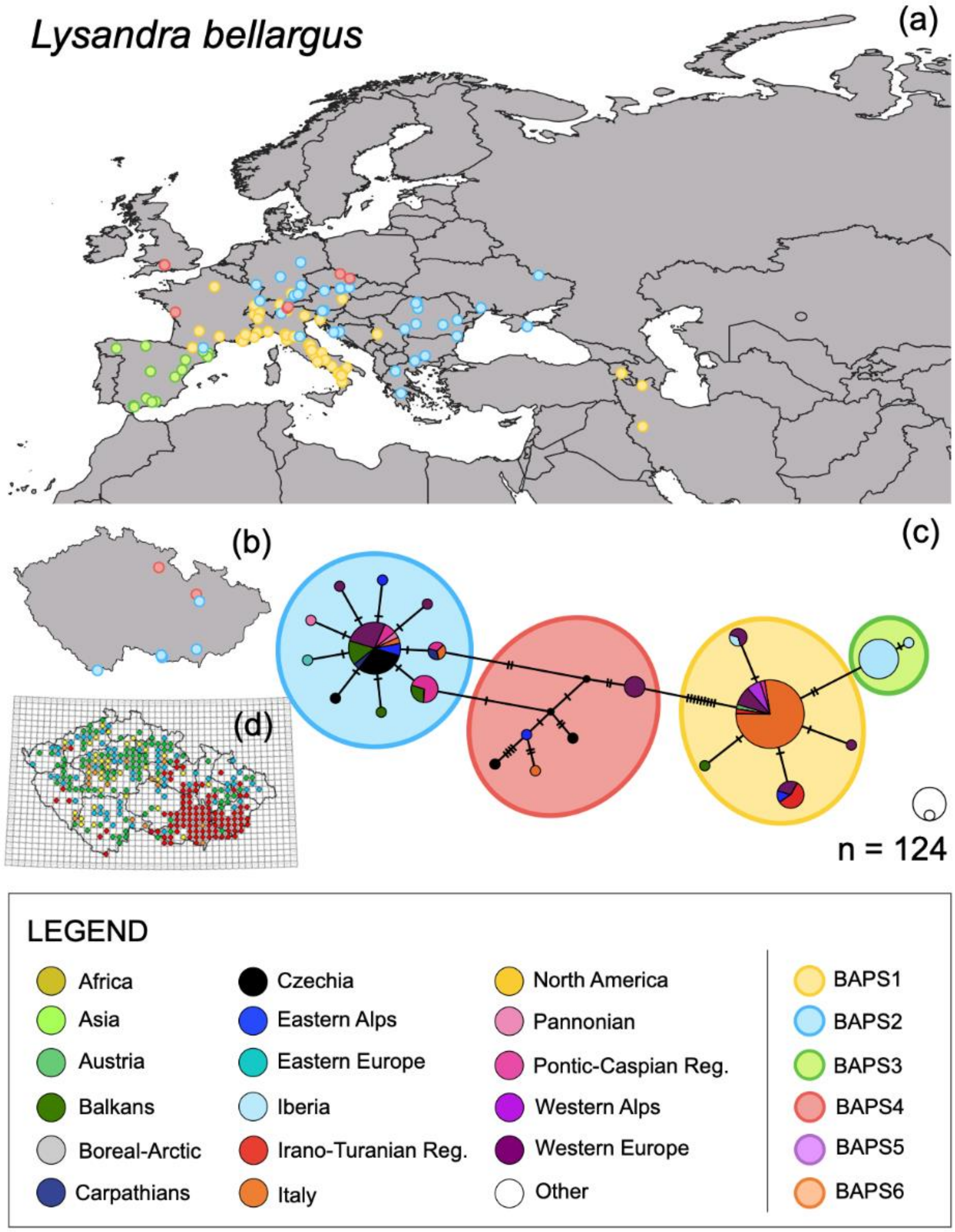


*Lysandra bellargus* (Rottemburg, 1775): **Habitat:** xerothermophilic 1. **Red List:** VU
**BAPS:** BAPS-Italy-Balkans + BAPS-temperate. **HAP:** HAP-Italy-Balkans
**Range:** Pontic-Mediterranean. **Distribution CZ:** Short dry grasslands. Moravian lowlands, scarcer in Bohemia, currently re-expanding. BAPS4 may represent a hybrid or misidentified *L. coridon*.

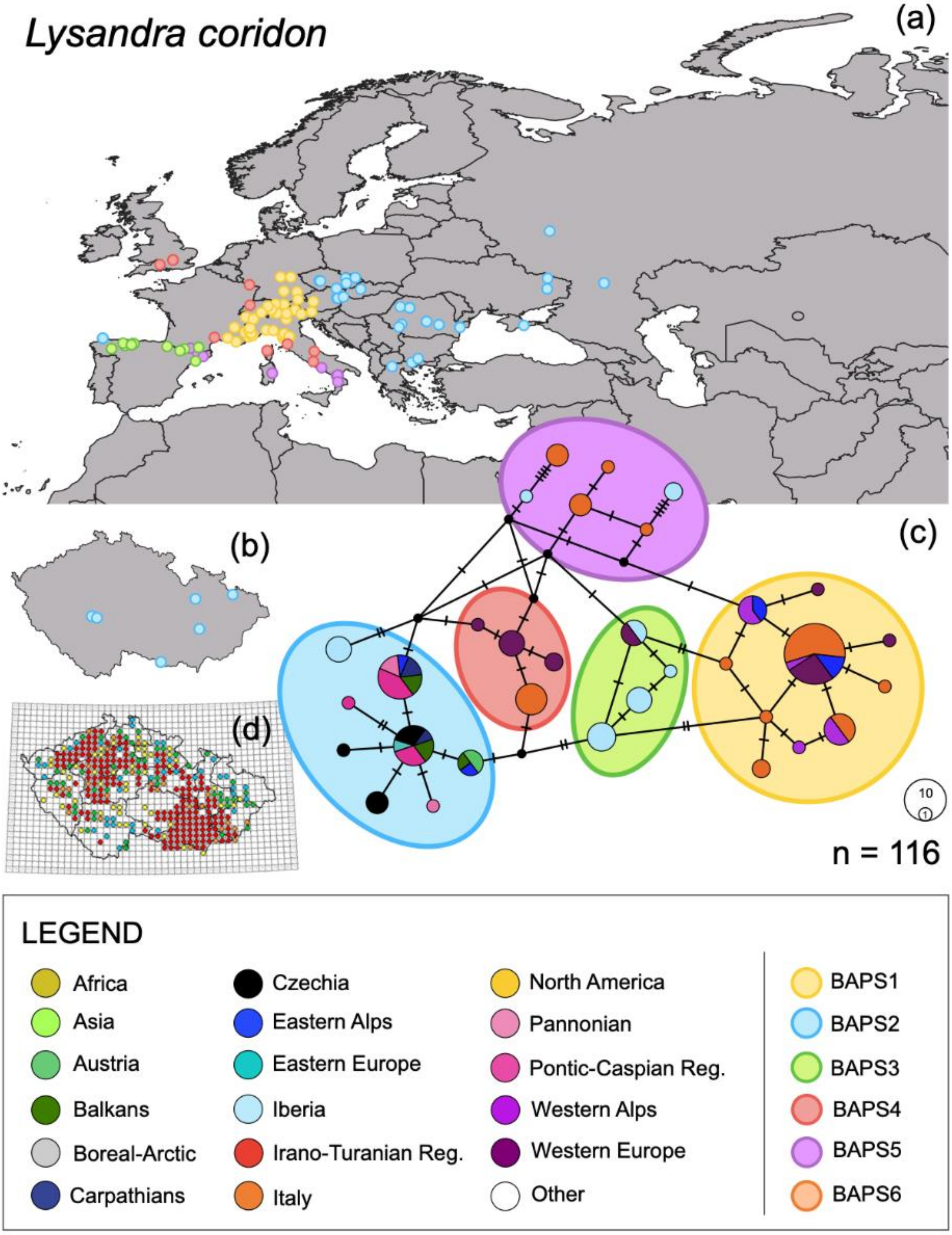


*Lysandra coridon* (Poda, 1761): **Habitat:** xerothermophilic 1. **Red List:** VU

**BAPS:** BAPS-Balkans-East. **HAP:** HAP-Balkans-East

**Range:** European. **Distribution CZ:** dry grasslands and shrublands in warm regions across the country.

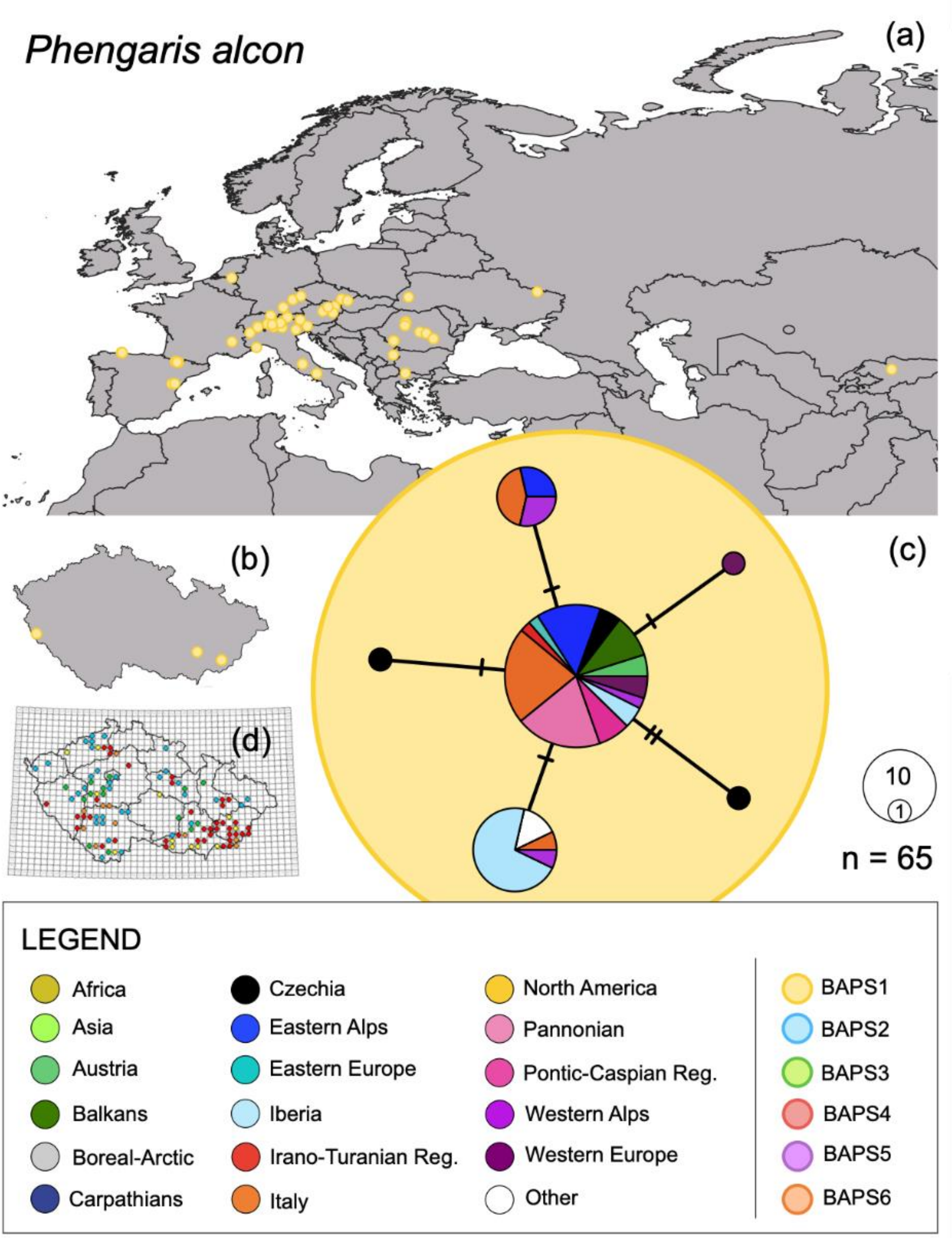


*Phengaris alcon* ([Denis & Schiffermüller], 1775): **Habitat:** mesophilic 1 + hydrophilic (f. *alcon*) + xerothermophilic 1 (f. *rebeli*). **Red List:** CR

**BAPS:** BAPS-low. **HAP:** HAP-low

**Range:** Western Palearctic. **Distribution CZ:** both forms are scarce and critically endangered, surviving in a few last localities across the country.

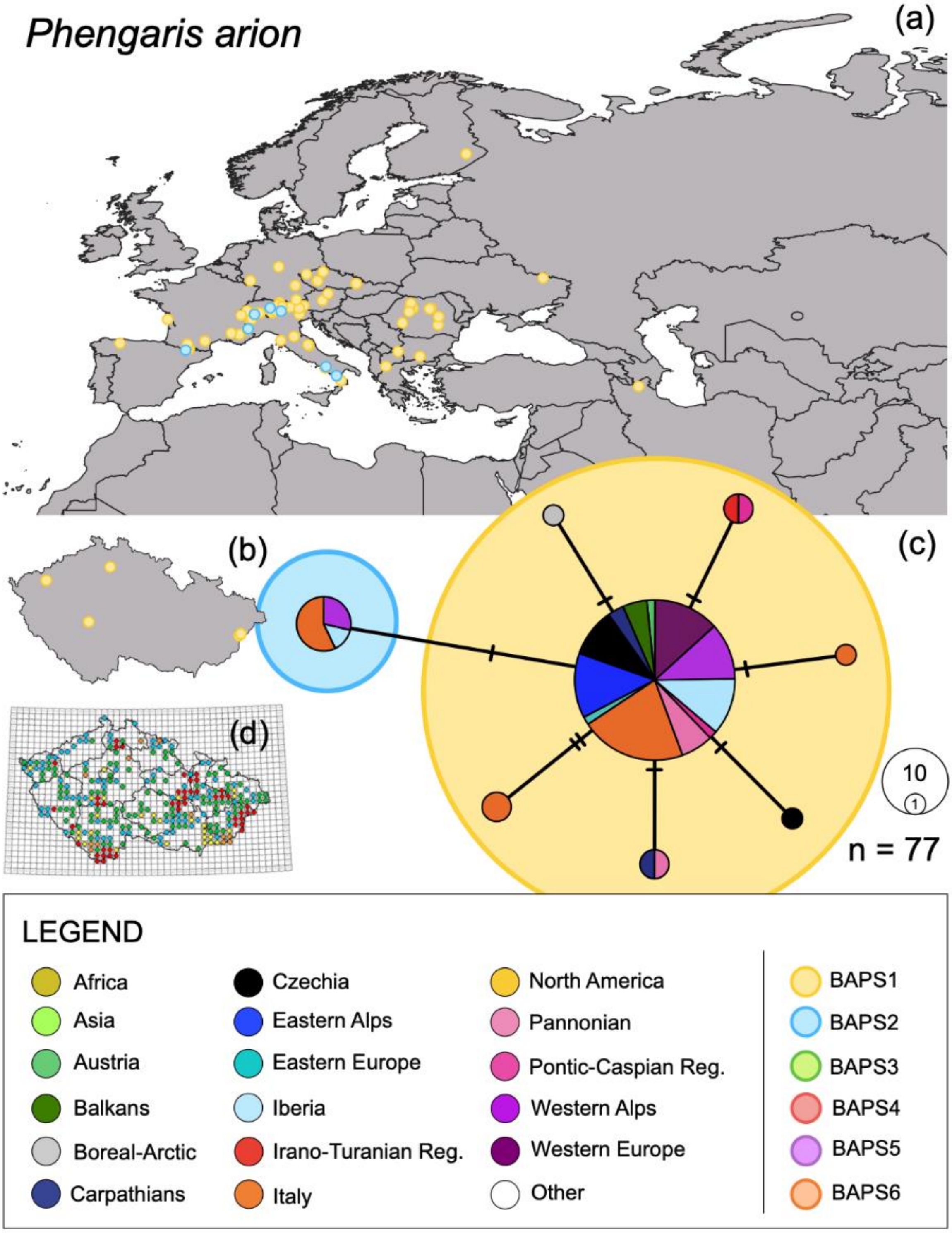


*Phengaris arion* (Linnaeus, 1758): **Habitat:** xerothermophilic 1. **Red List:** EN

**BAPS:** BAPS-wide. **HAP:** HAP-low

**Range:** Palearctic. **Distribution CZ:** short dry grasslands and pastures in both lowlands and piedmont areas. Severely decreased from majority of its localities due to agricultural intensification. Isolated localities across CZ, most viable population system persists in Vsetín District (Spitzer, Benes, Dandova, Jaskova, & Konvicka, 2009).

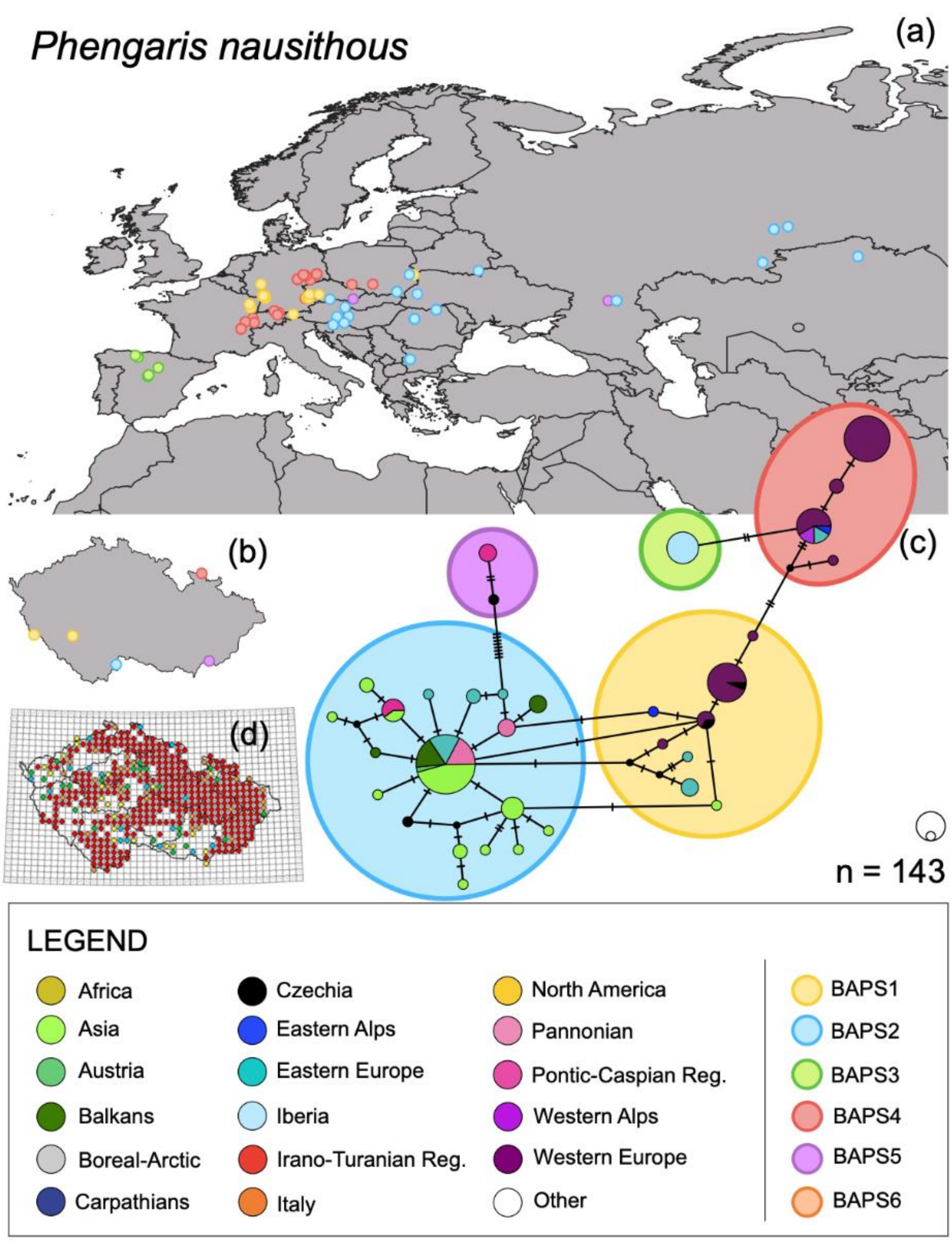


*Phengaris nausithous* (Bergsträsser, 1779): **Habitat:** hydrophilic. **Red List:** NT

**BAPS:** BAPS-mix. **HAP:** HAP-mix

**Range:** Western Palearctic. **Distribution CZ:** wet meadows, road margins with the host plant great burnet (*Sanguisorba officinalis*). Widespread across the country, including some urban regions.

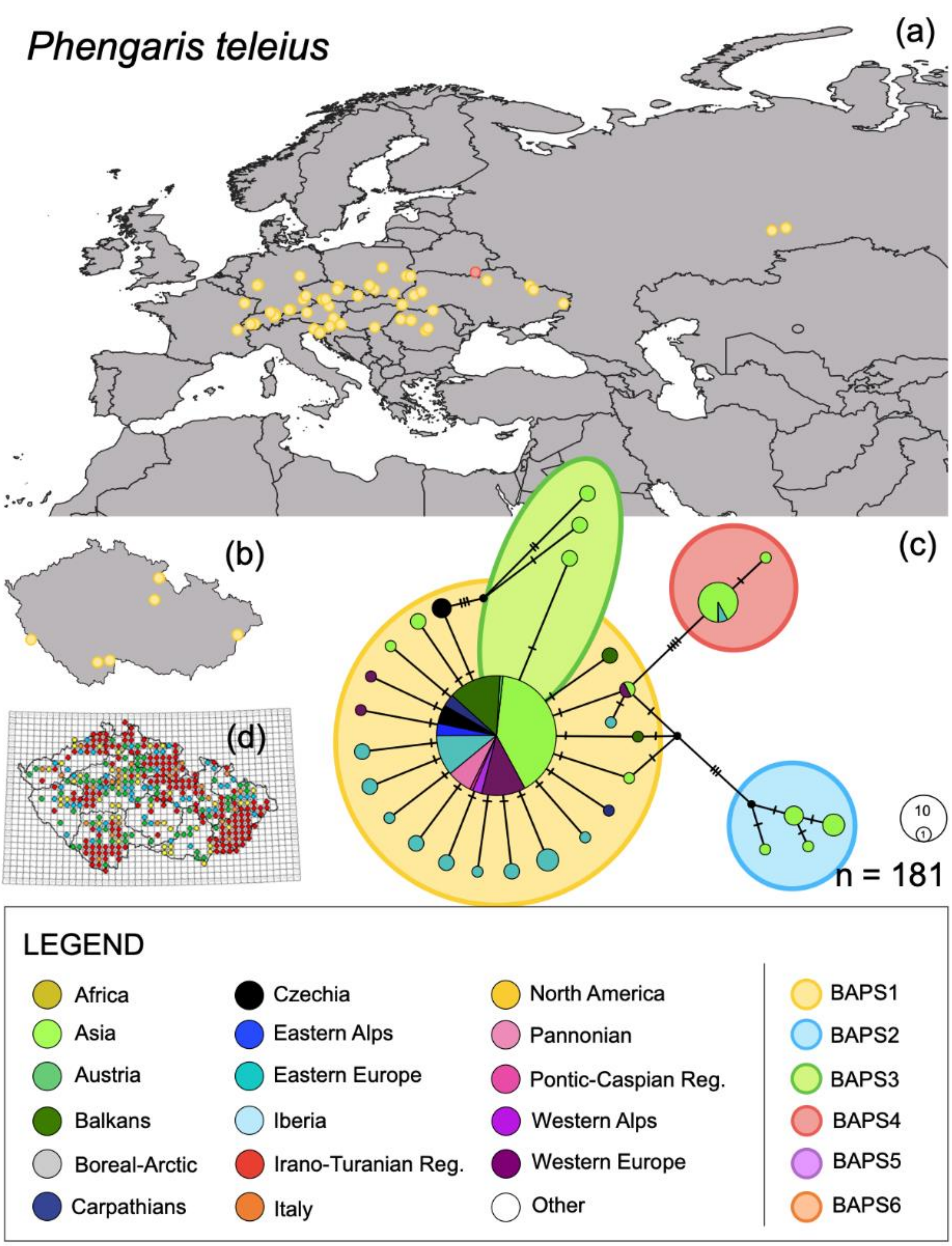


*Phengaris teleius* (Bergsträsser, 1779): **Habitat:** hydrophilic. **Red List:** VU

**BAPS:** BAPS-temperate. **HAP:** HAP-temperate (such distribution)

**Range:** Palearctic. **Distribution CZ:** non-intensively managed wet meadows with great burnet. Scattered distribution across the country, but decreased with drainage and afforestation activities.

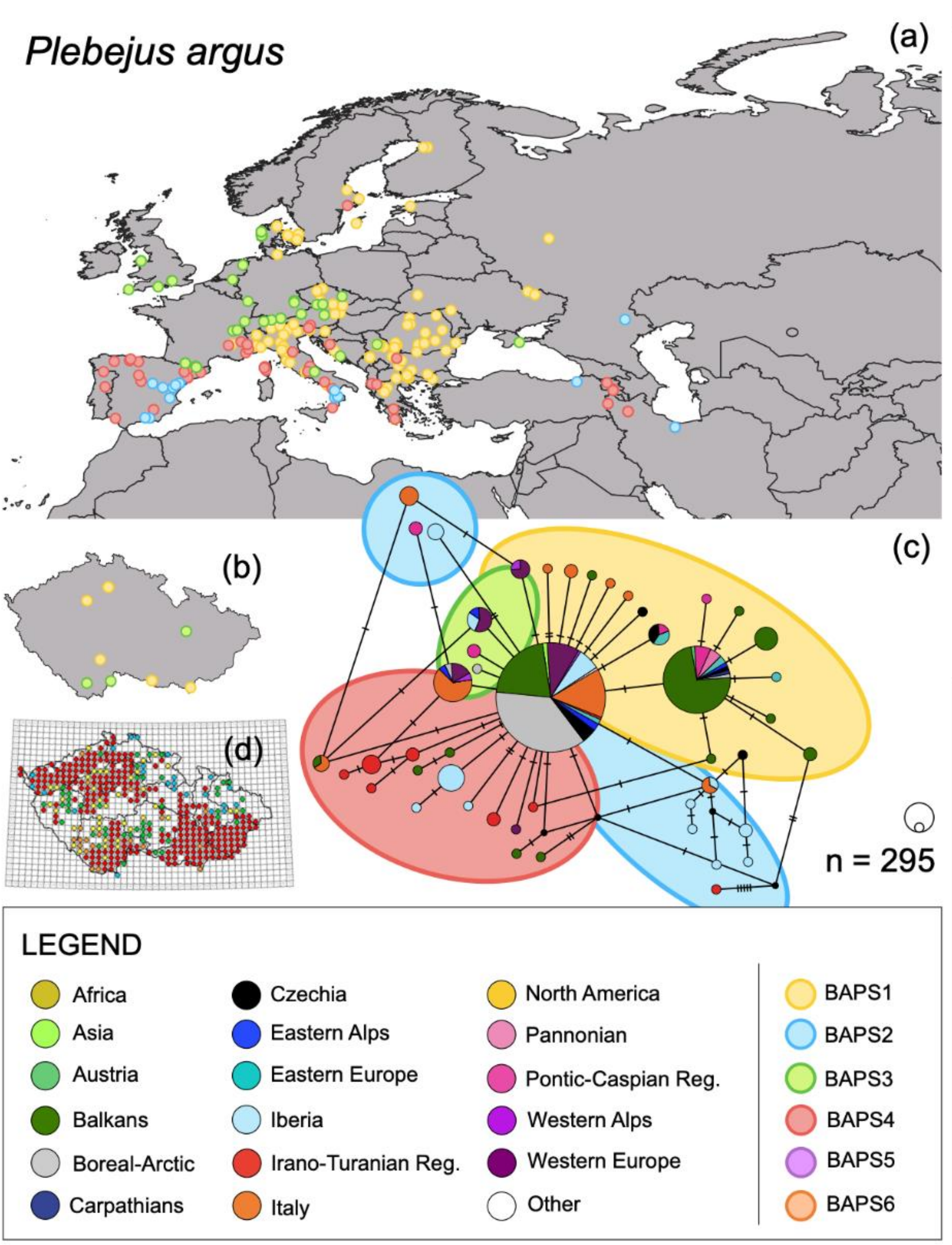


*Plebejus argus* (Linnaeus, 1758): **Habitat:** xerothermophilic 1. **Red List:** NT

**BAPS:** BAPS-wide + BAPS-Italy-Balkans. **HAP:** HAP-wide-complex

**Range:** Palearctic. **Distribution CZ:** dry grasslands, quarries, open sandy woodlands, and heaths. Widespread across the country, but retreated with traditional land use abandonment.

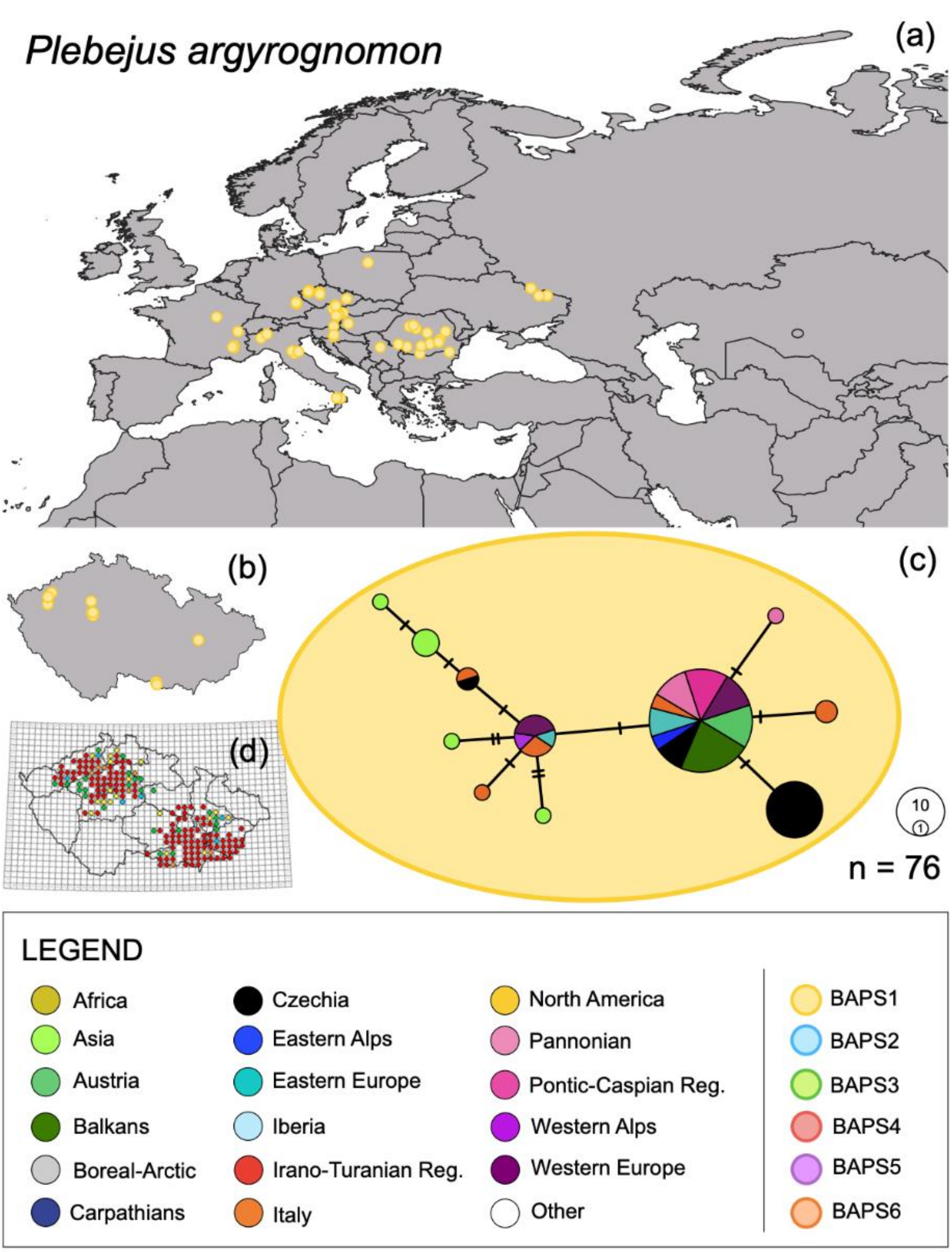


*Plebejus argyrognomon* (Bergsträsser, 1779): **Habitat:** xerothermophilic 1. **Red List:** LC

**BAPS:** BAPS-low. **HAP:** HAP-Italy-Balkans (such distribution)

**Range:** European. **Distribution CZ:** dry grasslands in thermophilic regions of Bohemia and Moravian lowlands.

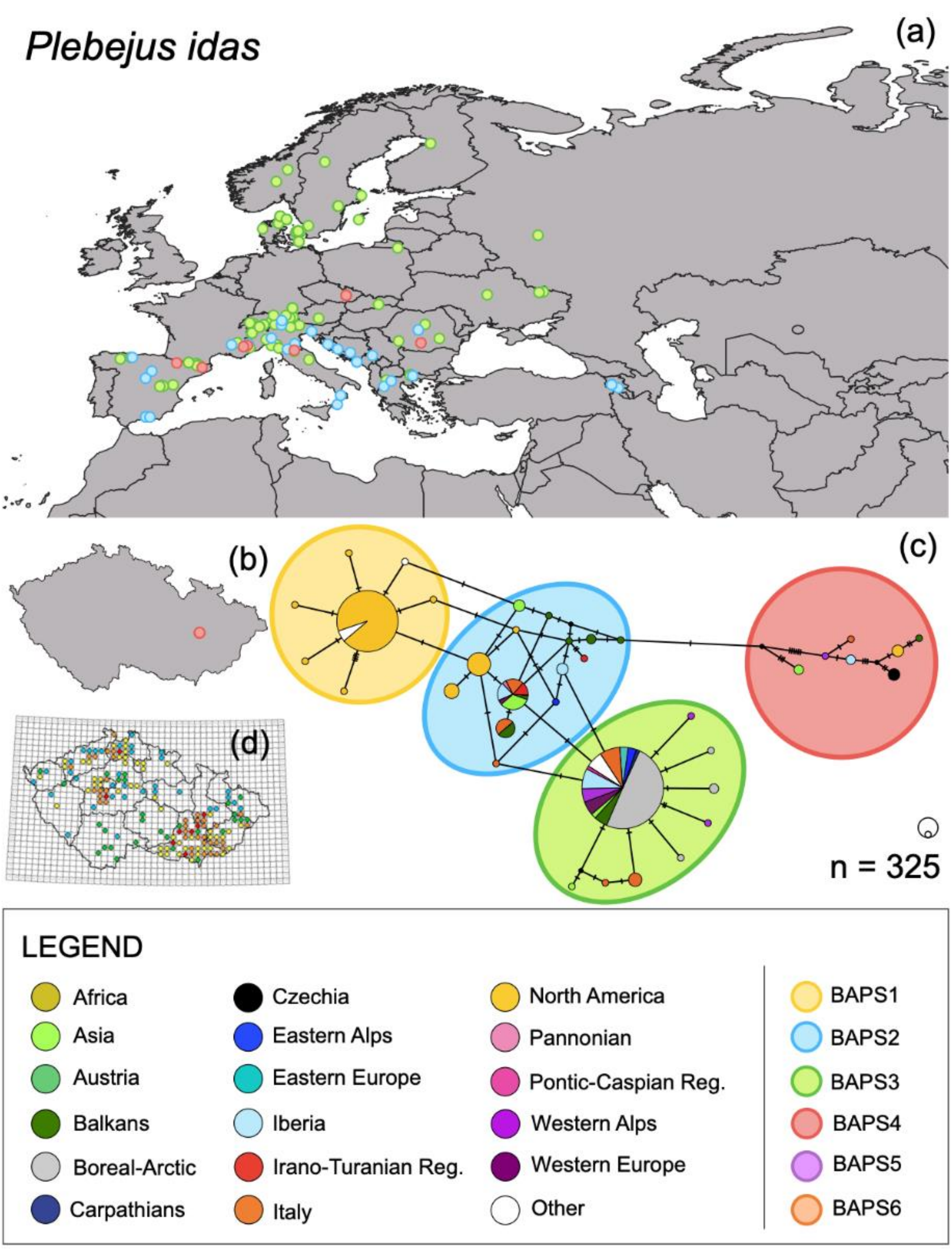


*Plebejus idas* (Linnaeus, 1760): **Habitat:** xerothermophilic 1. **Red List:** CR

**BAPS:** BAPS-wide. **HAP:** HAP-Central

**Range:** Holarctic. **Distribution CZ:** dry grasslands and heaths. In the past, scattered across the country. Currently extinct from majority of its former localities, surviving in a few last localities in north Bohemia and central Moravia.

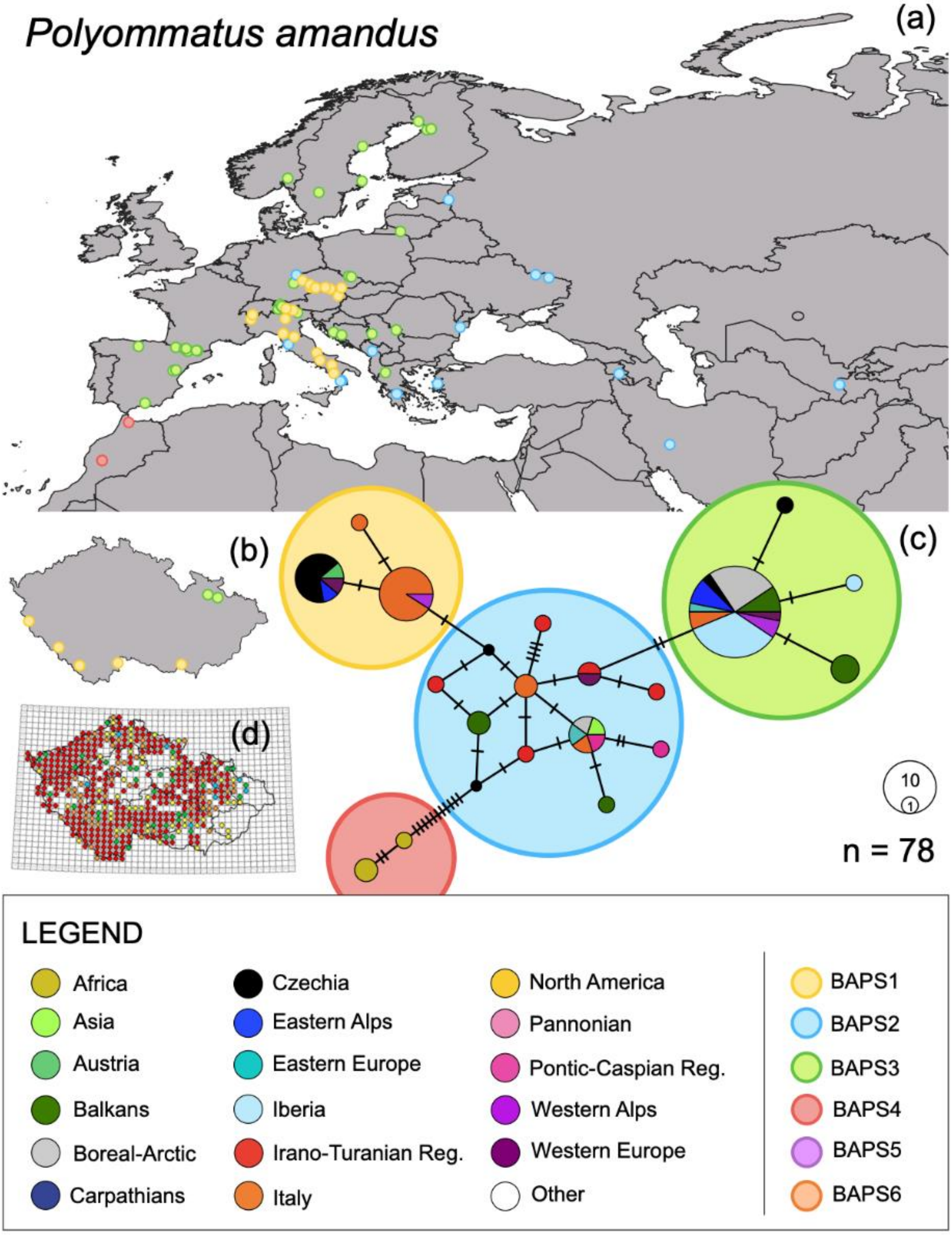


*Polyommatus amandus* (Schneider, 1792): **Habitat:** mesophilic 1 + tyrphophilic. **Red List:** NT

**BAPS:** BAPS-wide + BAPS-Italy-Balkans. **HAP:** HAP-wide-low + HAP-Italy-Balkans

**Range:** European. **Distribution CZ:** flowery meadows and pastures, also colder habitats including bogs, often in piedmont areas. Widespread, but missing from east of the country.

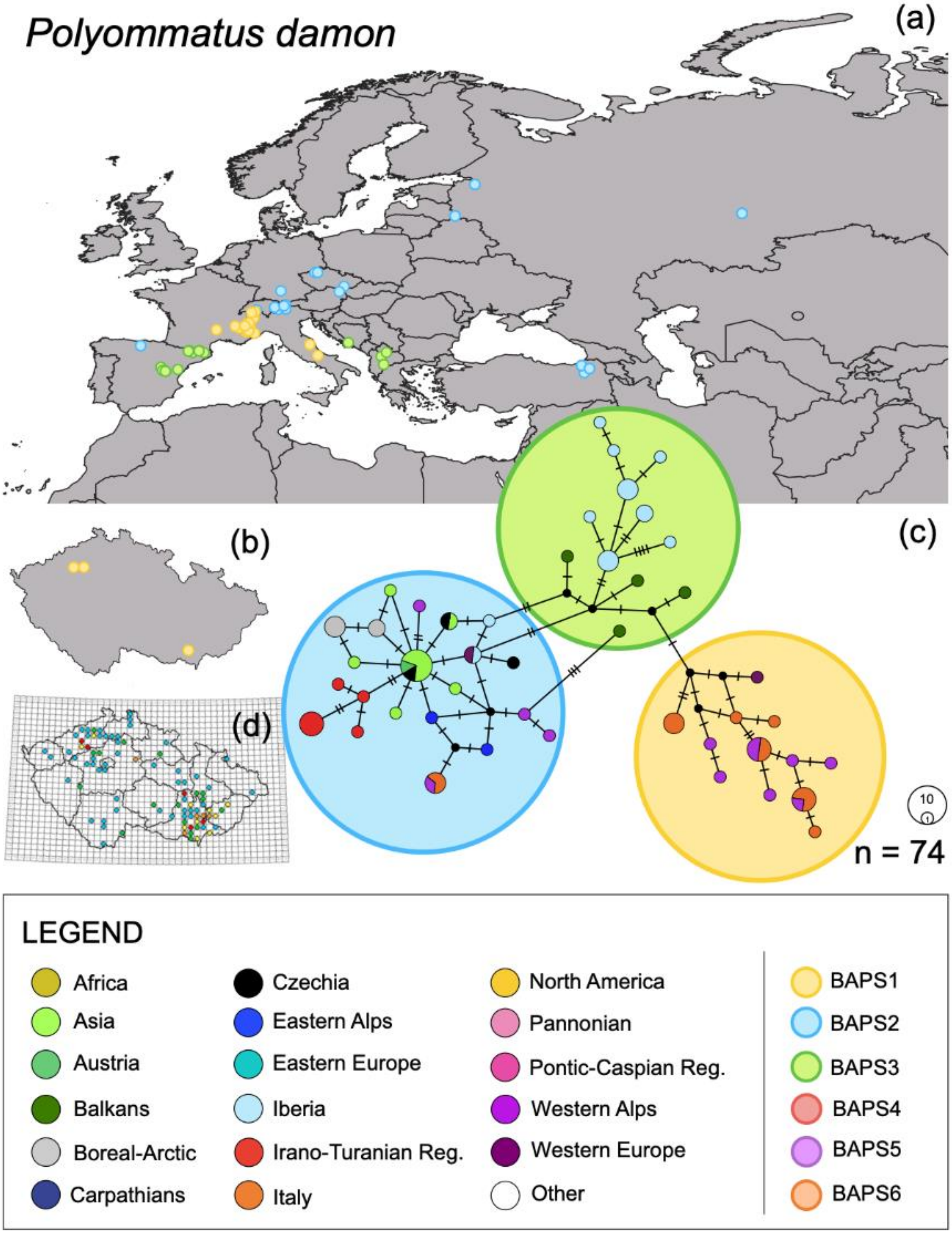


*Polyommatus damon* ([Denis & Schiffermüller], 1775): **Habitat:** xerothermophilic 1. **Red List:** CR

**BAPS:** BAPS-temperate. **HAP:** HAP-temperate

**Range:** Palearctic (reaching Mongolia). **Distribution CZ:** dry grasslands. Very close to extinction on its two last localities (Central Bohemian Uplands and South Moravia), only very few individuals or eggs recorded in recent years (Sucháčková Bartoňová, Konvička, Marešová, Bláhová, et al., 2021).

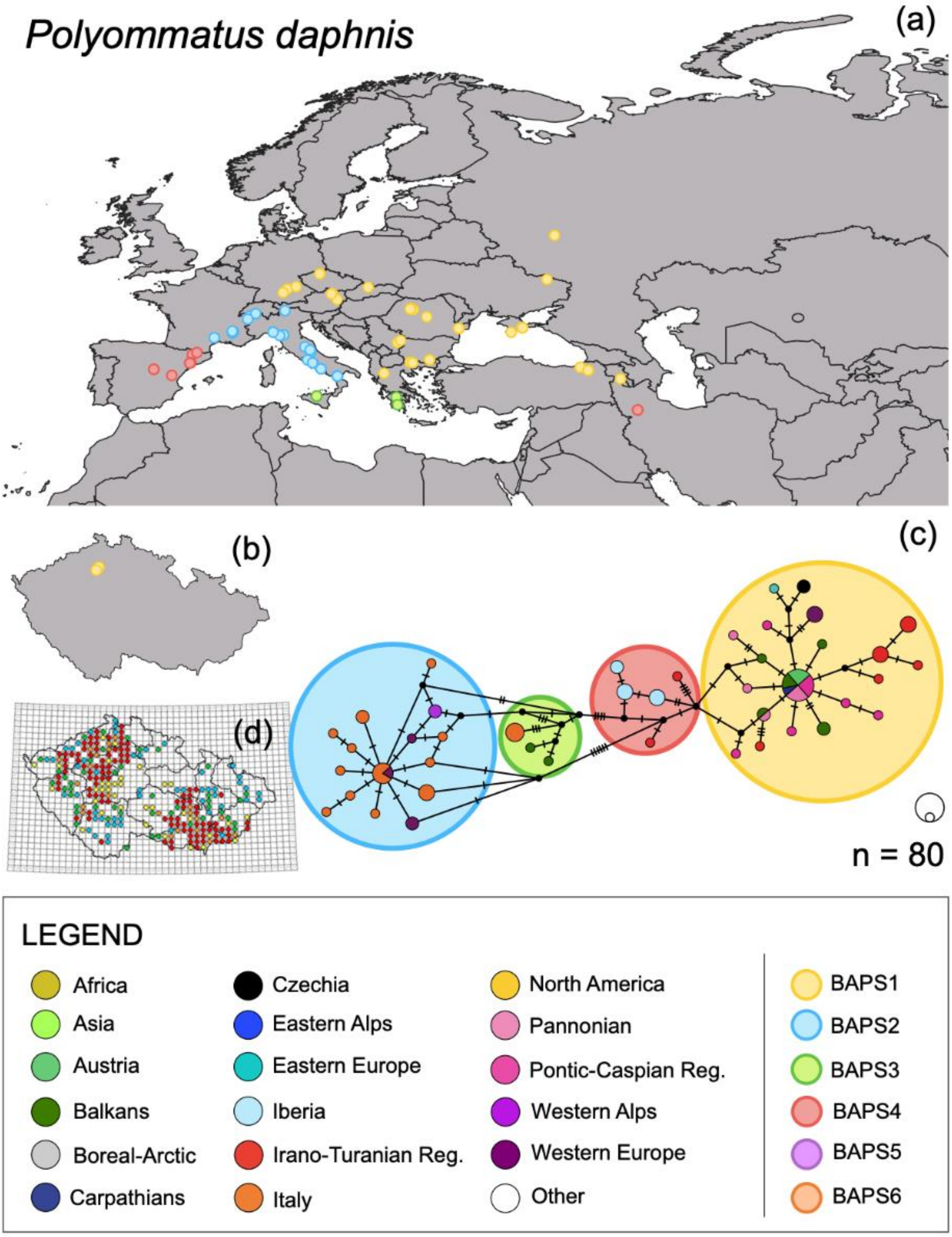


*Polyommatus daphnis* ([Denis & Schiffermüller], 1775): **Habitat:** xerothermophilic 1. **Red List:** VU

**BAPS:** BAPS-East-Balkans. **HAP:** HAP-temperate.

**Range:** Pontic-Mediterranean. **Distribution CZ:** dry grasslands in warm areas across the country, extinct from several regions.

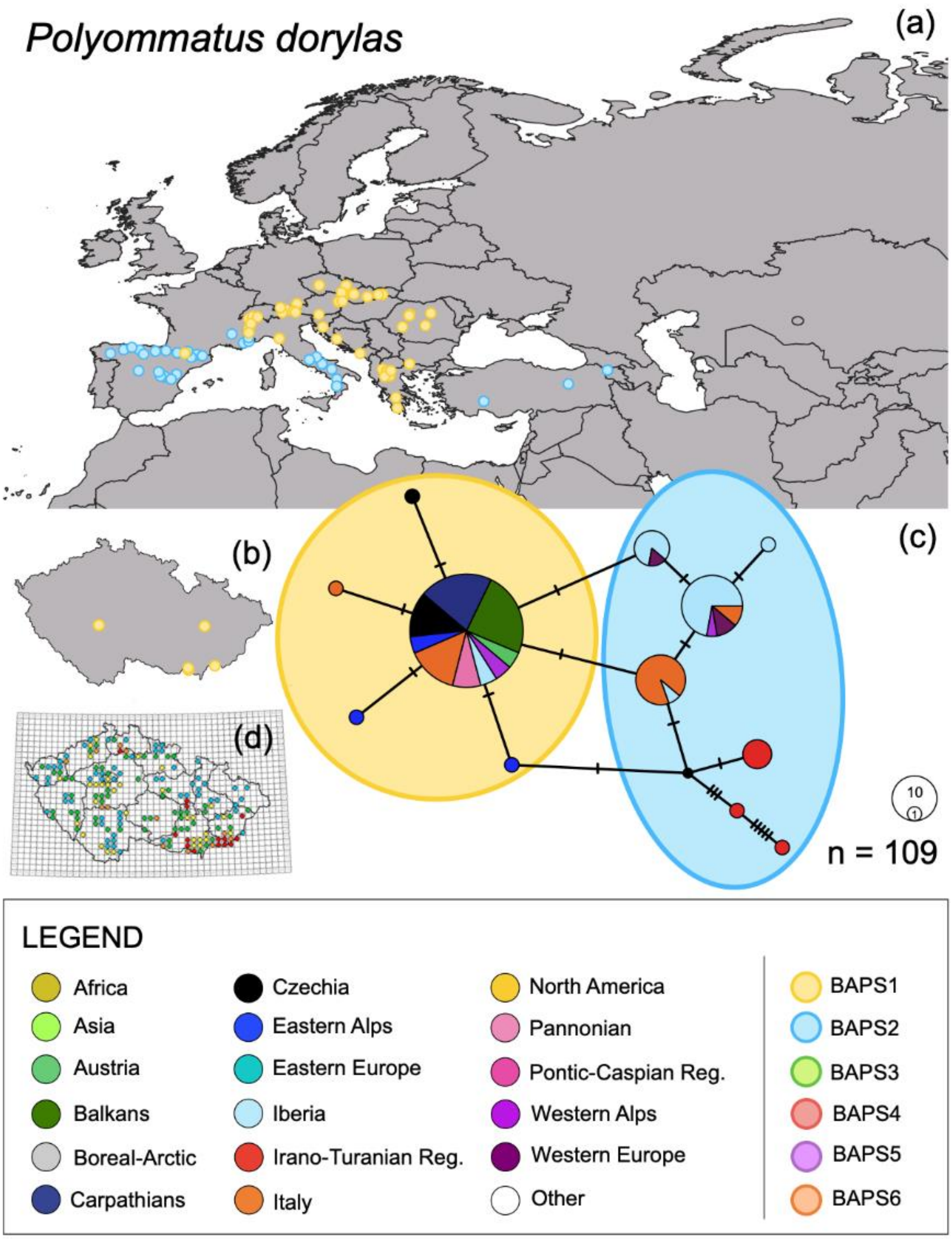


*Polyommatus dorylas* ([Denis & Schiffermüller], 1775): **Habitat:** xerothermophilic 1. **Red List:** CR

**BAPS:** BAPS-Italy-Balkans. **HAP:** HAP-Italy-Balkans

**Range:** European. **Distribution CZ:** dry grasslands and scrublands. In the past, distributed across all warm areas, currently a few small populations are surviving in the White Carpathians. Reintroduced to several areas from the White Carpathians (e.g., Týnčany Karst), unauthorized transfers elsewhere.

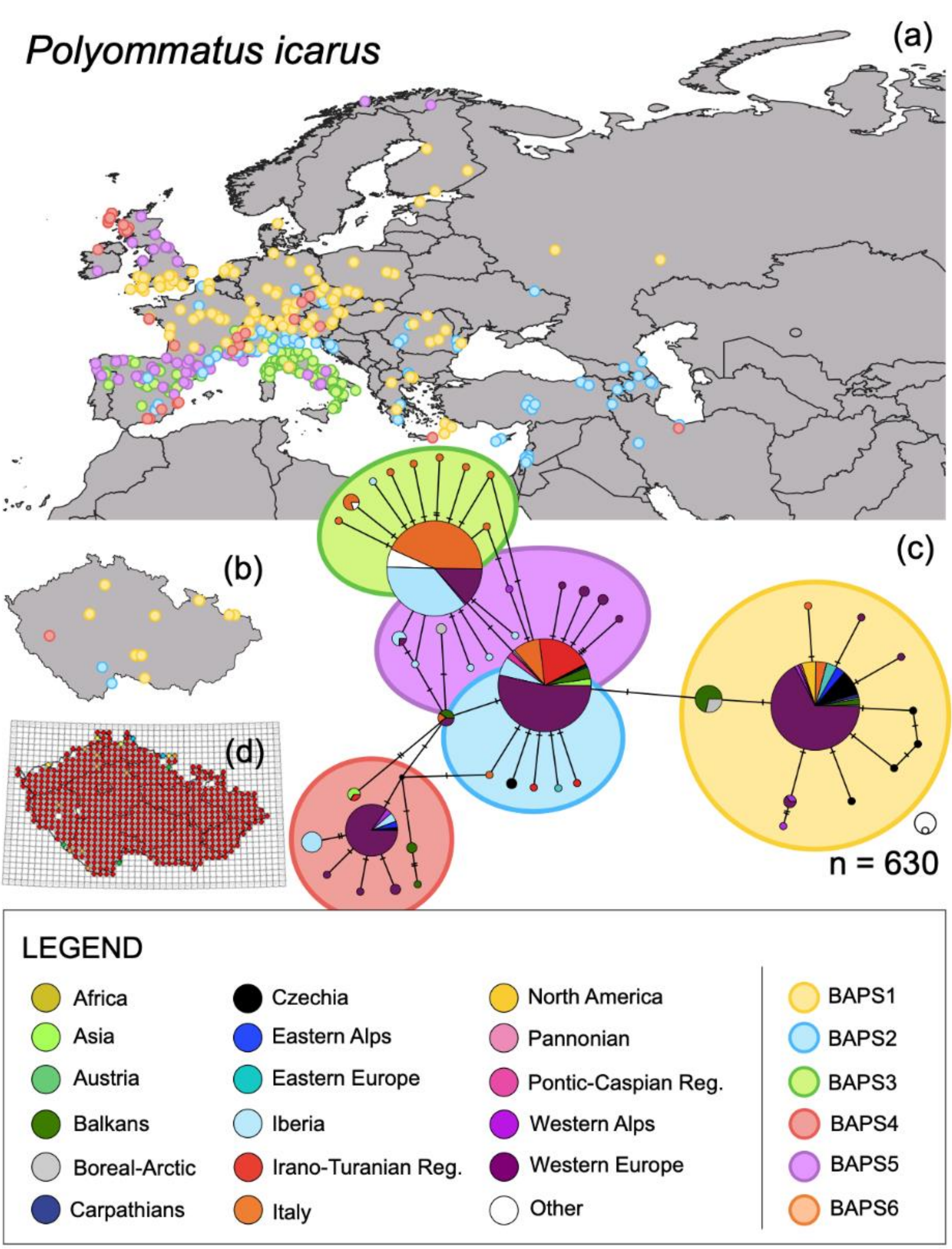


*Polyommatus icarus* (Rottemburg, 1775): **Habitat:** ubiquitous. **Red List:** LC

**BAPS:** BAPS-mix. **HAP:** HAP-mix

**Range:** Palearctic. **Distribution CZ:** various open habitats. Widespread and common.

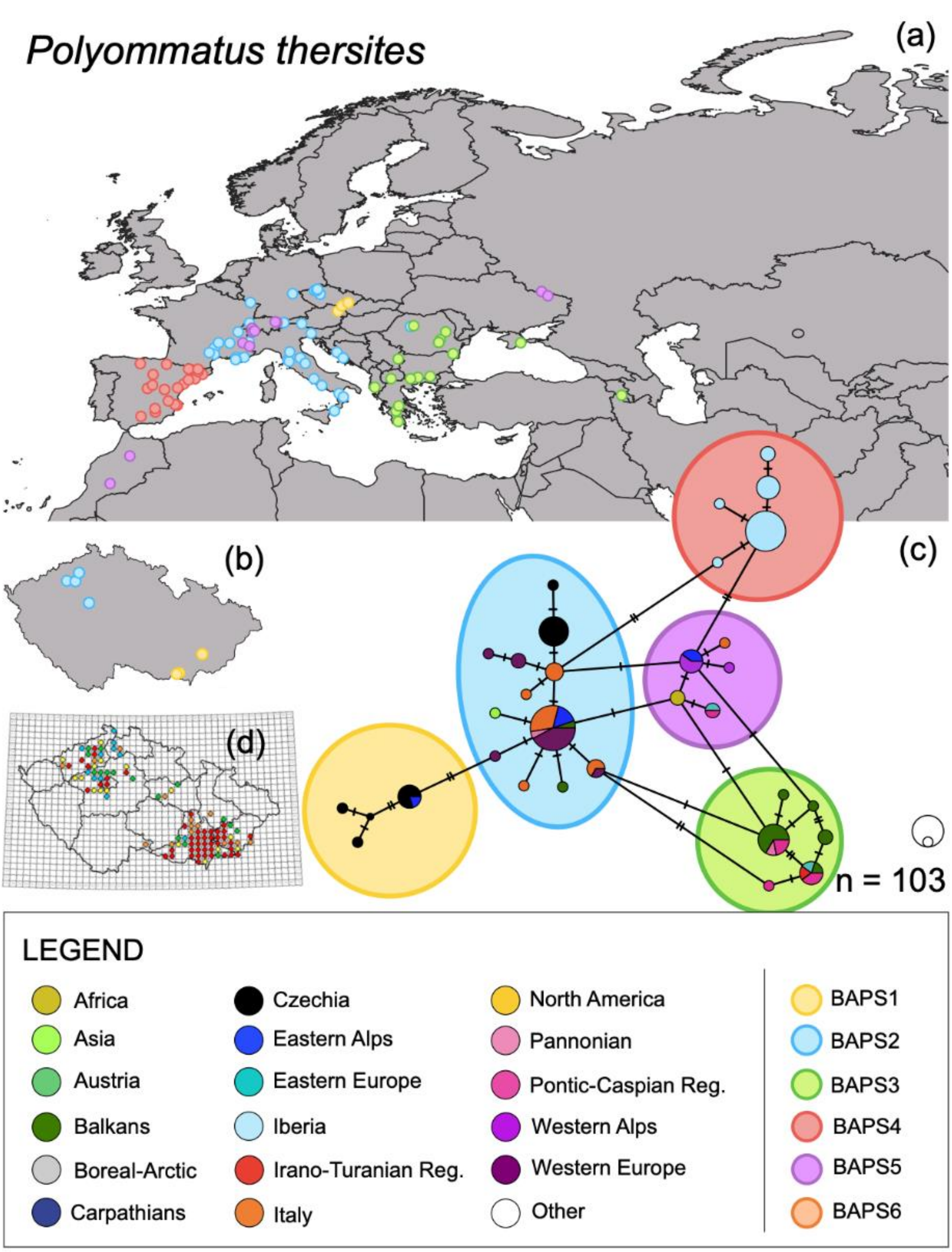


*Polyommatus thersites* (Cantener, 1835): **Habitat:** xerothermophilic 1. **Red List:** VU
**BAPS:** BAPS-Italy-Balkans + BAPS-Central. **HAP:** HAP-Italy-Balkans + HAP-Central
**Range:** Western Palearctic. **Distribution CZ:** rocky steppes and forest steppes in thermophilic regions of Bohemia and Moravia, decreasing.

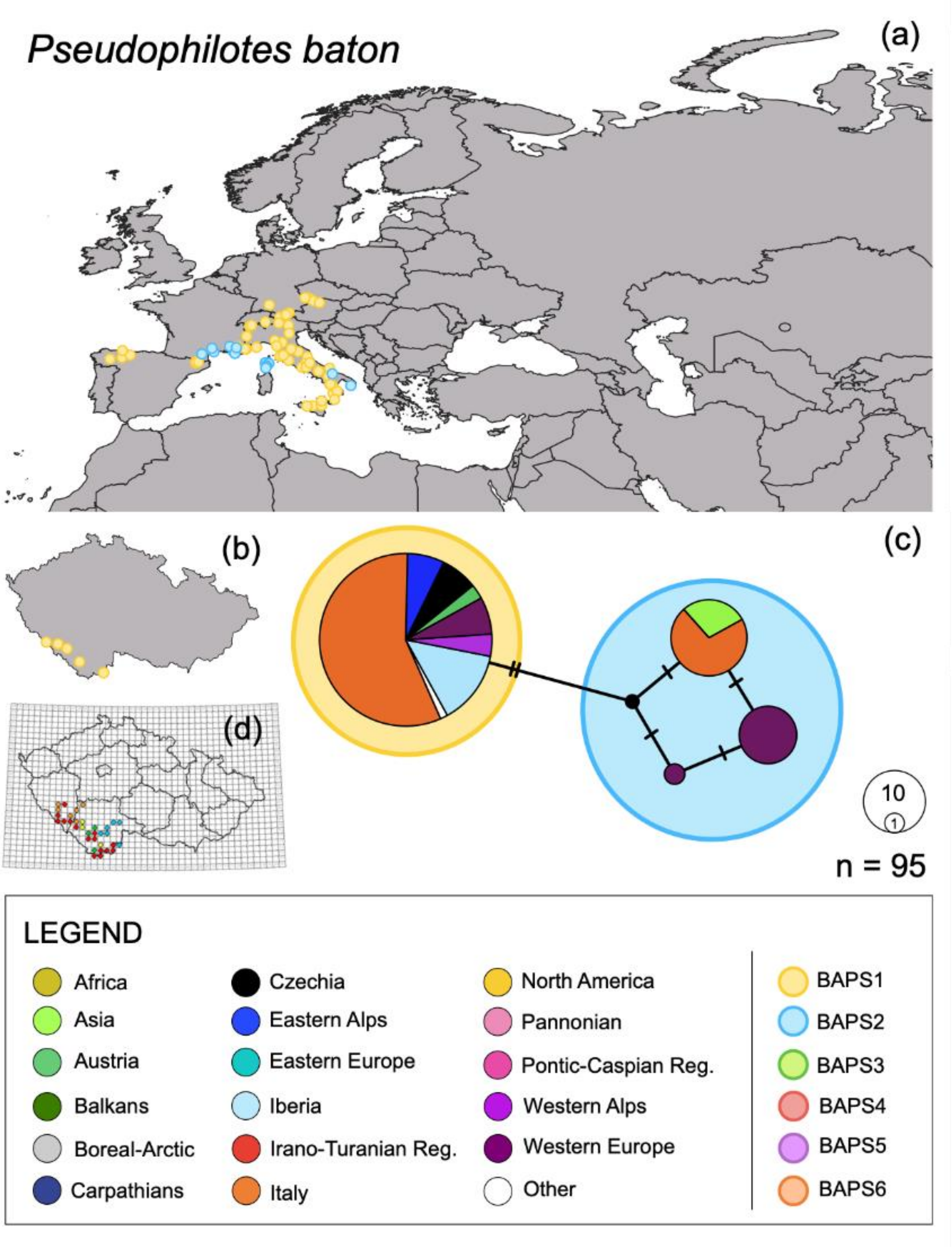


*Pseudophilotes baton* (Bergsträsser, 1779): **Habitat:** xerothermophilic 1. **Red List:** CR

**BAPS:** BAPS-West. **HAP:** HAP-West (such distribution)

**Range:** Atlantic-Mediterranean. **Distribution CZ:** dry short grassland. Historically distributed only in South and southeast Bohemia, but extinct from majority of its former sites. Only few individuals observed in recent years. Across Eurasia, several species of *Pseudophilotes* share haplotypes. The mitochondrial lineage contact zone of *P. baton* with *P. vicrama* stretches through Croatia and lowland Austria, where individuals of *P. vicrama* with the barcode of *P. baton* occur (Sucháčková Bartoňová, Konvička, Marešová, Wiemers, et al., 2021).

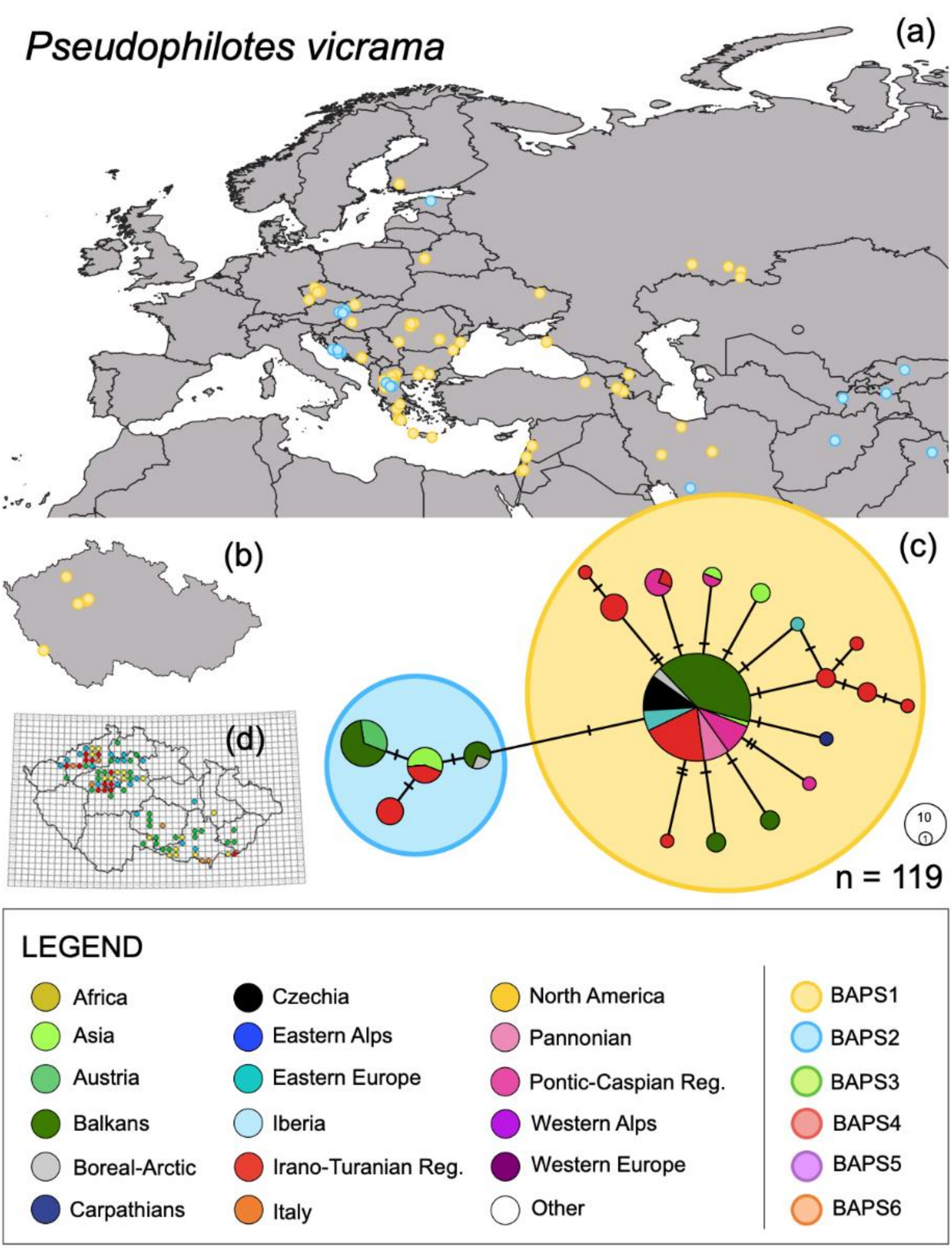


*Pseudophilotes vicrama* (Moore, 1865): **Habitat:** xerothermophilic 1. **Red List:** CR

**BAPS:** BAPS-East-Balkans. **HAP:** HAP-East-Balkans (such distribution)

**Range:** Western Palearctic. **Distribution CZ:** dry short grasslands and steppes in thermophilic regions, extinct from Moravia, only a few sites remain in central Bohemia. Decreased with successional changes.

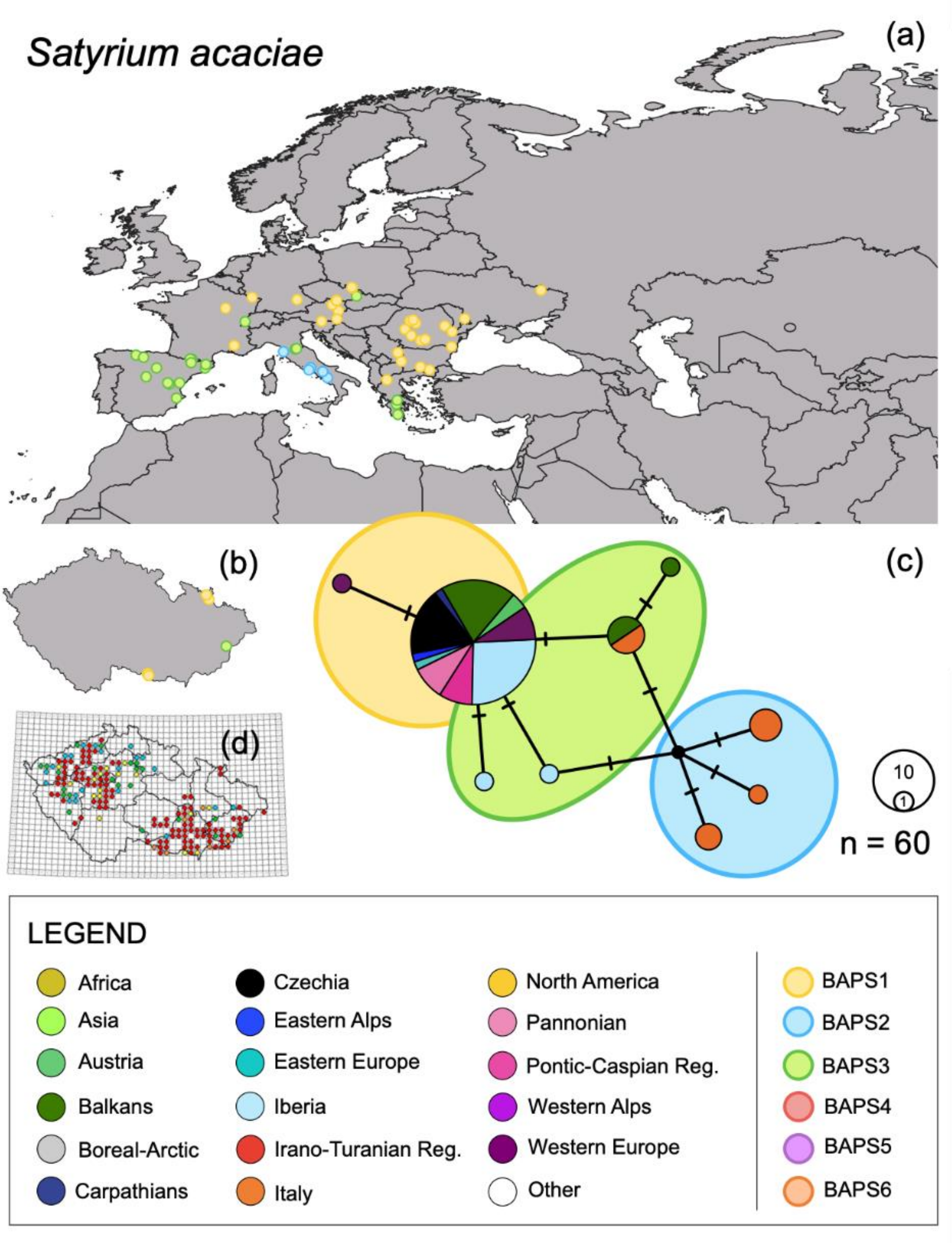


*Satyrium acaciae* (Fabricius, 1787): **Habitat:** xerothermophilic 2. **Red List:** LC

**BAPS:** BAPS-mix. **HAP:** HAP-wide-low

**Range:** Pontic-Mediterranean. **Distribution CZ:** warm regions of the country, dry warm shrublands. Expanding in recent years.

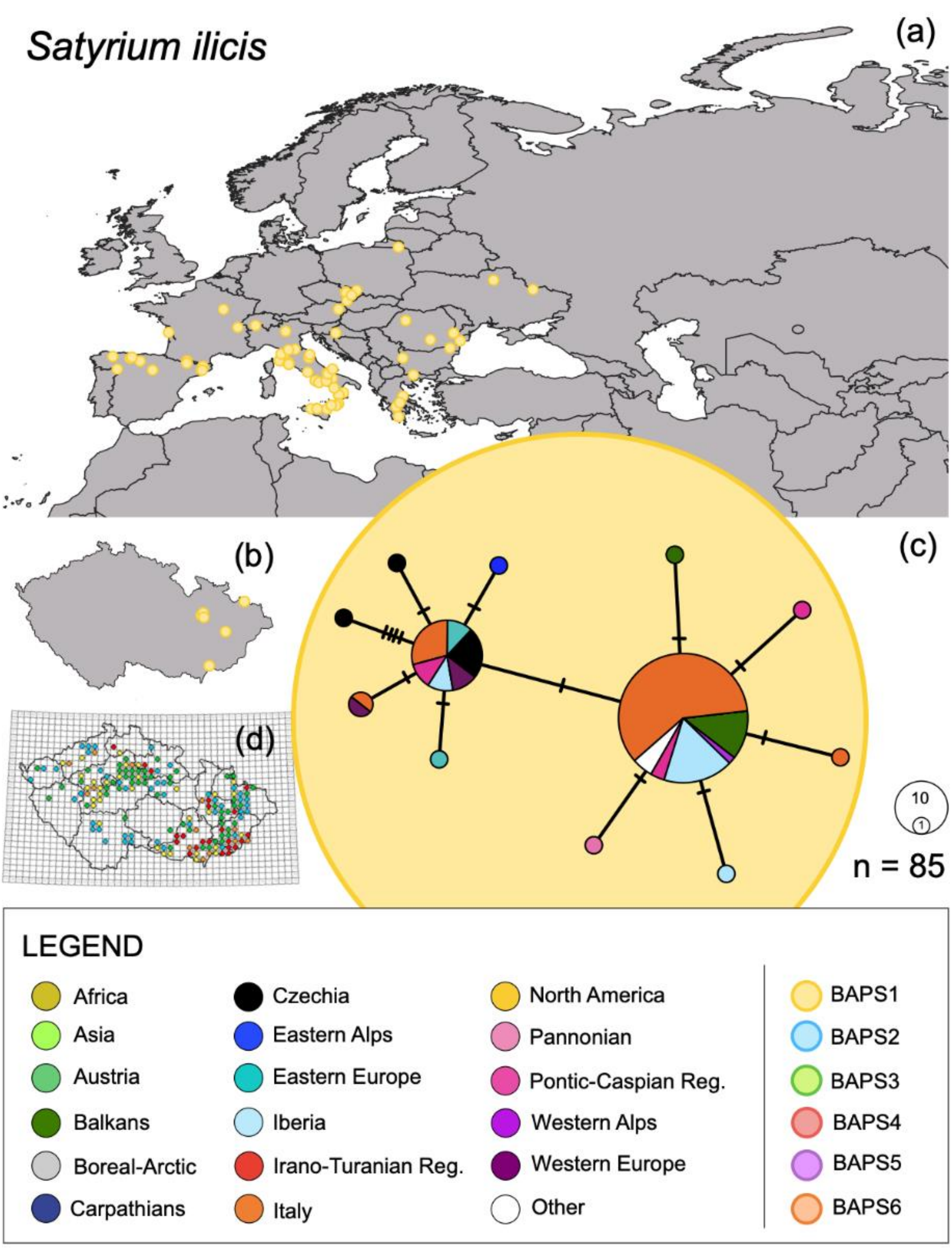


*Satyrium ilicis* (Esper, 1779): **Habitat:** mesophilic 2 + xerothermophilic 2. **Red List:** EN

**BAPS:** BAPS-low. **HAP:** HAP-wide-low

**Range:** Western Palearctic. **Distribution CZ:** woodland openings, scrublands. Oak woodlands in lowlands and highlands, extinct from several regions (loss of open woodlands).

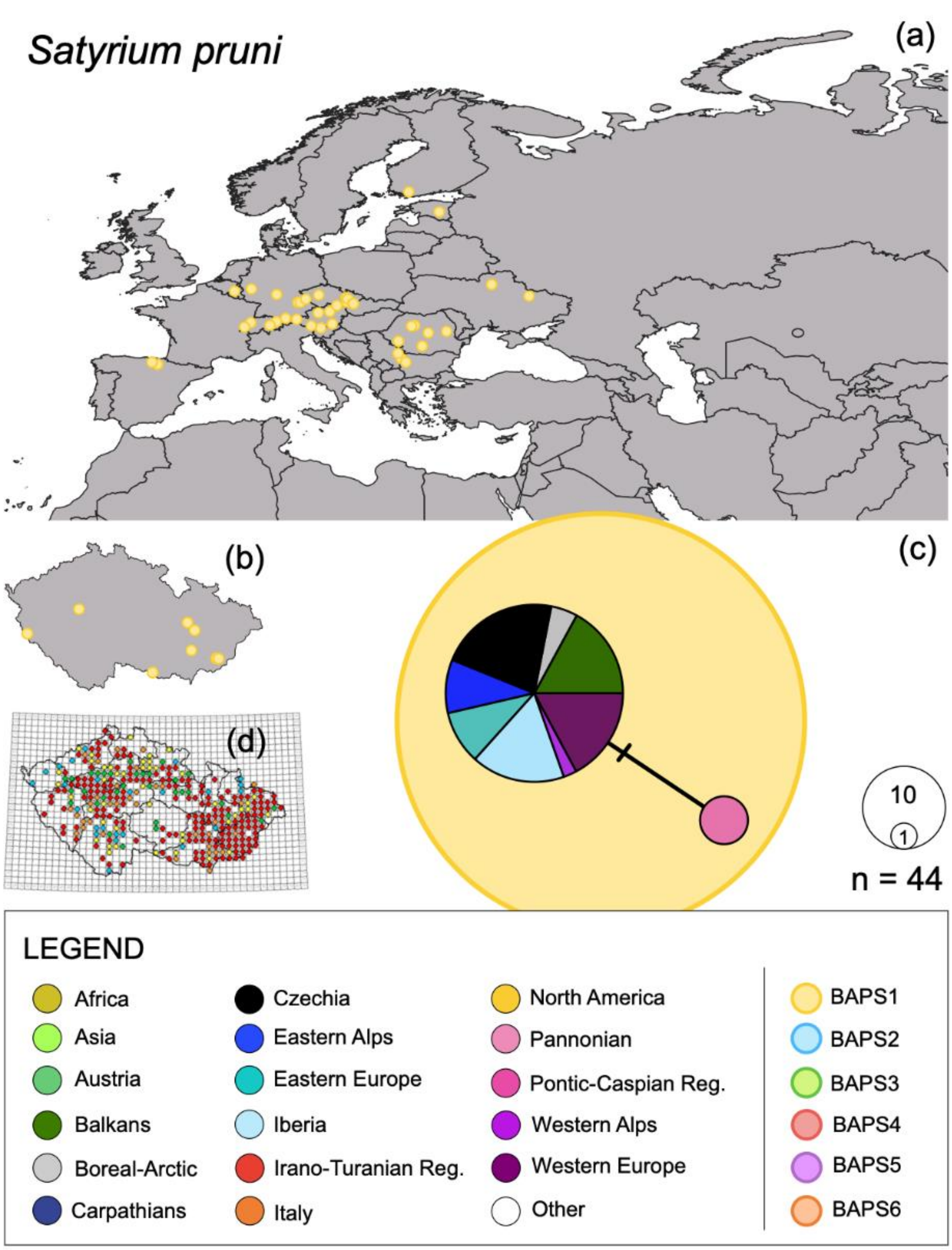


*Satyrium pruni* (Linnaeus, 1758): **Habitat:** mesophilic 2 + xerothermophilic 2. **Red List:** NT
**BAPS:** BAPS-low. **HAP:** HAP-low
**Range:** Palearctic. **Distribution CZ:** scrublands in lowlands and highlands across the country.

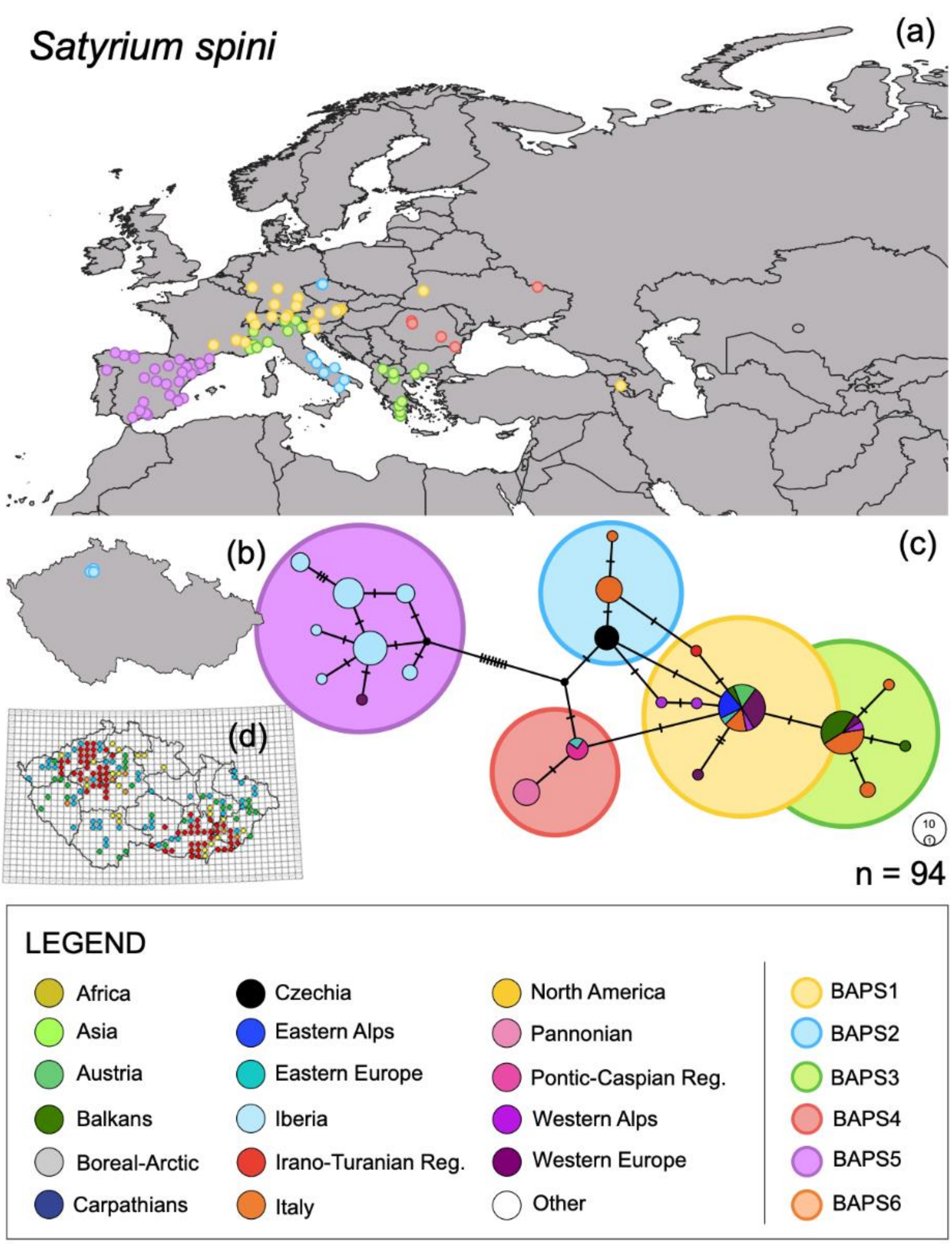


*Satyrium spini* ([Denis & Schiffermüller], 1775): **Habitat:** xerothermophilic 2. **Red List:** VU
**BAPS:** BAPS-Italy-Balkans. **HAP:** HAP-Italy-Balkans
**Range:** Western Palearctic. **Distribution CZ:** dry warm scrublands in thermophilic regions of Bohemia and Moravia.

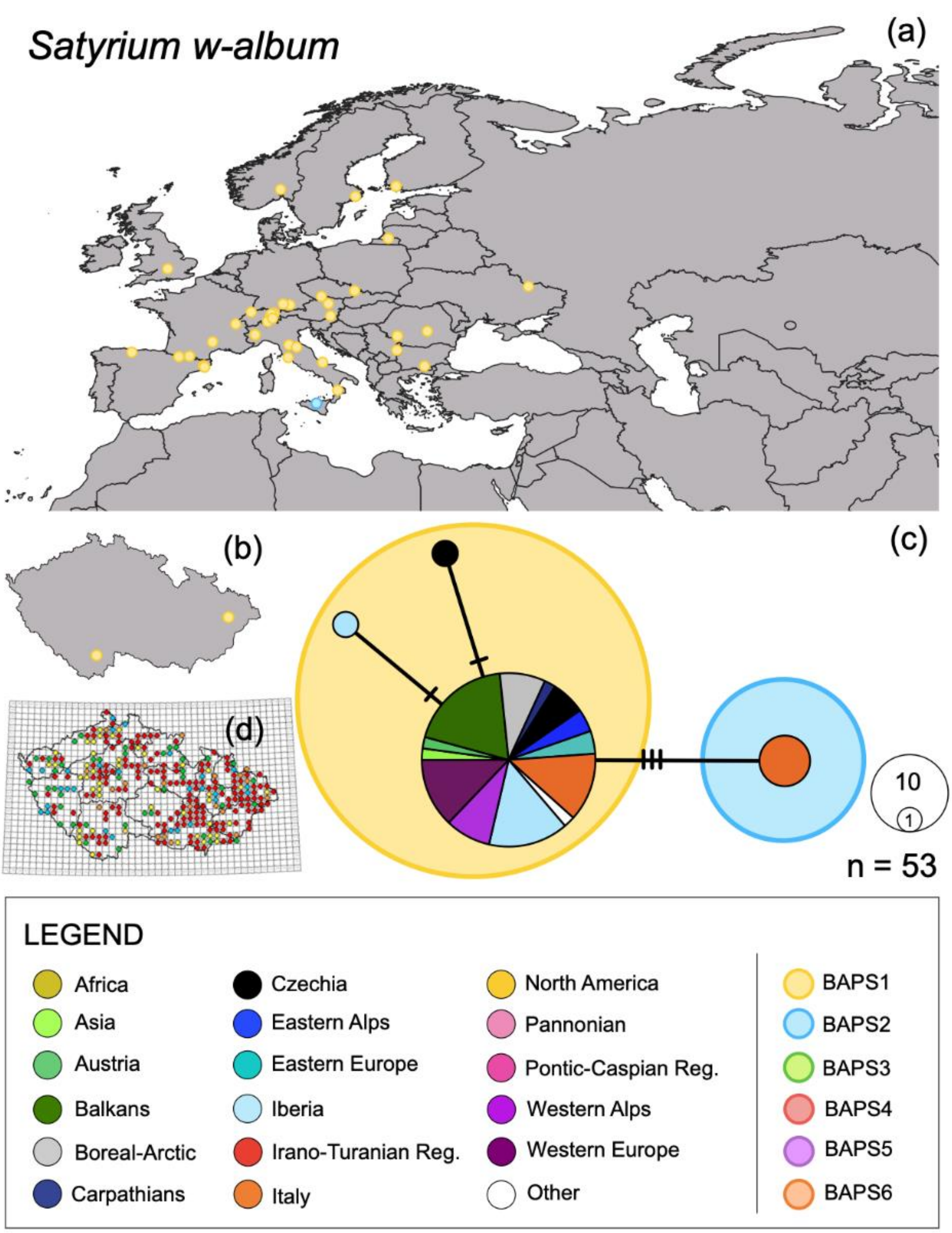


*Satyrium w-album* (Knoch, 1782): **Habitat:** mesophilic 3. **Red List:** NT

**BAPS:** BAPS-wide. **HAP:** HAP-wide-low

**Range:** Palearctic. **Distribution CZ:** various woodland margins and river banks. Widespread across the country but rare.

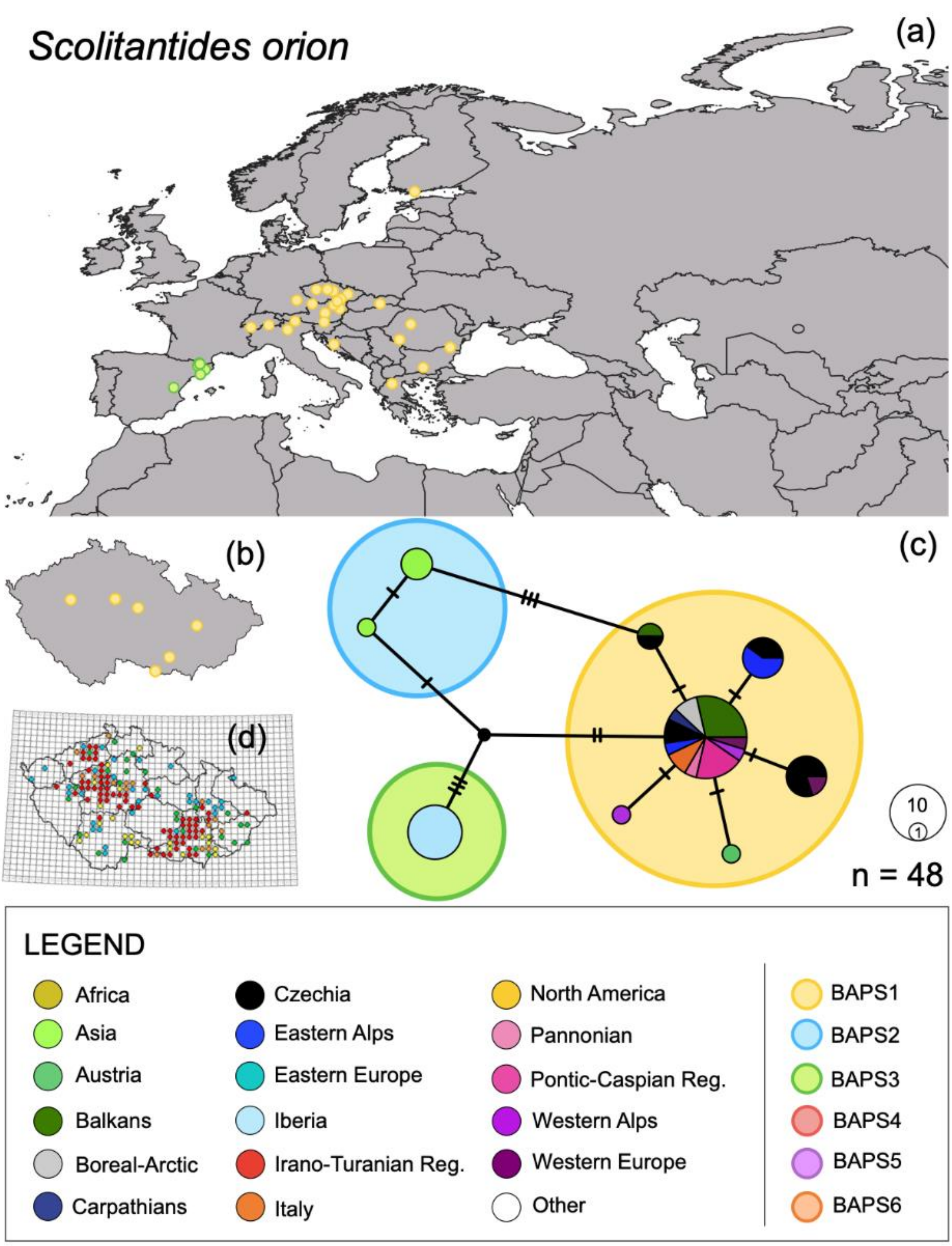


*Scolitantides orion* (Pallas, 1771): **Habitat:** xerothermophilic 1. **Red List:** VU

**BAPS:** BAPS-East-Balkans. **HAP:** HAP-East-Balkans

**Range:** Palearctic. **Distribution CZ:** dry rocky habitats in thermophilic regions of Bohemia and Moravia. Decreased with overgrown of rocky habitats.

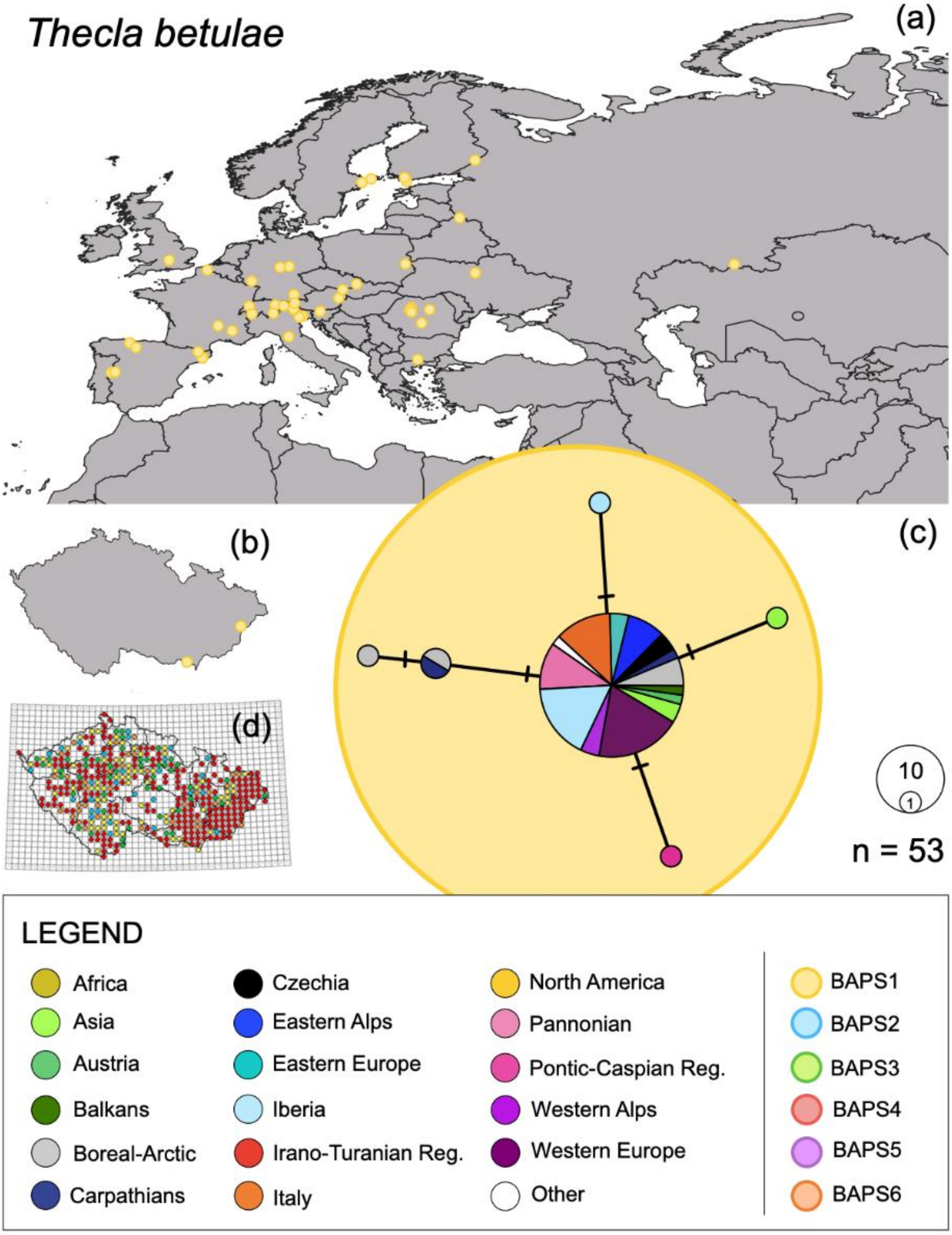


*Thecla betulae* (Linnaeus, 1758): **Habitat:** mesophilic 2 + xerothermophilic 2. **Red List:** LC
**BAPS:** BAPS-low. **HAP:** HAP-low

**Range:** Palearctic. **Distribution CZ:** dry warm shrublands and orchards across the entire country except for mountain regions.

**Nymphalidae**

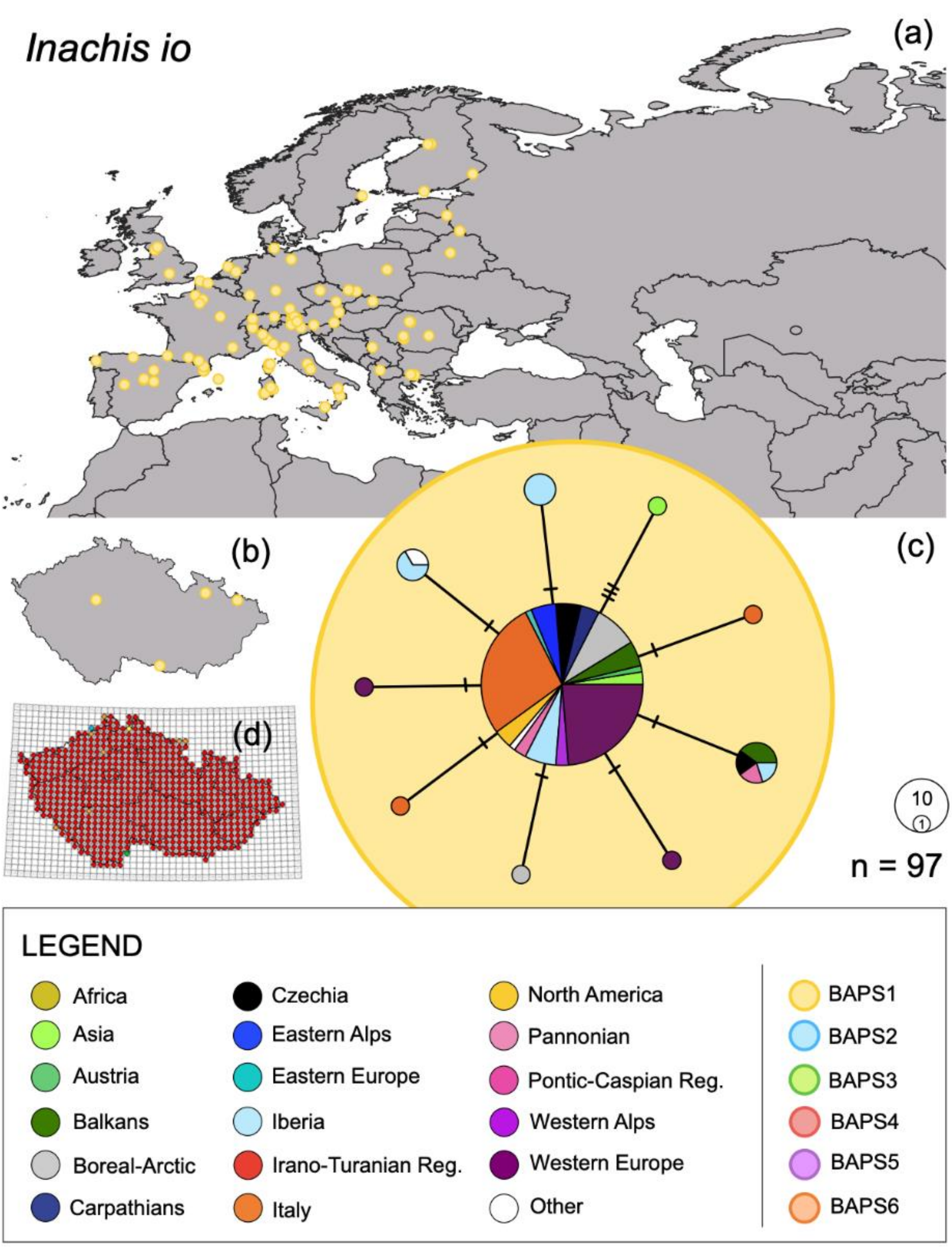


*Aglais io* (Linnaeus, 1758): **Habitat:** ubiquitous. **Red List:** LC

**BAPS:** BAPS-low. **HAP:** HAP-wide-expand

**Range:** Palearctic. **Distribution CZ:** various habitats, widespread and common.

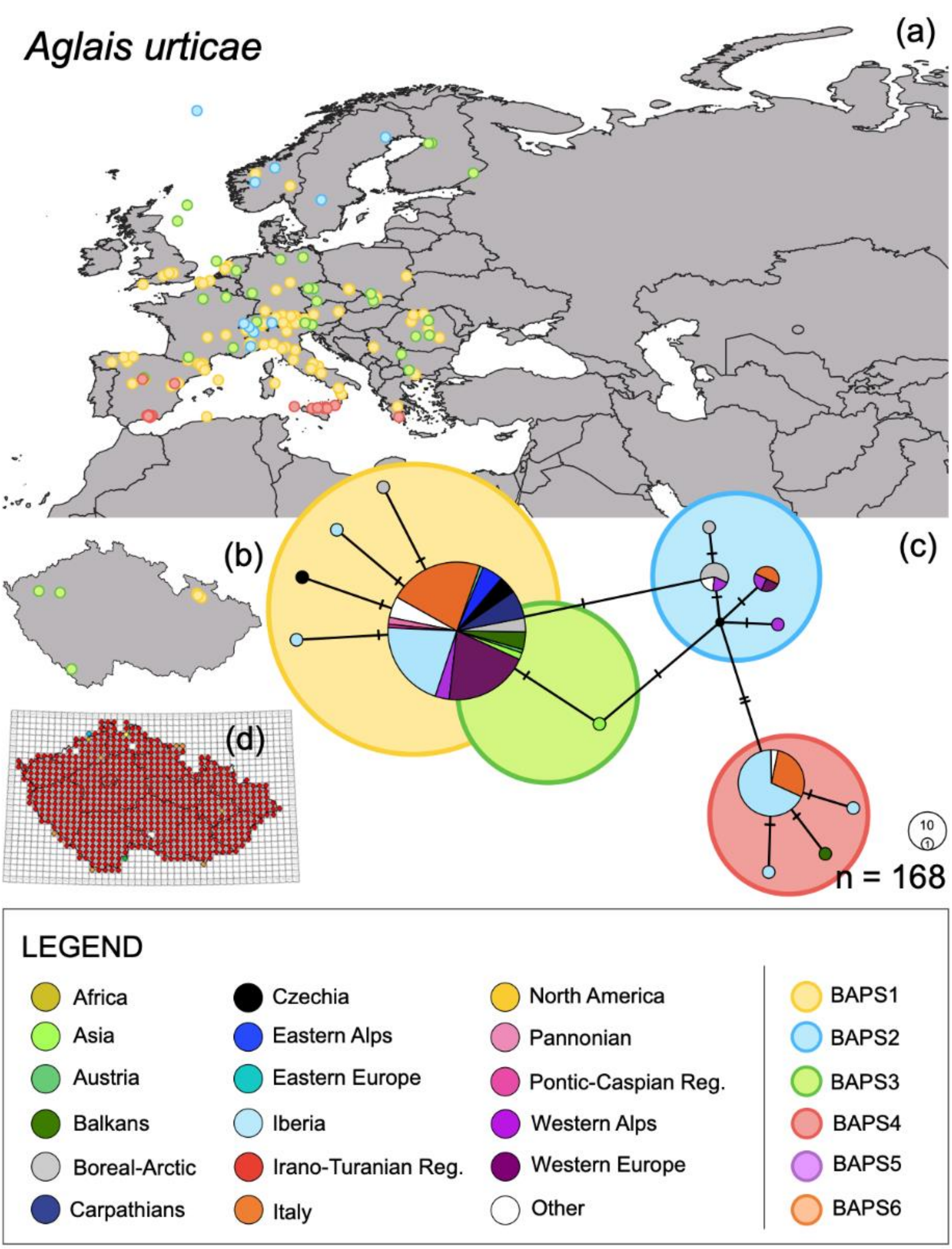


*Aglais urticae* (Linnaeus, 1758): **Habitat:** ubiquitous. **Red List:** LC

**BAPS:** BAPS-mix. **HAP:** HAP-wide-low

**Range:** Palearctic. **Distribution CZ:** various habitats, widespread and common.

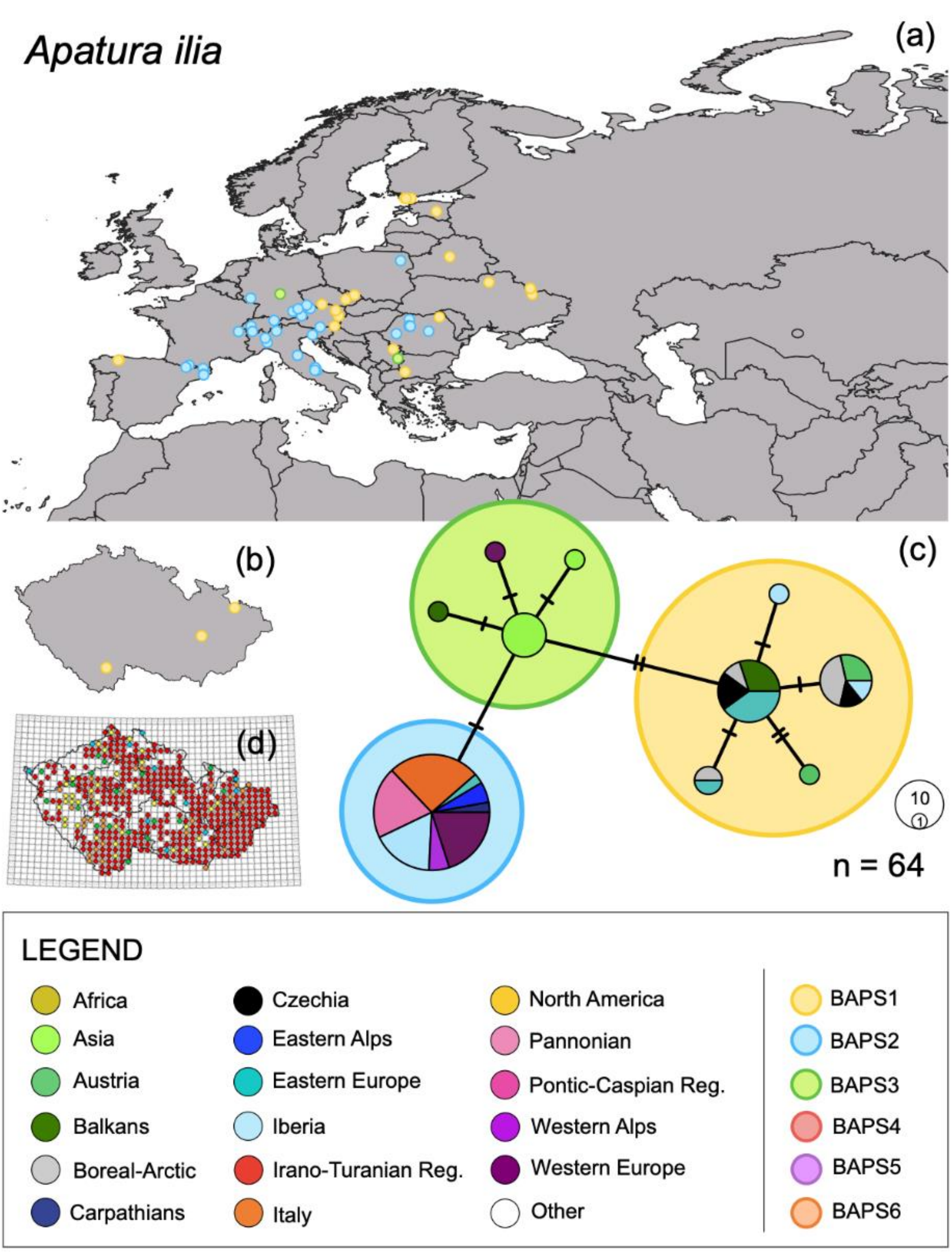


*Apatura ilia* ([Denis & Schiffermüller], 1775): **Habitat:** mesophilic 3. **Red List:** LC

**BAPS:** BAPS-wide. **HAP:** HAP-wide-low

**Range:** Euro-Siberian, disjunctive. **Distribution CZ:** woodland openings in alluvia. Widespread across the country.

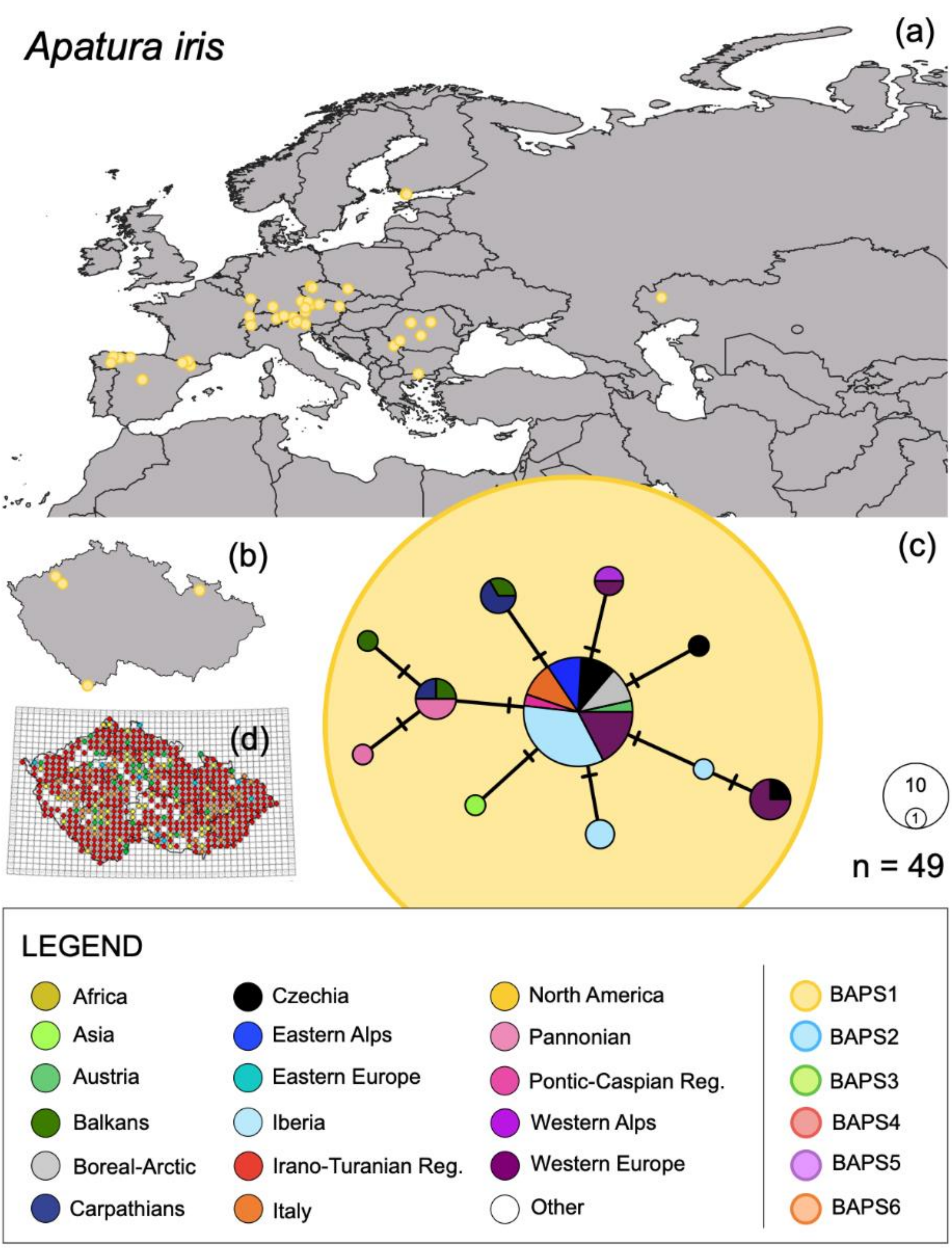


*Apatura iris* (Linnaeus, 1758): **Habitat:** mesophilic 3. **Red List:** LC

**BAPS:** BAPS-low. **HAP:** HAP-wide-low

**Range:** Euro-Siberian. **Distribution CZ:** woodland openings in alluvia. Widespread across the country, but more common in mountains.

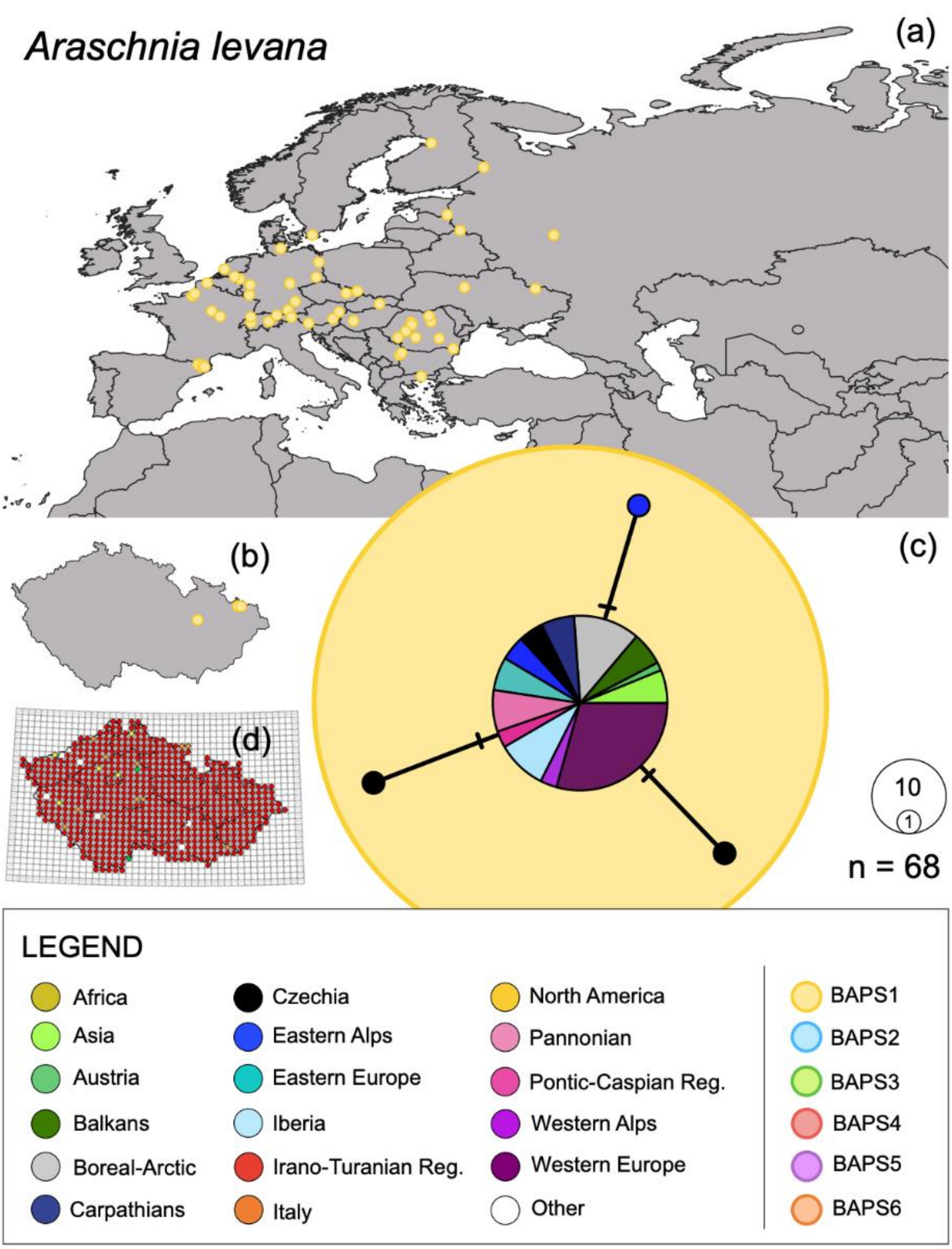


*Araschnia levana* (Linnaeus, 1758): **Habitat:** mesophilic 2. **Red List:** LC

**BAPS:** BAPS-low. **HAP:** HAP-low

**Range:** Palearctic. **Distribution CZ:** woodland openings and margins. Widespread and common.

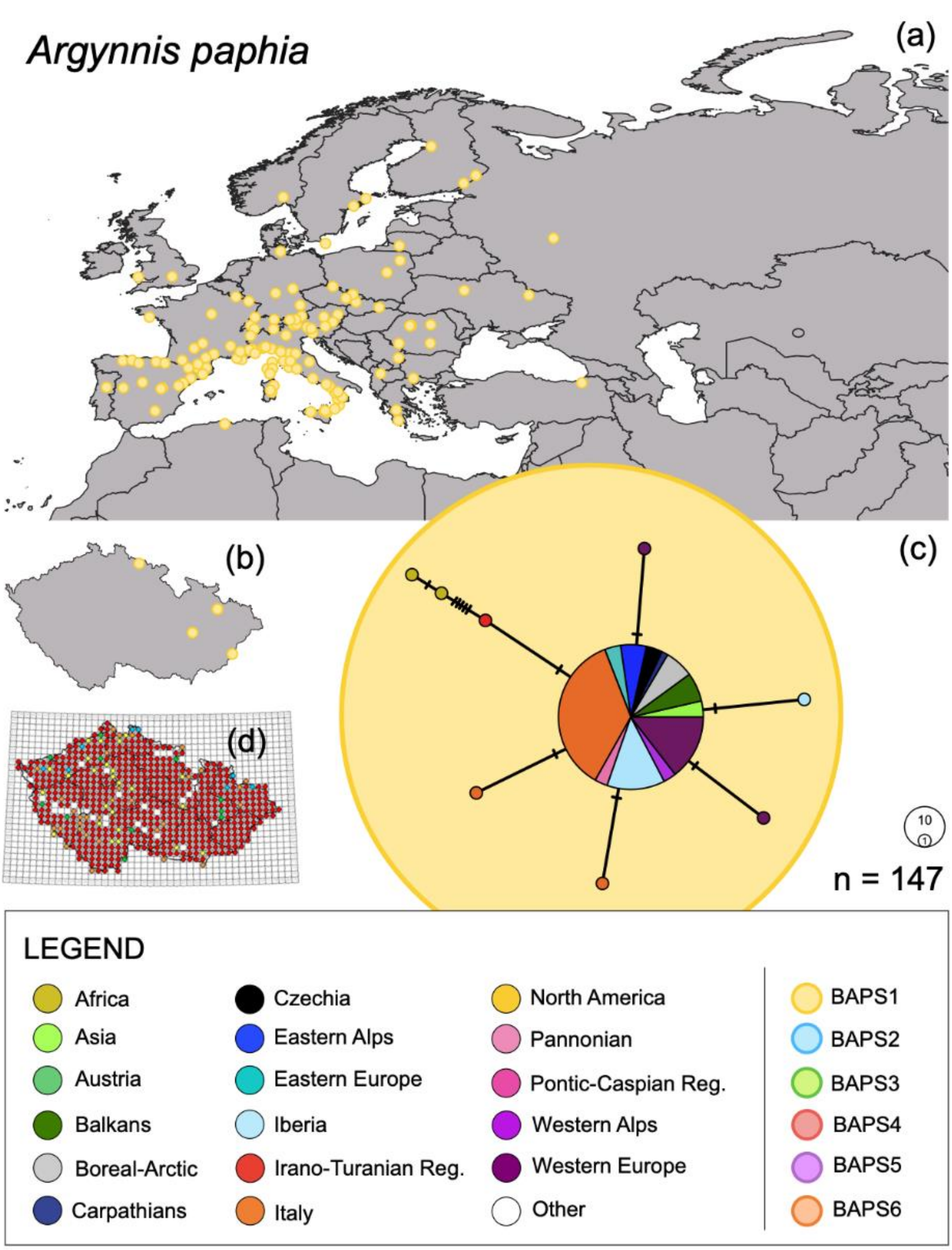


*Argynnis paphia* (Linnaeus, 1758): **Habitat:** mesophilic 3. **Red List:** LC

**BAPS:** BAPS-low. **HAP:** HAP-wide-low

**Range:** Euro-Siberian. **Distribution CZ:** woodland openings and margins. Widespread across the country.

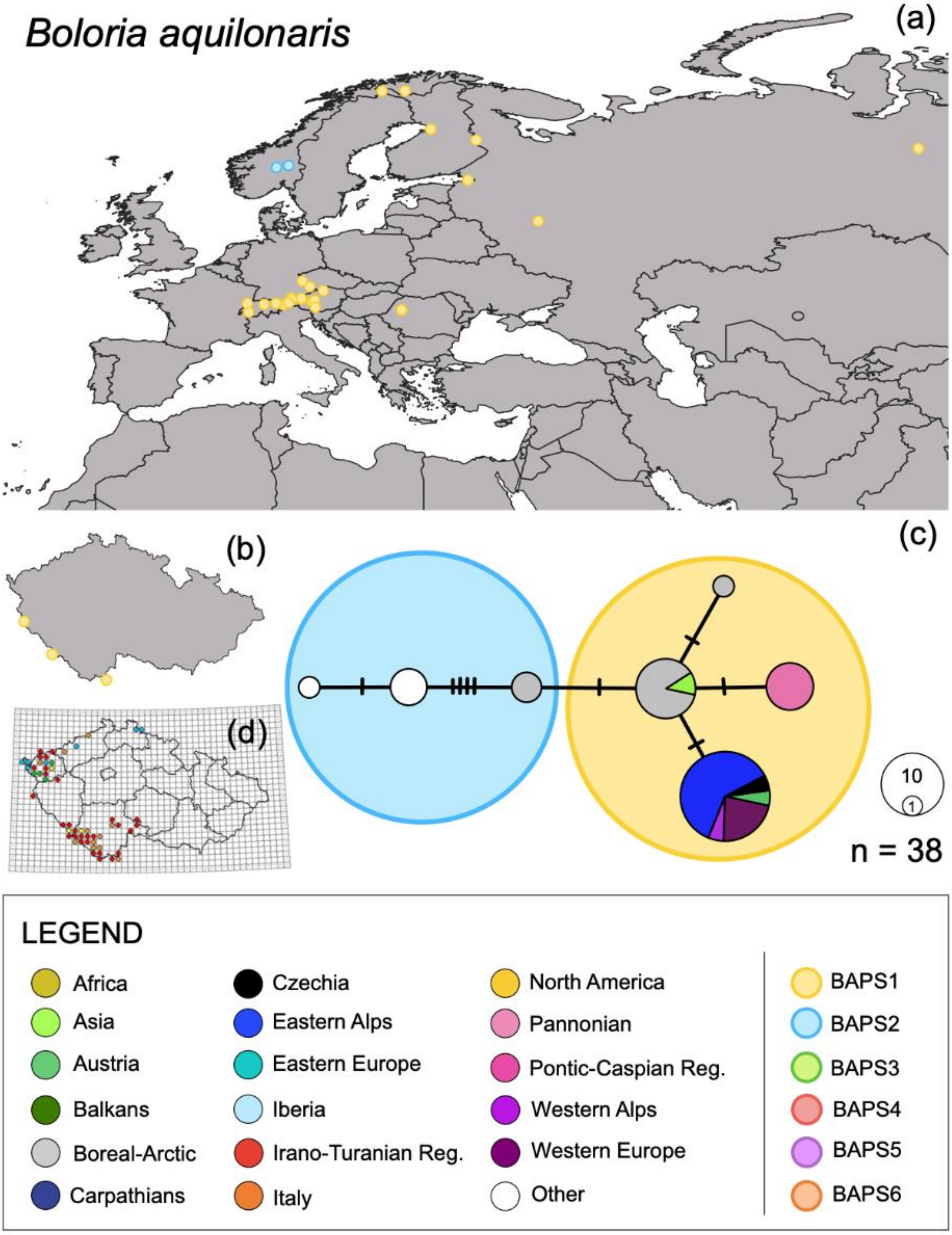


*Boloria aquilonaris* (Stichel, 1908): **Habitat:** tyrphophilic. **Red List:** VU

**BAPS:** BAPS-temperate. **HAP:** HAP-Central

**Range:** Euro-Siberian. **Distribution CZ:** bogs with the host plant *Oxycoccus palustris*. Only mountains in West and South Bohemia, isolated populations in Bohemian-Moravian Highlands. Extinct from other borderland mountains, never recorded from Moravia.

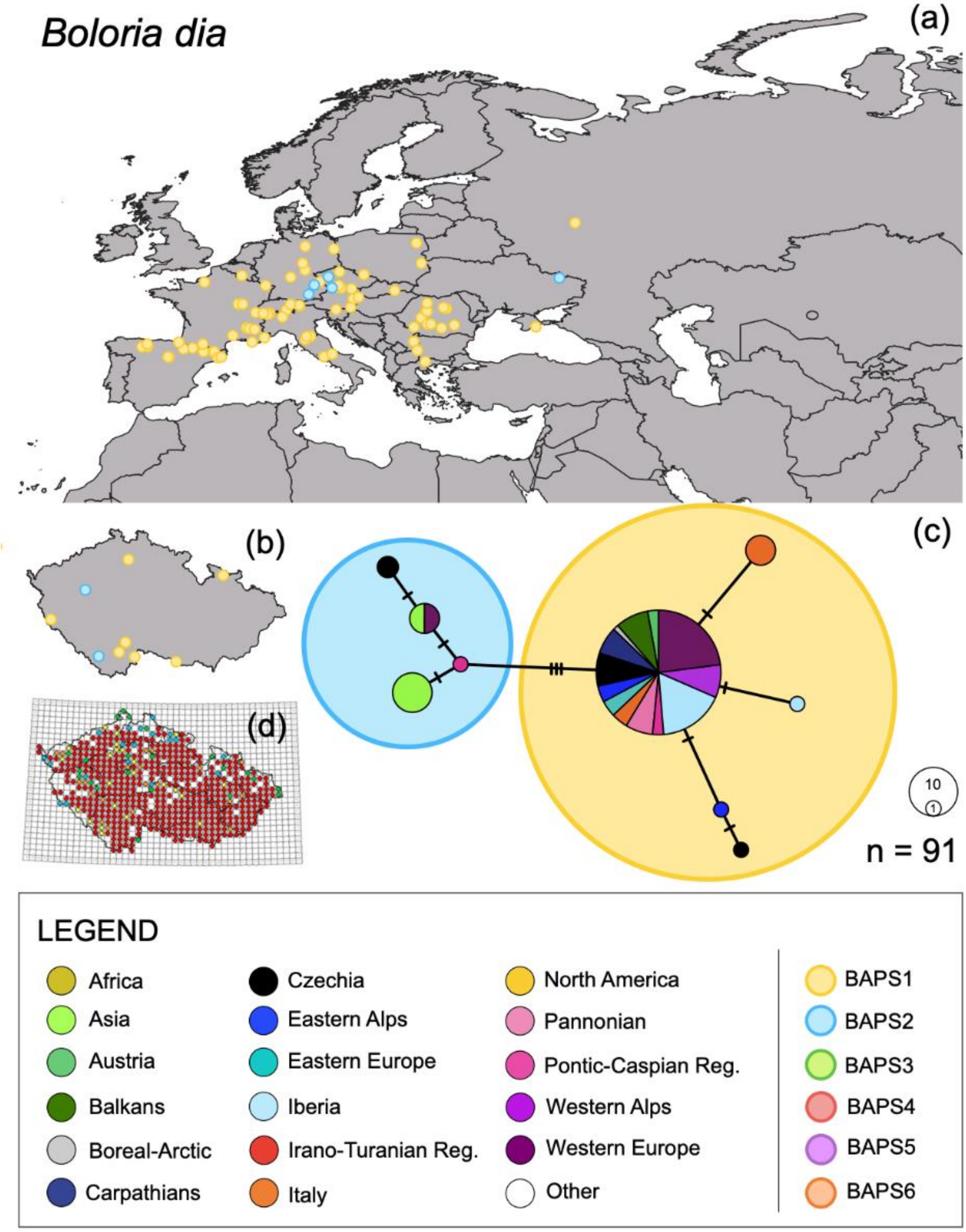


*Boloria dia* (Linnaeus, 1767): **Habitat:** mesophilic 2 + xerothermophilic 2. **Red List:** LC
**BAPS:** BAPS-wide + BAPS-temperate. **HAP:** HAP-wide-low + HAP-temperate
**Range:** Euro-Siberian. **Distribution CZ:** dry scrublands, warm overgrown sites. Widespread across CZ, missing from cold mountains.

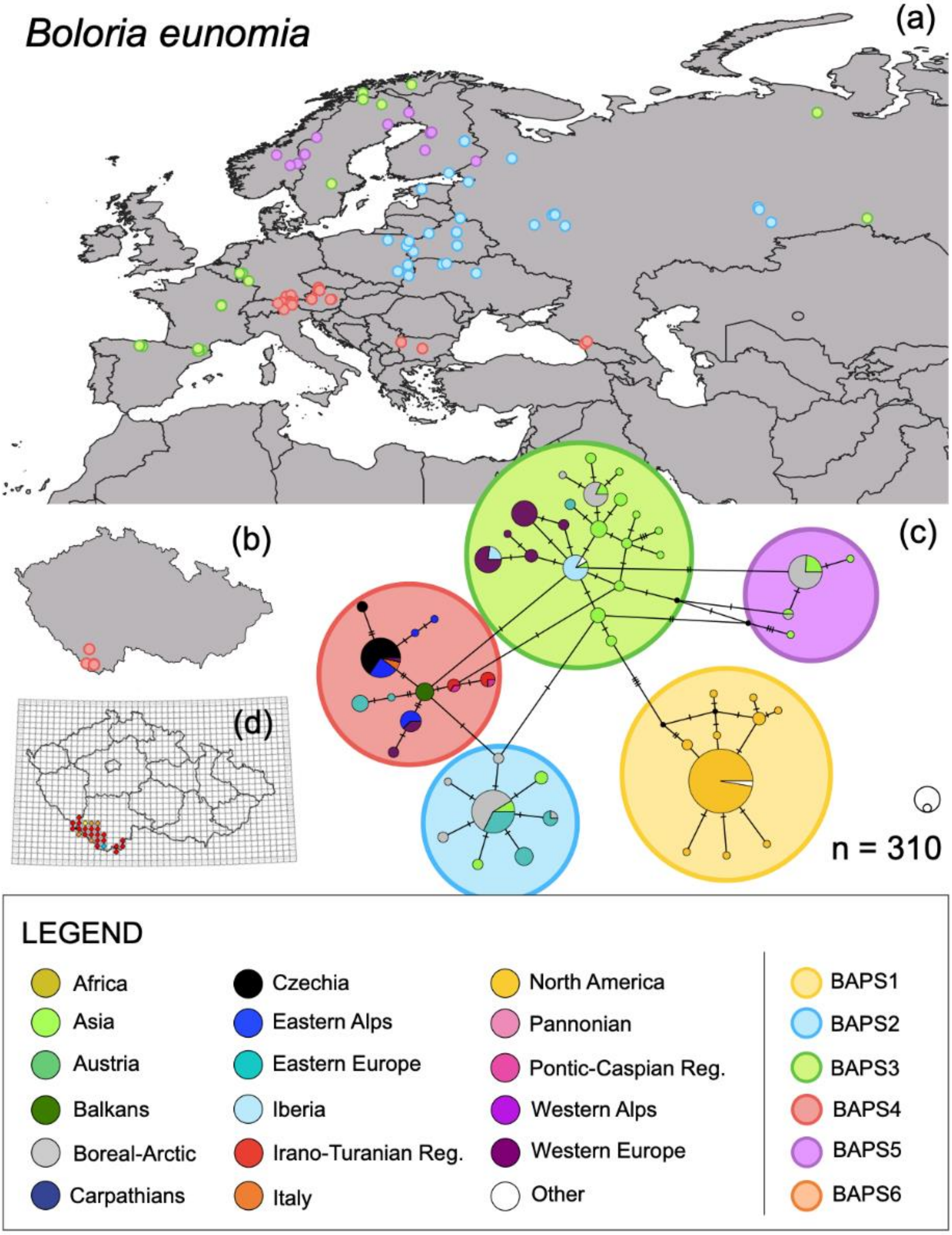


*Boloria eunomia* (Esper, 1800): **Habitat:** tyrphophilic. **Red List:** LC

**BAPS:** BAPS-temperate. **HAP:** HAP-Central

**Range:** Holarctic. **Distribution CZ:** mountain bogs and wet meadows. Only in Bohemian Forest and its foothills, currently expanding.

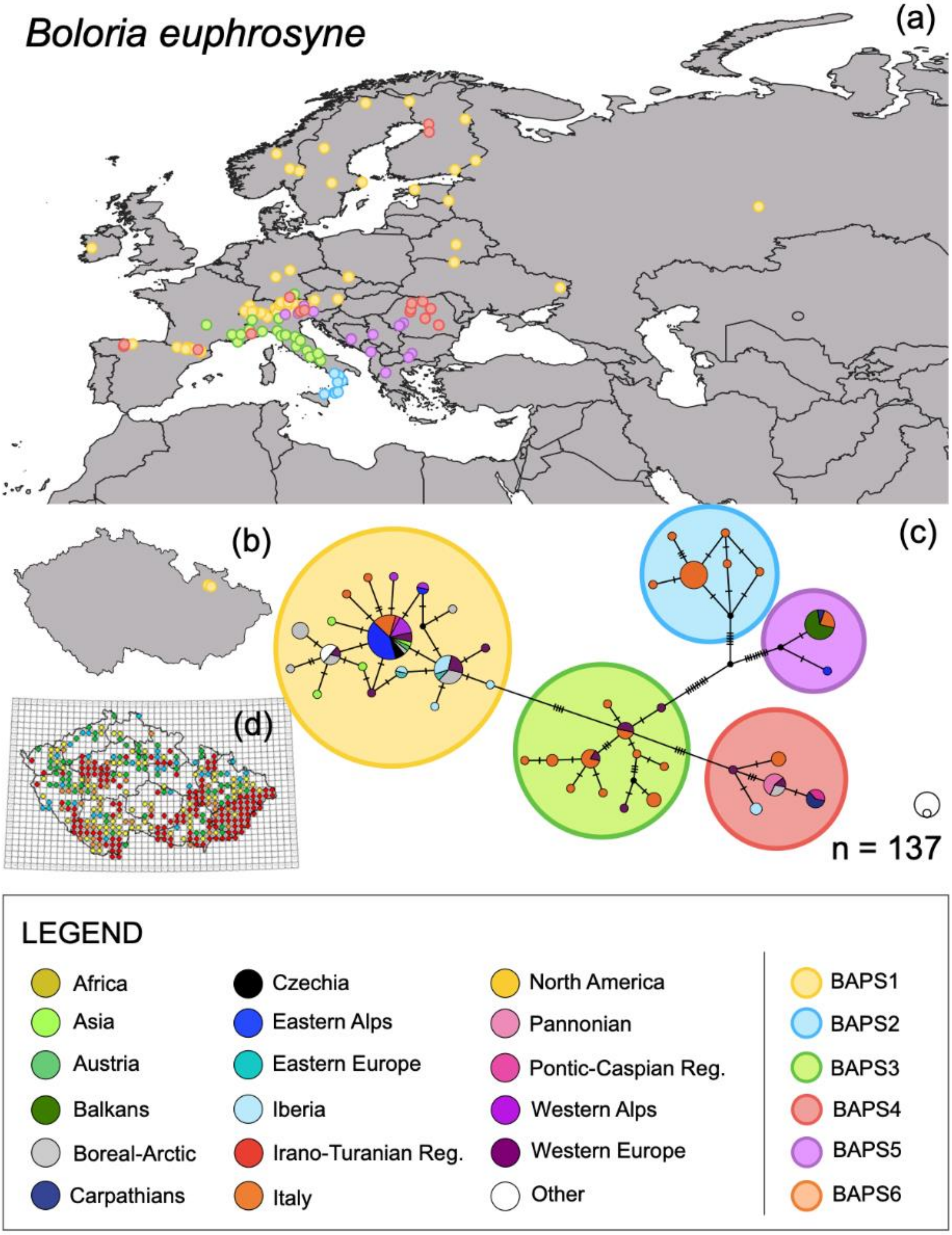


*Boloria euphrosyne* (Linnaeus, 1758): **Habitat:** mesophilic 2. **Red List:** VU

**BAPS:** BAPS-temperate. **HAP:** HAP-temperate

**Range:** Euro-Siberian. **Distribution CZ:** clearings and woodland margins, scrublands. Widespread across the country, expanding after diebacks of spruce plantations.

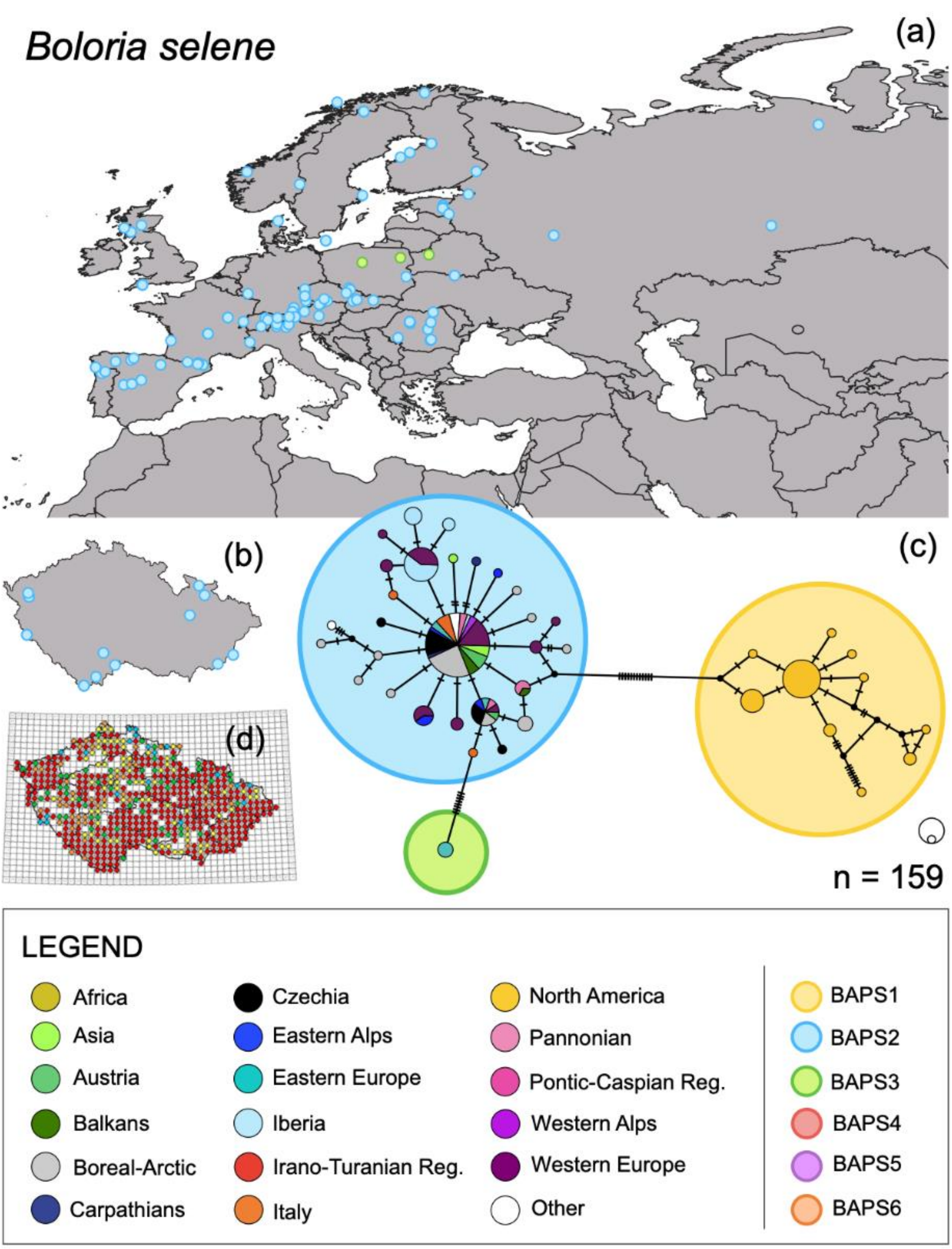


*Boloria selene* ([Denis & Schiffermüller], 1775): **Habitat:** mesophilic 2 + tyrphophilic. **Red List:** NT

**BAPS:** BAPS-wide. **HAP:** HAP-wide-complex

**Range:** Holarctic. **Distribution CZ:** meadows and open woodlands, bogs and wetlands. Widespread across the country, decreased in areas of intensive agriculture.

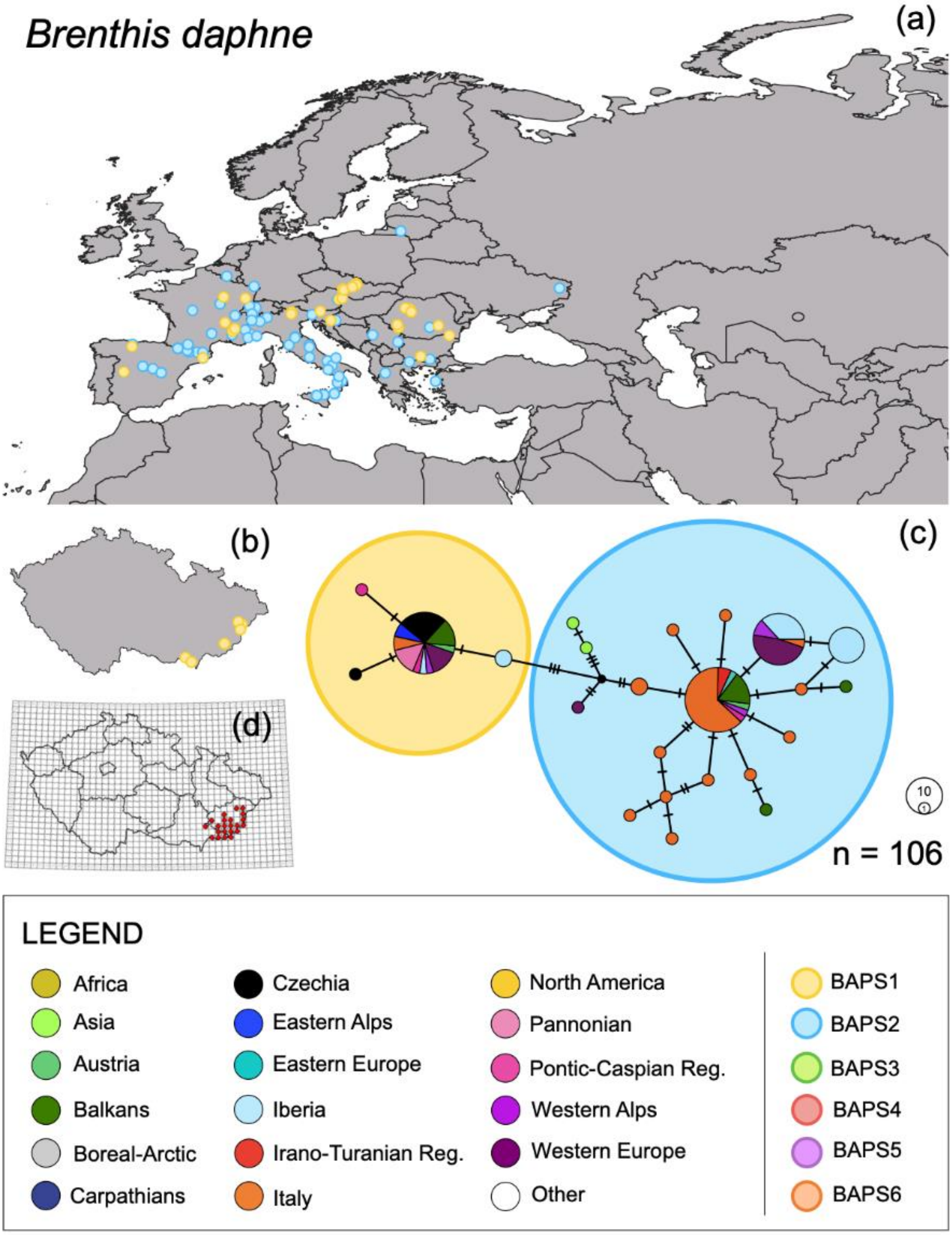


*Brenthis daphne* (Bergsträsser, 1780): **Habitat:** mesophilic 2 + xerothermophilic 2. **Red List:** LC

**BAPS:** BAPS-wide. **HAP:** HAP-temperate

**Range:** Palearctic. **Distribution CZ:** scrublands, woodland clearings and margins. Started to expand from Slovakia in 1995, now inhabiting most of the Carpathian Mountains system in Moravia.

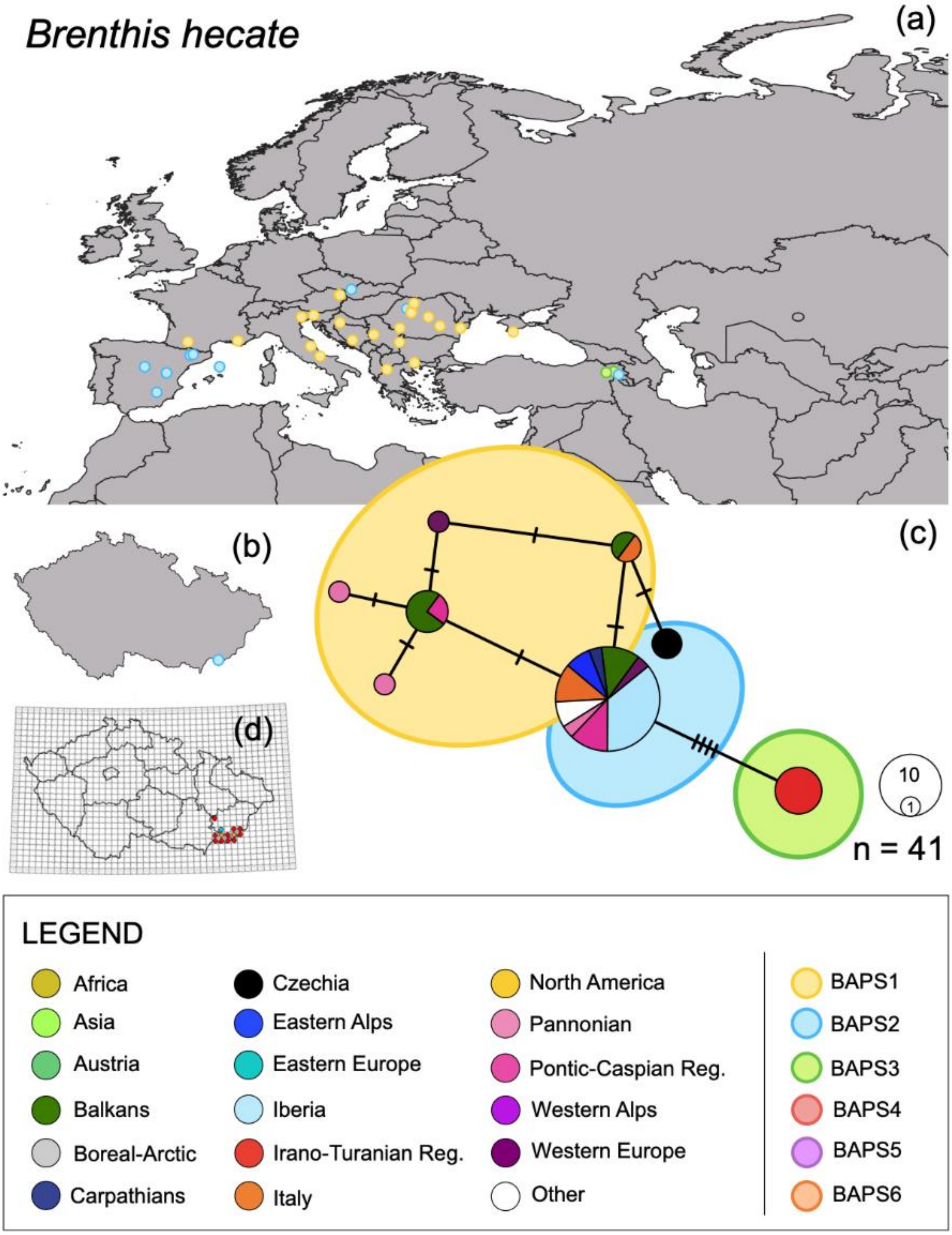


*Brenthis hecate* ([Denis & Schiffermüller], 1775): **Habitat:** xerothermophilic 2. **Red List:** NT

**BAPS:** BAPS-wide. **HAP:** HAP-Italy-Balkans

**Range:** Pontic-Mediterranean. **Distribution CZ:** dry scrublands and meadows with scattered trees. Both historically and recently present only in southeast Moravia (mostly White Carpathians).

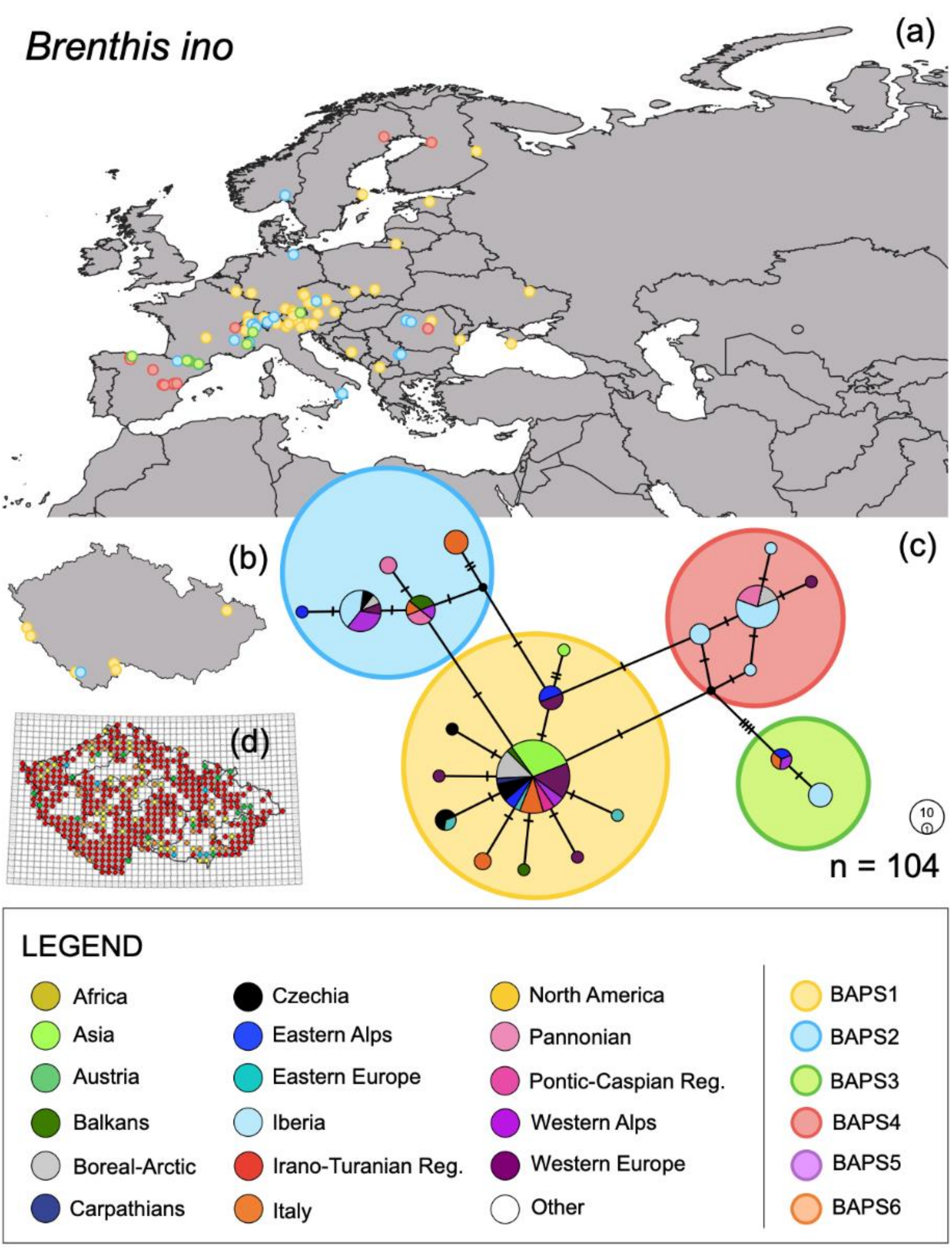


*Brenthis ino* (Rottemburg, 1775): **Habitat:** mesophilic 2 + hydrophilic. **Red List:** LC

**BAPS:** BAPS-wide + BAPS-temperate. **HAP:** HAP-wide-complex

**Range:** Euro-Siberian. **Distribution CZ:** wet, often abandoned meadows. Widespread across the country, expanding with general land abandonment.

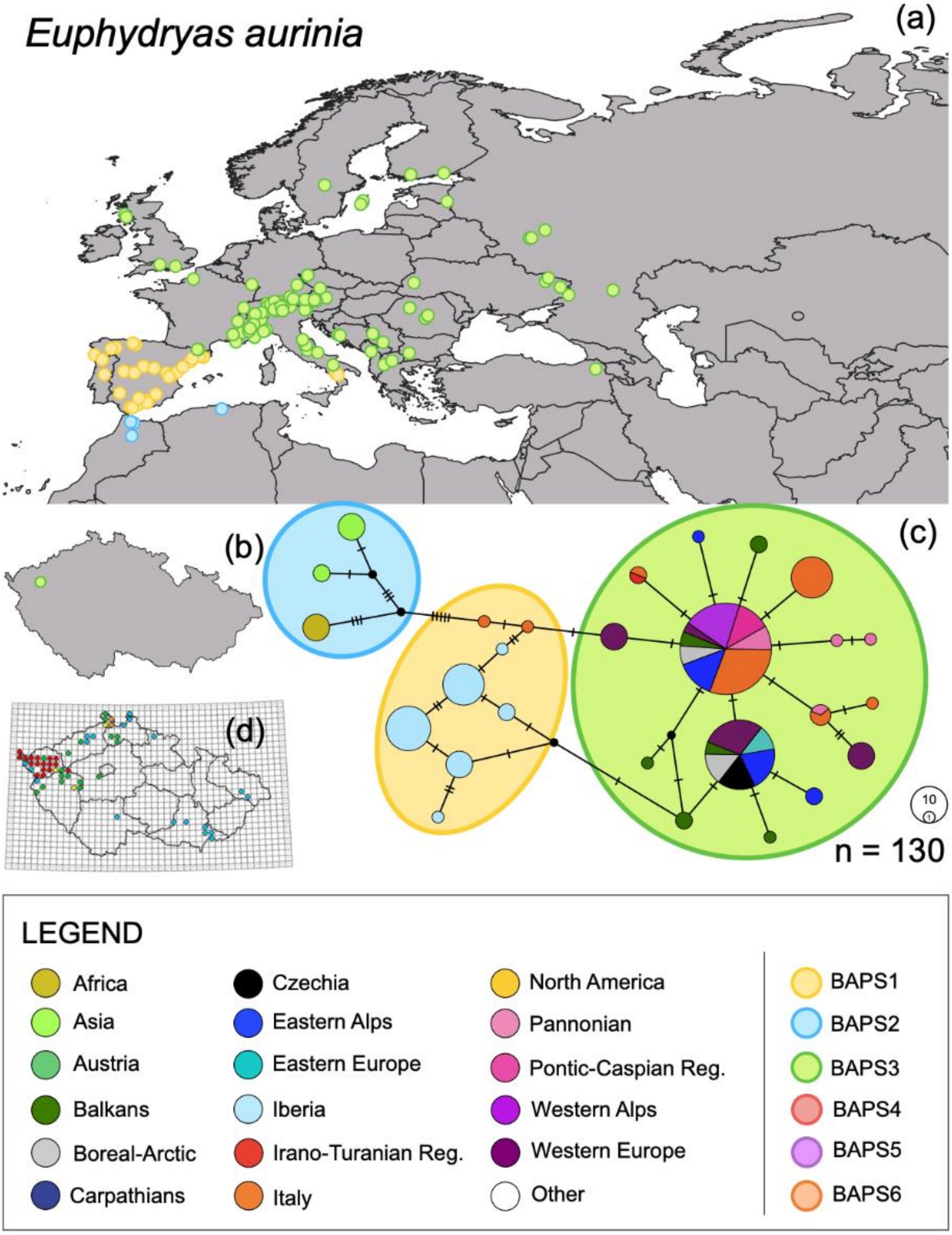


*Euphydryas aurinia* (Rottemburg, 1775): **Habitat:** hydrophilic. **Red List:** EN

**BAPS:** BAPS-wide. **HAP:** HAP-temperate

**Range:** Palearctic. **Distribution CZ:** wet and peat meadows in piedmont areas. It is currently present only in West Bohemia, extinct from several other parts of the country, decreasing and endangered.

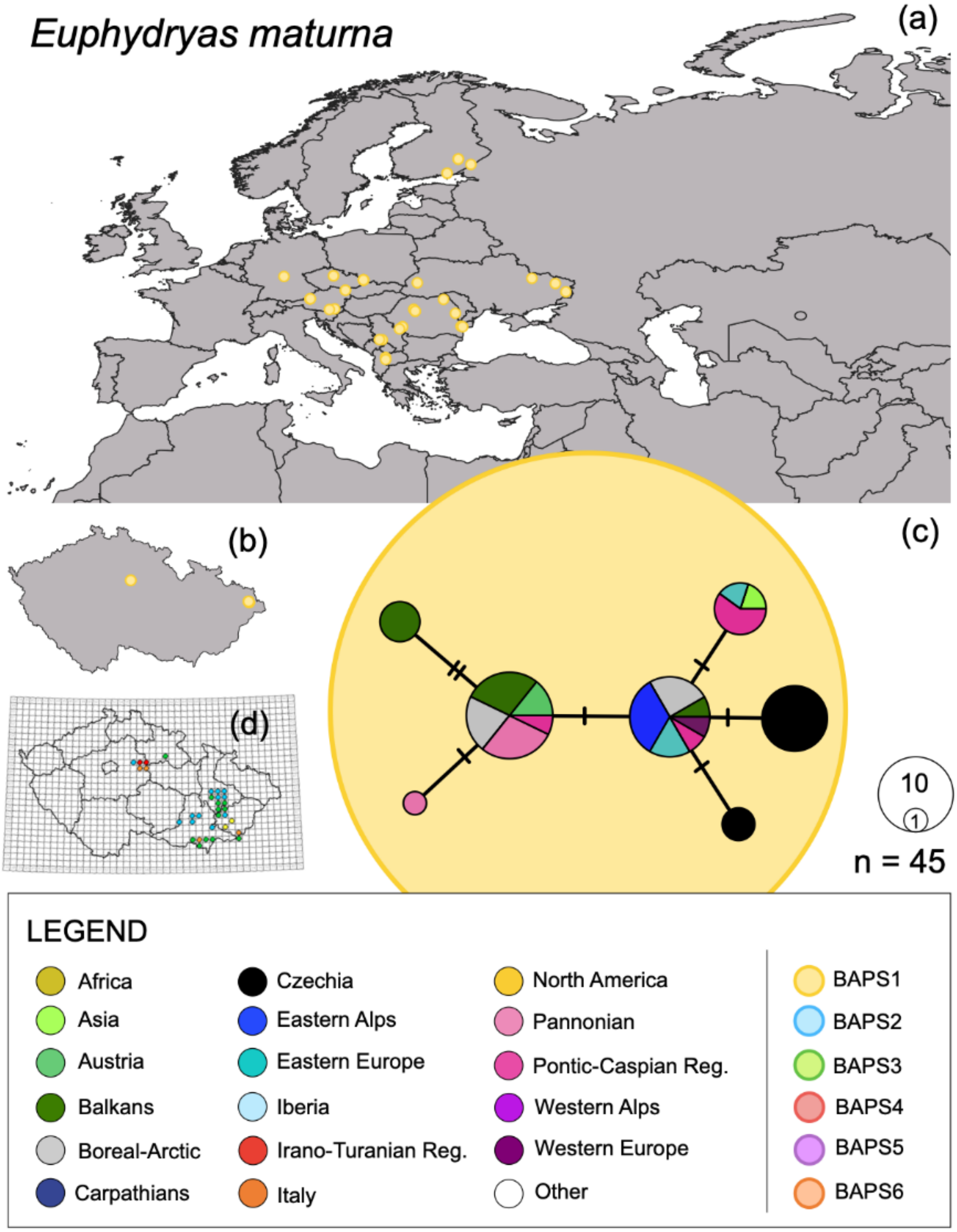


*Euphydryas maturna* (Linnaeus, 1758): **Habitat:** mesophilic 3. **Red List:** CR

**BAPS:** BAPS-temperate. **HAP:** HAP-temperate (such distribution)

**Range:** Euro-Siberian. **Distribution CZ:** open deciduous woodlands. Extinct from Moravia where it used to occur in south and central parts, retreated after the 1950s. Currently present on a single locality in Elbe alluvium, the population is conservation-dependent and supported by a conservation breeding program (John et al., 2021). Unauthorized introduction to northeastern Moravia (noticed in 2018, surviving there since then).

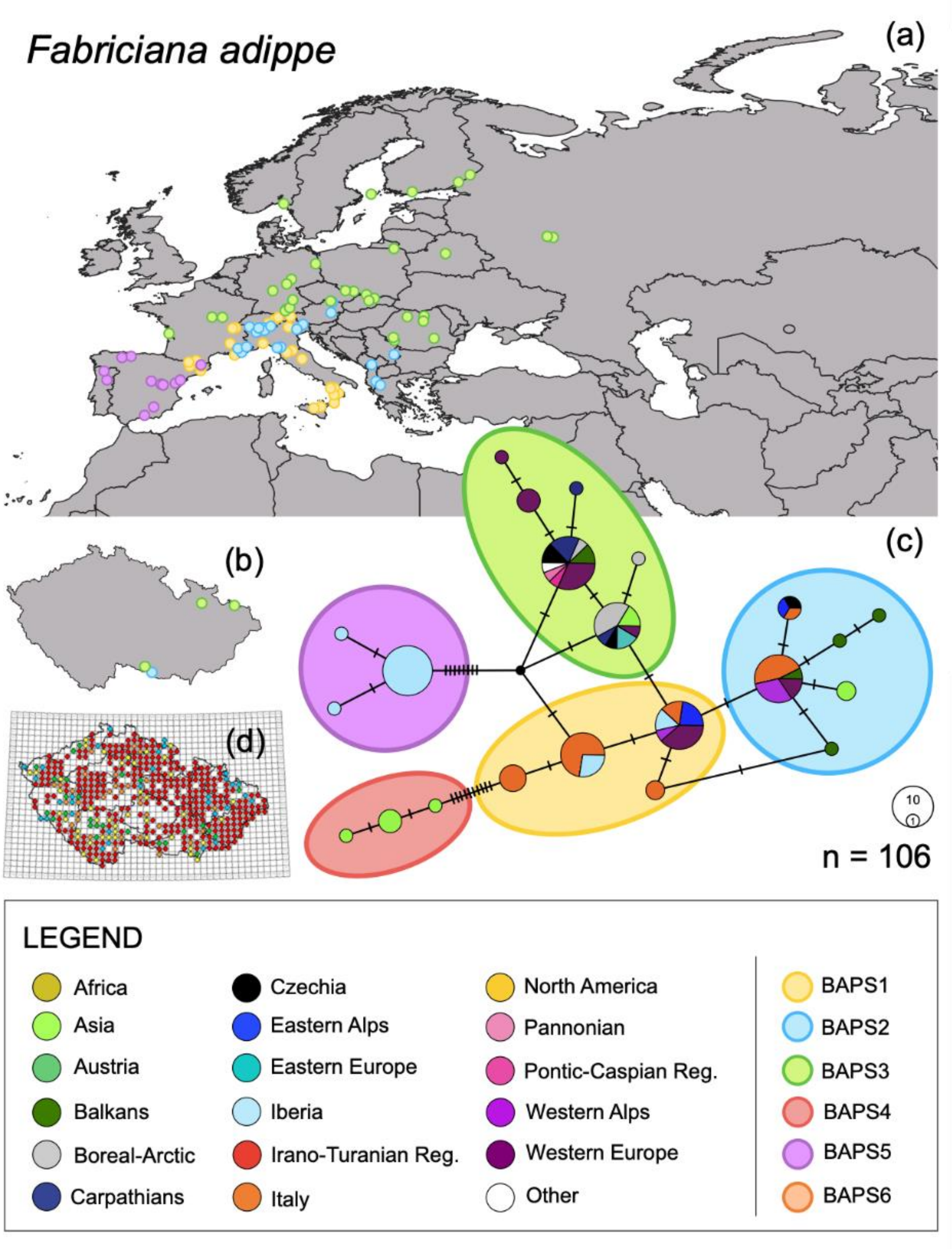


*Fabriciana adippe* ([Denis & Schiffermüller], 1775): **Habitat:** mesophilic 2. **Red List:** LC **BAPS:** BAPS-Italy-Balkans + BAPS-temperate. **HAP:** HAP-Italy-Balkans + HAP-temperate **Range:** Palearctic. **Distribution CZ:** scrubland and woodland clearings. Widespread across the country, decreased after the 1980s and currently re-expanding (Konvickova et al., 2023).

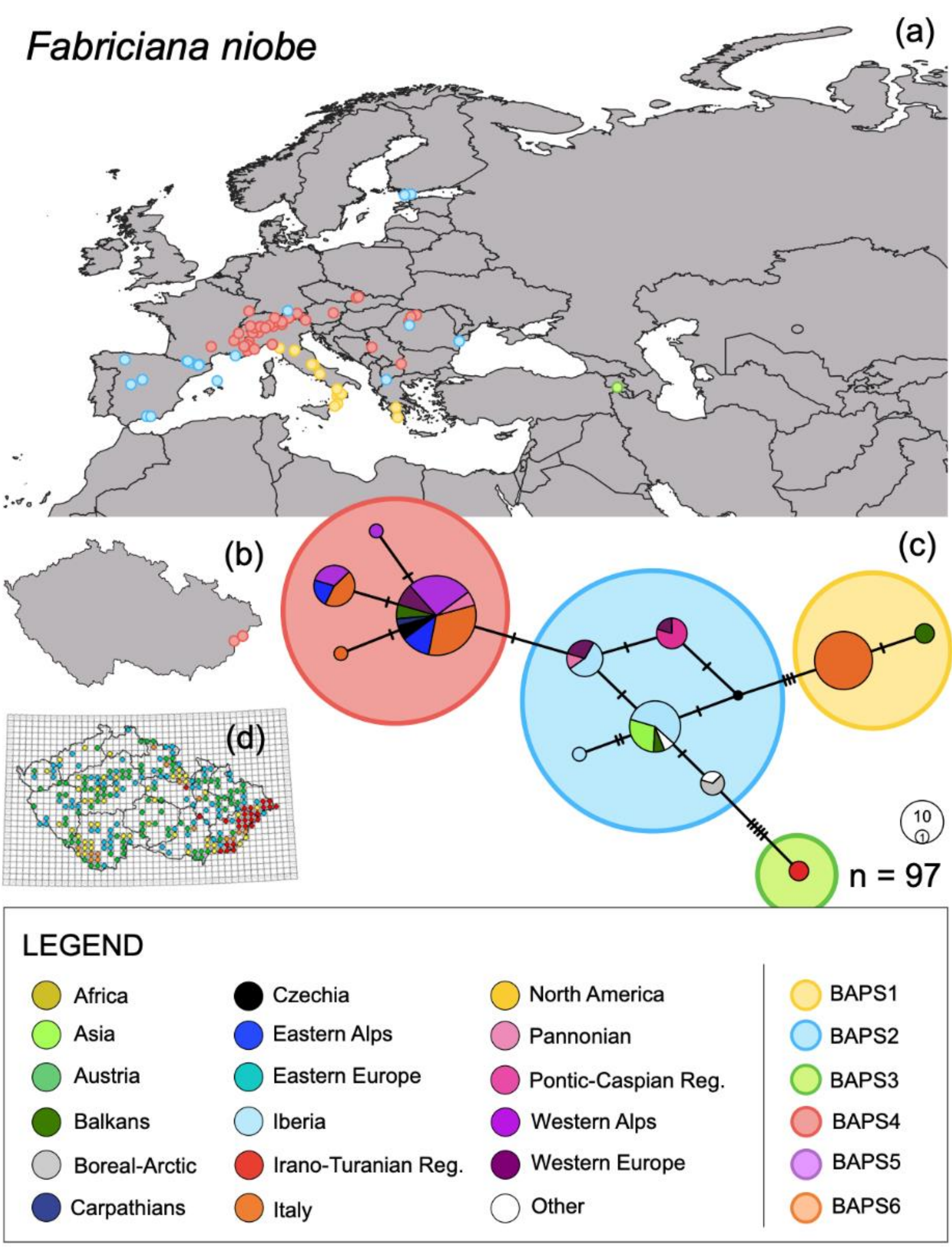


*Fabriciana niobe* (Linnaeus, 1758): **Habitat:** mesophilic 2. **Red List:** CR

**BAPS:** BAPS-Central. **HAP:** HAP-wide-low

**Range:** Palearctic. **Distribution CZ:** pastures and meadows in piedmont areas. In the past, the species was widespread across the country, but severely decreased. Currently persisting only in east Moravia.

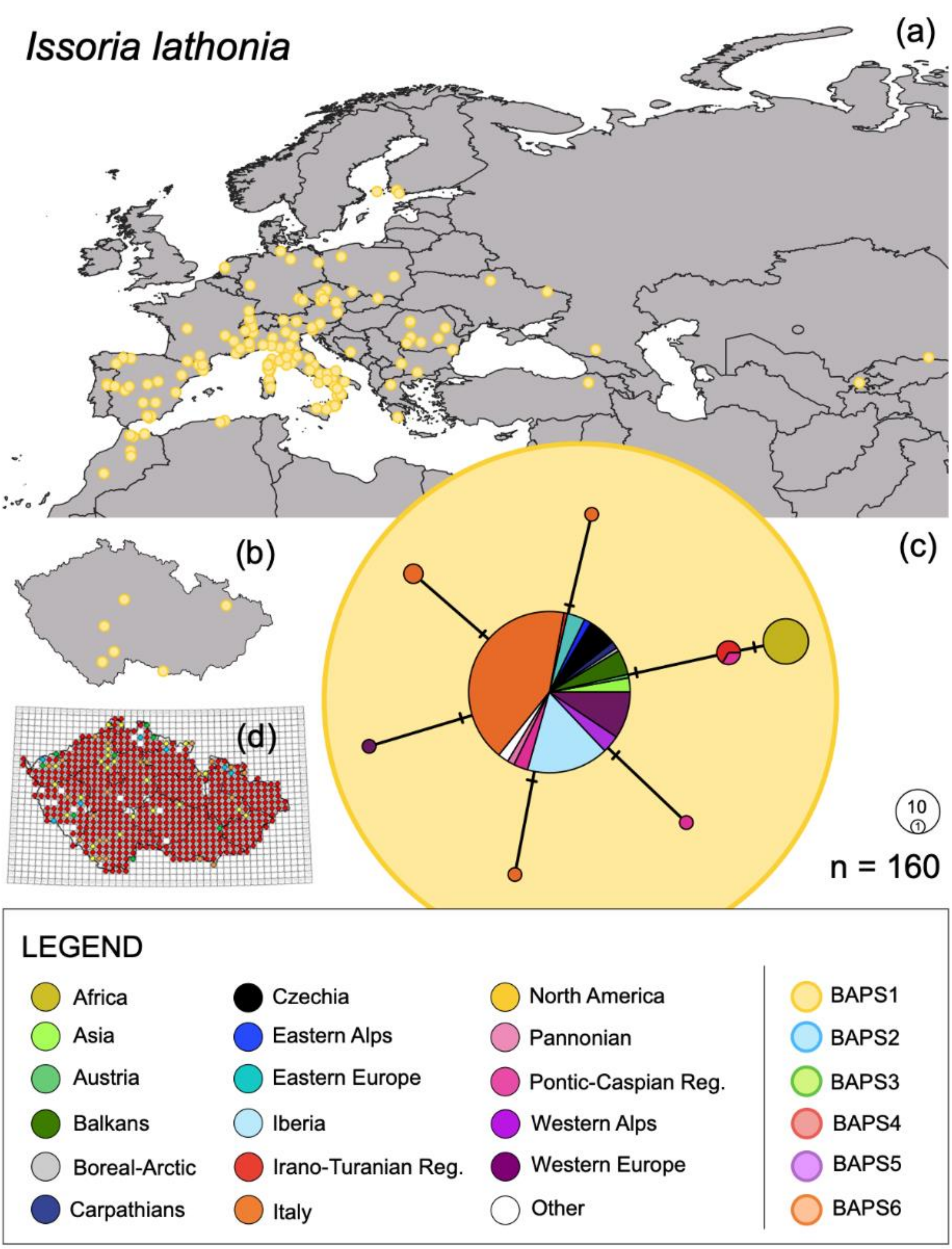


*Issoria lathonia* (Linnaeus, 1758): **Habitat:** ubiquitous. **Red List:** LC

**BAPS:** BAPS-low. **HAP:** HAP-low

**Range:** Western Palearctic. **Distribution CZ:** widespread and common migrant.

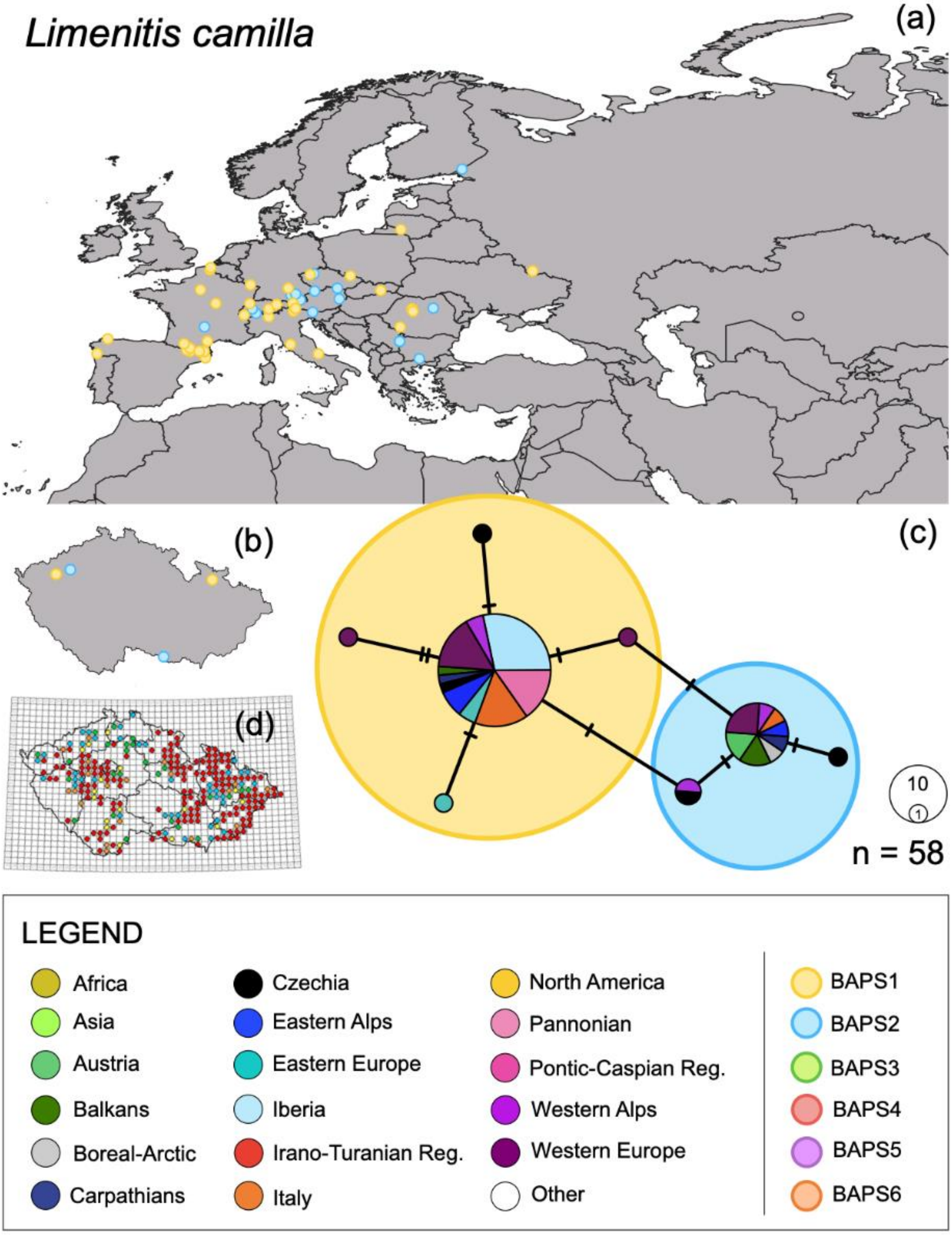


*Limenitis camilla* (Linnaeus, 1764): **Habitat:** mesophilic 3. **Red List:** NT

**BAPS:** BAPS-wide + BAPS-temperate. **HAP:** HAP-wide-low + HAP-temperate

**Range:** Palearctic. **Distribution CZ:** woodlands, margins and clearings. Two altitudinal optima reflecting the distribution of its host-plants, one in thermophilic woodlands with *Lonicera xylosteum*, the other, newly colonised, in mountain forests with *L. nigra*.

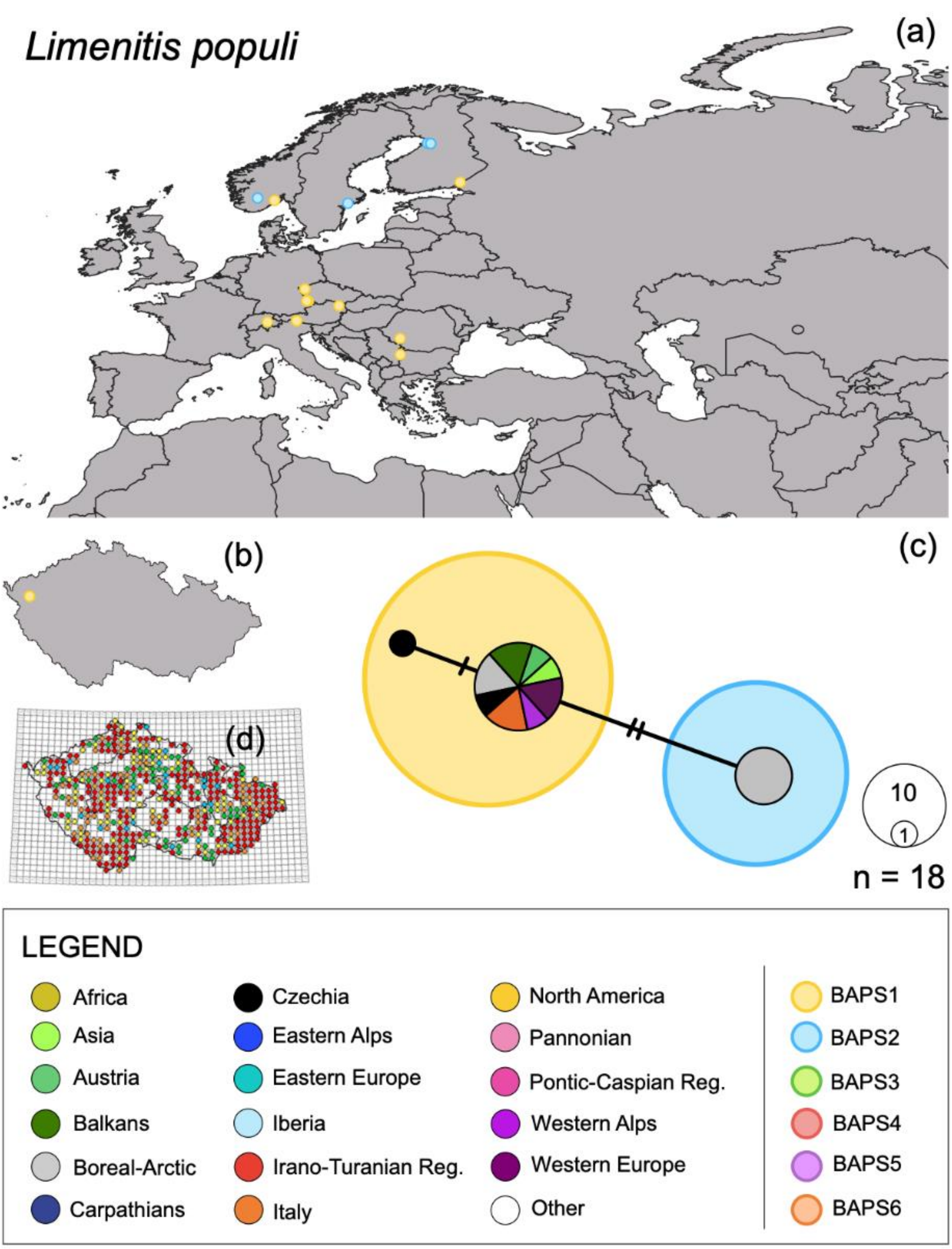


*Limenitis populi* (Linnaeus, 1758): **Habitat:** mesophilic 2. **Red List:** VU

**BAPS:** BAPS-temperate. **HAP:** HAP-temperate

**Range:** Euro-Siberian. **Distribution CZ:** woodland margins and clearings. In occurs across entire area, rare in agricultural landscapes.

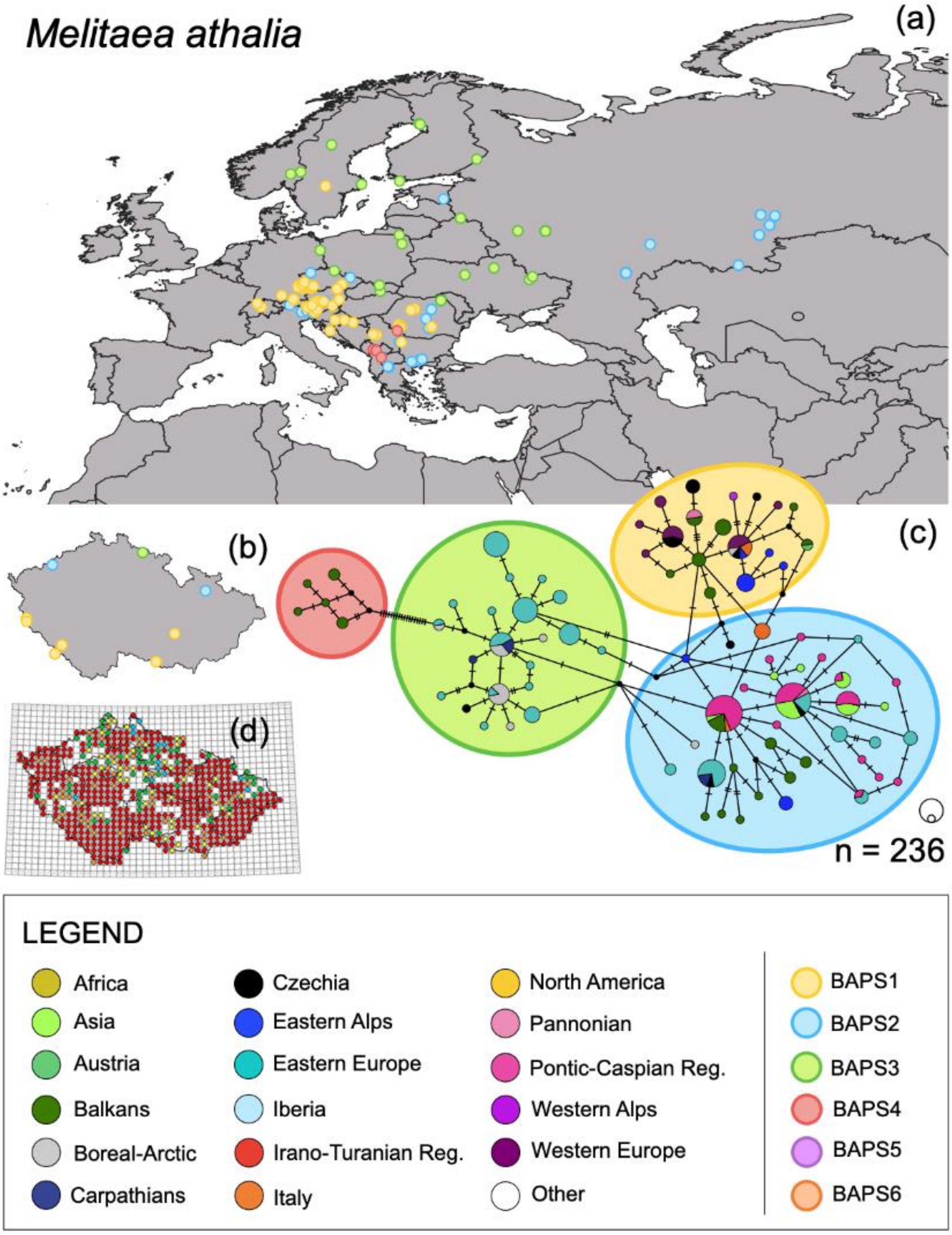


*Melitaea athalia* (Rottemburg, 1775): **Habitat:** mesophilic 2. **Red List:** NT

**BAPS:** BAPS-mix-East. **HAP:** HAP-mix

**Range:** Euro-Siberian. **Distribution CZ:** various meadow-woodland habitats, widespread across entire country, although decreasing. In CZ, three genetic lineages are present: BAPS1, occurring in Central/Eastern Europe and reaching Scandinavia, BAPS2 reaching further to the east, and BAPS3, occurring more northerly than BAPS1 and 2. The species is vicariant with *M. celadussa* inhabiting Iberia and Italy, and a highly divergent Balkan clade (both in mtDNA and genome-wide markers) is present (Tahami et al., 2021).

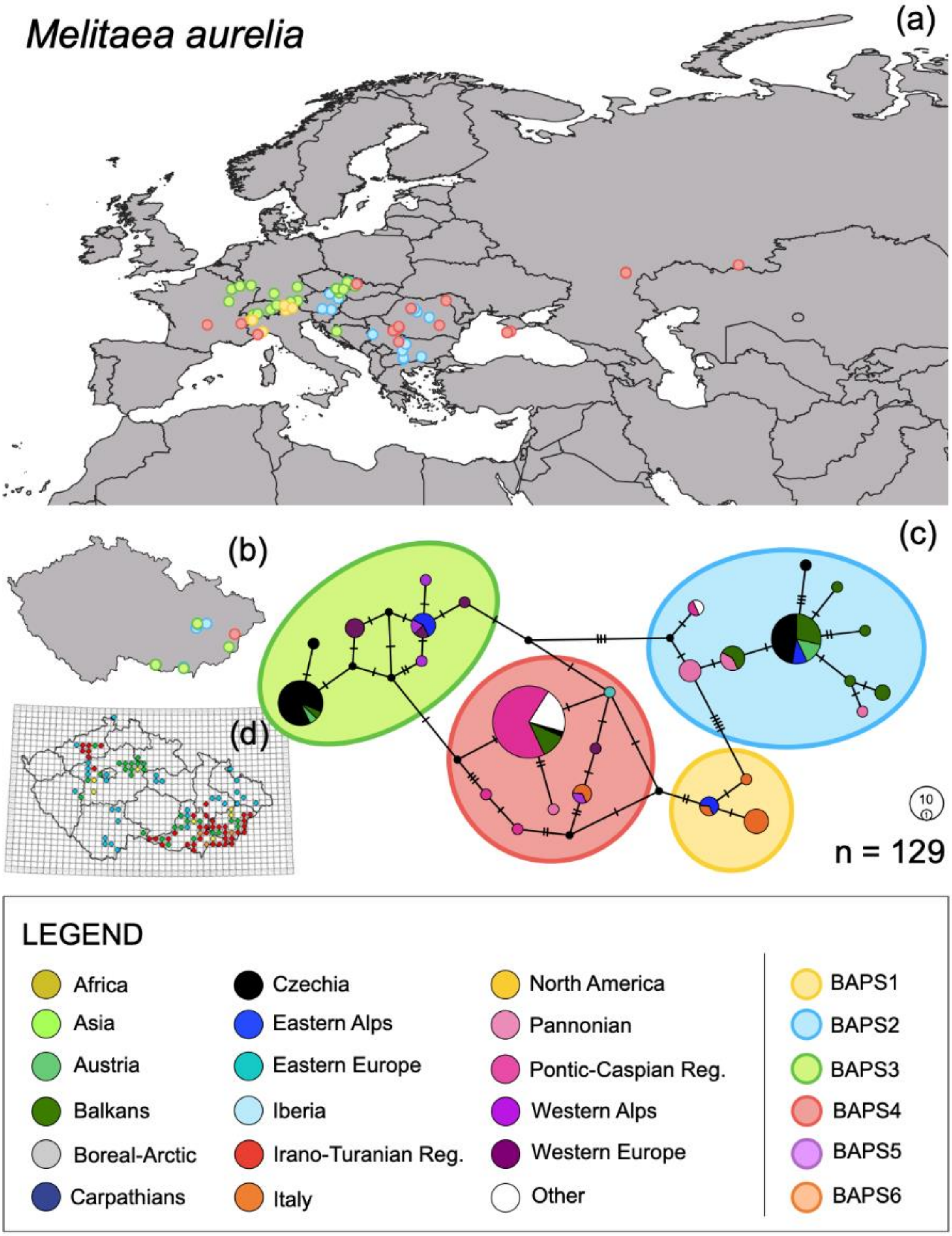


*Melitaea aurelia* Nickerl, 1850: **Habitat:** xerothermophilic 1. **Red List:** EN

**BAPS:** BAPS-mix. **HAP:** HAP-mix

**Range:** Western Palearctic. **Distribution CZ:** dry short grasslands. It was distributed in all warm regions in the past, but decreased severely. It currently occurs only in Central Bohemian Highlands and in several regions in southern and central Moravia. Several different mitochondrial lineages co-occur in CZ.

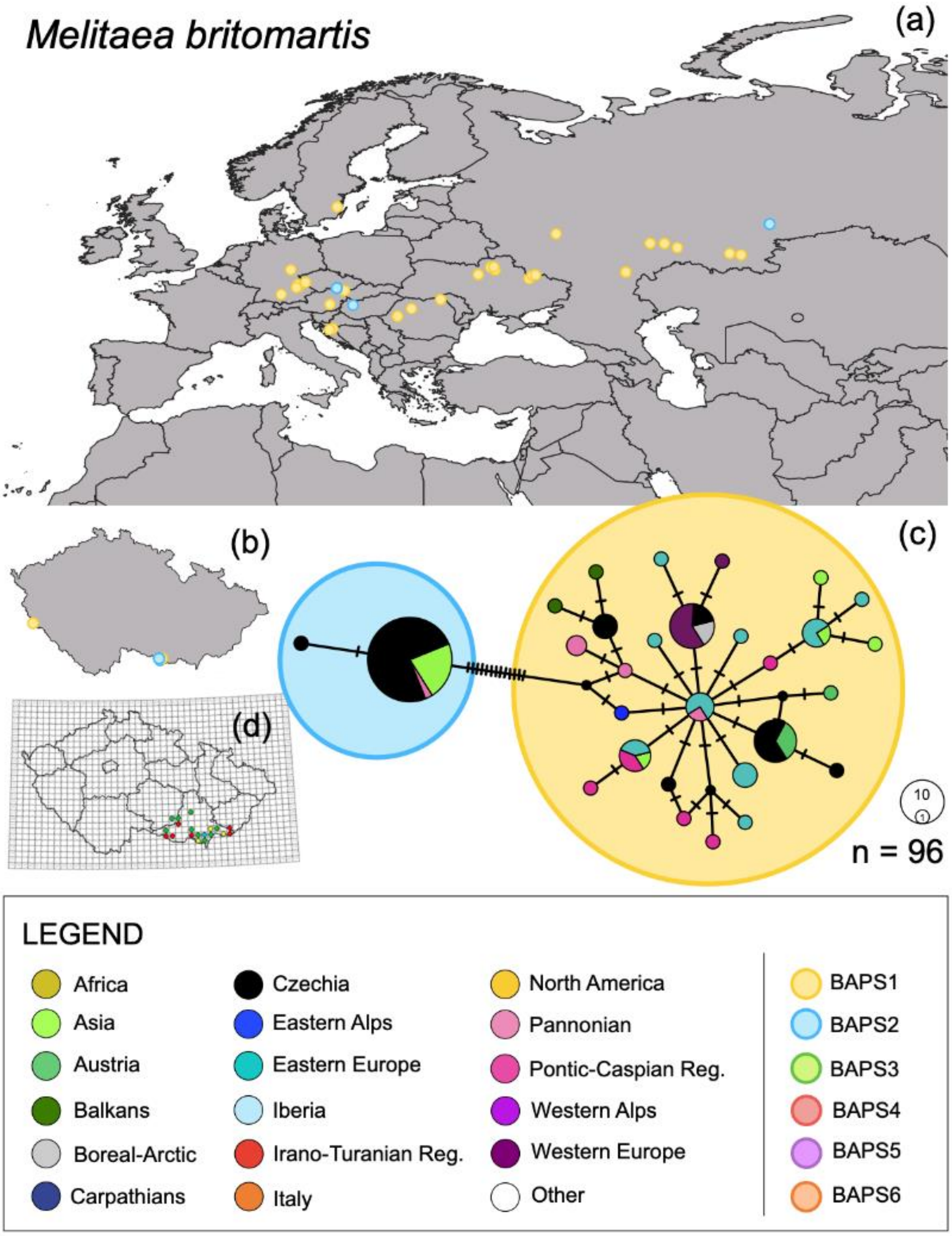


*Melitaea britomartis* Assmann, 1847: **Habitat:** xerothermophilic 2. **Red List:** CR

**BAPS:** BAPS-mix-East. **HAP:** HAP-mix

**Range:** Euro-Siberian. **Distribution CZ:** forest steppes, dry scrublands. Used to be more widespread in South Moravia, never confirmed from Bohemia in the past. It decreased severely in Moravia, and is present only at a few last localities (Hajkova et al., 2023; Vodičková et al., 2019). In the recent years, the species was recorded in West Bohemia (Upper Palatine Forest) for the first time, likely expanding from Germany (haplotypes shared with Western Europe – Germany, and Scandinavia) In Moravia, the two mitochondrial lineages with deep splits (BAPS1 and 2) are present in sympatry.

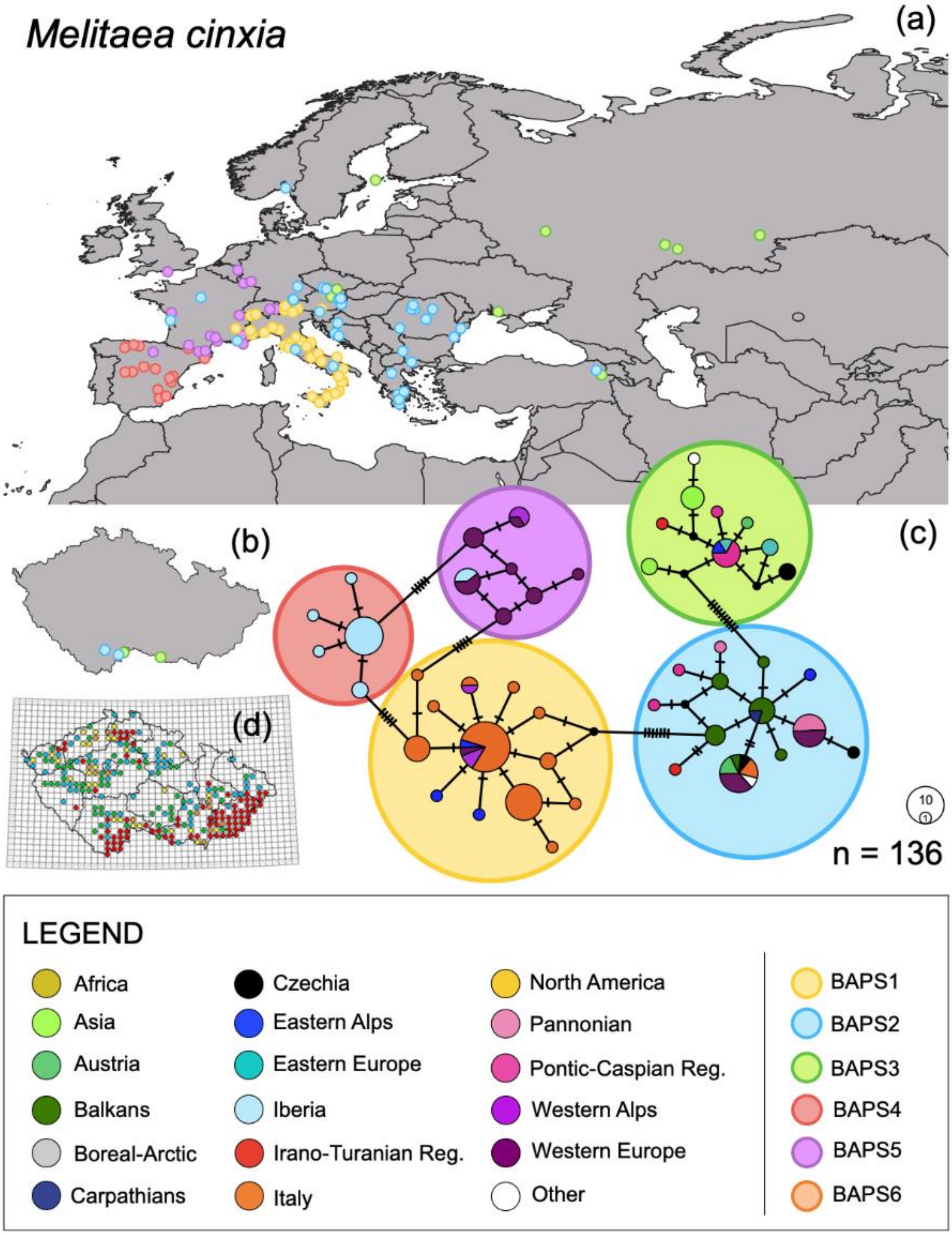


*Melitaea cinxia* (Linnaeus, 1758): **Habitat:** mesophilic 1. **Red List:** VU

**BAPS:** BAPS-Italy-Balkans + BAPS-temperate. **HAP:** HAP-Italy-Balkans + HAP-temperate

**Range:** Palearctic. **Distribution CZ:** dry grasslands and forest steppes in warm regions. Used to be more widely distributed in lowlands and highlands across the country, decreased due to land use changes. Now re-expanding again with warming climate. Two different mitochondrial lineages found in CZ, occurring likely in sympatry.

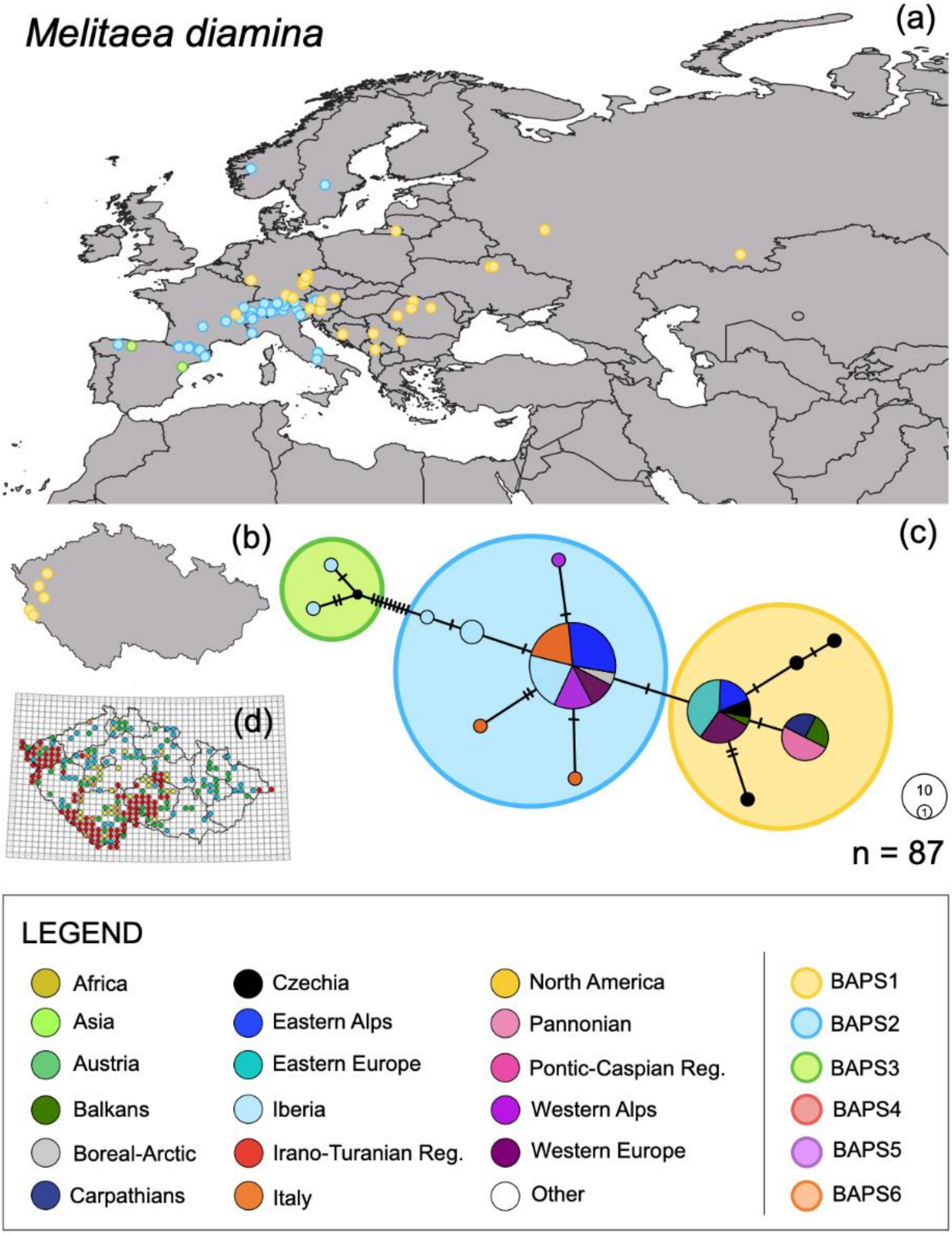


*Melitaea diamina* (Lang, 1789): **Habitat:** hydrophilic + tyrphophilic. **Red List:** VU
**BAPS:** BAPS-East-Balkans. **HAP:** HAP-East-Balkans
**Range:** Euro-Siberian. **Distribution CZ:** wet and peat meadows, more in highlands and mountains. Decreased with land use intensification, still present in mountainous parts of Bohemia; in Moravia present only in the east (Beskid Mts.).

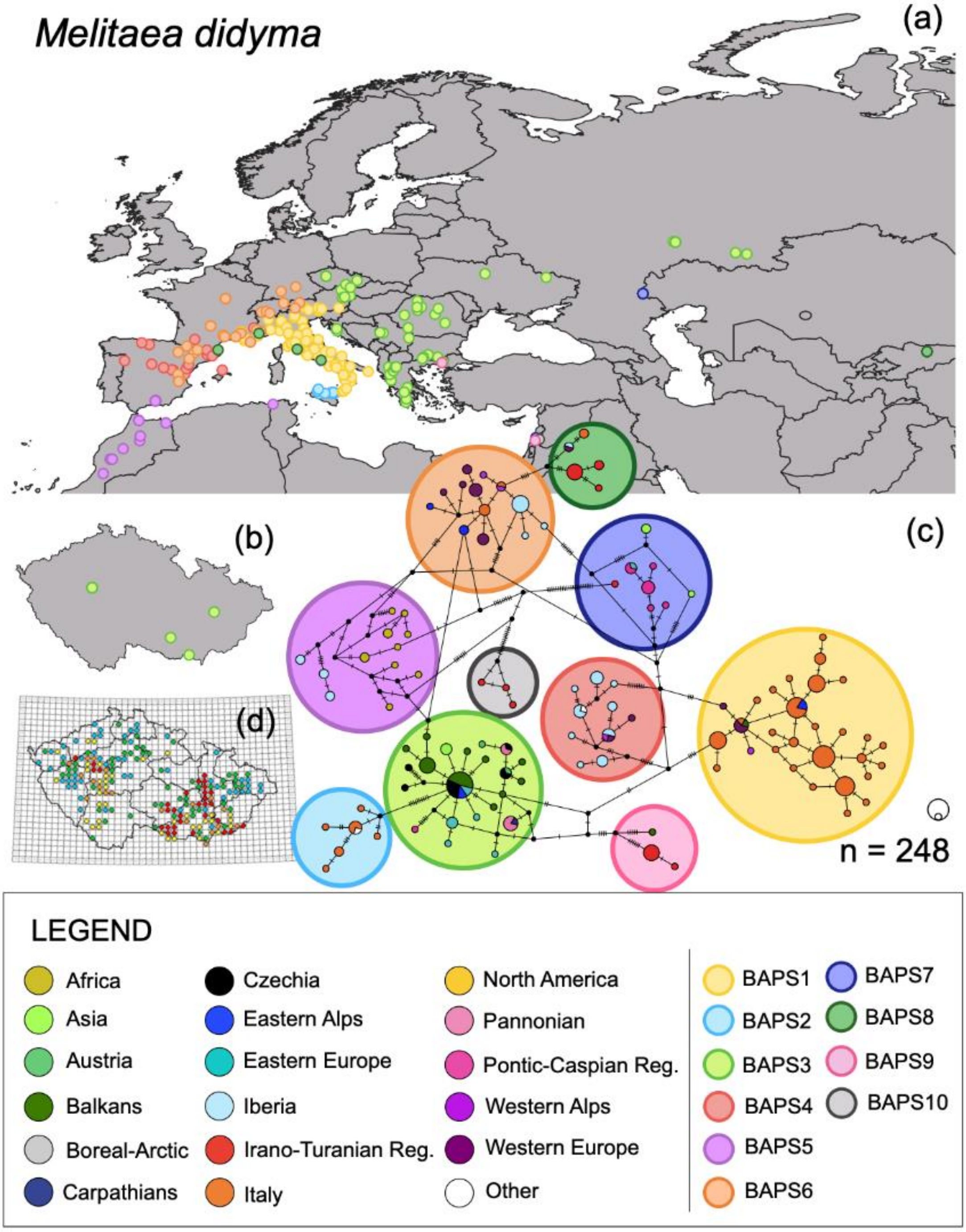


*Melitaea didyma* (Esper, 1778): **Habitat:** xerothermophilic 1. **Red List:** CR

**BAPS:** BAPS-East-Balkans. **HAP:** HAP-East-Balkans

**Range:** Western Palearctic. **Distribution CZ:** flowery dry short grasslands. Widespread across the country in the past, extinct from majority of the area. Now present in central Bohemia and in southern and central Moravia, currently re-expanding again with warmer climate. Czech samples belong to sp./ssp. *neera* (Pazhenkova & Lukhtanov, 2016). The East/Balkans lineage inhabiting CZ was also recovered by a genome-wide marker (Dincă, Lee, Vila, & Mutanen, 2019).

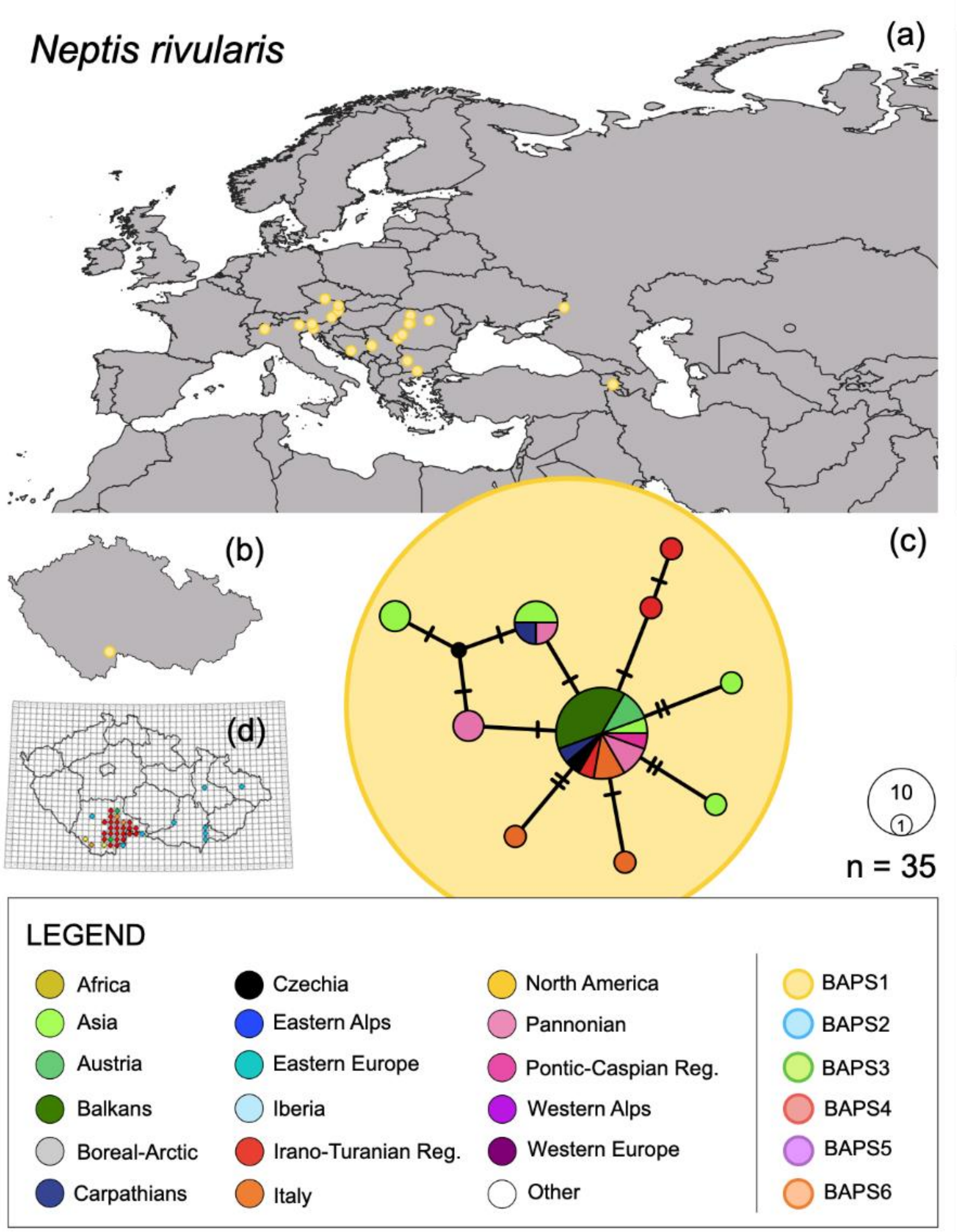


*Neptis rivularis* (Scopoli, 1763): **Habitat:** hydrophilic. **Red List:** NT

**BAPS:** BAPS-low. **HAP:** HAP-wide-low

**Range:** Euro-Siberian. **Distribution CZ:** alluvial meadows and forest margins. In CZ, specific for South Bohemia and closest surroundings, where it expands with its host plant *Spiraea* spp. used as ornamental bushes; extinct from Moravia. From the Czech population, also specimens with double peaks (degenerate bases) were obtained (N=3), these are not included in analyses, as this probably represent heteroplasmy or similar issue.

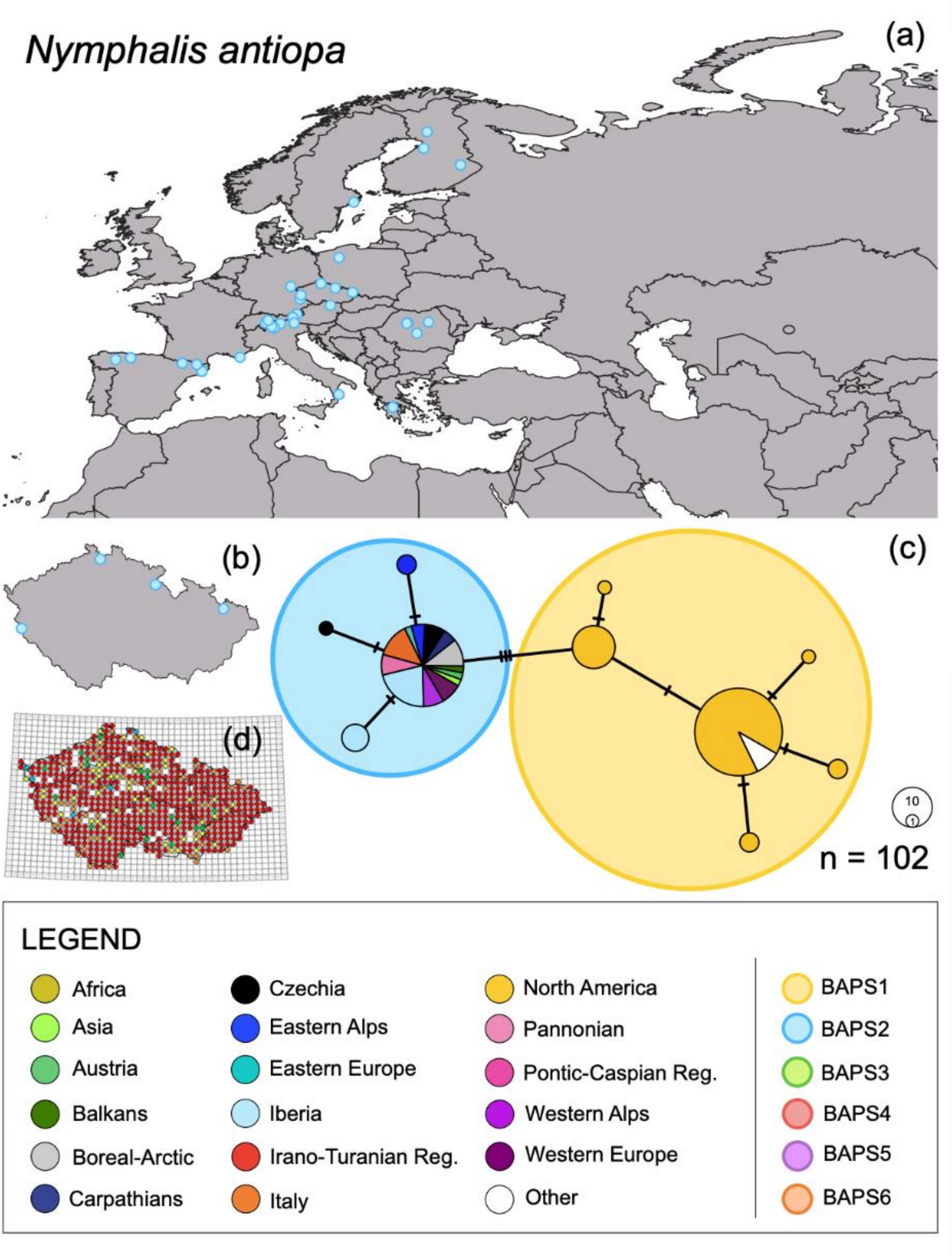


*Nymphalis antiopa (Linnaeus, 1758)*: **Habitat:** mesophilic 3. **Red List:** LC

**BAPS:** BAPS-low. **HAP:** HAP-low

**Range:** Holarctic. **Distribution CZ:** woodland clearings and margins. Widespread across the country.

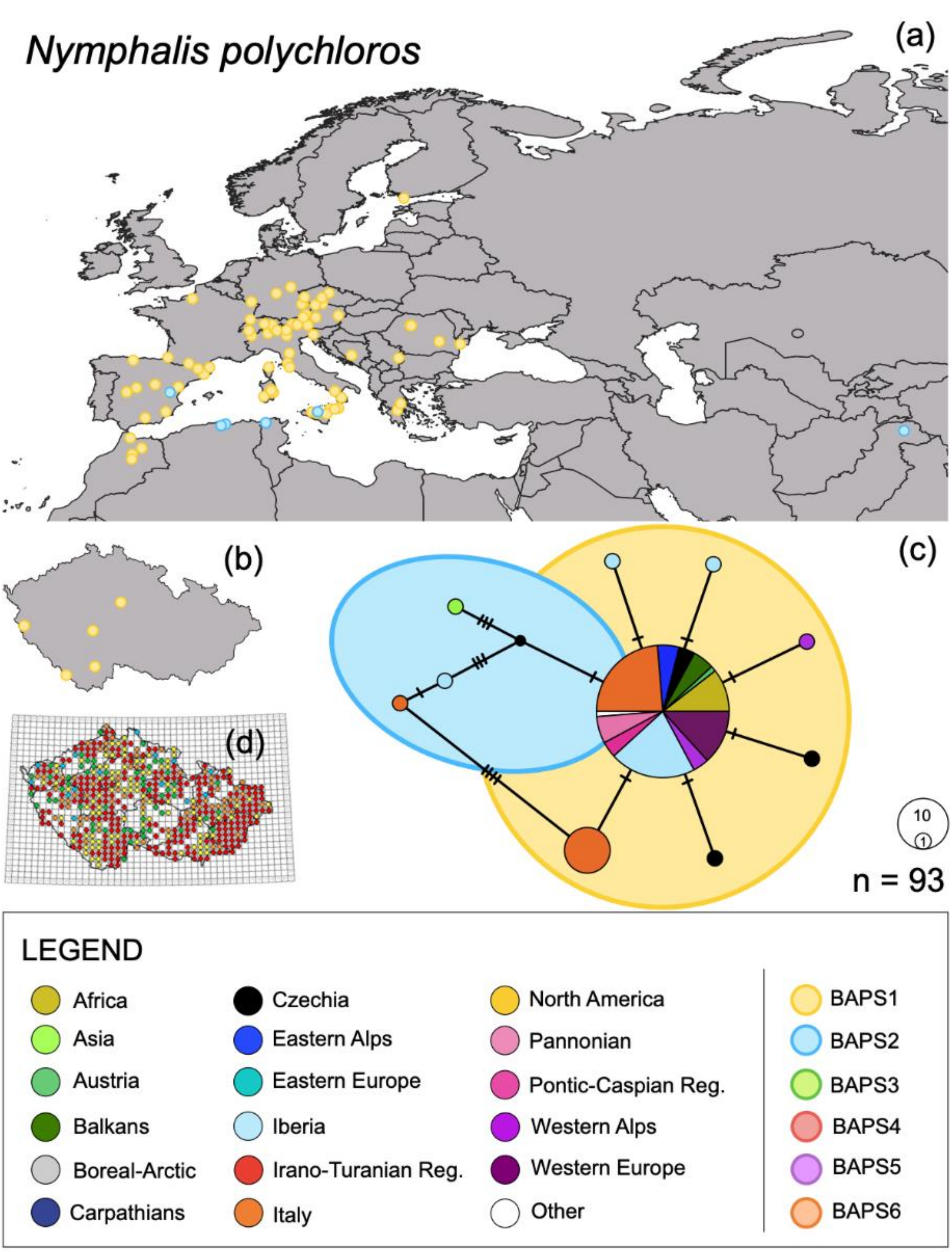


*Nymphalis polychloros* (Linnaeus, 1758): **Habitat:** mesophilic 3. **Red List:** LC

**BAPS:** BAPS-wide. **HAP:** HAP-wide-low

**Range:** Western Palearctic. **Distribution CZ:** woodland clearings and margins, alluvial woodlands. Widespread across the country.

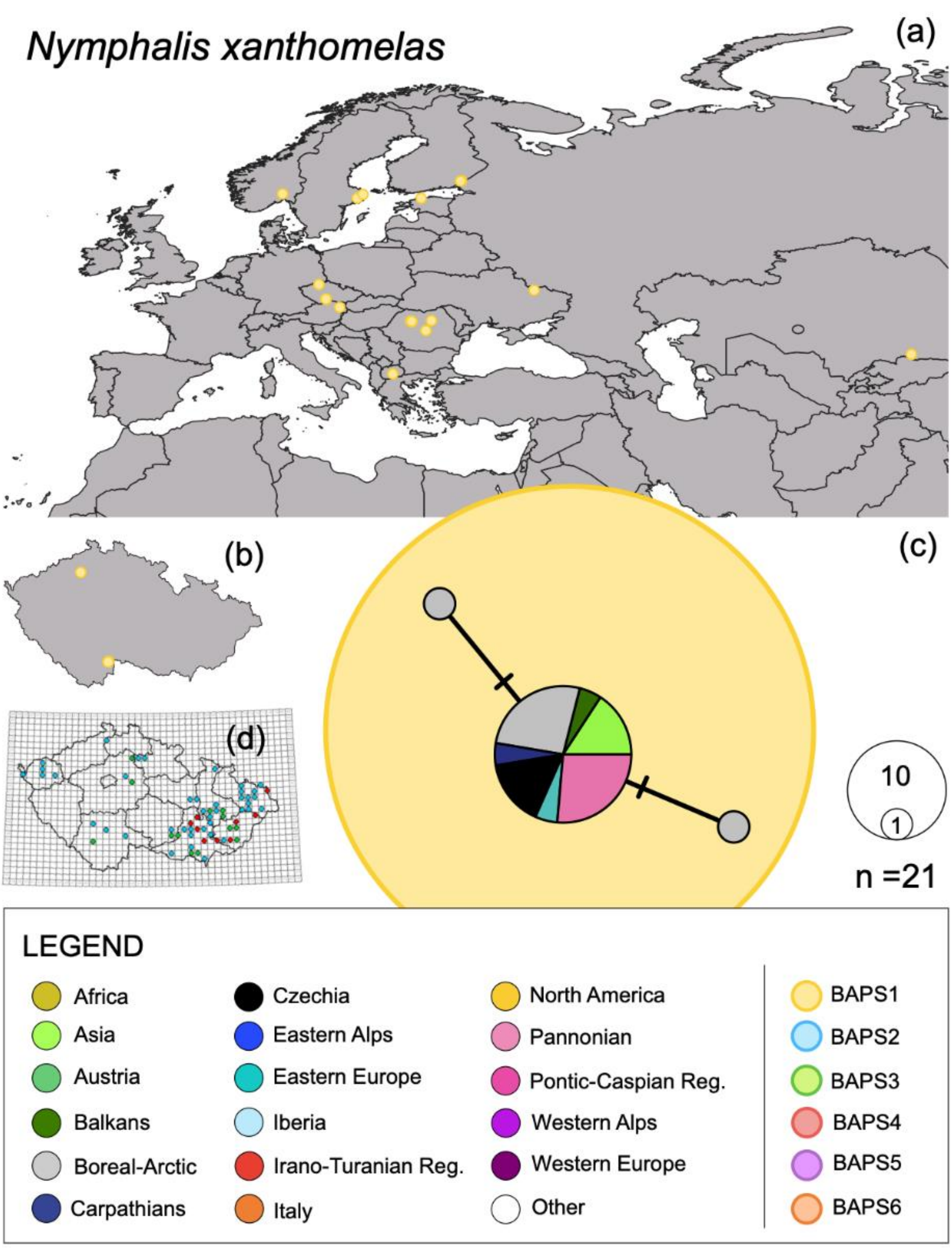


*Nymphalis xanthomelas* (Esper, 1781): **Habitat:** mesophilic 3. **Red List:** LC

**BAPS:** BAPS-low. **HAP:** HAP-low

**Range:** Euro-Siberian. **Distribution CZ:** alluvial woodlands. Extinct in the past (1960s), but recently recolonizing the country, with records across the country. Most regularly in South Bohemia (Třeboň Basin).

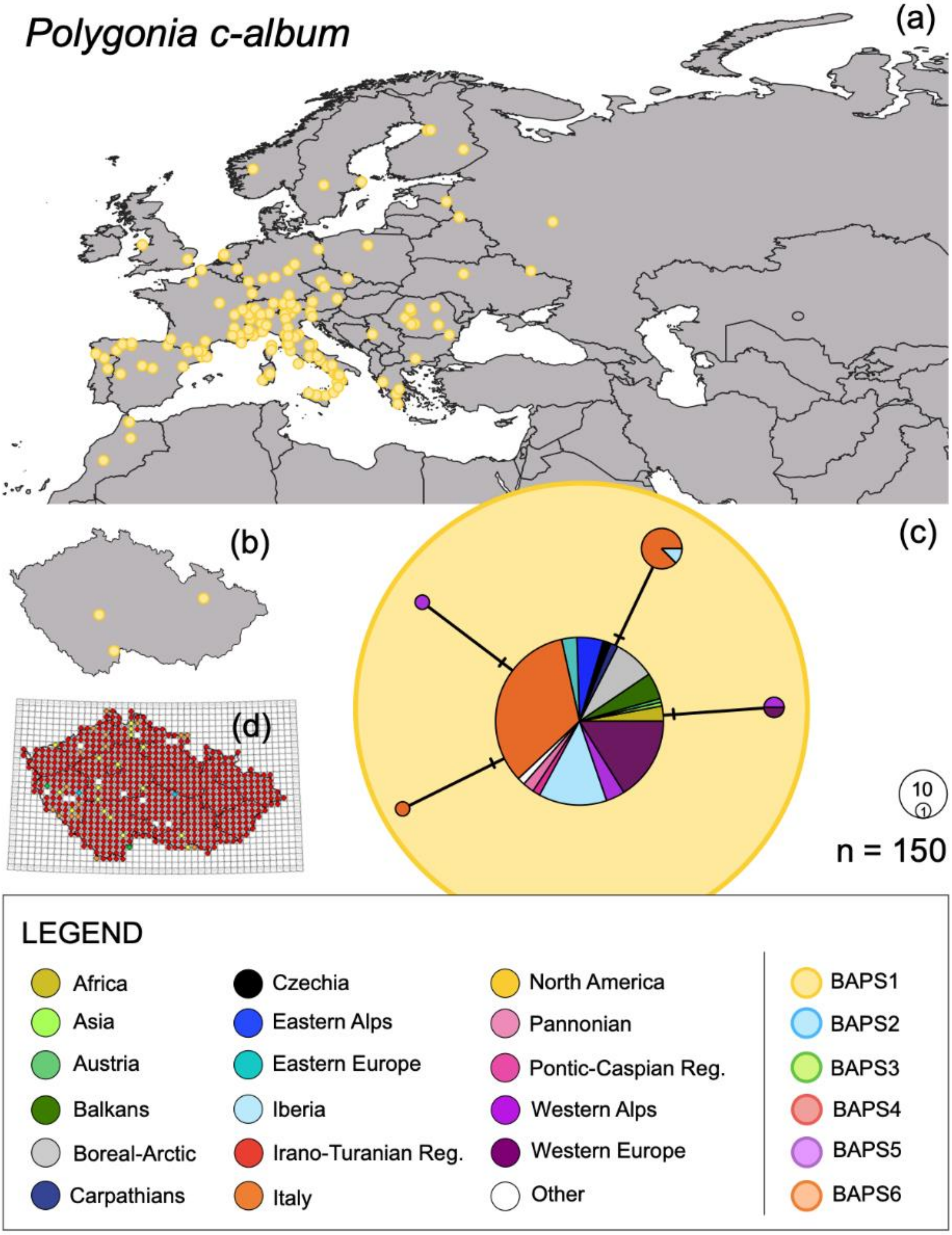


*Polygonia c-album* (Linnaeus, 1758): **Habitat:** mesophilic 3. **Red List:** LC

**BAPS:** BAPS-low. **HAP:** HAP-low

**Range:** Palearctic. **Distribution CZ:** woodland clearings and margins, gallery vegetation. Widespread across the country.

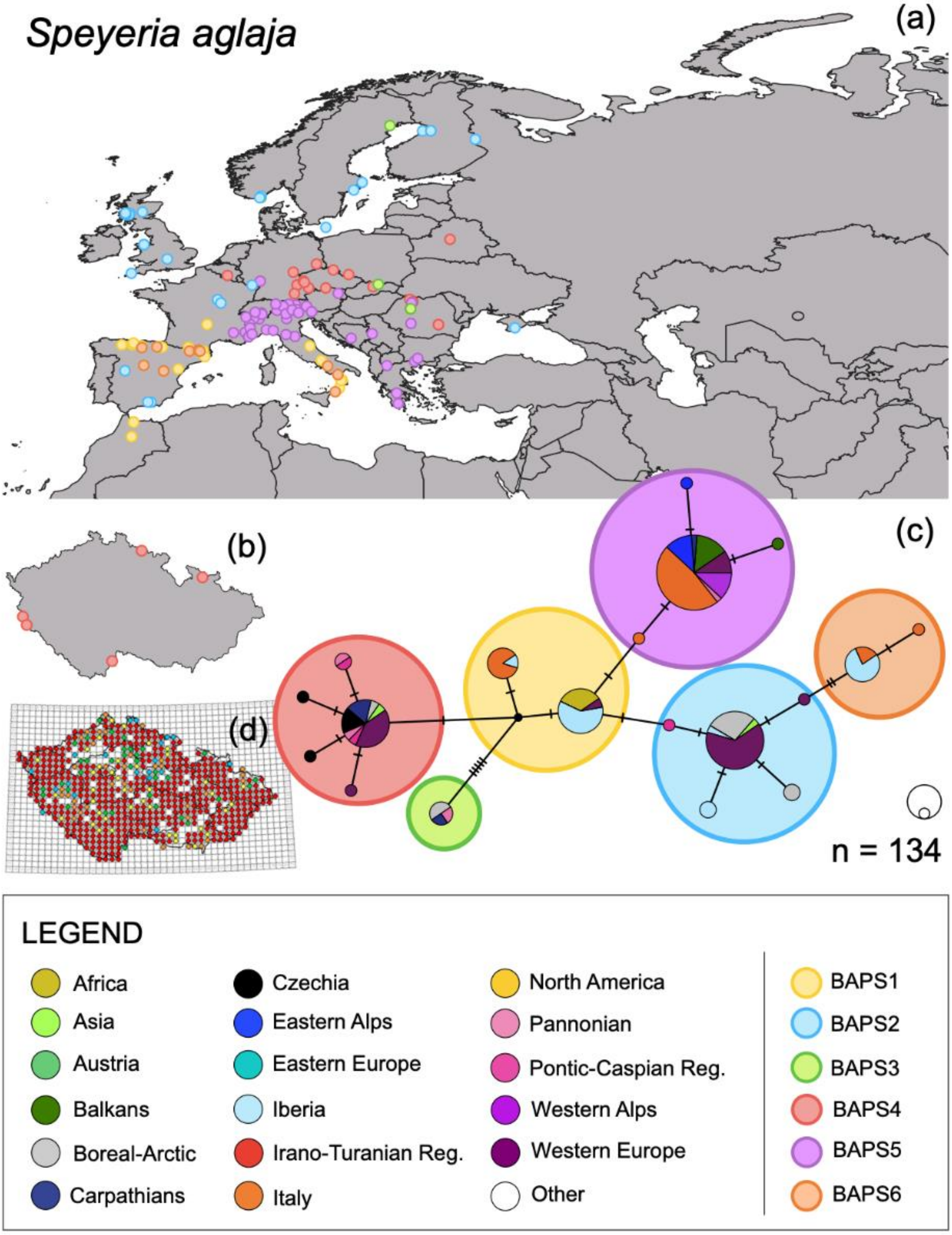


*Speyeria aglaja* (Linnaeus, 1758): **Habitat:** mesophilic 2. **Red List:** LC

**BAPS:** BAPS-Central. **HAP:** HAP-Central

**Range:** Palearctic. **Distribution CZ:** woodland clearings and margins, meadows, scrublands. Widespread across the country but decreasing.

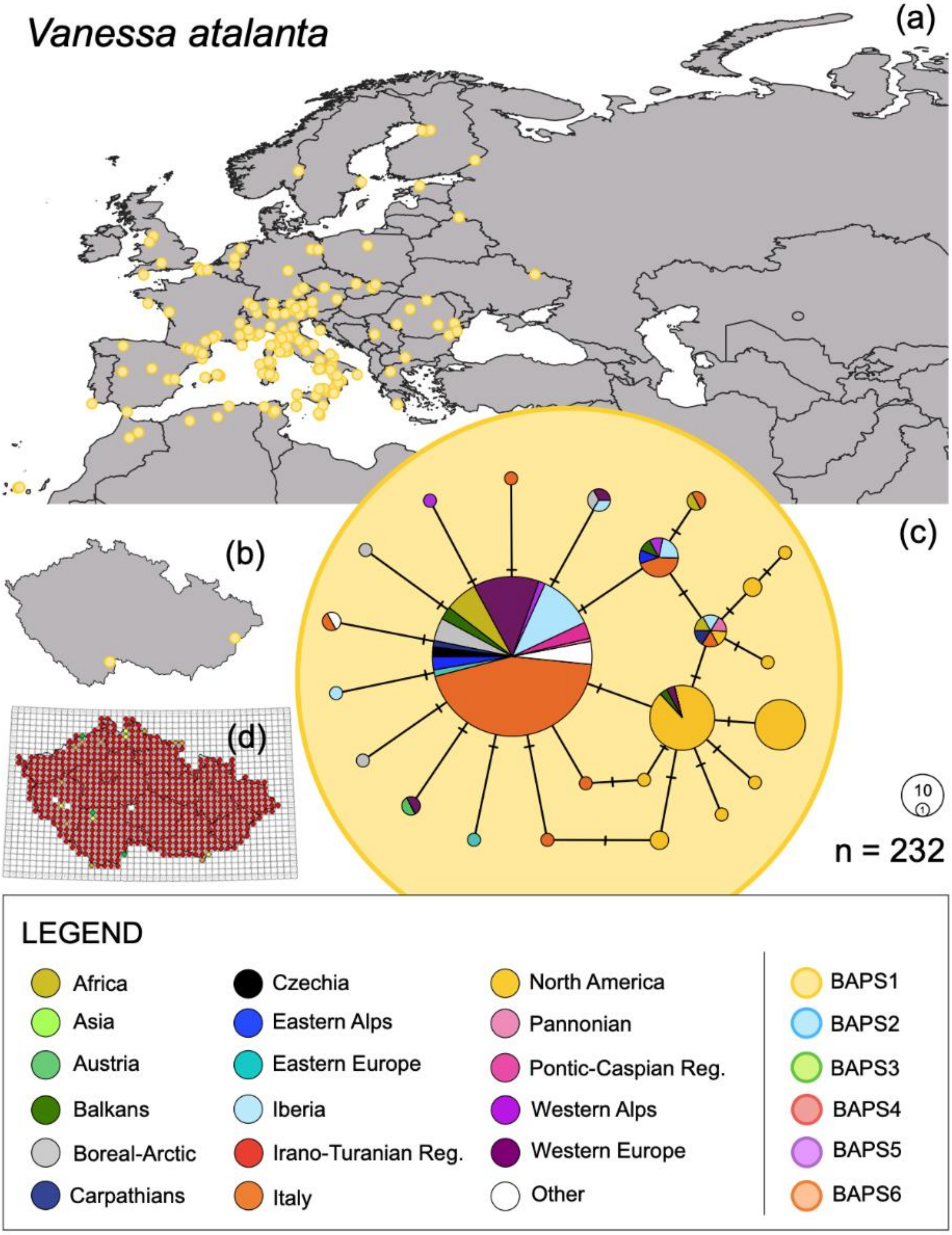


*Vanessa atalanta* (Linnaeus, 1758): **Habitat:** ubiquitous. **Red List:** LC

**BAPS:** BAPS-low. **HAP:** HAP-wide-complex

**Range:** Holarctic. **Distribution CZ:** widespread and common generalist, migrant from the Mediterranean.

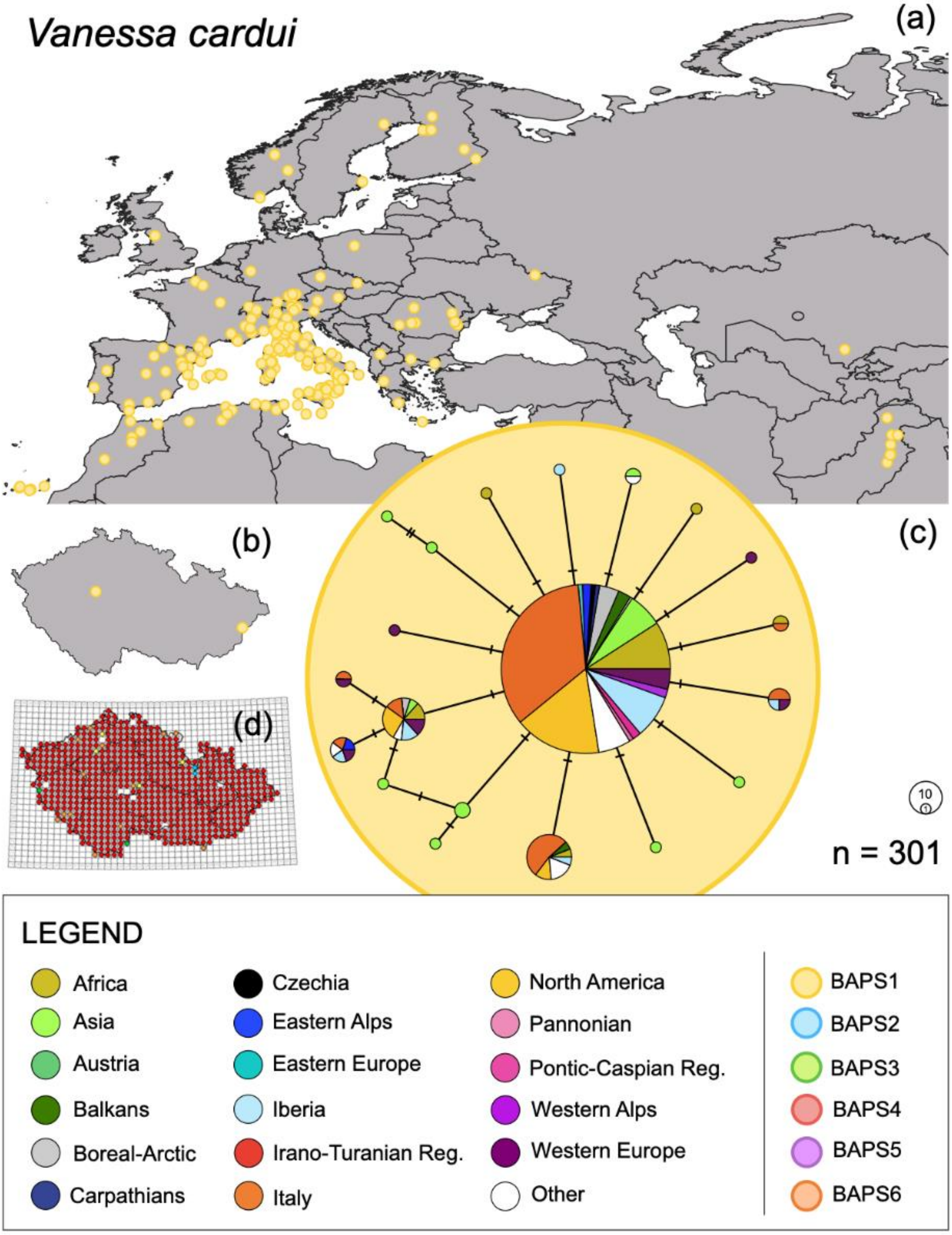


*Vanessa cardui* (Linnaeus, 1758): **Habitat:** ubiquitous. **Red List:** LC

**BAPS:** BAPS-low. **HAP:** HAP-wide-expand

**Range:** Cosmopolite. **Distribution CZ:** widespread and common migrant from the Mediterranean and Africa.

### Nymphalidae: Satyrinae

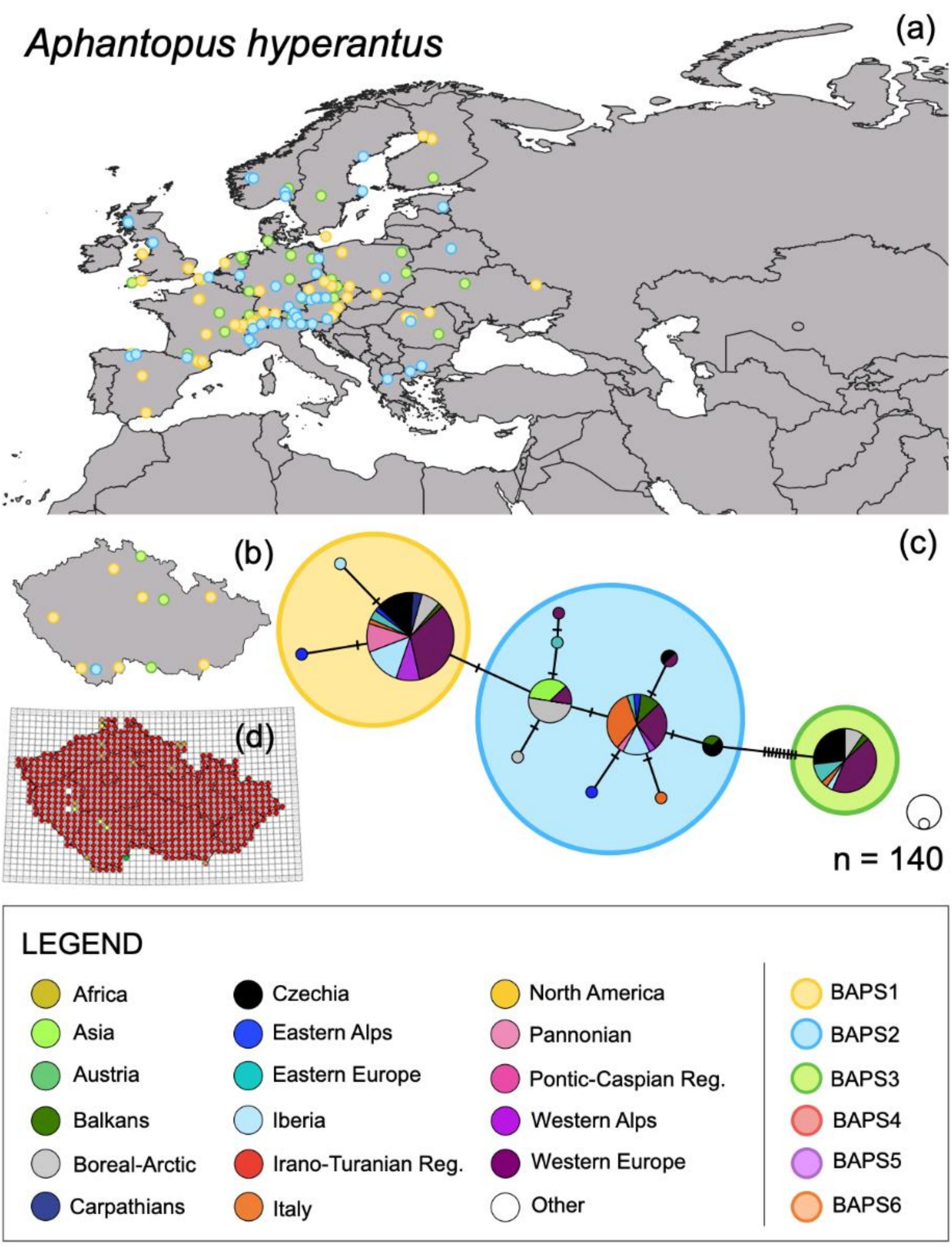


*Aphantopus hyperantus* (Linnaeus, 1758): **Habitat:** mesophilic 1. **Red List:** LC

**BAPS:** BAPS-mix. **HAP:** HAP-mix

**Range:** Palearctic. **Distribution CZ:** various meadow habitats. Widespread and common.

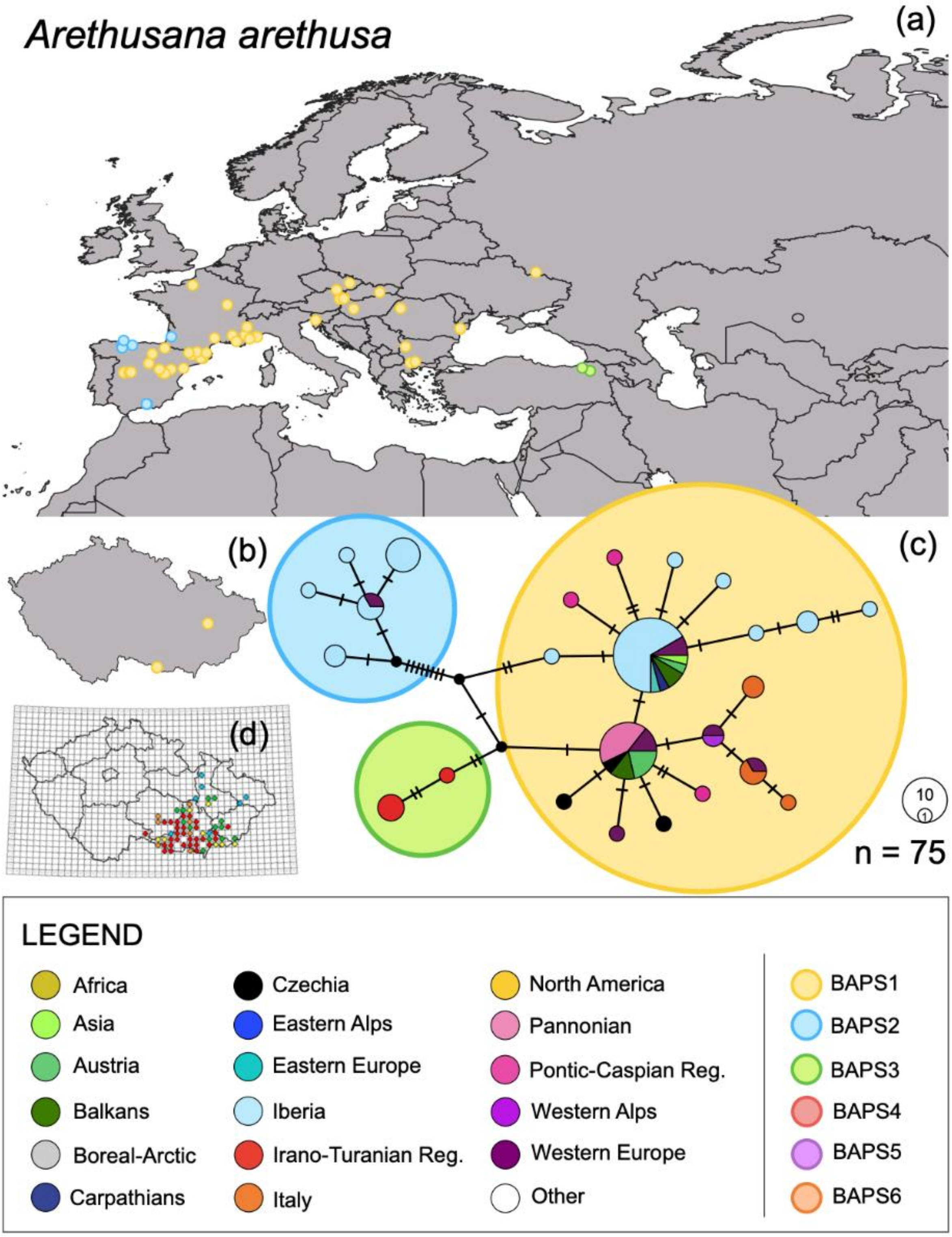


*Arethusana arethusa* ([Denis & Schiffermüller], 1775): **Habitat:** xerothermophilic 2. **Red List:** VU

**BAPS:** BAPS-wide. **HAP:** HAP-wide-complex

**Range:** Western Palearctic. **Distribution CZ:** dry grasslands and scrublands. It occurred historically and recently only in Moravia (Pannonia), decreasing.

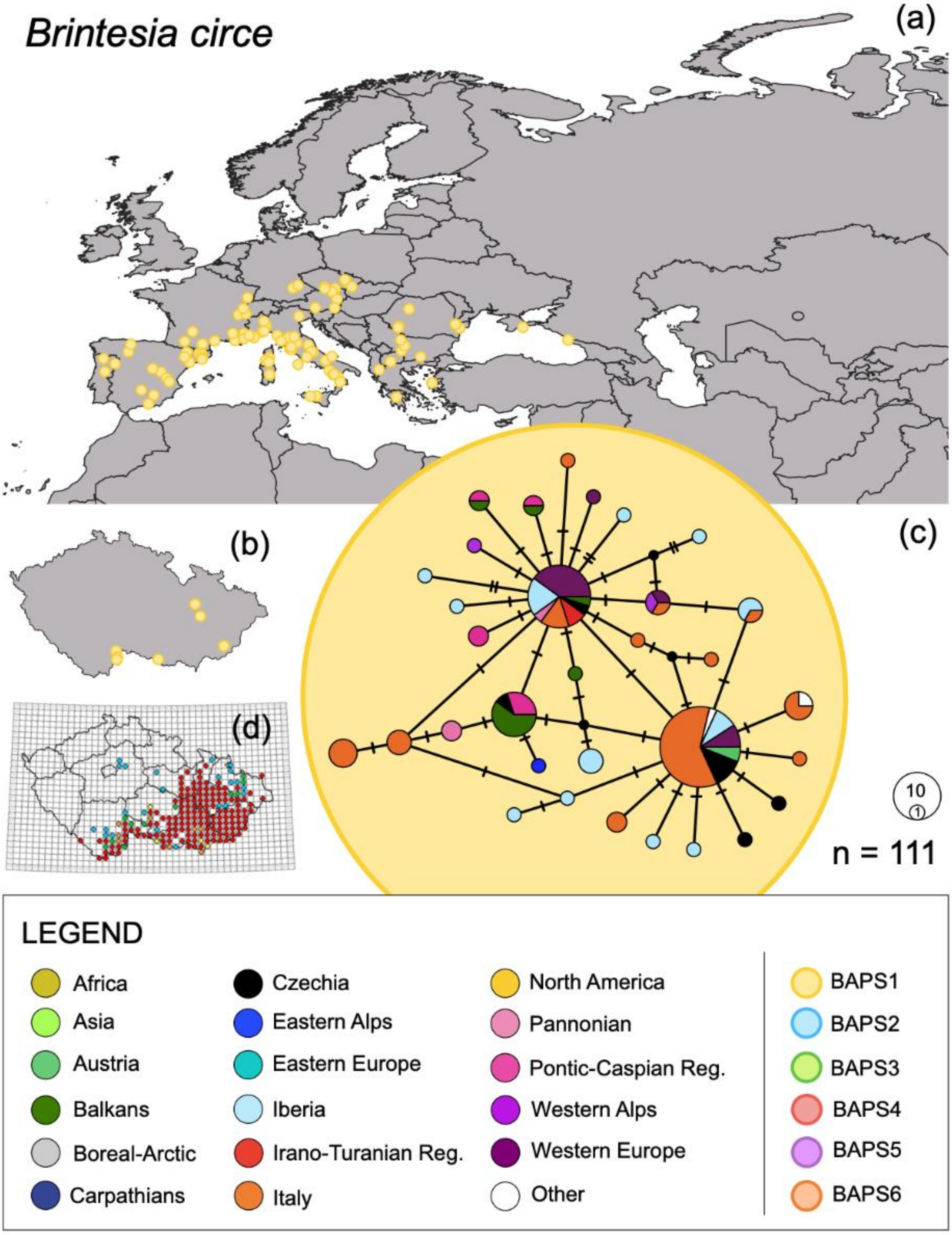


*Brintesia circe* (Fabricius, 1775): **Habitat:** xerothermophilic 2. **Red List:** LC

**BAPS:** BAPS-low. **HAP:** HAP-mix

**Range:** Western Palearctic. **Distribution CZ:** dry grasslands and forest steppes, park landscapes. Decreased during 20th century to the south and east, but expanding again. Widespread across Moravia and South Bohemia, mostly absent from other parts of Bohemia.

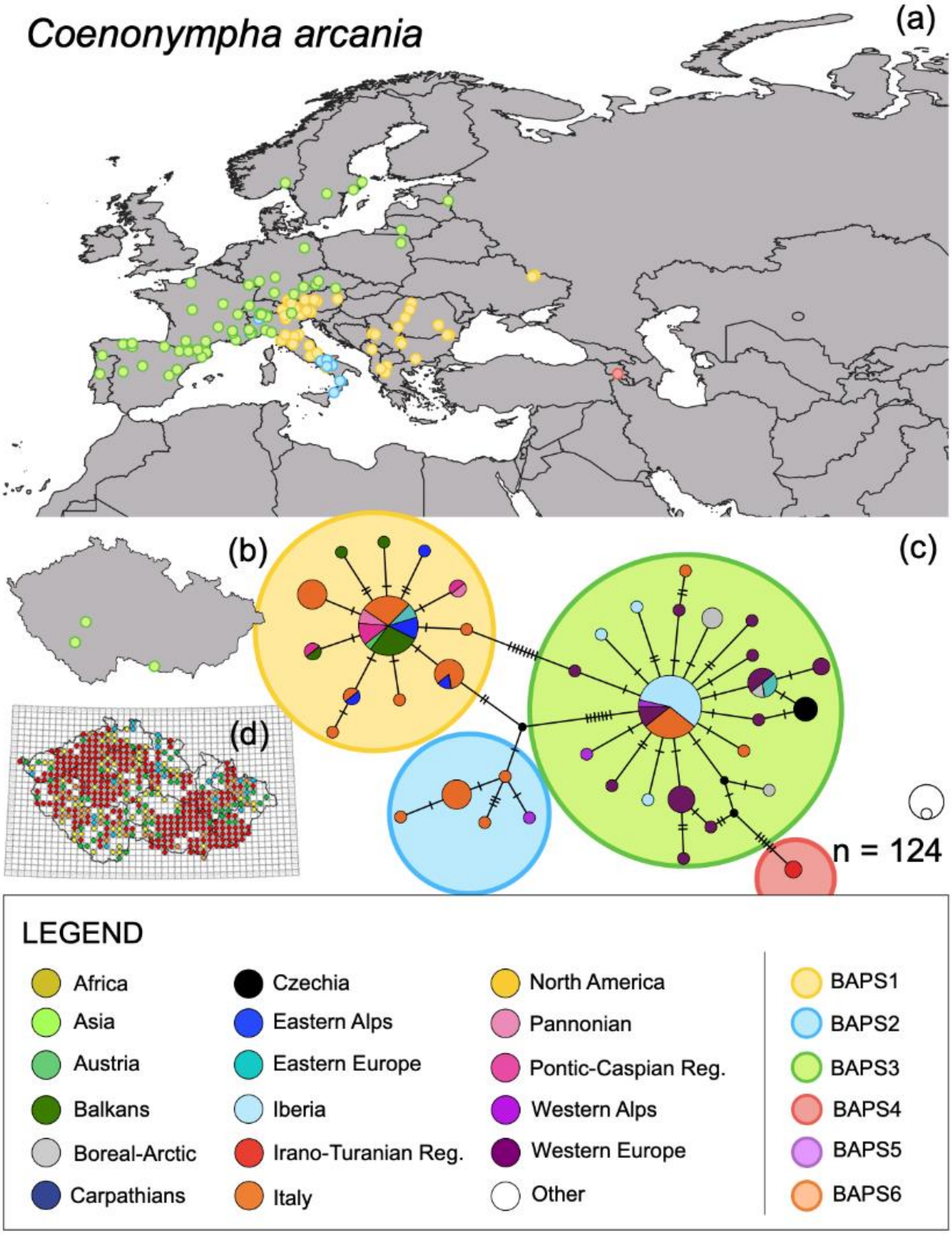


*Coenonympha arcania* (Linnaeus, 1760): **Habitat:** mesophilic 2 + xerothermophilic 2. **Red List:** NT

**BAPS:** BAPS-West. **HAP:** HAP-West

**Range:** European. **Distribution CZ:** woodland margins, deciduous open woodlands. Distributed across entire country in lowlands and highlands, but decreased from agricultural landscapes. The contact zone between BAPS1 (Italy-Balkans lineage) and BAPS2 (West lineage) is probably situated in Moravia or Slovakia – further sampling is needed to confirm the location of the contact zone.

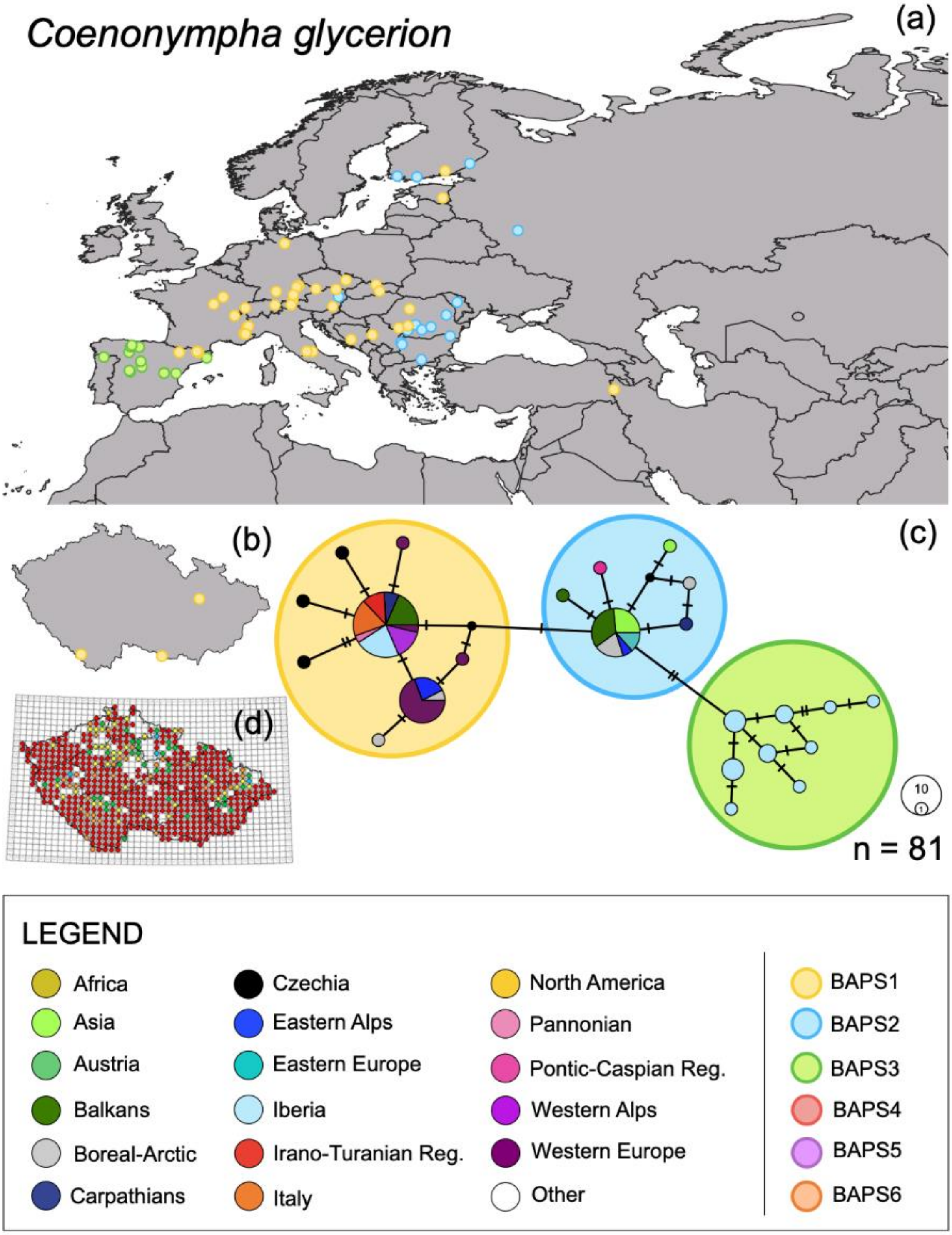


*Coenonympha glycerion* (Borkhausen, 1788): **Habitat:** xerothermophilic 2 + hydrophilic.

**Red List:** LC

**BAPS:** BAPS-wide. **HAP:** HAP-wide-low

**Range:** Euro-Siberian. **Distribution CZ:** shrublands and open woodlands, but also wet and boggy meadows. Widespread across almost entire country, decreased from intensive agricultural landscapes.

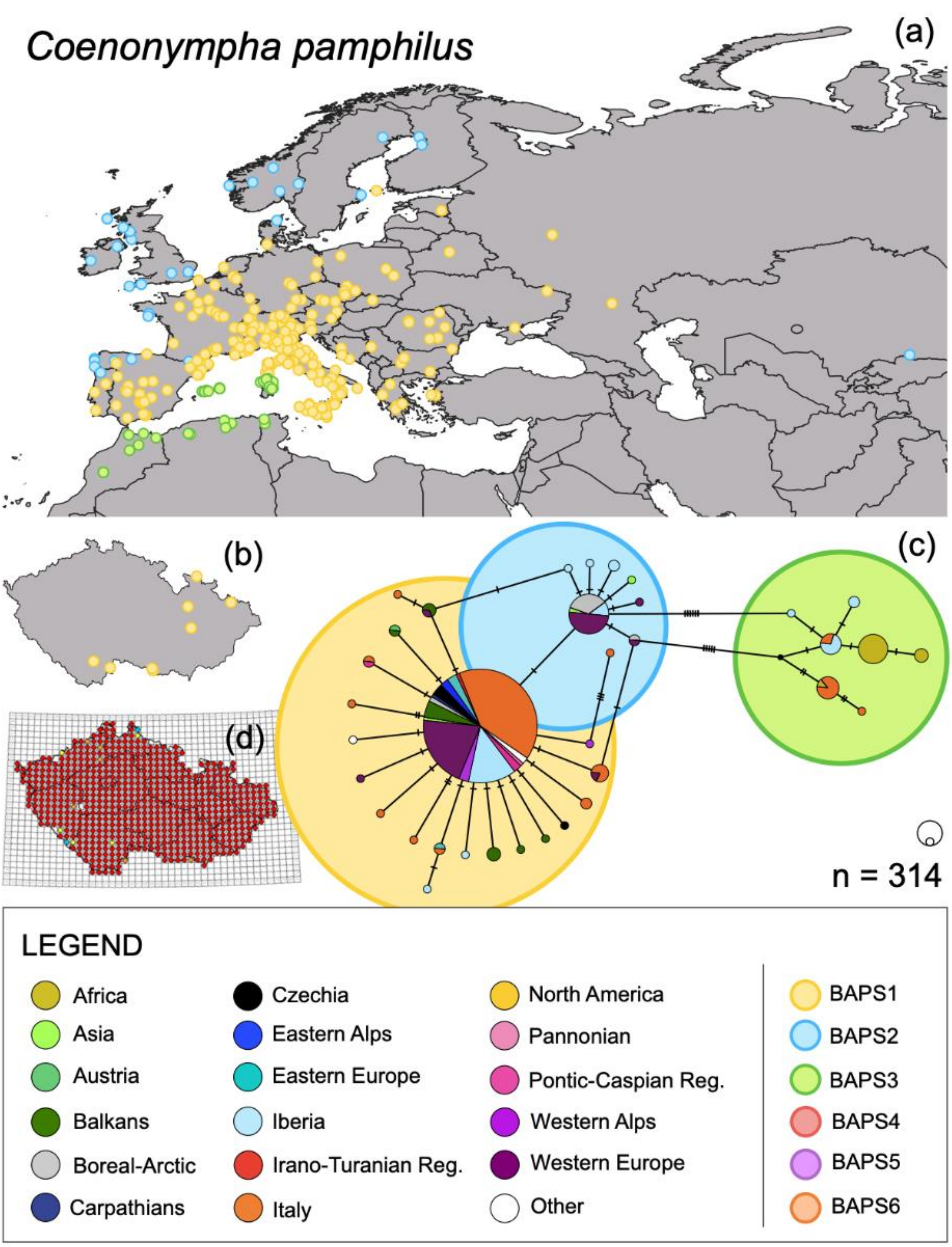


*Coenonympha pamphilus* (Linnaeus, 1758): **Habitat:** ubiquitous. **Red List:** LC

**BAPS:** BAPS-wide. **HAP:** HAP-wide-expand

**Range:** Western Palearctic. **Distribution CZ:** all types of meadow habitats. Widespread and common.

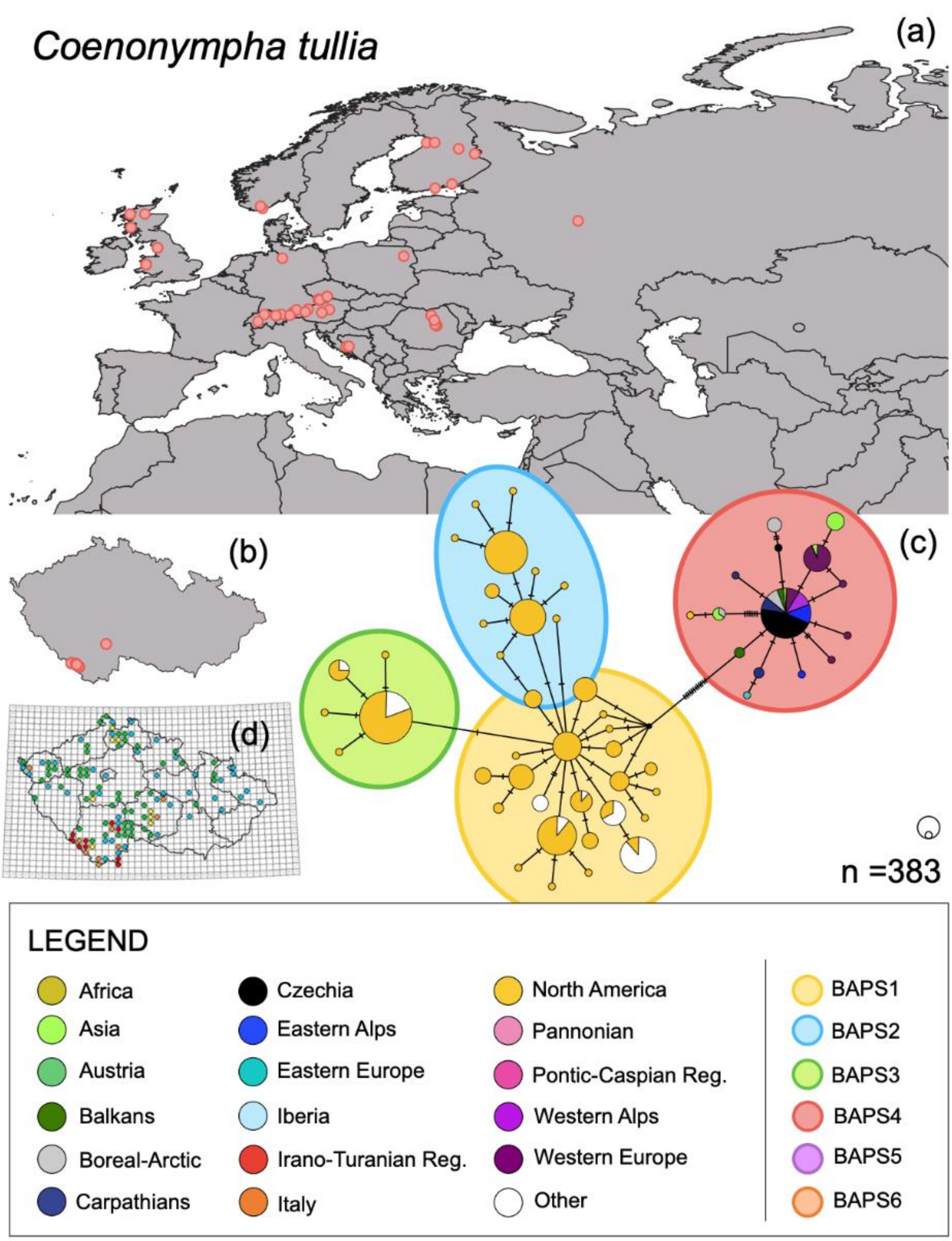


*Coenonympha tullia* (Müller, 1764): **Habitat:** tyrphophilic. **Red List:** CR

**BAPS:** BAPS-low. **HAP:** HAP-temperate (such distribution)

**Range:** Holarctic. **Distribution CZ:** Peat bogs. It used to be spread in such habitats across the country, but extinct from almost all its former localities due to afforestation and meliorations. Currently present only in Bohemian Forest and Třeboň Basin.

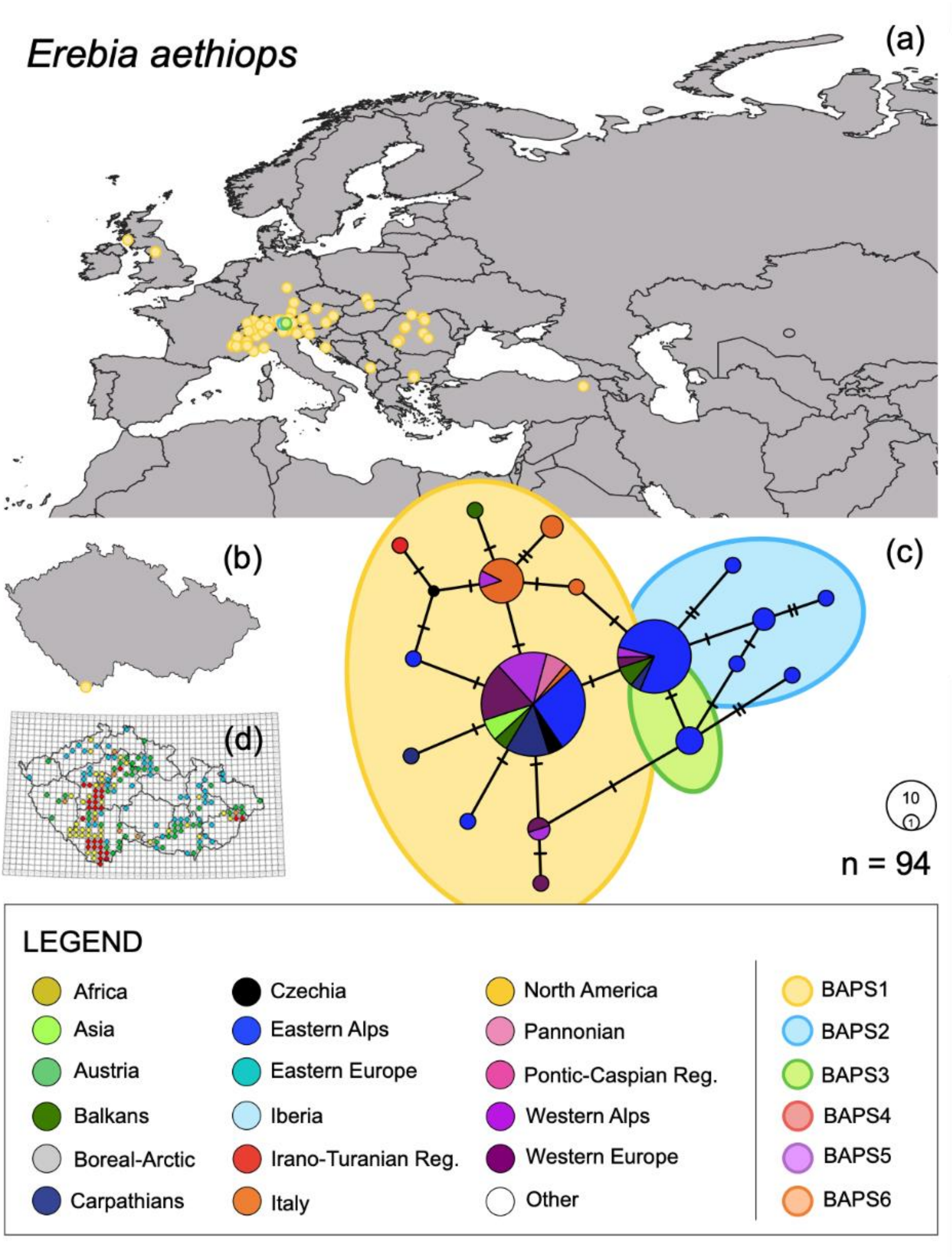


*Erebia aethiops* (Esper, 1777): **Habitat:** mesophilic 2 + xerothermophilic 2. **Red List:** EN

**BAPS:** BAPS-wide. **HAP:** HAP-wide-complex

**Range:** Euro-Siberian. **Distribution CZ:** forest steppes and scrublands in piedmont areas. Decreasing, extinct from Moravia, present only in southern and central Bohemia.

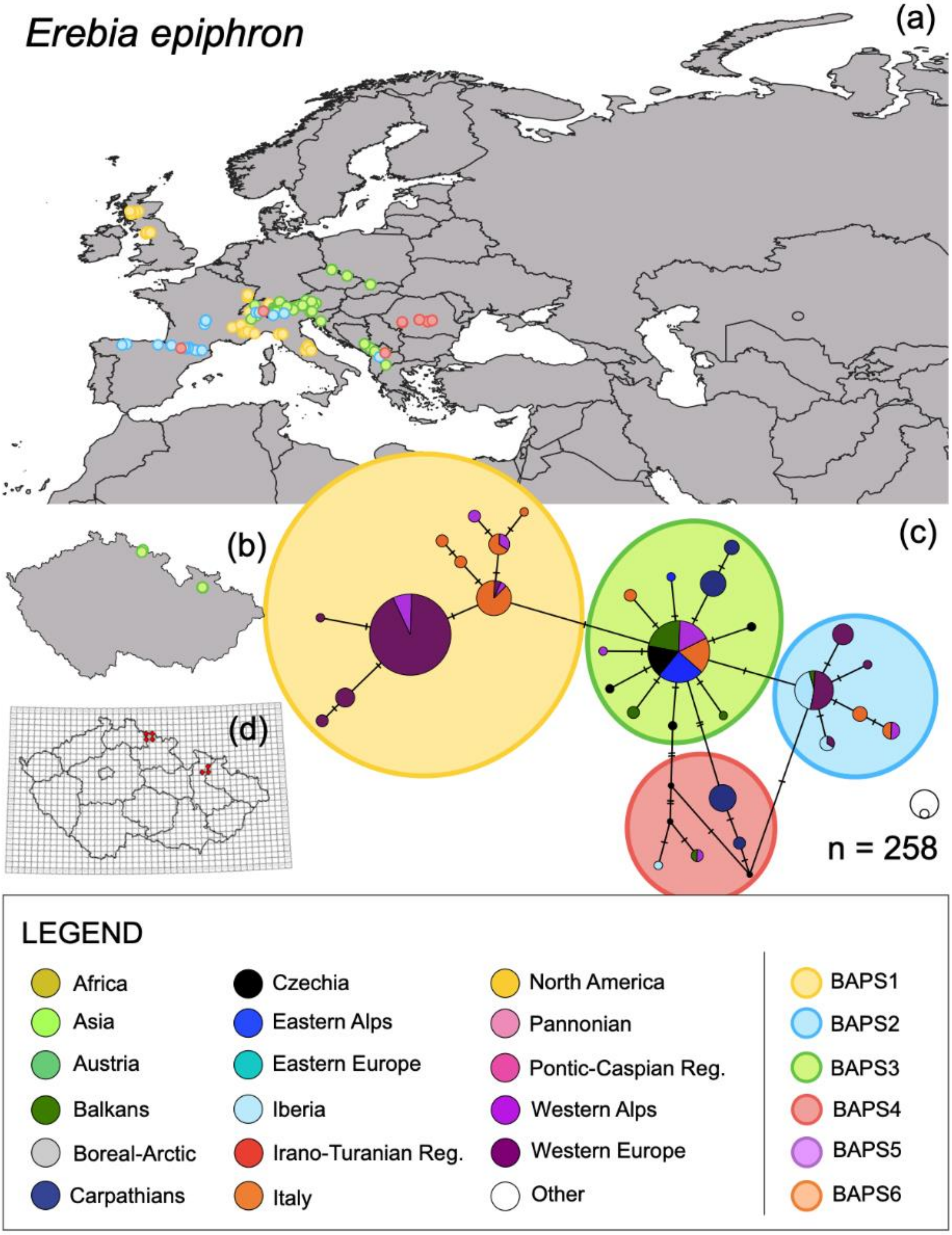


*Erebia epiphron* (Knoch, 1783): **Habitat:** alpine. **Red List:** LC

**BAPS:** BAPS-Italy-Balkans. **HAP:** HAP-Italy-Balkans

**Range:** European. **Distribution CZ:** alpine zone of mountains. It naturally occurs in High Ash Mountains (Hrubý Jeseník), from where it was introduced to Giant Mountains (Krkonoše) in 1930s (Schmitt, Cizek, & Konvicka, 2005).

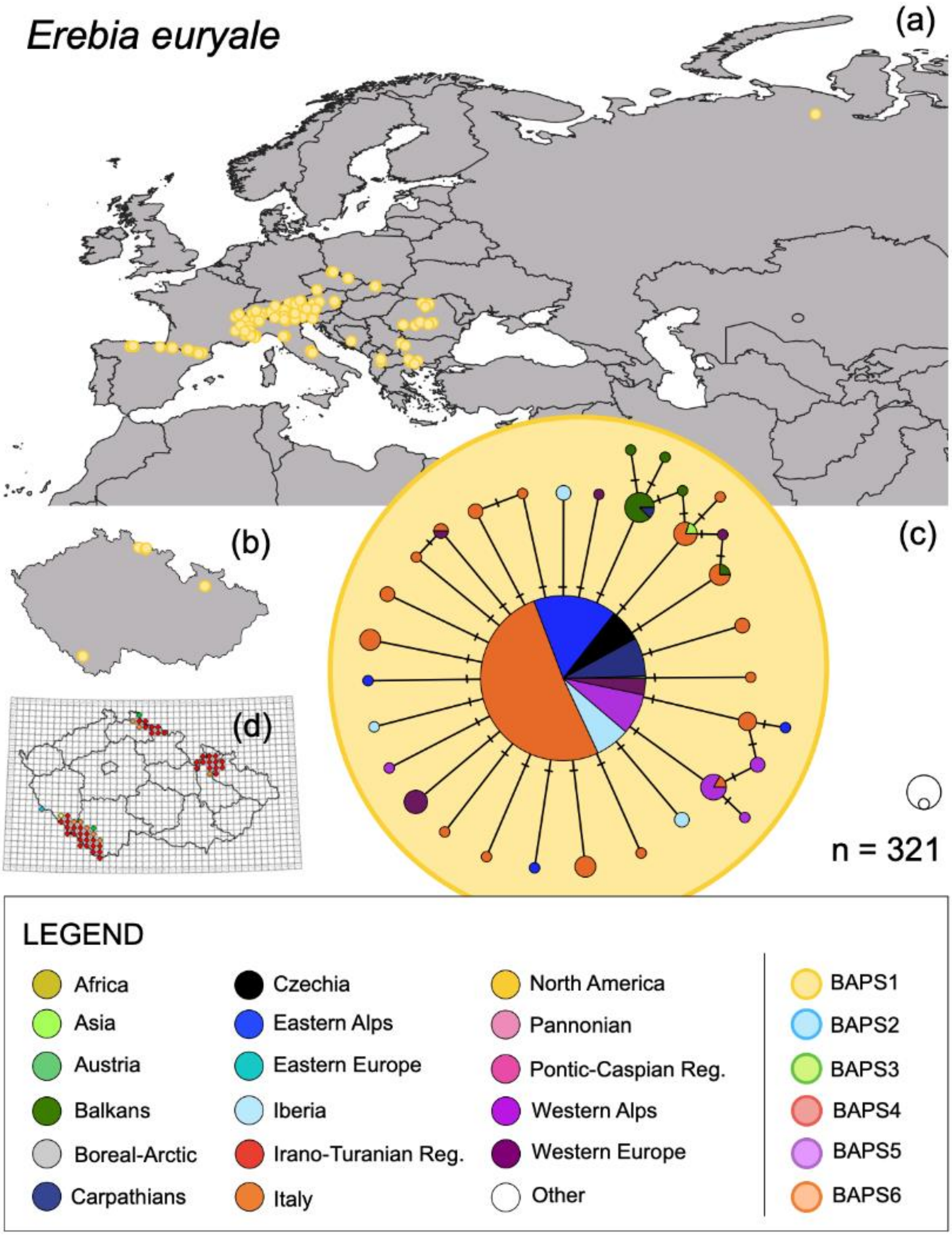


*Erebia euryale* (Esper, 1805): **Habitat:** mesophilic 3. **Red List:** LC

**BAPS:** BAPS-low. **HAP:** HAP-wide-expand

**Range:** European. **Distribution CZ:** clearings in mountain woodlands, mountain forest meadows. Restricted to several mountain systems along northern and southwestern borders of the country. *Erebia euryale* shares haplotypes with *E. ligea*. When treated together in the analysis, the BAPS revealed two groups: one consisting only of *E. ligea* (Fennoscandia, Italy, Alps, Balkans, Carpathians and Sudeten Mts.), the other of a combination of *E. euryale* and *E. ligea* (Pyrenees, Alps, Carpathians, Balkans, Sudeten Mts.)

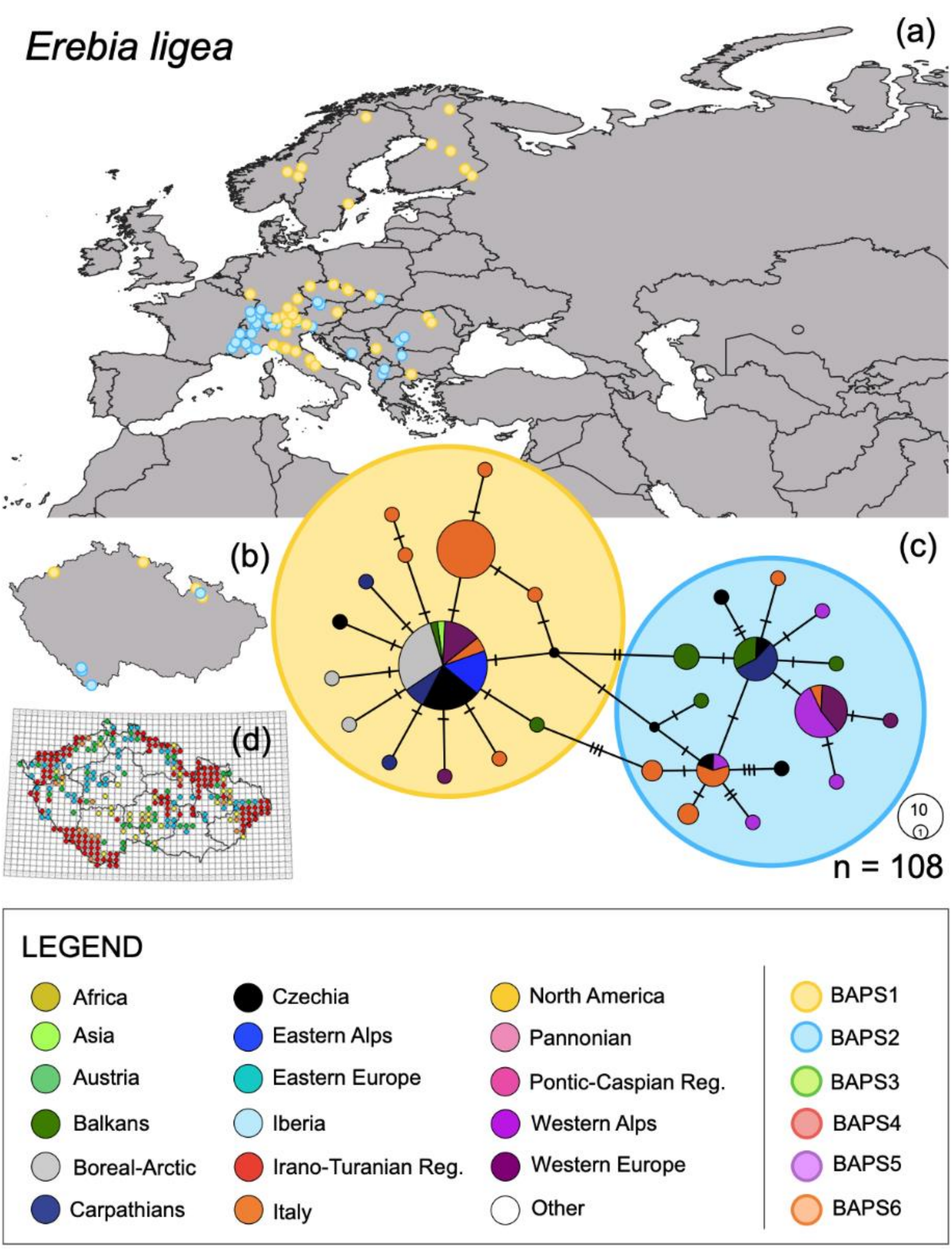


*Erebia ligea* (Linnaeus, 1758): **Habitat:** mesophilic 3. **Red List:** NT

**BAPS:** BAPS-mix-East. **HAP:** HAP-mix

**Range:** Euro-Siberian. **Distribution CZ:** clearings in mountain woodlands, mountain river alluvia. Present in borderland mountains, but also in highlands. Decreased in lower altitudes.

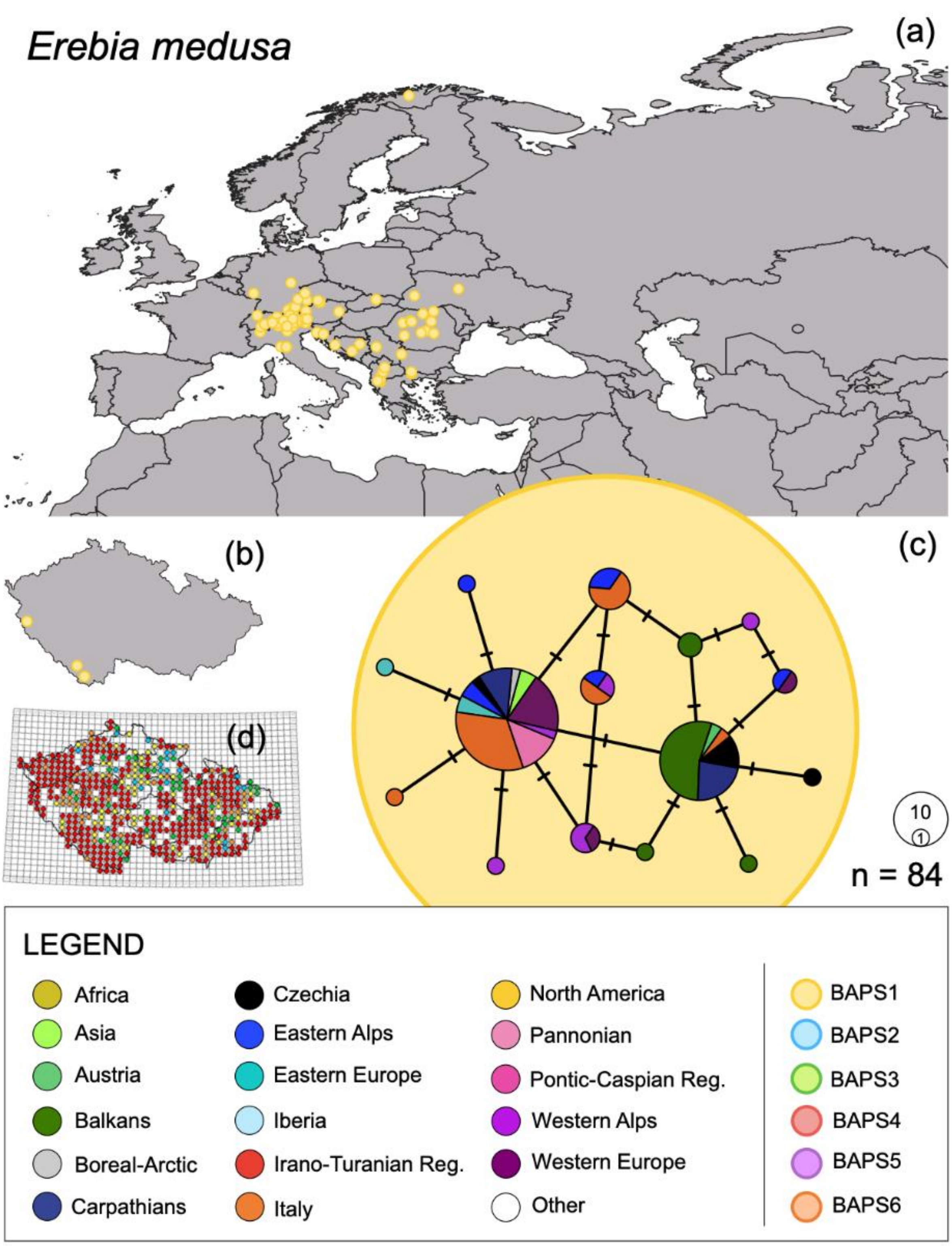


*Erebia medusa* ([Denis & Schiffermüller], 1775): **Habitat:** mesophilic 2. **Red List:** NT
**BAPS:** BAPS-low. **HAP:** HAP-Italy-Balkans (such distribution)
**Range:** Euro-Siberian. **Distribution CZ:** overgrown meadows with bushes, woodland meadows. Distributed across the country, but decreased with habitat alteration.

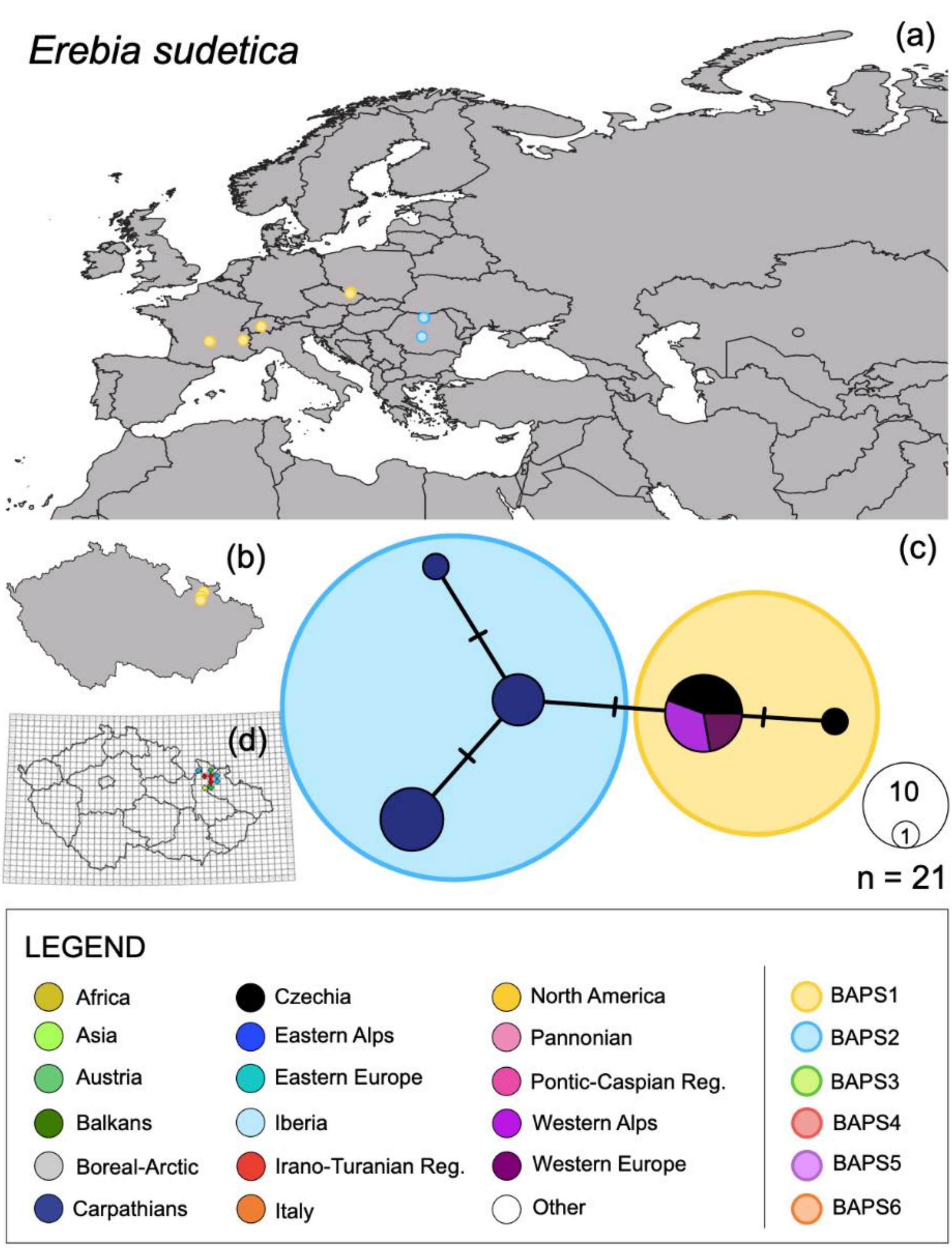


*Erebia sudetica* Staudinger, 1861: **Habitat:** alpine. **Red List:** VU

**BAPS:** BAPS-West. **HAP:** HAP-West

**Range:** European. **Distribution CZ:** mountain woodlands openings, subalpine habitats. It occurs only in Hrubý Jeseník Mts., where it has retreated from lower altitudes.

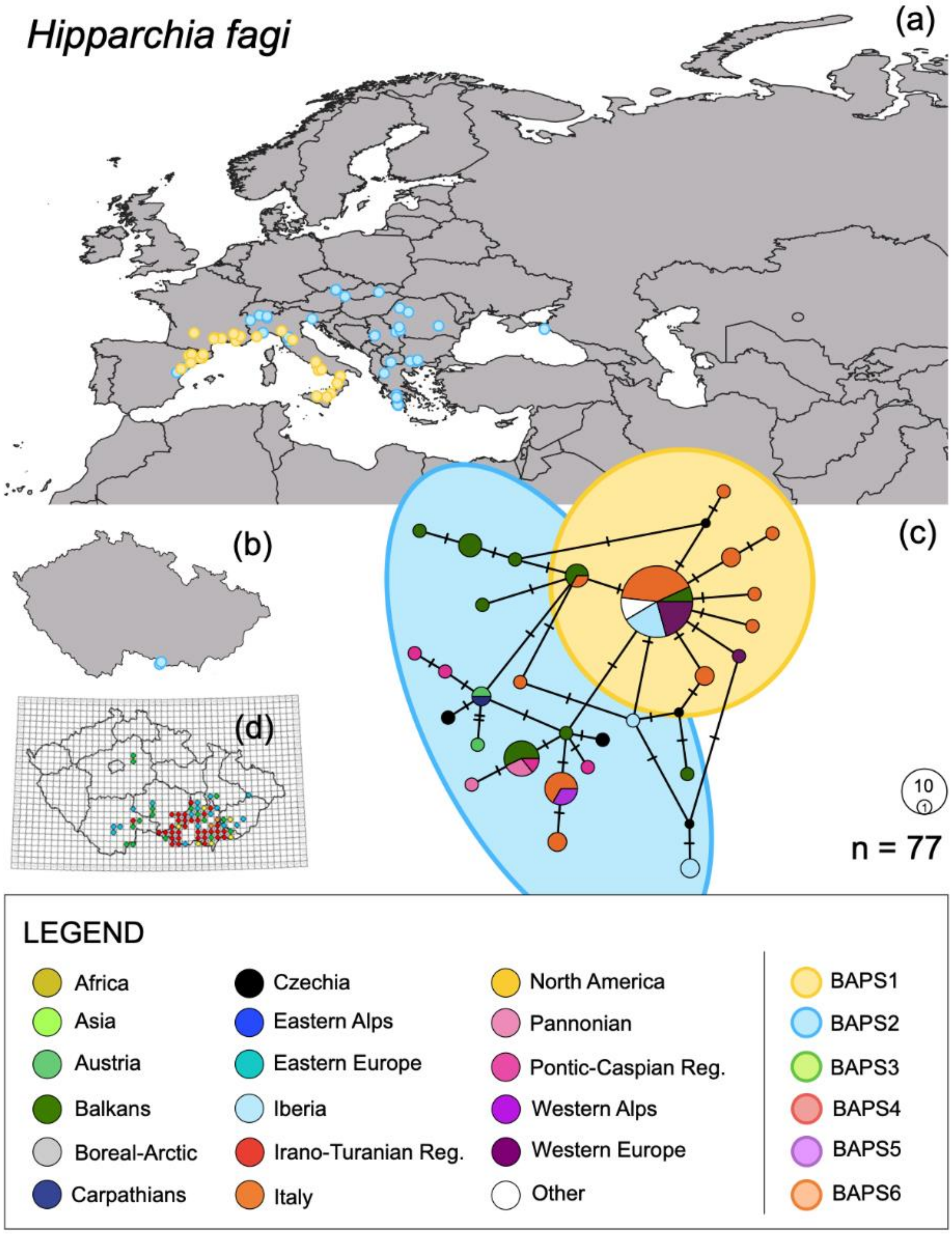


*Hipparchia fagi* (Scopoli, 1763): **Habitat:** xerothermophilic 2. **Red List:** VU

**BAPS:** BAPS-wide. **HAP:** HAP-East-Balkans

**Range:** Pontic-Mediterranean. **Distribution CZ:** dry open woodland margins, forest steppes, rocky places. Extinct from Bohemia, present in South Moravia.

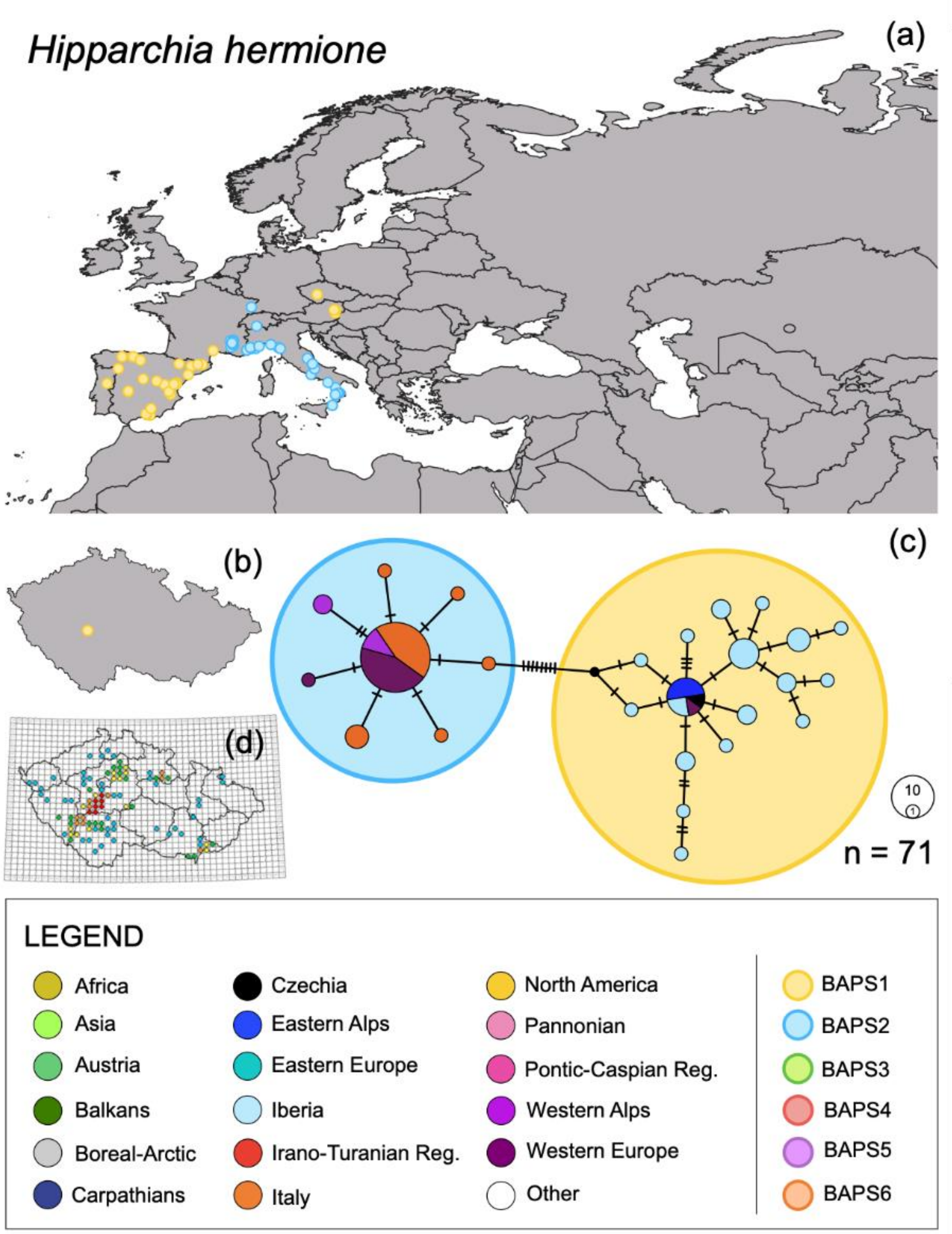


*Hipparchia hermione* (Linnaeus, 1764): **Habitat:** mesophilic 2 + xerothermophilic 2. **Red List:** CR

**BAPS:** BAPS-West. **HAP:** HAP-West

**Range:** European. **Distribution CZ:** dry open woodlands in warm areas, rock outcrops. Extinct from majority of its former distribution (loss of open woodlands), now surviving only in Vltava river canyon (Kysela, Konvička, Pokorný, & Sedláček, 2021).

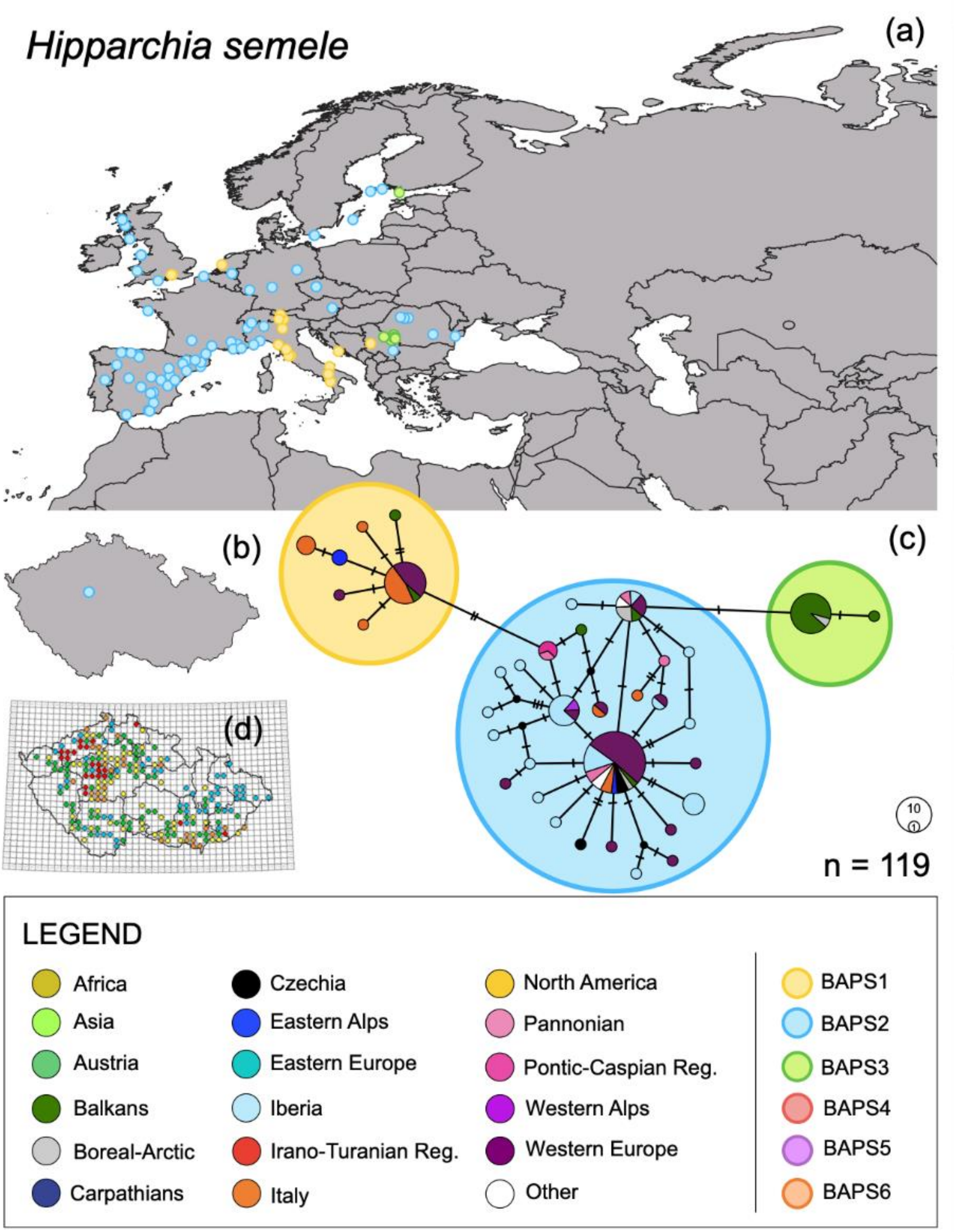


*Hipparchia semele* (Linnaeus, 1758): **Habitat:** xerothermophilic 2. **Red List:** CR

**BAPS:** BAPS-wide. **HAP:** HAP-wide-complex

**Range:** European. **Distribution CZ:** dry open woodlands in warm areas, rocky places. Severely declined from the entire area, extinct from Moravia, which is currently recolonizing from the south. Still present in a few localities in central parts of Bohemia. It uses post-industrial areas as an alternative habitat (Tropek, Cizek, Kadlec, & Klecka, 2017).

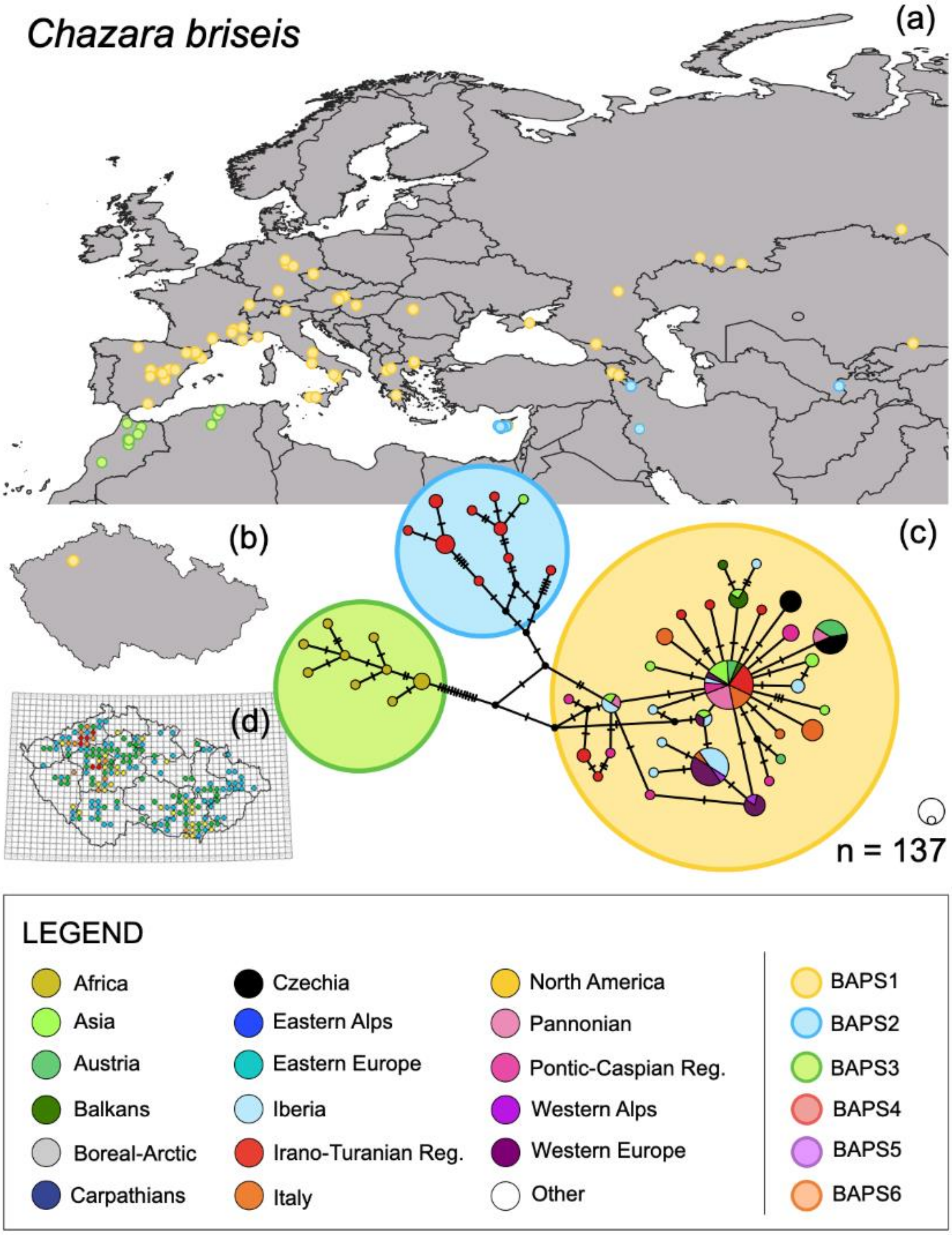


*Chazara briseis* (Linnaeus, 1764): **Habitat:** xerothermophilic 1. **Red List:** CR

**BAPS:** BAPS-wide. **HAP:** HAP-Italy-Balkans

**Range:** Pontic-Mediterranean. **Distribution CZ:** dry short grasslands on rocky substrates. Historically present on rocky sites across the country, but extinct from most of its range. In recent years, its range has contracted to a single site in Central Bohemian Uplands. However, an extensive conservation breeding combined with habitat management allowed several reintroductions, aiming to restore a metapopulation dynamics (John, Beneš, Číp, Andres, & Konvička, 2020; Sucháčková Bartoňová, Konvička, Marešová, Bláhová, et al., 2021).

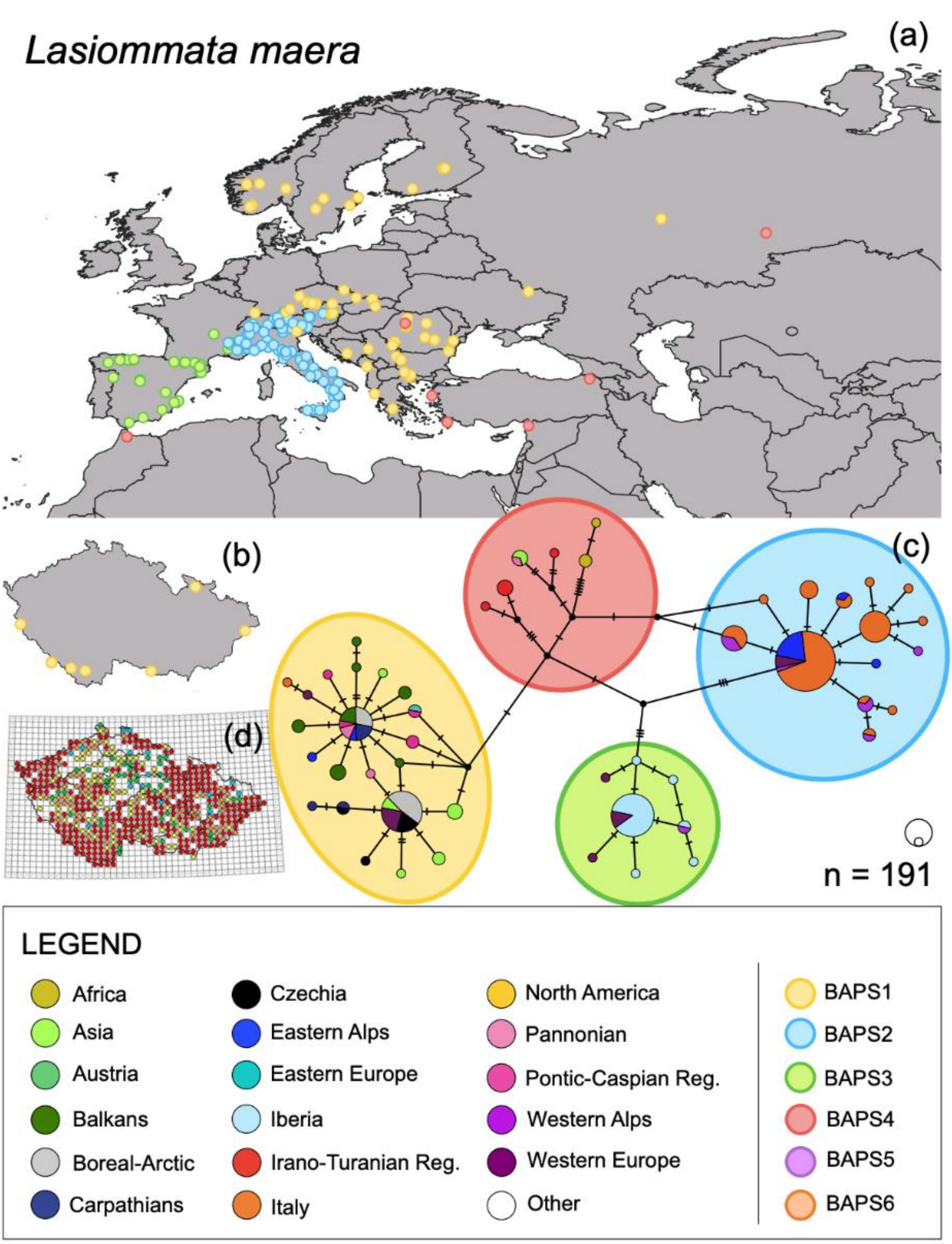


*Lasiommata maera* (Linnaeus, 1758): **Habitat:** mesophilic 2 + xerothermophilic 2. **Red List:** NT

**BAPS:** BAPS-East-Balkans. **HAP:** HAP-East-Balkans

**Range:** Western Palearctic. **Distribution CZ:** woodland margins and clearings, and also forest steppes and dry scrublands. Widespread, but absent from open farmlands.

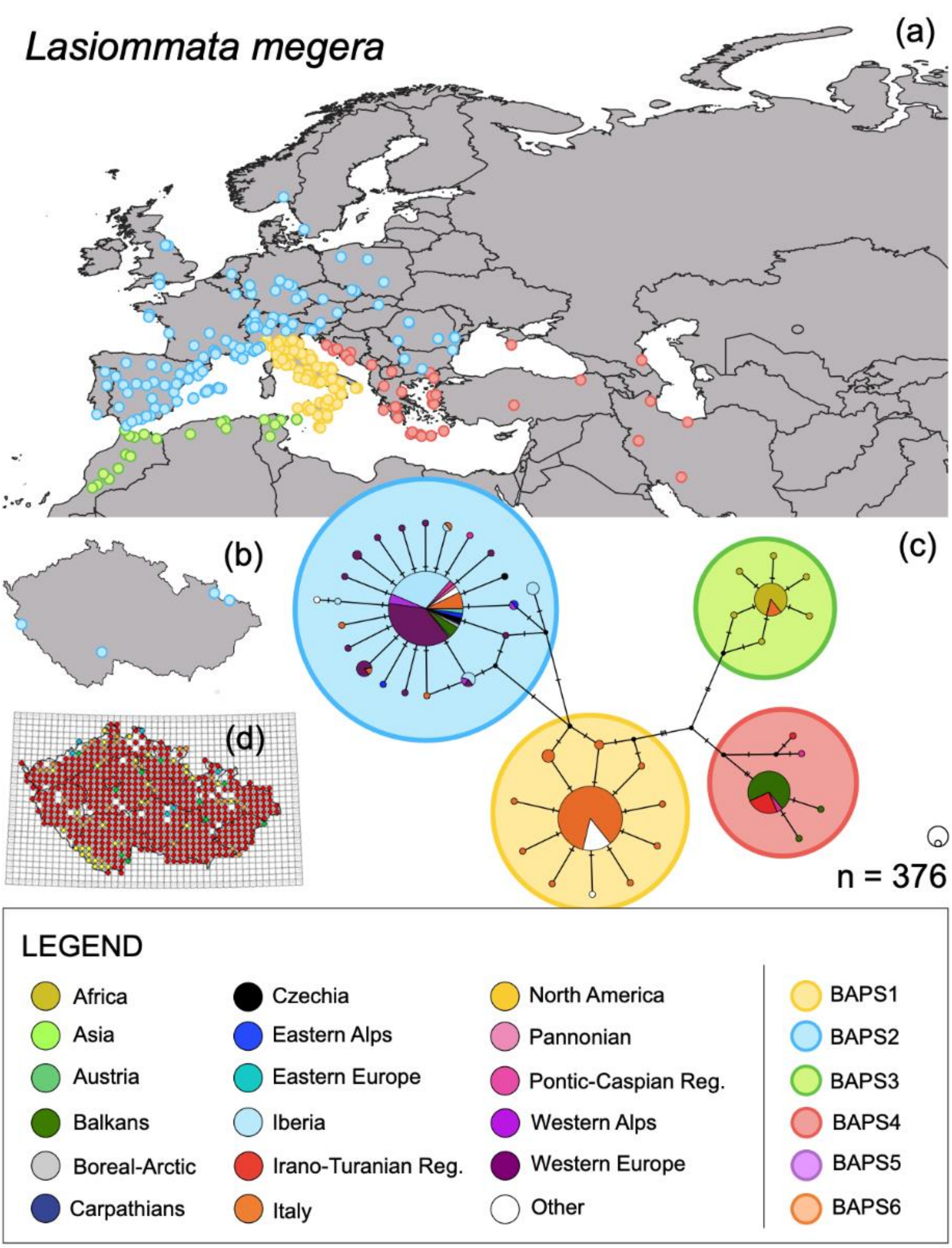


*Lasiommata megera* (Linnaeus, 1767): **Habitat:** ubiquitous. **Red List:** LC

**BAPS:** BAPS-West. **HAP:** HAP-West

**Range:** Western Palearctic. **Distribution CZ:** diverse habitats, it occurs close to garden walls, rocky places, field roads. Widespread, avoiding high altitudes.

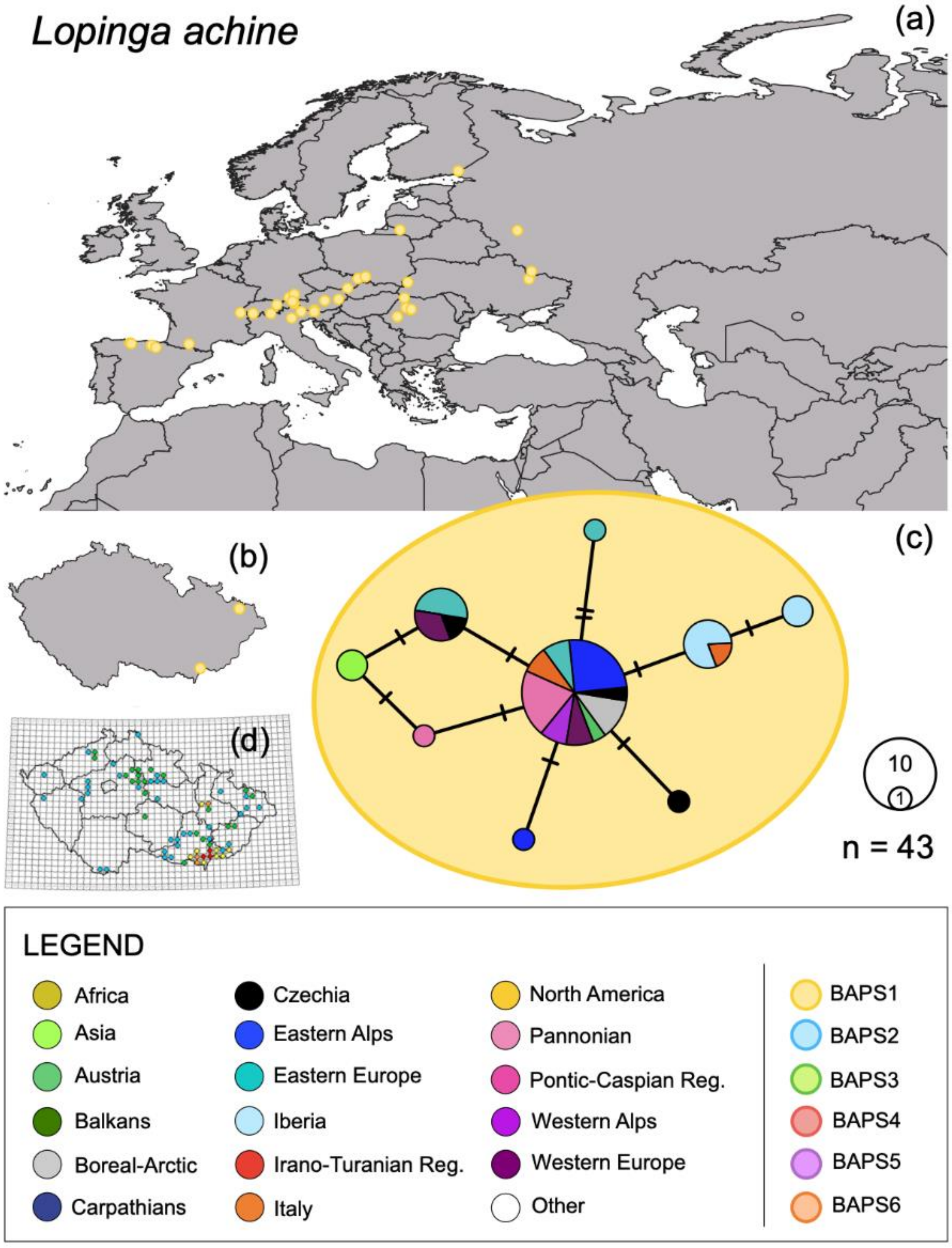


*Lopinga achine* (Scopoli, 1763): **Habitat:** mesophilic 3. **Red List:** CR

**BAPS:** BAPS-low. **HAP:** HAP-temperate

**Range:** Euro-Siberian. **Distribution CZ:** open woodlands. Extinct from majority of its range (loss of open woodlands), surviving at a single site southern Moravia, plus at a site in northeastern Moravia established by unauthorised reintroduction.

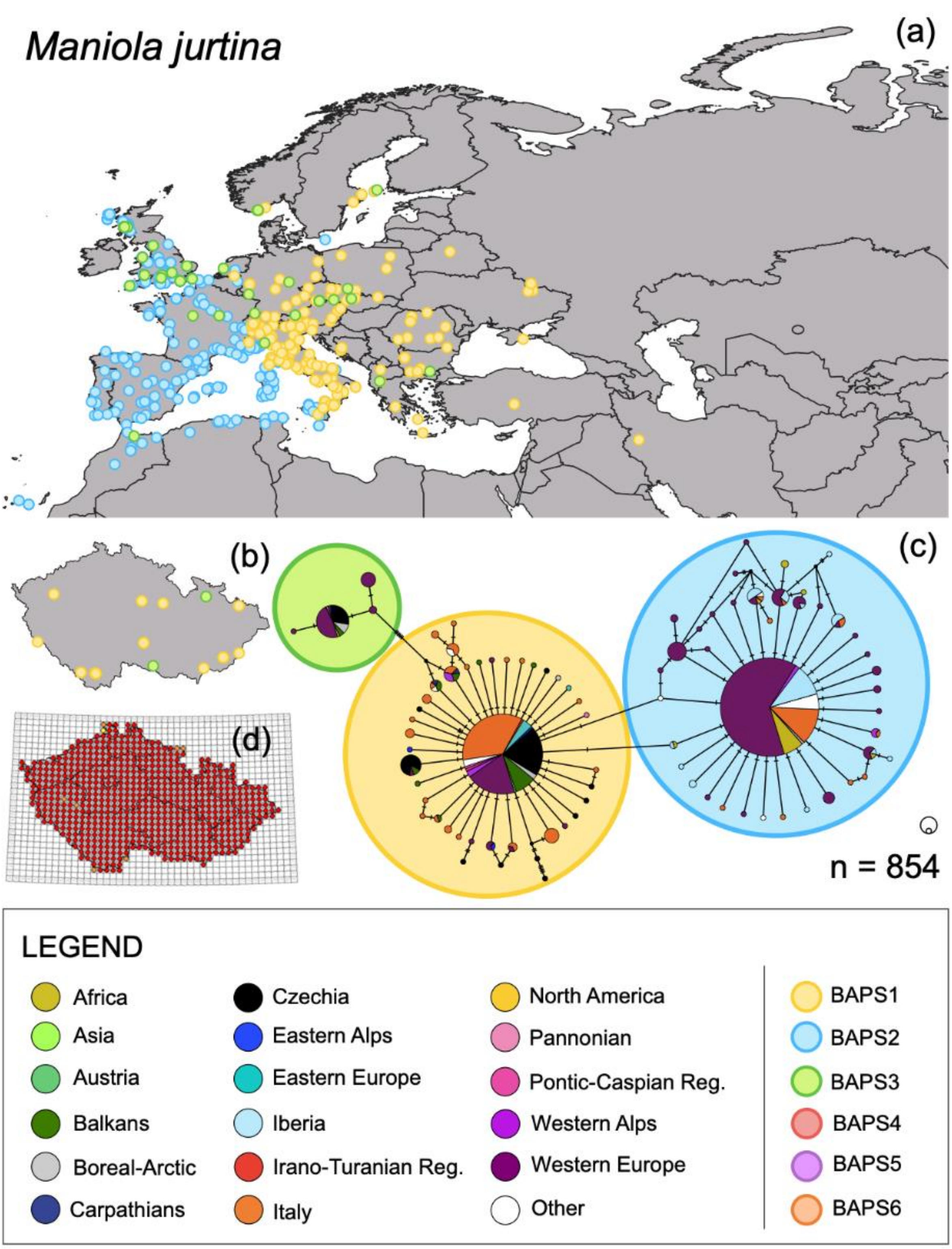


*Maniola jurtina* (Linnaeus, 1758): **Habitat:** ubiquitous. **Red List:** LC

**BAPS:** BAPS-mix-East. **HAP:** HAP-mix

**Range:** Western Palearctic. **Distribution CZ:** various meadow habitats. Widespread and common.

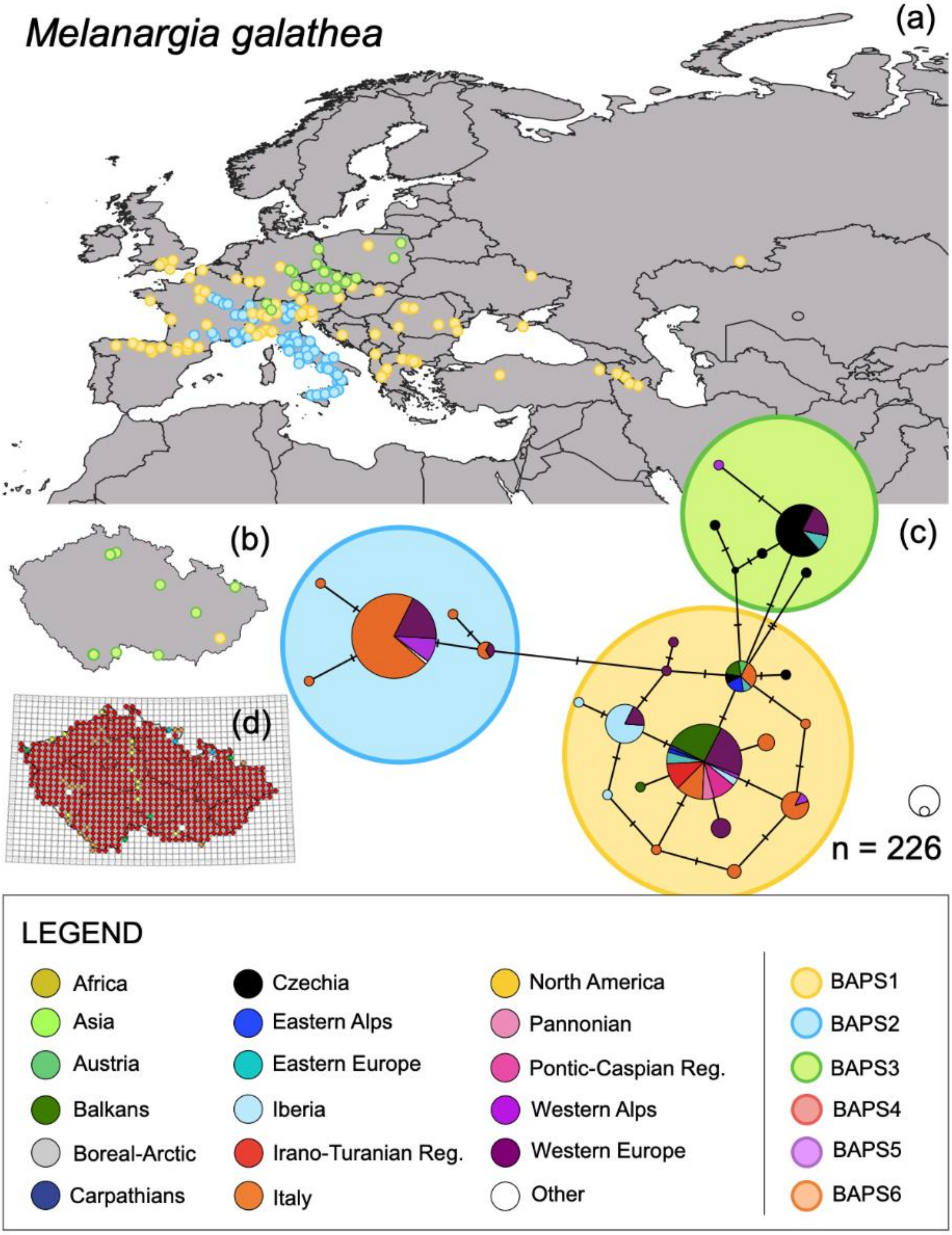


*Melanargia galathea* (Linnaeus, 1758): **Habitat:** mesophilic 1. **Red List:** LC

**BAPS:** BAPS-wide + BAPS-Central. **HAP:** HAP-Central

**Range:** Western Palearctic. **Distribution CZ:** various grassland habitats. Widespread and common.

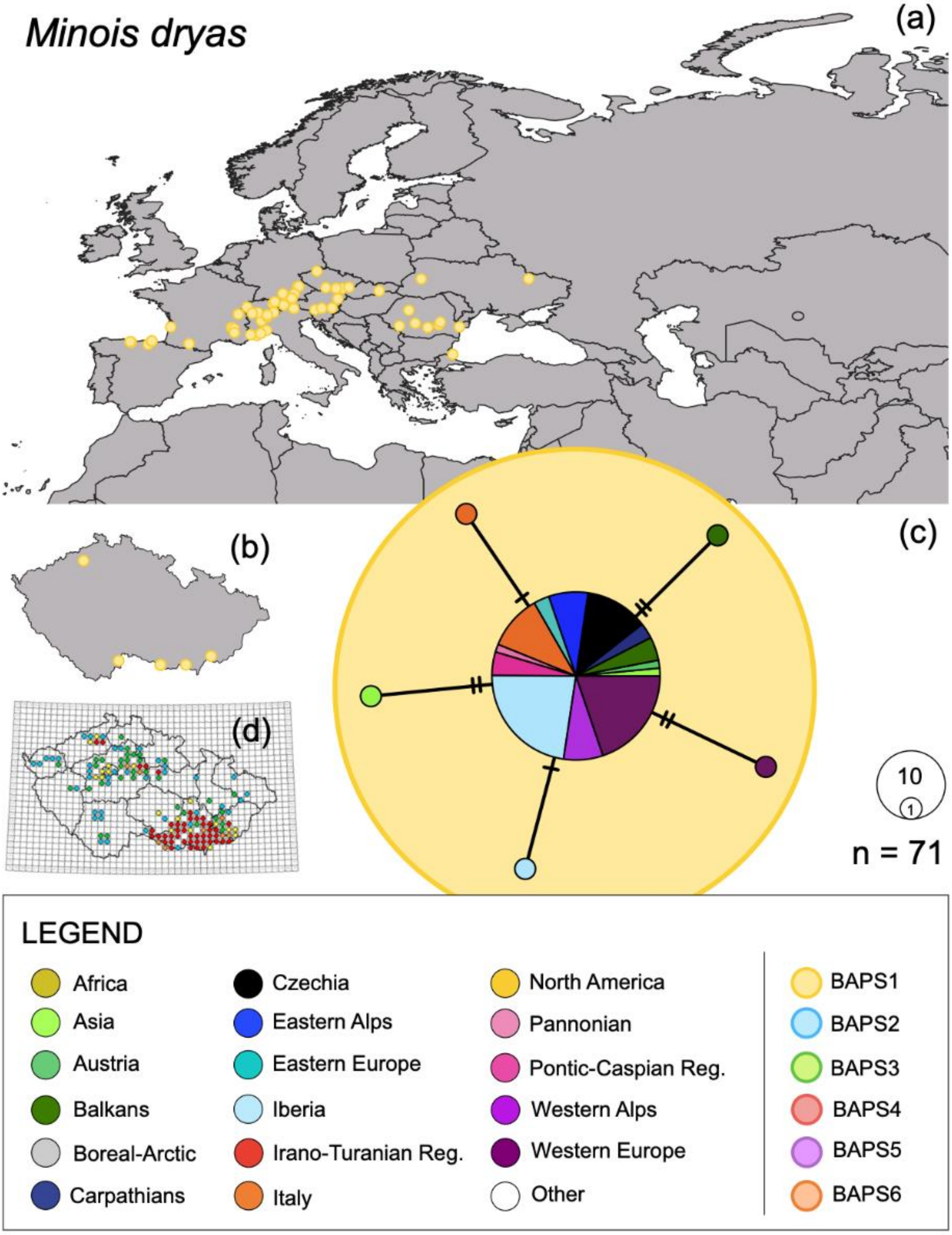


*Minois dryas* (Scopoli, 1763): **Habitat:** xerothermophilic 2 + hydrophilic. **Red List:** VU

**BAPS:** BAPS-low. **HAP:** HAP-low

**Range:** Euro-Siberian. **Distribution CZ:** warm dry scrublands and open woodlands, sometimes on wet bushy meadows. Decreased, locally common in south and central Moravia, in central and northern parts of Bohemia, and southern Bohemia.

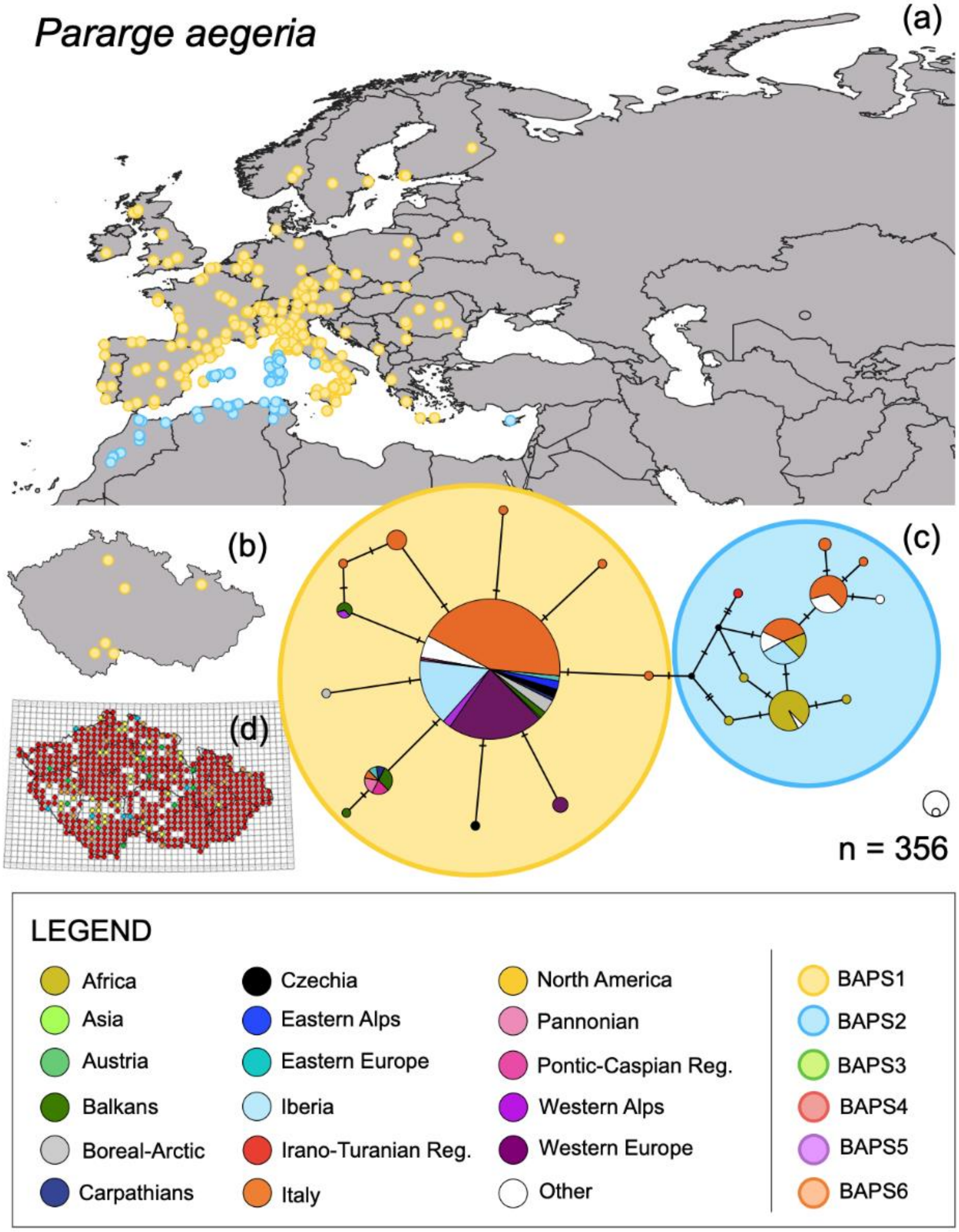


*Pararge aegeria* (Linnaeus, 1758): **Habitat:** mesophilic 3. **Red List:** LC

**BAPS:** BAPS-wide. **HAP:** HAP-wide-low

**Range:** Western Palearctic. **Distribution CZ:** all types of woodlands, including close-canopy forest plantations. Widespread and common.
